# Benchmarking principal component analysis for large-scale single-cell RNA-sequencing

**DOI:** 10.1101/642595

**Authors:** Koki Tsuyuzaki, Hiroyuki Sato, Kenta Sato, Itoshi Nikaido

## Abstract

Principal component analysis (PCA) is an essential method for analyzing single-cell RNA-seq (scRNA-seq) datasets, but large-scale scRNA-seq datasets require long computational times and a large memory capacity.

In this work, we review 21 fast and memory-efficient PCA implementations (10 algorithms) and evaluate their application using 4 real and 18 synthetic datasets. Our benchmarking showed that some PCA algorithms are faster, more memory efficient, and more accurate than others. In consideration of the differences in the computational environments of users and developers, we have also developed guidelines to assist with selection of appropriate PCA implementations.

## Background

The emergence of single-cell RNA sequencing (scRNA-seq) technologies [1], has enabled the examination of many types of cellular heterogeneity. For example, cellular subpopulations consisting of various tissues [2–6], rare cells and stem cell niches [7], continuous gene expression changes related to cell cycle progression [8], spatial coordinates [9–11], and differences in differentiation maturity [12, 13] have been captured by many scRNA-seq studies. As the measurement of cellular heterogeneity is highly dependent on the number of cells measured simultaneously, a wide variety of large-scale scRNA-seq technologies have been developed [14], including those using cell sorting devices [15–17], Fludigm C1 [18–21], droplet-based technologies (Drop-Seq [2–4], inDrop RNA-Seq [5, 6], the 10X Genomics Chromium system [22]), and single-cell combinatorial-indexing RNA-sequencing (sci-RNA-seq [23]). Such technologies have encouraged the establishment of several large-scale genomics consortiums, such as the Human Cell Atlas [24–26], Mouse Cell Atlas [27], and Tabula Muris [28]. These projects are analyzing a tremendous number of cells by scRNA-seq and tackling basic life science problems such as the number of cell types comprising an individual, cell-type-specific marker gene expression and gene functions, and molecular mechanisms of diseases at a single-cell resolution.

Nevertheless, the analysis of scRNA-seq datasets poses a potentially difficult problem; the cell type corresponding to each data point is unknown *a priori* [1, 29–35]. Accordingly, researchers perform unsupervised machine learning (UML) methods, such as dimensionality reduction and clustering, to reveal the cell type corresponding to each individual data point. In particular, principal component analysis (PCA [36–38]) is a commonly used UML algorithm applied across many situations.

Despite its wide use, there are several reasons why it is unclear how PCA should be conducted for large-scale scRNA-seq. First, because the widely used PCA algorithms and implementations load all elements of a data matrix into memory space, for large-scale datasets such as the 1.3 million cells measured by 10X Genomics Chromium [39] or the 2 million cells measured by sci-RNA-seq [23], the calculation is difficult unless the memory size of the user’s machine is very large. Furthermore, the same data analysis workflow is performed repeatedly, with deletions or additions to the data or parameter changes for the workflow, and under such trial-and-error cycles, PCA can become a bottleneck for the workflow. Therefore, some fast and memory-efficient PCA algorithms are required.

Second, there are indeed some PCA algorithms that are fast and memory efficient, but their practicality for use with large-scale scRNA-seq datasets is not fully understood. Generally, there are trade-offs between the acceleration of algorithms by some approximation methods and the accuracy of biological data analysis. Fast PCA algorithms might overlook some important differential gene expression patterns. In the case of large-scale scRNA-seq studies aiming to find novel cell types, this property may cause a loss of clustering accuracy and not be acceptable.

Finally, actual computational time and memory efficiency are highly dependent on the specific implementation, including the programming language, the method for loading input files, and the data format. However, there is no benchmarking to evaluate these properties. Such information is directly related to the practicality of the software and is useful as a guideline for users and developers.

For the above reasons, in this research, we examine the practicality of fast and memory-efficient PCA algorithms for use with large-scale scRNA-seq datasets. This work provides four key contributions. First, we review the existing PCA algorithms and their implementations (Figure 1). Second, we present a benchmark test with selected PCA algorithms and implementations. To our knowledge, this is the first comprehensive benchmarking of PCA algorithms and implementations with large-scale scRNA-seq datasets. Third, we provide some original implementations of some PCA algorithms and utility functions for quality control (QC), filtering, and feature selection. All commands are implemented in a fast and memory-efficient Julia package. Finally, we propose guidelines for end-users and software developers.

**Figure 1.**
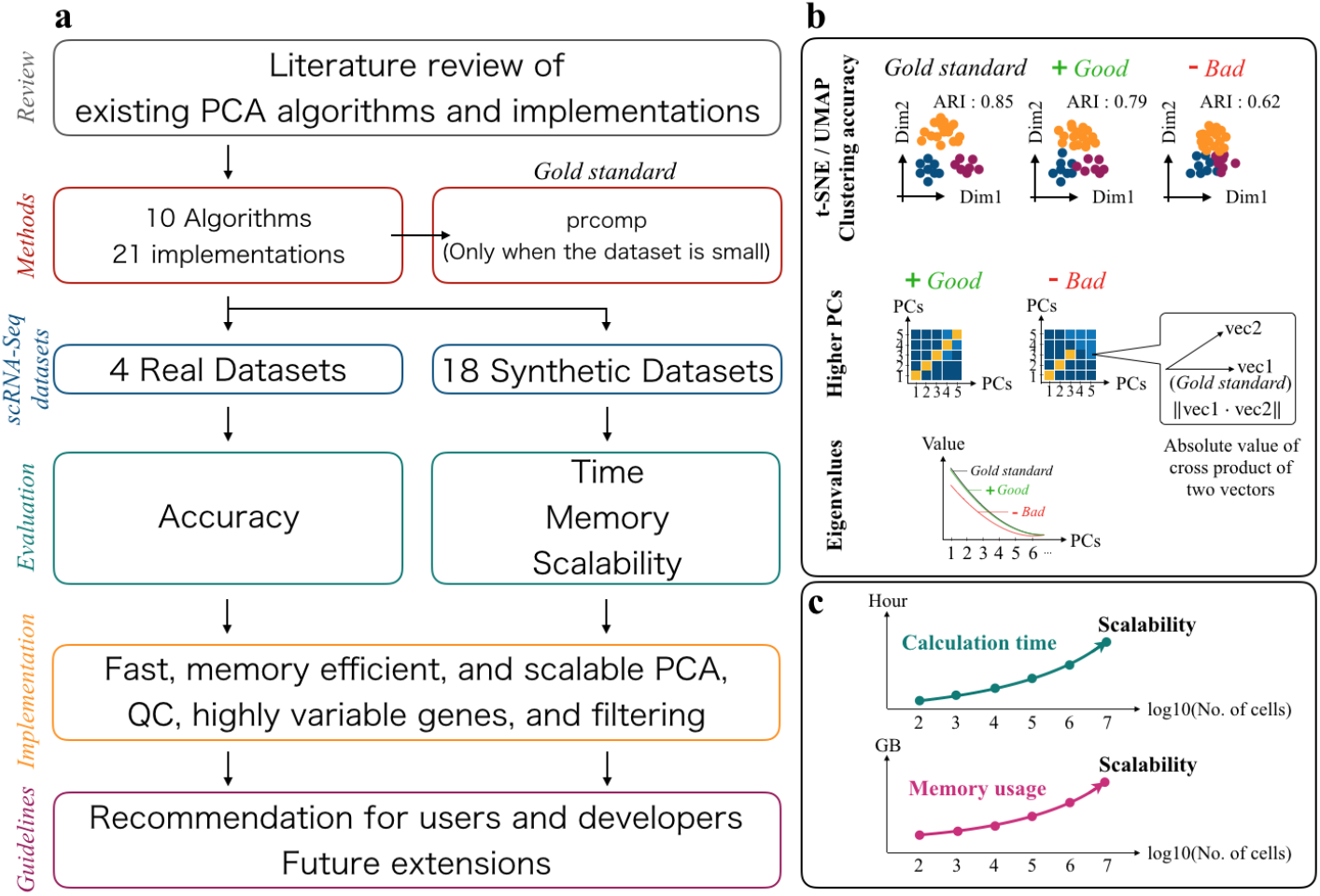
Overview of benchmarking in this work. (a) Schematic overview of this work. (b) Evaluation metrics of the benchmarking with real-world datasets. (c) Evaluation metrics of the benchmarking with synthetic datasets.

## Results

### Review of PCA algorithms and implementations

PCA is widely used for data visualization [39–41], data QC [42], feature selection [13, 43–49], de-noising [50, 51], imputation [52–54], confirmation and removal of batch effects [55–57], confirmation and estimation of cell-cycle effects [58], rare cell type detection [59, 60], cell type and cell state similarity search [61], pseudotime inference [13, 62–66], and spatial reconstruction [9].

Additionally, principal component (PC) scores are also used as the input of other non-linear dimensionality reduction [67–73] and clustering methods [74–77] in order to preserve the global structure, avoid the “curse of dimensionality” [78–81], and save memory space. A wide variety of scRNA-seq data analysis tools actually include PCA as an internal function or utilize PC scores as input for down-stream analyses [22, 82–89].

We reviewed the existing PCA algorithms and implementations and classified the algorithms into six categories, namely, similarity transformation-based (SimT), downsampling-based (DS), singular value decomposition (SVD) update-based (SU), Krylov subspace-based (Krylov), gradient descent-based (GD), and random projection-based (Rand) (Additional file 1 [22, 42–44, 49–52, 55–61, 63, 65, 69, 74–77, 82, 85, 89–113]). We have listed 21 PCA implementations (comprising 10 algorithms) that are freely available and easy to download, install, and use for analyses. The correspondence of the reviewed PCA implementations and scRNA-seq studies are summarized in Table 1.

**Table 1.**
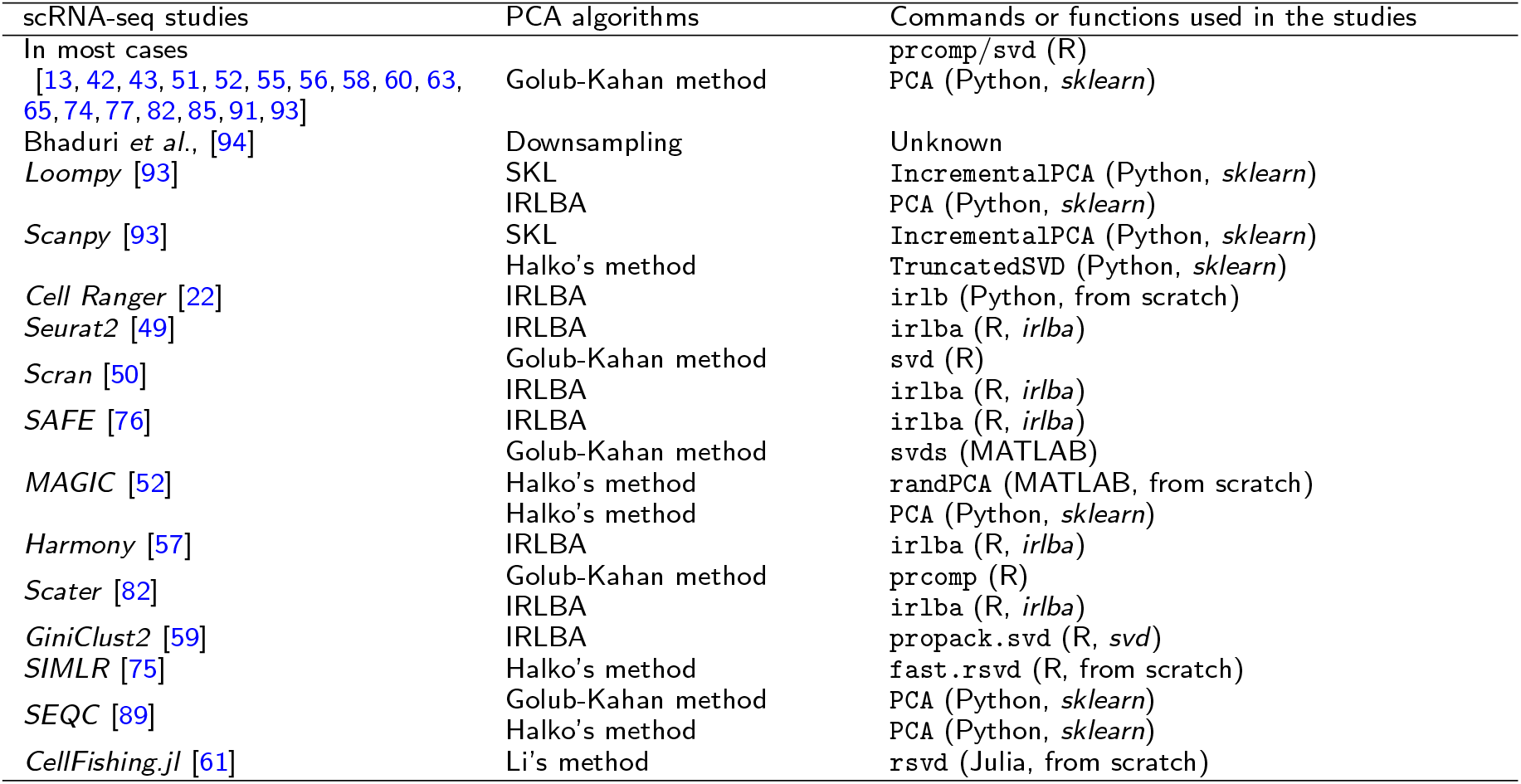
Use cases of PCA implementations in scRNA-seq studies.

To extend the scope of the algorithms used in the benchmarking, we originally implemented some PCA algorithms in an out-of-core manner (Additional file 1). The pseudo-code and source code of all the algorithms benchmarked in this study are summarized in Additional file 2 and Additional file 3, respectively.

### Benchmarking of PCA algorithms and implementations

Next, we performed the benchmarking tests of the PCA algorithms and implementations. The results of the benchmarking are summarized in Figure 2 [69, 90, 92, 94–99, 107–109, 114, 115].

**Figure 2.**
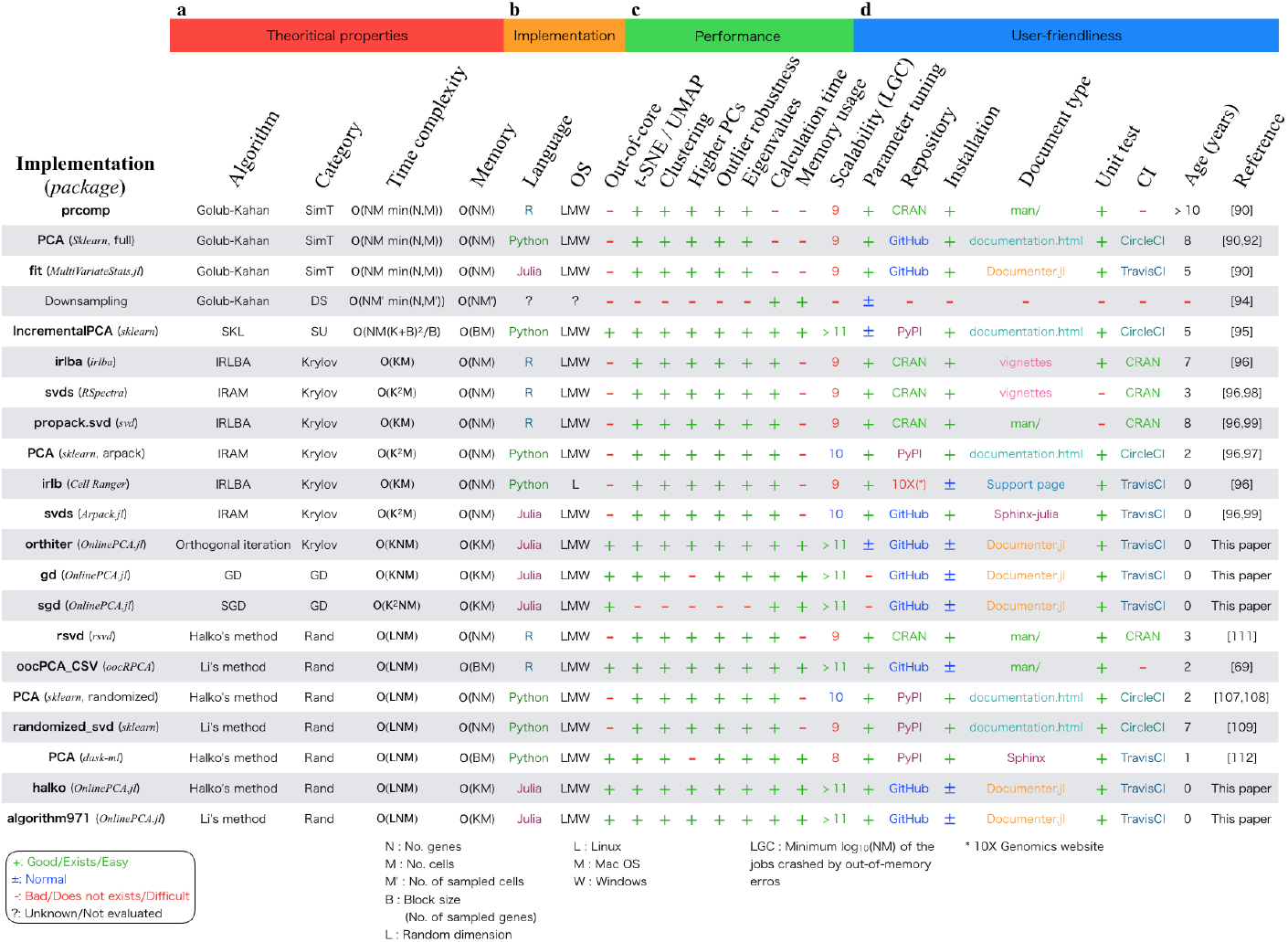
Summary of results. (a) Theoretical properties summarized by our literature review. (b) Properties related to each implementation. (c) Performance evaluated by benchmarking with real-world and synthetic datasets. (d) User-friendliness evaluated by some metrics.

#### Real-world datasets

In consideration of the trade-offs among the large number of methods evaluated with our limited time, computational resources, and manpower, we carefully selected real-world datasets for the benchmarking. The latest scRNA-seq methods are divided into two categories, namely, full-length scRNA-seq methods and high-throughput scRNA-seq methods with specific cell dissociation and cellular/molecular barcoding technologies such as droplet-based and split-and-pool experiments [34, 35]. Because the number of cells measured by scRNA-seq has been increased by the latter technology, we selected the following four datasets generated by such technologies: human peripheral blood mononuclear cells (PBMCs), human pancreatic cells (Pancreas), mouse brain and spinal cord (BrainSpinalCord), and mouse cells from the cortex, hippocampus, and ventricular zone (Brain) (Table 2). These datasets have been used in many previous scRNA-seq studies [61, 76, 94, 116–122].

**Table 2.**
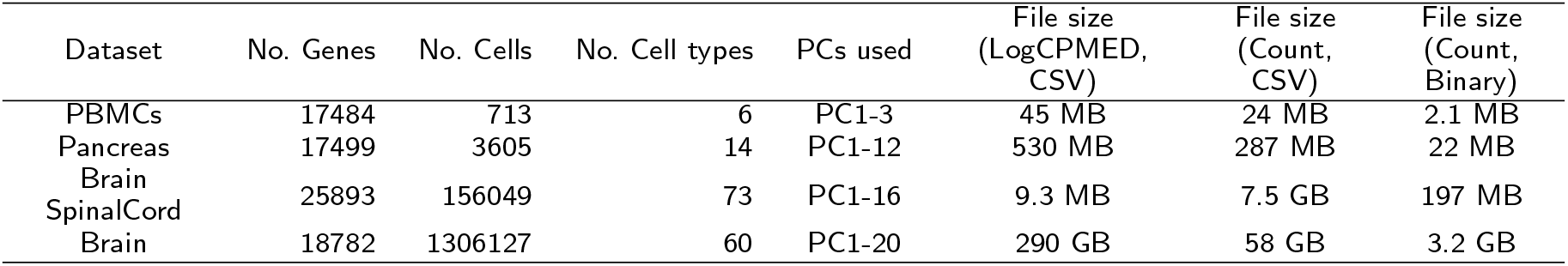
Real-world datasets for benchmarking

#### The accuracy of PCA algorithms

Here, we evaluate the accuracy of the various PCA algorithms by using the four real-world datasets. For the analyses of the PBMCs and Pancreas datasets, we set the result of prcomp as the gold standard, which is a wrapper function for performing SVD with LAPACK subroutines (Additional file 1). The other implementations are compared with this result (Figures 1b and 2). For the BrainSpinalCord and Brain datasets analyses, full-rank SVD by LAPACK is computationally difficult. According to the benchmarking guidelines developed by Mark D. Robinson’s group [123], comparing the methods against each other is recommended when the ground truth cannot be defined. Therefore, we just compared the results of the methods against each other using several different criteria, such as the magnitude of the eigenvalues and the clustering accuracy.

First, we performed t-stochastic neighbor embedding (t-SNE [67,68]) and uniform manifold approximation and projection (UMAP [71,72]) for the results of each PCA algorithm and compared the clarity of the cluster structures detected by the original studies (Figures 1b, 3, Additional file 4, and Additional file 5). For the Brain-SpinalCord and Brain datasets, only downsampling, `IncrementalPCA` (*sklearn*), `orthiter/gd/sgd/halko/algorithm971` (*OnlinePCA.jl*), and `oocPCA CSV` (*oocR-PCA*) could be performed, while the other implementations were terminated by out-of-memory errors on 96 and 128 GB RAM machines. For the PBMCS and Pancreas datasets, compared with the gold standard cluster structures, the structures detected by downsampling were unclear, and some distinct clusters determined by the original studies were incorrectly combined into single clusters. In the realistic situation when the cellular labels were unavailable *a priori*, the labels were exploratorily estimated by confirming differentially expressed genes, known markergenes, or related gene functions of clusters. In such a situation, downsampling may overlook subgroups hiding in a cluster.

**Figure 3.**
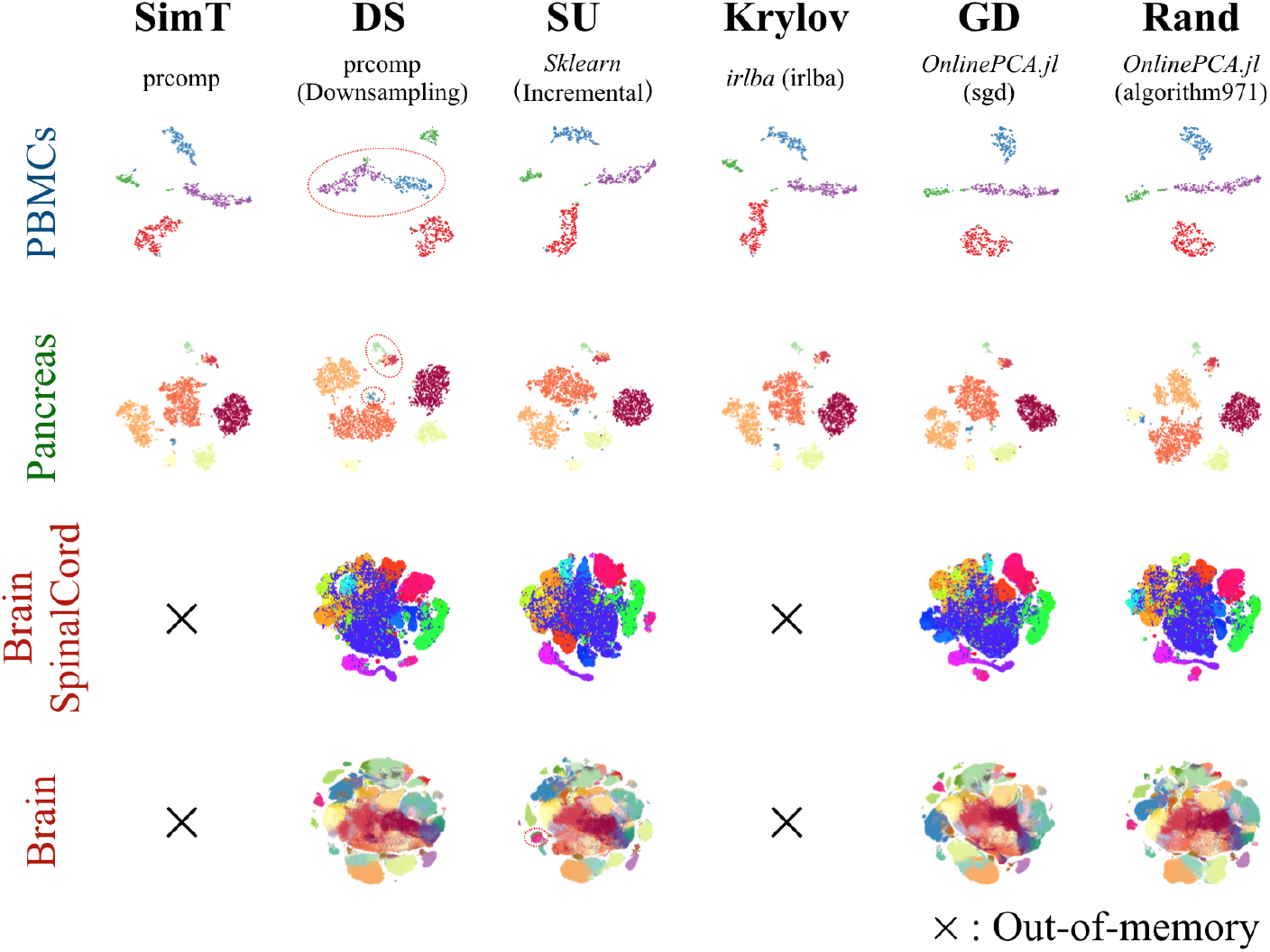
The comparison of t-stochastic neighbor embedding (t-SNE) plots. Comparison of multiple principal component analysis (PCA) implementations performed with empirical datasets: PBMCs (10^2^ cells), Pancreas (10^3^ cells), BrainSpinalCord (10^5^ cells), and Brain datasets (10^6^ cells). t-SNE was performed with the result of each PCA implementation.

We also performed four clustering algorithms on all the results of the PCA implementations and calculated the adjusted Rand index (ARI [124]) to evaluate clustering accuracy (Additional file 6). Here, we only show the result of Louvain clustering [125] (Figures 1b and 4). The ARI values show that the results of downsampling and sgd (*OnlinePCA.jl*) were worse compared with the gold standard or other implementations.

**Figure 4.**
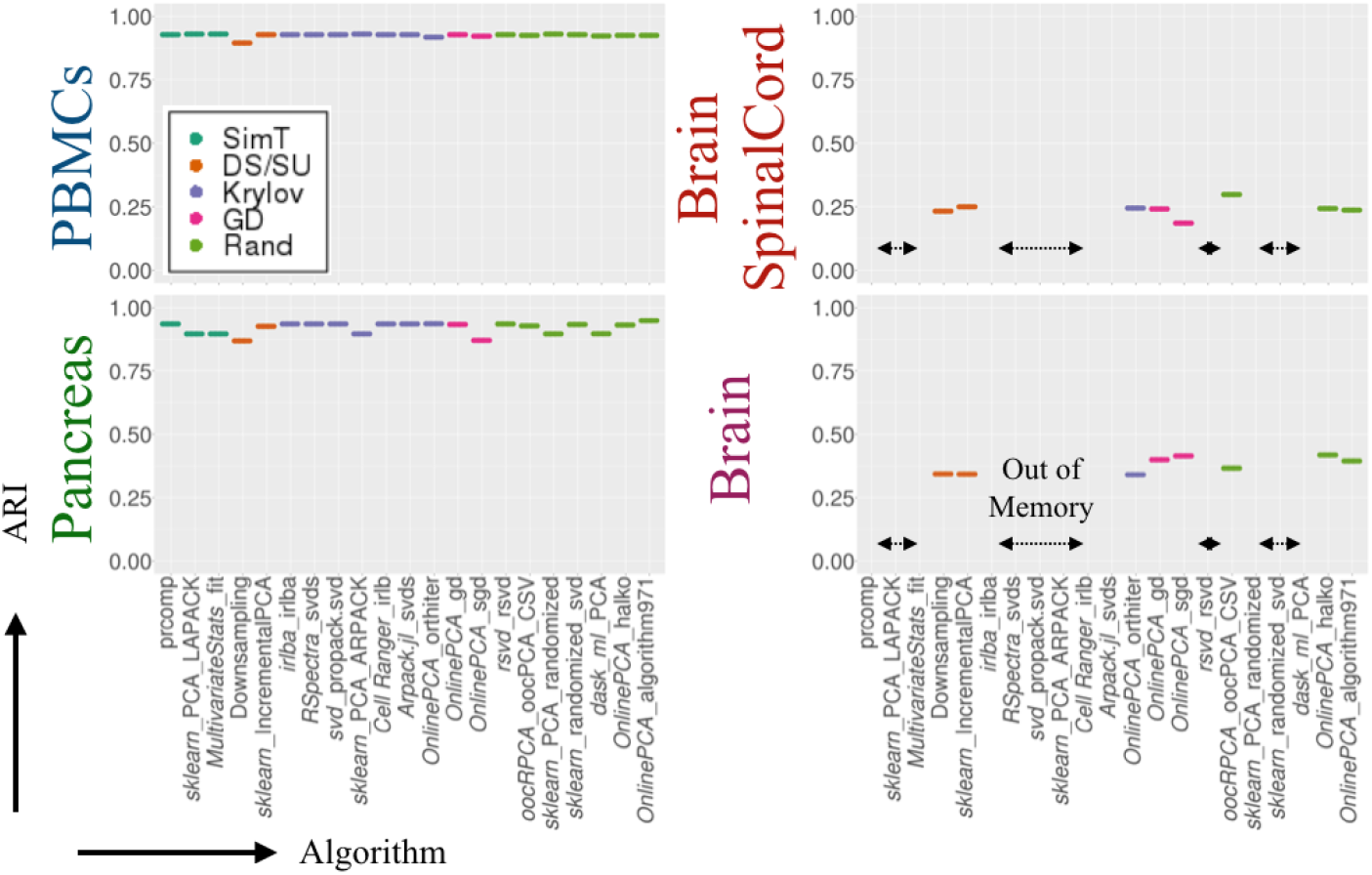
Clustering accuracy comparison. Clustering accuracy was evaluated by the adjusted Rand index (ARI) of the result of Louvain clustering. Multiple principal component analysis (PCA) implementations were performed for PBMCs (10^2^ cells), Pancreas (10^3^ cells), BrainSpinalCord (10^5^ cells), and Brain datasets (10^6^ cells); Louvain clustering was performed for the PCA results. For each PCA result, Louvain clustering calculations were performed ten times. The cluster labels are the same as those of the respective original papers.

Next, we performed an all-to-all comparison between PCs from the gold standard and the other PCA implementations (Figures 1b, 5a, and Additional file 7). Because the PCs are unit vectors, when two PCs are directed in the same or opposite direction, their cross product becomes 1 or *−*1, respectively. Both the same and opposite direction vectors are mathematically identical in PCA optimization, and different PCA implementations may yield PCs with different signs. Accordingly, we calculated the absolute value of the cross product ranging from 0 to 1 for the all-to-all comparison and evaluated whether higher PCs, which correspond to lower eigenvalues, are accurately calculated. The Figure 5a and Additional file 7 show that the higher PCs based on downsampling, `orthiter/gd/sgd` (*OnlinePCA.jl*), and `PCA` (*dask-ml* [115]) become inaccurate as the dimensionality of a PC increases. The higher PCs of these implementations also appear noisy and unclear in pair plots of PCs between each implementation and seem uninformative (Additional file 8, Additional file 9, Additional file 10, and Additional file 11). In particular, the higher PCs calculated by downsampling and `sgd` (*OnlinePCA.jl*) are sometimes influenced by the existence of outlier cells (Additional file 8 and Additional file 9). When performing some clustering methods, such as *k*-means and Gaussian mixture model (GMM [126]) methods, such outlier cells are also detected as singleton clusters having only a single cell as their cluster member (Additional file 12). Contrary to these results, all the implementations of IRLBA and IRAM, as well as the randomized SVD approaches except for `PCA` (*dask-ml*), are surprisingly accurate regardless of the language in which they are written or their developers. Although `PCA` (*dask-ml*) is based on Halko’s method and is nearly identical to the other implementations of Halko’s method, this function uses the direct tall-and-skinny QR algorithm [127] (https://github.com/dask/dask/blob/a7bf545580c5cd4180373b5a2774276c2ccbb573/dask/array/linalg.py#L52), and this characteristic might be related to the inaccuracy of the implementations. Because there is no gold standard in the case of the BrainSpinalCord and Brain datasets, we compared the eigenvectors of the PCA implementations in all possible combinations (Additional file 13) and found that the higher PCs of downsampling and `sgd` differed from those of the other PCA implementations.

**Figure 5.**
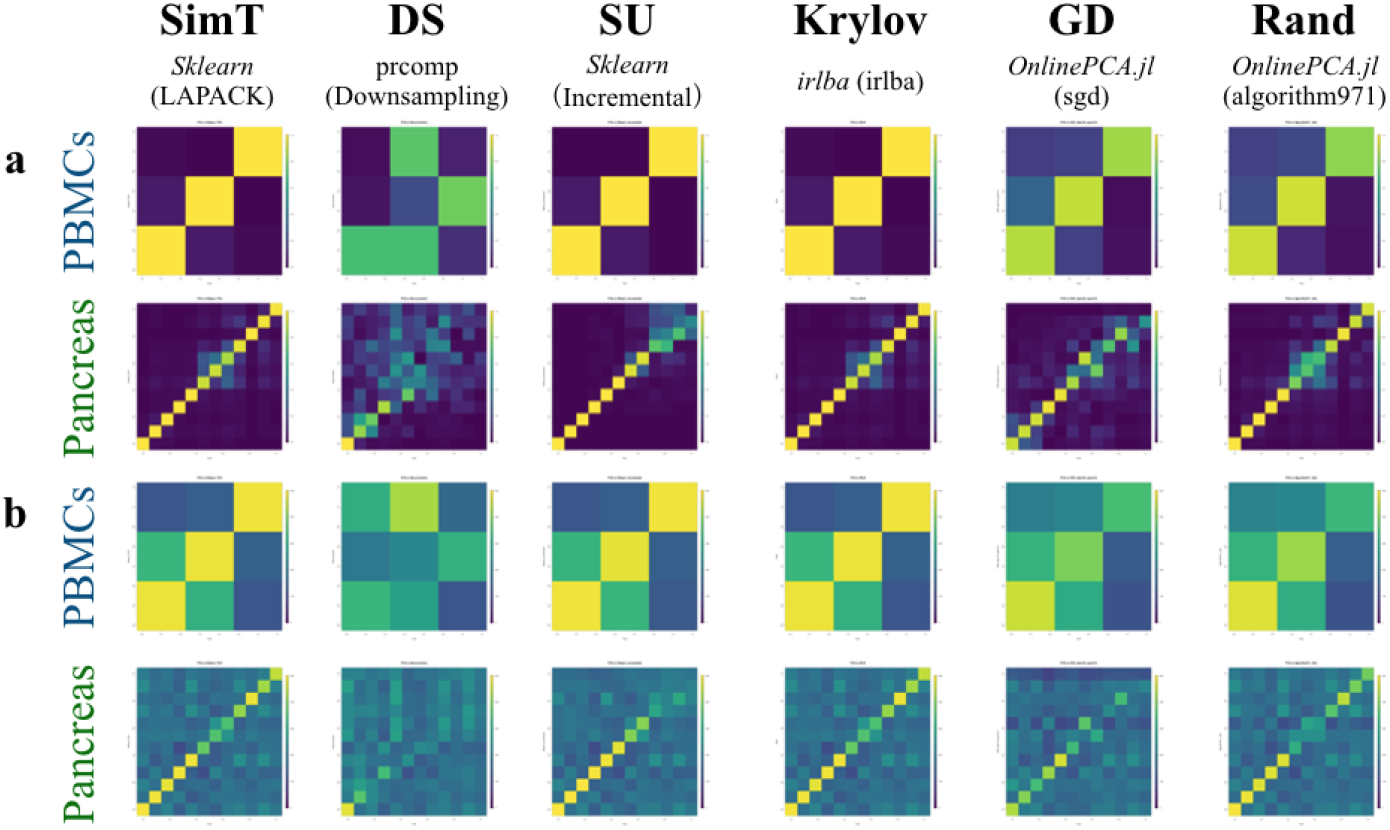
Comparison of all combinations of eigenvectors. Absolute values of the cross products of all combinations between the eigenvectors of the gold standard methods and those of the other principal component analysis (PCA) implementations were calculated. The closer the value is to 1, the closer the two corresponding eigenvectors are to each other. If two PCA results are equal without considering differences in sign, the matrix in this figure becomes an identity matrix.

Because gene-wise eigenvectors (i.e., loading vectors) are also retrieved from the data matrix and cell-wise eigenvectors (i.e., PCs), we also compared the loading vectors (Figure 5b and Additional file 14). We extracted the top 500 genes in terms of the largest absolute values of loading vectors and calculated the number of genes in common between the two loading vectors. As is the case with the eigenvectors, even for loading vectors, downsampling, `orthiter/gd/sgd` (*OnlinePCA.jl*), and `PCA` (*dask-ml* [115]) become inaccurate as the dimensionality of the PC increases. Because the genes with large absolute values for loading vectors are used as feature values in some studies [43–48], inaccurate PCA implementations may lower the accuracy of such an approach.

The distributions of the eigenvalues of downsampling, `IncrementalPCA` (*sklearn*), and `sgd` (*OnlinePCA.jl*) also differ from those of the other implementations (Figure 6).

**Figure 6.**
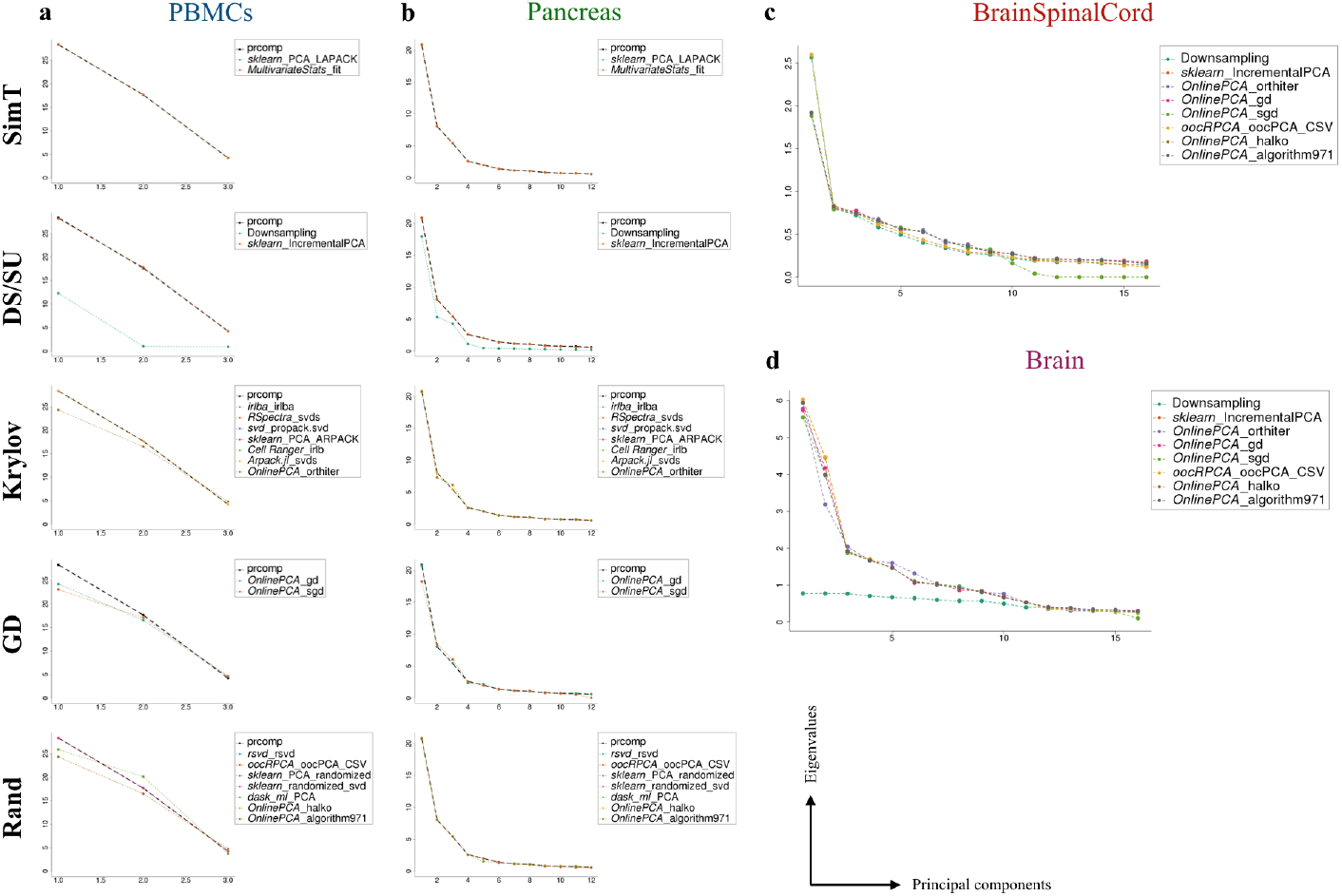
Comparison of eigenvalues. Distribution of eigenvalues of all the principal component analysis (PCA) implementations for each real dataset.

#### Calculation time, memory usage, and scalability

We compared the computational time and memory usage of all the PCA implementations (Figure 7). For the BrainSpinalCord dataset, downsampling itself was faster than most of the PCA implementations, but other preprocessing steps, such as matrix transposition and multiplication of the transposed data matrix and loading vectors to calculate PCs, were slow and had high memory space requirements (Additional file 3). For the Brain dataset, downsampling became slower than most of the PCA implementations, and such a tendency is noticeable as the size of the data matrix increases, because downsampling is based on the full-rank SVD in LAPACK.

**Figure 7.**
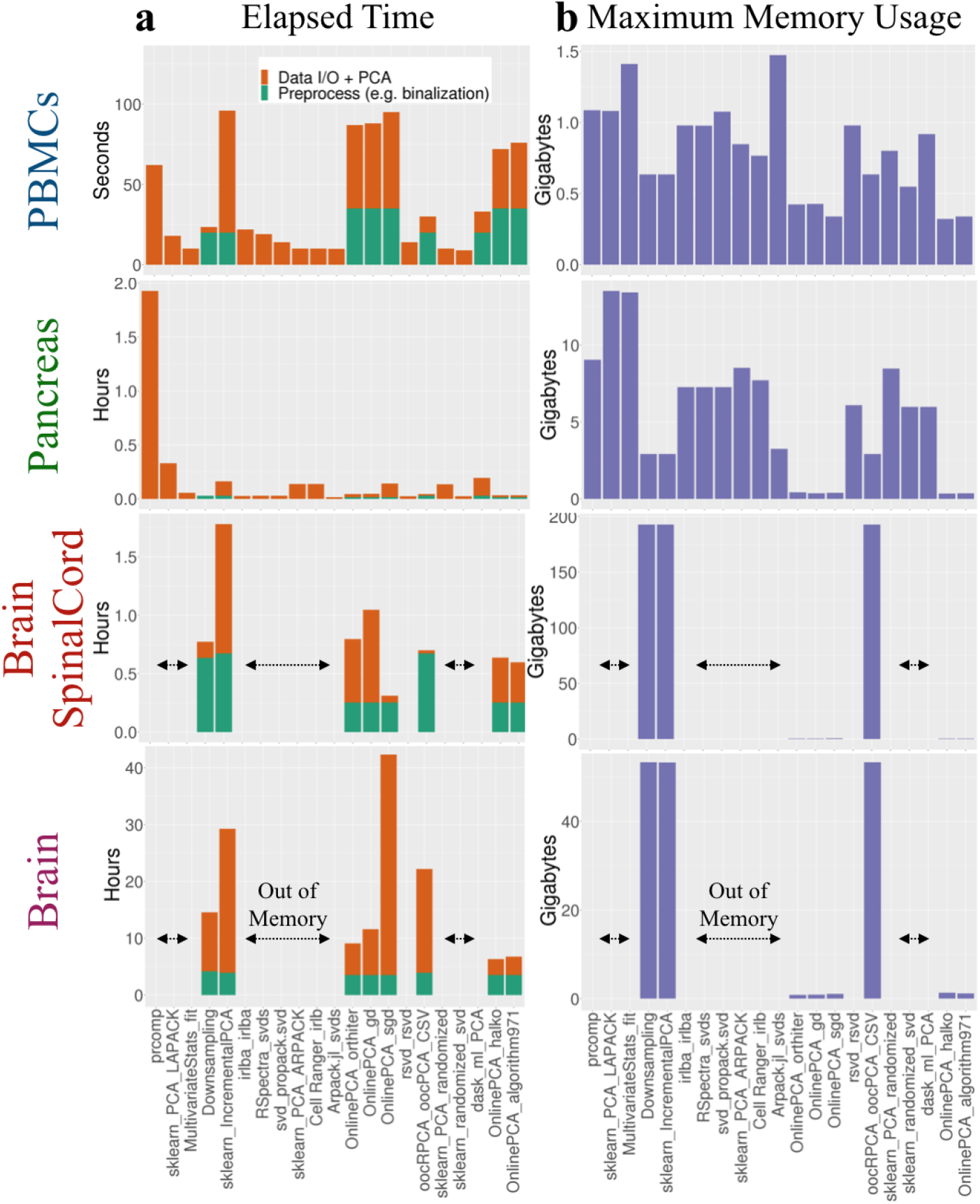
Comparison of the elapsed time and maximum memory usage for empirical datasets. (a) Elapsed time and (b) memory usage of all principal component analysis (PCA) implementations calculated for each empirical dataset. We used our in-house Julia script to preprocess the Brain dataset.

We also found that the calculation time of `PCA` (*dask-ml*) was not as fast in spite of its out-of-core implementation; for the BrainSpinalCord and Brain datasets, this implementation could not finish the calculation within three days in our computational environment. The other out-of-core PCA implementations, such as `IncrementalPCA` (*sklearn*), `orthiter/gd/sgd/halko/algorithm971` (*OnlinePCA.jl*), and `oocPCA CSV` (*oocRPCA*), were able to finish those calculations.

We also systemically estimated the calculation time, memory usage, and scalability of all the PCA implementations using 18 synthetic datasets consisting of {10^2^, 10^3^, 10^4^} gene × {10^2^, 10^3^, 10^4^, 10^5^, 10^6^, 10^7^} cell matrices (see Materials and methods). We evaluated whether the calculations could be finished or were interrupted by out-of-memory errors (Figure 1b). We also manually terminated a PCA process that was unable to generate output files within three days (i.e., *dask-ml*). All the terminated jobs are summarized in Additional file 15. To evaluate only the scalability and computability, we set the number of epochs (also known as passes) in `orthiter/gd/sgd` (*OnlinePCA.jl*) to one. However, in actual data analysis, a value several times larger should be used.

Figures 8 and 9 show the calculation time and the memory usage of all the PCA implementations, which can be scaled to a 10^4^ × 10^7^ matrix. `IncrementalPCA` (*sklearn*) and `oocPCA CSV` (*oocRPCA*) were slightly slower than the other implementations (Figure 8), and this was probably because the inputs of these implementations were CSV files while the other implementations used compressed binary files (Zstd). The memory usage of all the implementations were almost the same, except for `IncrementalPCA` (*sklearn*) and `oocPCA CSV` (*oocRPCA*). `oocPCA CSV` (*oocRPCA*) has a parameter that controls the maximum memory usage (*mem*), and we set the value to 10 GB (Additional file 3). Indeed, the memory usage had converged to around 10 GB (Figure 9). This property is considered an advantage of this implementation; users can specify a different value to suit their computational environment.

**Figure 8.**
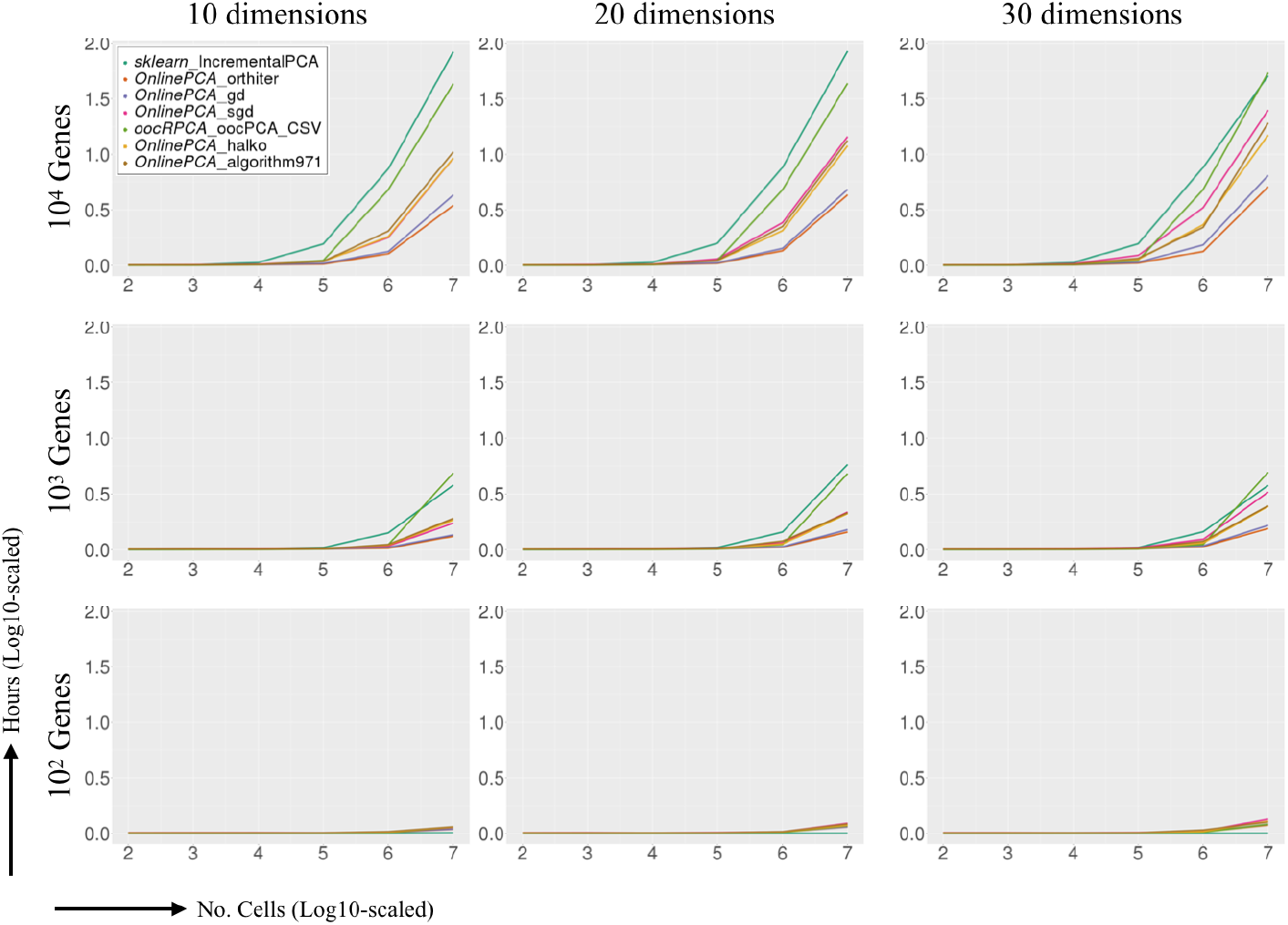
Comparison of the elapsed time for simulated datasets. Synthetic datasets ({10^2^, 10^3^, 10^4^} gene × {10^2^, 10^3^, 10^4^, 10^4^, 10^5^, 10^6^, 10^7^} cell matrices) were randomly generated, and all the out-of-core principal component analysis (PCA) implementations were performed. In each panel, the logarithm of the number of cells is indicated along each *x*-axis, and the elapsed time (hours) is shown along each *y*-axis.

**Figure 9.**
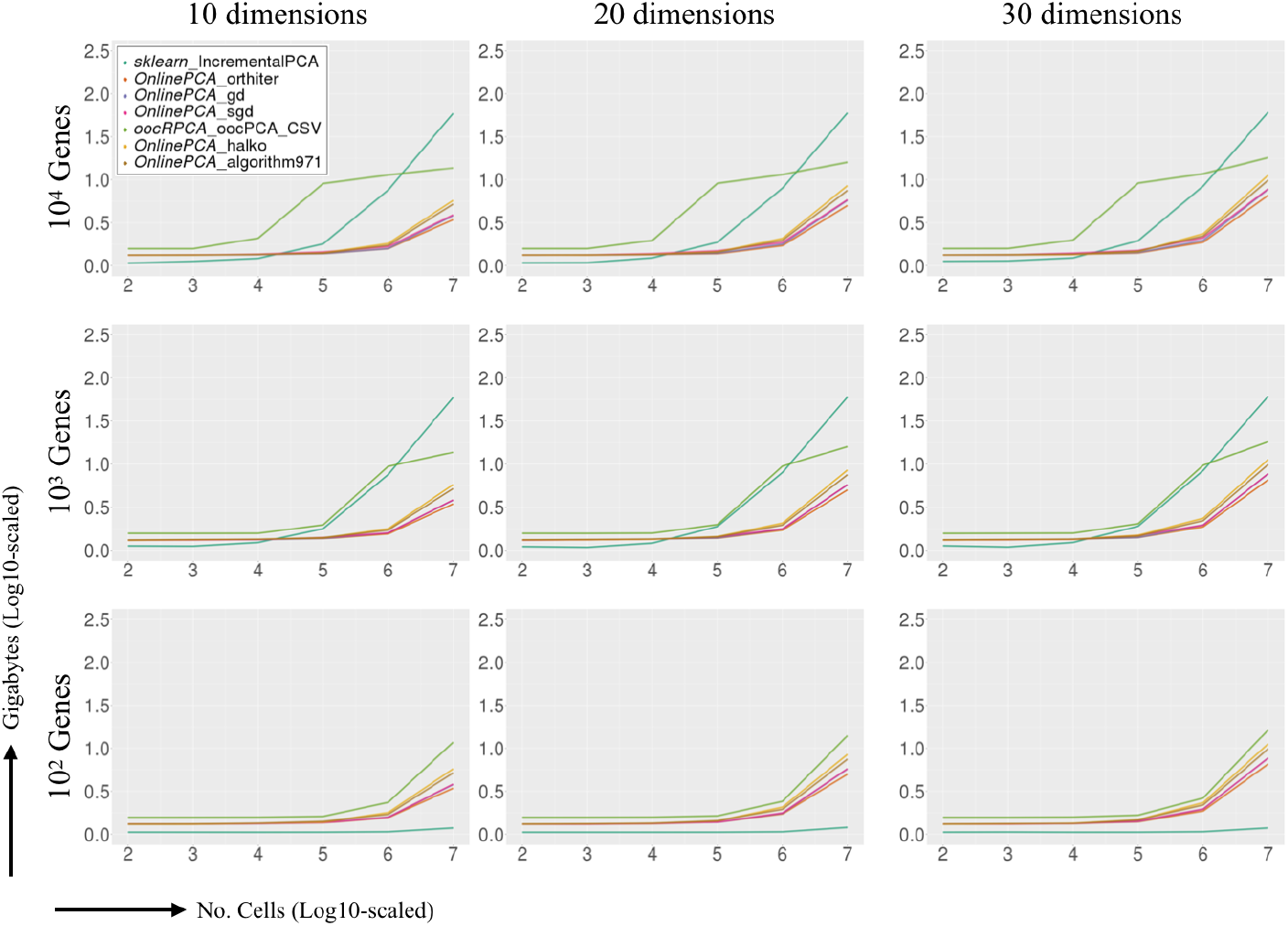
Comparison of the maximum memory usage for simulated datasets. Synthetic datasets ({10^2^, 10^3^, 10^4^} gene × {10^2^, 10^3^, 10^4^, 10^4^, 10^5^, 10^6^, 10^7^} cell matrices) were randomly generated, and all the out-of-core principal component analysis (PCA) implementations were performed. In each panel, the logarithm of the number of cells is indicated along the *x*-axis, and memory usage (GB) is shown along the *y*-axis.

### The relationship between file format and performance

We also counted the passes of the Brain matrix in the out-of-core implementations such as `oocPCA CSV` (R, *oocRPCA*), `IncrementalPCA` (Python, *sklearn*), and `orthiter/gd/sgd/halko/algorithm971` (Julia, *OnlinePCA.jl*) (Figure 10a). In the `oocPCA CSV` (R, *oocRPCA*), `IncrementalPCA` (Python, *sklearn*), the data matrix was passed to these function as the CSV format and in the other out-of-core implementations, the data matrix was firstly binarized and compressed in the Zstd file format. We found that the calculation time was correlated with the number of passes of the implementation. Furthermore, binarizing and data compression substantially accelerated the calculation time. This suggests that the data loading process is very critical for out-of-core implementation and that the overhead for this process has a great effect on the overall calculation time and memory usage.

**Figure 10.**
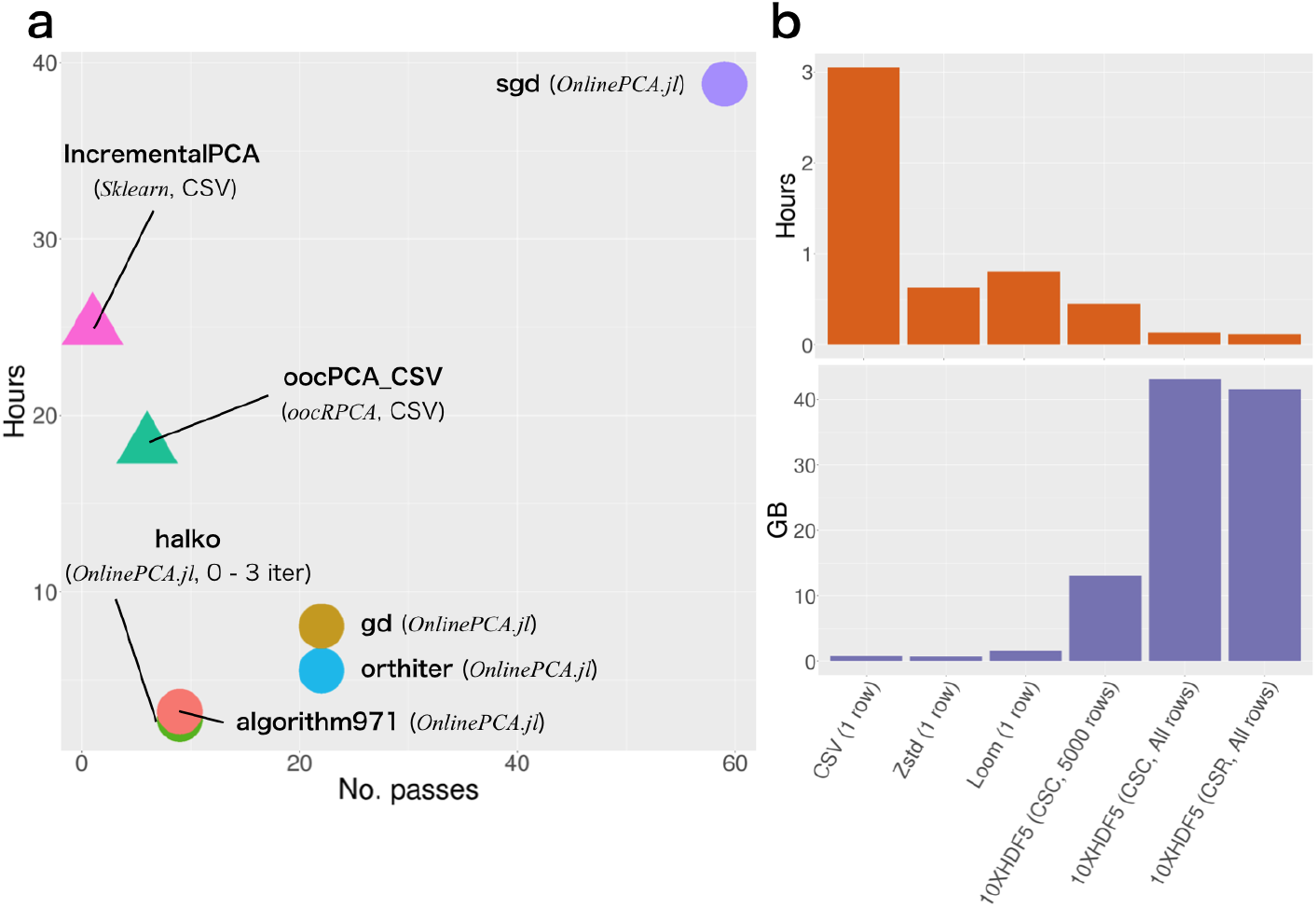
Relationships of the algorithms/implementations, the number of passes, and the file format with the elapsed time for performing principal component analysis (PCA) with the Brain dataset. (a) Number of passes for the data matrix and the computation time for each algorithms/implementations were calculated. (b) Elapsed time and memory usage for one-pass orthogonal iteration were calculated.

Accordingly, using different data formats, such as CSV, Zstd, Loom [93], and hierarchical data format 5 (HDF5), provided by the 10X Genomics (10X-HDF5) for the Brain dataset, we evaluated the calculation time and the memory usage for the simple one-pass orthogonal iteration (qr(XW)), where qr is the QR decomposition, *X* is the data matrix, and *W* represents the 30 vectors to be estimated as the eigenvectors (Figure 10b). For this algorithm algorithm, incremental loading of large block matrices (e.g., 5000 rows) from a sparse matrix was faster than incremental loading of row vectors from a dense matrix, although the memory usage of the former was lower.

While it is not obvious that the usage of a sparse matrix accelerates the PCA with scRNA-seq datasets because scRNA-seq datasets are not particularly sparse compared with data from other fields (cf. recommender systems or social networks [128, 129]), we showed that it has the potential to speed up the calculation time for scRNA-seq datasets.

When all row vectors stored in 10X-HDF5 are loaded at once, the calculation is fastest, but the memory usage is also highest. Because the calculation time and the memory usage have a trade-off and the user’s computational environment is not always high-spec, the block size should be optionally specified as a command argument. For the above reasons, we also developed `tenxpca`, which is a new implementation that performs Li’s method for a sparse matrix stored in the 10X-HDF5 format. Using all the genes in the CSC matrix incrementally, `tenxpca` was able to finish the calculation in 1.3 hours with a maximum memory usage of 83.0 GB. This is the fastest analysis of the Brain dataset in this study.

In addition to `tenxpca`, some algorithms used in this benchmarking, such as orthogonal iteration, GD, SGD, Halko’s method, and Li’s method, are implemented as Julia functions and command line tools, which have been published as a Julia package *OnlinePCA.jl* (Figure 11). When data are stored as a CSV file, they are binarized and compressed in the Zstd file format (Figure 11a) and then some out-of-core PCA implementations are performed. When data are in 10X-HDF5 format, Li’s method is directly performed with the data by `tenxpca` (Figure 11b). We also implemented some functions and command line tools to extract row-wise/column-wise statistics such as mean and variance as well as highly variable genes (HVGs) [130] in an out-of-core manner. Because such statistics are saved as small vectors, they can be loaded by any programming language without out-of-core implementation and used for QC, and the users can select only informative genes and cells. After QC, the filtering command removes low-quality genes/cells and generates another Zstd file.

**Figure 11.**
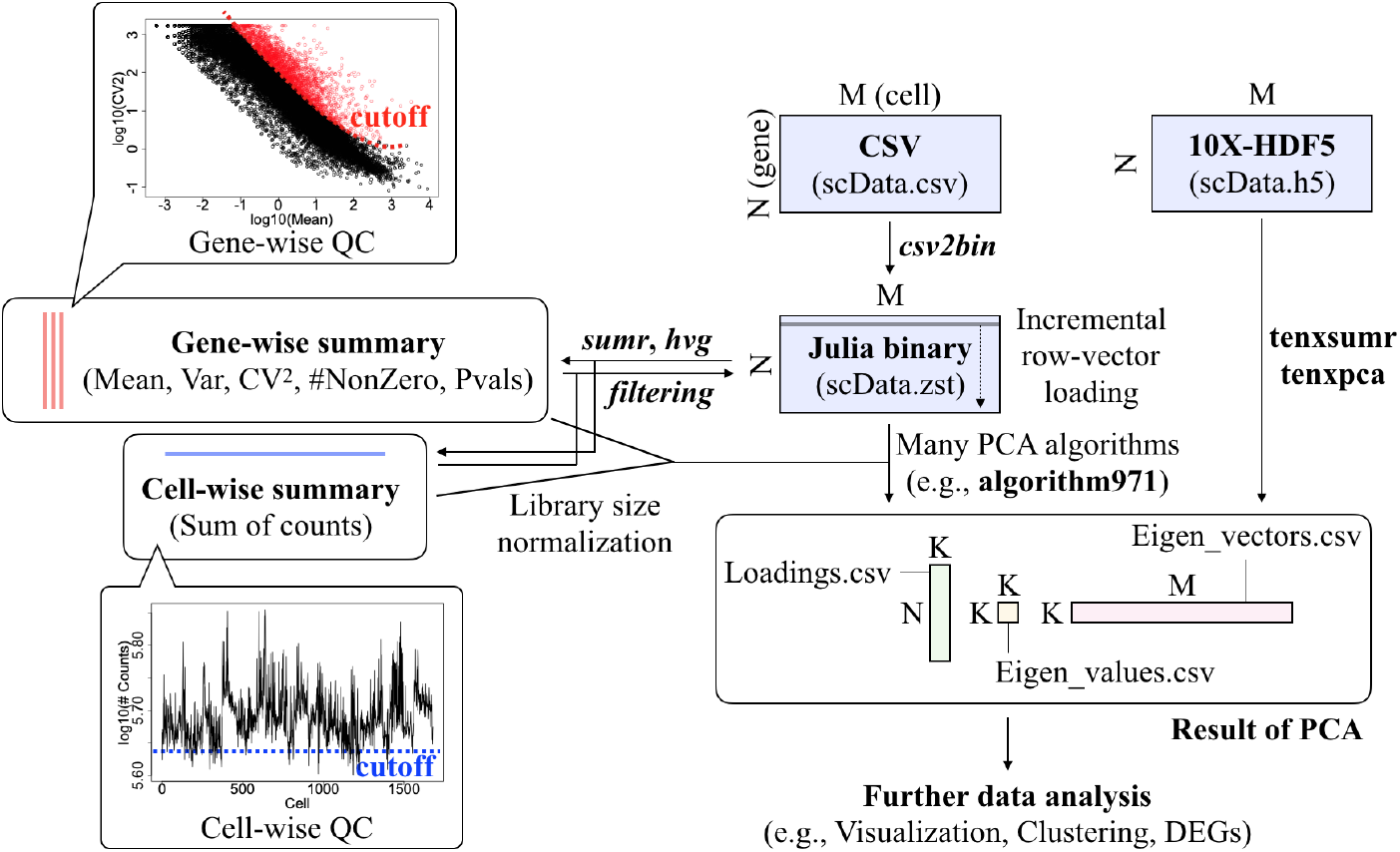
OnlinePCA.jl schematic. Input CSV files were first saved as a binary file with the csv2bin command and analyzed with the `gd` and `sgd` commands, which perform incremental principal component analysis (PCA). When using the HDF5 file format defined by 10X Genomics, we converted the file to CSV format using an in-house Python script. Gene-wise or cell-wise summary statistics were calculated using the `sumr` command. Highly variable genes can also be calculated with the hvg command. Because the gene-wise and cell-wise summary statistics are expressed as small vectors, they can be used to perform precise data quality control (QC) with any programming language without out-of-core implementations. After QC, the `filtering` command removed low-quality genes and cells using a user-specified index. Combined with the small size vectors, some out-of-core PCA implementations, such as `orthiter/gd/sgd/halko/algorithm971` have commands to incrementally update eigenvectors from the row vector of the data matrix. The `tenxpca` command directly performed `algorithm971` on 10X-HDF files.

## Discussion

### Guidelines for users

Based on all the benchmarking results and our implementation in this work, we propose some user guidelines (Figure 12). Considering that bioinformatics studies combine multiple tools to construct a user’s specific workflow, the programming language is an important factor in selecting the right PCA implementation. Therefore, we categorized the PCA implementations according to language (i.e., R [111], Python [112], and Julia [113]; Figure 12, column-wise). In addition to the data matrix size, we also categorized implementations according to the way they load data (in-memory or out-of-core) as well as their input matrix format (dense or sparse, Figure 12, row-wise). Here, we define the GC-value of a data matrix as the number of genes *×* the number of cells.

**Figure 12.**
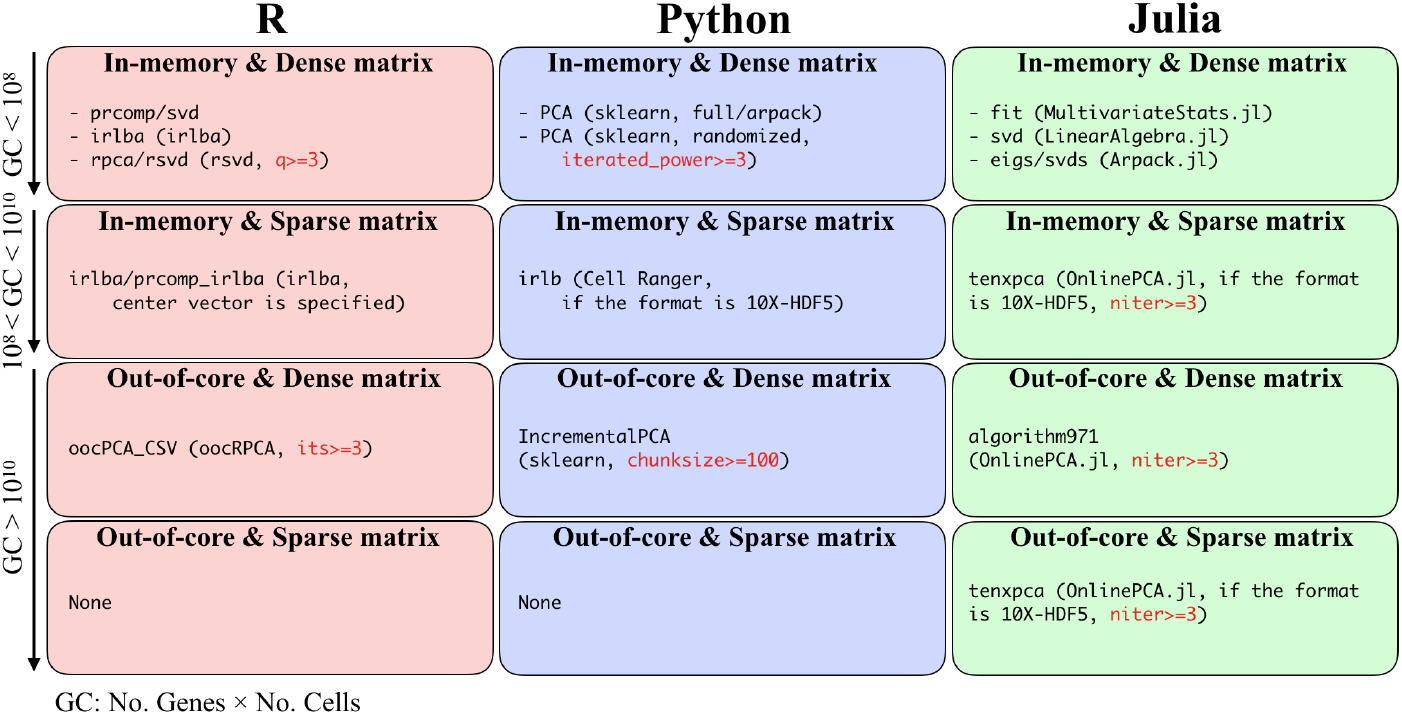
User guidelines. Recommended PCA implementations categorized based on written language and matrix size.

If the data matrix is not too large (e.g., GC *≤* 10^7^), the data matrix can be loaded as a dense matrix, and full-rank SVD in LAPACK is then accurate and optimal (in-memory & dense matrix). In such a situation, the wrapper functions for the full-rank SVD written in each language are suitable. However, if the data matrix is much larger (e.g., GC *≥* 10^8^), an alternative to the full-rank SVD is needed. Based on the benchmarking results, we recommend IRLBA, IRAM, Halko’s method, and Li’s method as alternatives to the full-rank SVD. For intermediate GC-values (10^8^ *≤* GC *≤* 10^10^), if the data matrix can be loaded into memory as a sparse matrix, some implementations for these algorithms are available (in-memory & sparse matrix). In particular, such implementations are effective for large data matrices stored in 10X-HDF5 format using CSC format. Seurat2 [49] also introduces this approach by combining the matrix market format (R, *Matrix*) and `irlba` function (R, *irlba*). When the data matrix is dense and cannot be loaded into memory space (e.g., GC *≥* 10^10^), the out-of-core implementations, such as `oocPCA CSV` (R, *oocRPCA*), `IncrementalPCA` (Python, *sklearn*), and `algorithm971` (Julia, *OnlinePCA.jl*), are useful (dense matrix & out-of-core). If the data matrix is extremely large and cannot be loaded into memory even if the data are formatted as a sparse matrix, out-of-core PCA implementations for sparse matrix are needed. Actually, R cannot load the Brain dataset, even if the data is formatted as a sparse matrix (https://github.com/satijalab/seurat/issues/1644). Hence, in such a situation, `tenxpca` can be used if the data is stored in the 10X-HDF5 format.

The PCA implementations examined in this work are affected by various parameters. For example, in `gd` and `sgd` (*OnlinePCA.jl*), the result is sensitive to the value of learning parameters and the number of epochs. Therefore, a grid-search of such parameters is necessary (Additional file 17). When using `IncrementalPCA` (*sklearn*), the user specifies the chunk size of the input matrix, and a larger value slightly improves the accuracy of PCA (Additional file 16) and the calculation time (Figure 8), although there is a trade-off between these properties and memory usage (Figure 9). Both Halko’s method and Li’s method have a parameter for specifying the number of power iterations (*niter*), and this iteration step sharpens the distribution of eigenvalues and enforces a more rapid decay of singular values ([114] and Additional file 3). In our experiments, the value of *niter* is critical for achieving accuracy, and we highly recommend a *niter* value of three or larger (Additional file 18). In some implementations, the default values of the parameters are specified as inappropriate values or cannot be accessed as a function parameter. Therefore, users should carefully set the parameter or select an appropriate implementation.

### Guidelines for developers

We have also established guidelines for developers. Many technologies such as data formats, algorithms, and computational frameworks and environments are available for developing fast, memory-efficient, and scalable PCA implementations (Additional file 19). Here, we focus on two topics.

The first topic is “loss of sparsity.” As described above, the use of a sparse matrix can effectively reduce memory space and accelerate calculation, but developers must be careful not to destroy the sparsity of a sparse matrix. PCA with a sparse matrix is not equivalent to SVD with a sparse matrix; in PCA, all sparse matrix elements must be centered by the subtraction of gene-wise average values. Once the sparse matrix *X* is centered (*X − X*_mean_), where *X*_mean_ has gene-wise average values as column vectors, it becomes a dense matrix and the memory usage is significantly increased. Obviously, the explicit calculation of the subtraction described above should be avoided. In such a situation, if multiplication of this centered matrix and a dense vector/matrix is required, the calculation should be divided into two parts, such as (*X − X*_mean_) *W* = *XW − X*_mean_*W*, where *W* represents the vectors to be estimated as eigenvectors, and these parts should be calculated separately. If one or both parts require more than the available memory space, such parts should be incrementally calculated in an out-of-core manner. There are actually some PCA implementations that can accept a sparse matrix, but they may require very long calculation times and large memory space because of a loss of sparsity (cf. `rpca` of *rsvd* https://github.com/cran/rsvd/blob/7a409fe77b220c26e88d29f393fe12a20a5f24fb/R/rpca.R#L158). To our knowledge, only `prcomp_irlba` in *irlba* (https://github.com/bwlewis/irlba/blob/8aa970a7d399b46f0d5ad90fb8a29d5991051bfe/R/irlba.R#L379), `irlb` in *Cell Ranger* (https://github.com/10XGenomics/cellranger/blob/e5396c6c444acec6af84caa7d3655dd33a162852/lib/python/cellranger/analysis/irlb.py#L118), `safe_sparse_dot` in *sklearn* (https://scikit-learn.org/stable/modules/generated/sklearn.utils.extmath.safe_sparse_dot.html), and `tenxpca` in *OnlinePCA.jl* (https://github.com/rikenbit/OnlinePCA.jl/blob/c95a2455acdd9ee14f8833dc5c53615d5e24b5f1/src/tenxpca.jl#L183) deal with this issue. Likewise, as an alternative to the centering calculation, `MaxAbsScaler` in *sklearn* (https://scikit-learn.org/stable/modules/generated/sklearn.preprocessing.MaxAbsScaler.html) introduces a scaling method in which the maximum absolute value of each gene vector becomes one, thereby avoiding the loss of sparsity.

The second topic is “lazy loading.” The out-of-core PCA implementations used in this benchmarking explicitly calculate centering, scaling, and all other relevant arithmetic operations from the extracted blocks of the data matrix. However, to reduce the complexity of the source code, it is desirable to calculate such processes as if the matrix was in memory and only when the data are actually required, so the processes are lazily evaluated on the fly. Some packages, such as `DeferredMatrix` in *BiocSingular* (R/Bioconductor, https://bioconductor.org/packages/devel/bioc/html/BiocSingular.html), *CenteredSparseMatrix* (Julia, https://github.com/jsams/CenteredSparseMatrix), *Dask* [115] (Python, https://dask.org), and *Vaex* (Python, https://vaex.io/), support lazy loading.

### Future perspective

In this benchmarking study, we found that PCA implementations based on full-rank SVD are accurate but cannot be scaled for use with high-throughput scRNA-seq datasets such as the BrainSpinalCord and Brain datasets, and alternative implementations are thus required. Some methods approximate this calculation by using truncated SVD forms that are sufficiently accurate as well as faster and more memory-efficient than full-rank SVD. The actual memory usage highly depends on whether an algorithm is implemented as out-of-core and whether sparse matrix can be specified as input. Some sophisticated implementations, including our *OnlinePCA.jl*, can handle such issues. Other PCA algorithms, such as downsampling and SGD, are actually not accurate, and their use risks overlooking cellular subgroups contained within scRNA-seq datasets. These methods commonly update eigenvectors with small fractions of the data matrix, and this process may overlook subgroups or subgroup-related gene expression, thereby causing the observed inaccuracy. Our literature review, benchmarking, special implementation for scRNA-seq datasets, and guidelines provide important resources for new users and developers tackling the UML of high-throughput scRNA-seq.

Although the down-stream analyses of PCA vary widely, and we could not examine all the topics of scRNA-seq analyses, such as rare cell-type detection [59, 60] and pseudotime analysis [13,62–66], differences among PCA algorithms might also affect the accuracy of such analyses. Butler *et al*. showed batch effect removal can be formalized as canonical correlation analysis (CCA) [49], which is mathematically very similar to PCA. The optimization of CCA is also formalized in various ways, including randomized CCA [131] or SGD of CCA [132].

This work also sheds light on the effectiveness of randomized SVD. This algorithm is popular in population genetic studies [110]. In the present study, we also assessed its effectiveness with scRNA-seq datasets with high heterogeneity. This algorithm is relatively simple and some studies have implemented it from scratch (Table 1). Simplicity may be the most attractive feature of this algorithm.

There are also many focuses of recent PCA algorithms (Additional file 19). The randomized subspace iteration algorithm, which is a hybrid of Krylov and Rand methodologies, was developed based on randomized SVD [133,134]. In pass-efficient or one-pass randomized SVD, some tricks to reduce the number of passes have been considered [135, 136]. TeraPCA, which is a software tool for use in population genetics studies, utilizes the Mailman algorithm to accelerate the expectation– maximization algorithms for PCA [137, 138]. Townes *et al*. recently proposed the use of PCA for generalized linear models (GLM-PCA) and unified some PCA topics, such as log-transformation, size factor normalization, non-normal distribution, and feature selection, in their GLM framework [139, 140]. Although such topics are beyond the scope of the present work, the current discussion will be useful for the development and application of such methods above.

## Materials and methods

### Empirical datasets

The gene expression matrix and cell type labels for the PBMCs dataset and the Brain dataset [39] were downloaded from the 10X Genomics website (https://support.10xgenomics.com/single-cell-gene-expression/datasets/pbmc_1k_protein_v3 and https://support.10xgenomics.com/single-cell/datasets/1M_neurons, respectively). The gene expression matrix and cell type labels for the Pancreas dataset [40] and the BrainSpinalCord dataset [41] were retrieved from the GEO database (GSE84133 and GSE110823, respectively). For the Pancreas dataset, only the sample of GSM2230759 was used. The genes of all matrices with zero variance were removed because such genes are meaningless for PCA calculation. We also removed the ERCC RNA Spike-Ins, and the number of remaining genes and cells are summarized in Table 2. Additionally, we investigated the effect of feature selection on clustering accuracy (Additional file 20).

### Simulated datasets

All count datasets were generated by the R `rnbinom` (random number based on a negative binomial distribution) function with shape and rate parameters of 0.4 and 0.3, respectively. Matrices of {10^2^, 10^3^, 10^4^} genes *× {*10^2^, 10^3^, 10^4^, 10^5^, 10^6^, 10^7^} cells were generated.

### Benchmarking procedures

Assuming digital expression matrices of unique molecular identifier (UMI) counts, all the data files, including real and synthetic datasets, were in CSV format. When using the Brain dataset, the matrix stored in 10X-HDF5 format was converted to CSV using our in-house Python script (https://gist.github.com/kokitsuyuzaki/5b6cebcaf37100c8794bdb89c7135fd5).

After being loaded by each PCA implementation, the raw data matrix *X*_raw_ was converted to normalized values by count per median (CPMED [141–143]) normalization according to the formula 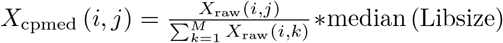, where *M* is the number of columns and Libsize is the column-wise sum of counts of *X*. After normalization, *X*_cpmed_ was transformed to *X* by the logarithm-transformation *X* = log_10_ (*X*_cpmed_ + 1), where log_10_ is the element-wise logarithm. In all the randomized PCA implementation, random seed was fixed.

When *X*_raw_ was extremely large and could not be loaded into the memory space all at once, we prepared two approaches to perform PCA with *X*. When PCA implementations are `orthiter, gd, sgd, halko`, or `algorithm971` (*OnlinePCA.jl*), each row-vector of *X*_raw_ is normalized using the pre-calculated Libsize by the `sumr` command, then log-transformed, and finally used for each of the PCA algorithms. When using other out-of-core PCA implementations such as `IncrementalPCA` (*sklearn*), `oocPCA CSV` (*oocRPCA*), or `PCA` (*dask-ml*), there is no option to normalize and log-transform each row-vector of *X*_raw_, so we first calculated *X*_cpmed_ using our in-house Python script (https://gist.github.com/kokitsuyuzaki/5b6cebcaf37100c8794bdb89c7135fd5), which was then used for the input matrix of the PCA implementations.

We also investigated the effect of differences in normalization methods on the PCA results (Additional file 21). When performing each PCA implementation based on the truncated SVD, the number of PCs was specified in advance (Table 2).

Although it is unclear how many cells should be used in downsampling, one empirical analysis [94] suggests that 20,000 to 50,000 cells are sufficient for clustering and detecting subpopulations in the Brain dataset. Thus 50, 000*/*1, 300, 000*×*100 = 3.8% of cells were sampled from each dataset and used for the downsampling method. When performing `IncrementalPCA` (*sklearn*), the row-vectors, which match the number of PCs, were extracted until the end of the lines of the files. When performing irlb(*Cell Ranger*), the loaded dataset was first converted to a scipy sparse matrix and passed to it because this function supports sparse matrix data stored in 10X-HDF5 format. When performing the benchmark, conversion time and memory usage were also recorded. When performing all the functions of *OnlinePCA.jl*, including `orthiter/gd/sgd/halko/algorithm971`, we converted the CSV data to Zstd format, and the calculation time and the memory usage were recorded in the benchmarking for fairness. For `orthiter, gd`, and `sgd` (*OnlinePCA.jl*), calculations were performed until they converged (Additional file 17). For all the randomized SVD implementations, the *niter* parameter value was set to 3 (Additional file 18). When performing `oocPCA_CSV`, the users can also use `oocPCA_BIN`, which performs PCA with binarized CSV files. The binarization is performed by the `csv2binary` function, which is also implemented in the *oocRPCA* package. Although data binarization accelerates the calculation time for PCA itself, we confirmed that `csv2binary` is based on in-memory calculation, and in our computing environment, `csv2binary` was terminated by an out-of-memory error. Accordingly, we only used `oocPCA_CSV`, and the CSV files were directly loaded by this function.

### Computational environment

All computations were performed on two-node machines with Intel Xeon E5-2697 v2 (2.70 GHz) processors and 128 GB of RAM, four-node machines with Intel Xeon E5-2670 v3 (2.30 GHz) processors and 96 GB of RAM, and four-node machines with Intel Xeon E5-2680 v3 (2.50 GHz) processors and 128 GB of RAM. Storage among machines was shared by NFS, connected using InfiniBand. All jobs were queued by the Open Grid Scheduler/Grid Engine (v2011.11) in parallel. The elapsed time and maximum memory usage were evaluated using the GNU `time` command (v1.7).

### Reproducibility

All the analyses were performed on the machines described above. We used R v3.5.0, Python v3.6.4, and Julia v1.0.1 in the benchmarking; for t-SNE and CSV conversion of the Brain dataset, we used Python v2.7.9. The *Sklearn* (Python) package was used to perform *k*-means and GMM clustering methods. The igraph (R), nn2 (R), and Matrix (R) packages were used to perform Louvain clustering (Additional file 6). The hdbscan (Python) package was used to perform HDBScan clustering. The bhtsne (Python) package was used to perform t-SNE. Lastly, the umap (Python) package was used to perform UMAP. All the programs used to perform the PCA implementations in the benchmarking are summarized in Additional file 3. Orthogonal iteration, GD, SGD, Halko’s method, and Li’s method are implemented as `orthiter, gd, sgd, halko`, and `algorithm971`, respectively, which are the Julia functions or commands for *OnlinePCA.jl* (https://github.com/rikenbit/OnlinePCA.jl). We also published the script files used to perform the benchmarking (https://github.com/rikenbit/onlinePCA-experiments).

## Abbreviations

PCA: principal component analysis
scRNA-seq: single-cell RNA sequencing
sci-RNA-seq: single-cell combinatorial-indexing RNA-sequencing analysis
UML: unsupervised machine learning
QC: quality control
PC: principal component
EVD: eigenvalue decomposition
SVD: singular value decomposition
SimT: similarity transformation-based
DS: downsampling-based
SU: SVD update-based
Krylov: Krylov subspace-based
GD: gradient descent-based
Rand: Random projection-based
*Sklearn*: *scikit-learn*
SKL: sequential Karhunen-Loeve transform
IRLBA: augmented implicitly restarted Lanczos bidiagonalization
IRAM: implicitly restarted Arnoldi method
GD: gradient descent
SGD: stochastic gradient descent
t-SNE: t-stochastic neighbor embedding
UMAP: uniform manifold approximation and projection
FIt-SNE: Fourier transform-accelerated interpolation-based t-stochastic neighbor embedding
oocPCA: out-of-core PCA
GMM: Gaussian mixture model
ARI: adjusted Rand index
Zstd: Zstandard
UMI: unique molecular identifier
CSV: comma-separated values
HDF5: hierarchical data format 5
10X-HDF5: HDF5 provided by 10X Genomics
CSC: compressed sparse column format
CSR: compressed sparse row format
CCA: canonical correlation analysis
GLM: generalized linear models
CPMED: Count per median
HVGs: highly variable genes

## Author’s contributions

KT and HS surveyed the PCA algorithms and implementations. KT and IN designed the benchmarking test. KT and KS implemented the Julia program and performed all the analyses. KT retrieved and preprocessed the test dataset to evaluate the proposed method. All the authors have written, read, and approved the manuscript.

## Acknowledgements

We thank Mr. Akihiro Matsushima and Mr. Manabu Ishii for their assistance with the IT infrastructure for the data analysis. We are also grateful to all member of the Laboratory for Bioinformatics Research, RIKEN Center for Biosystems Dynamics Research for their helpful advice. Computations were partially performed on the NIG supercomputer at ROIS National Institute of Genetics.

## Additional Files

Additional File 1 — Review of existing PCA algorithms and implementations. (PDF 271 KB)

Additional File 2 — Pseudo-code of all the PCA algorithms. (PDF 178 KB)

Additional File 3 — Source code of all the PCA implementations. (PDF 59 KB)

Additional File 4 — Results of t-SNE of all the PCA implementations. (PNG 623 KB)

Additional File 5 — Results of UMAP of all the PCA implementations. (PNG 368 KB)

Additional File 6 — Results of clustering methods of all the PCA implementations (PDF 3.6 MB)

Additional File 7 — Eigenvectors of all the PCA implementations (PBMCs and Pancreas). (PNG 308 MB)

Additional File 8 — Pair plots of all the PCA (PBMCs) implementations. (TAR.GZ 649 KB)

Additional File 9 — Pair plots of all the PCA (Pancreas) implementations. (TAR.GZ 4.9 MB)

Additional File 10 — Pair plots of all the PCA (BrainSpinalCord) implementations. (TAR.GZ 3.1 MB)

Additional File 11 — Pair plots of all the PCA (Brain) implementations. (TAR.GZ 5.8 MB)

Additional File 12 — Number of singleton clusters. (PNG 271 KB)

Additional File 13 — Eigenvectors of all the PCA implementations (BrainSpinalCord and Brain). (PNG 532 KB)

Additional File 14 — Loading vectors of all the PCA implementations (PBMCs and Pancreas). (PNG 349 KB)

Additional File 15 — Crashed jobs caused by out-of-memory errors. (TXT 882 B)

Additional File 16 — Parameter tuning of the `IncrementalPCA` implementations. (PDF 445 KB)

Additional File 17 — Parameter tuning of the orthogonal iteration, gradient descent, and stochastic gradient descent implementations. (PDF 1.3 MB)

Additional File 18 — Parameter tuning of the randomized SVD implementations. (PDF 734 KB)

Additional File 19 — Developer guidelines. (PNG 1.1 MB)

Additional File 20 — Effect of feature selection on clustering accuracy. (PDF 1 MB)

Additional File 21 — Comparison of normalizing size factors. (HTML 1.4 MB)

