## Additional File 1 for "Benchmarking principal component analysis for large-scale single-cell RNA-sequencing"

### Review of existing PCA algorithms and implementations

Here, we review the existing principal component analysis (PCA) algorithms and their implementations. PCA is formalized as the eigenvalue decomposition (EVD [90]) of the covariance matrix of the data matrix or the singular value decomposition (SVD [90]) of the data matrix. To perform PCA, the most widely used PCA function in the R language [111] is probably the `prcomp` function, which is a standard R function [42-44,51,55,60,63,65,74,77,82,85,91]. Likewise, users and developers of the Python language [112] may use the `PCA` function in *scikit-learn* (*sklearn*) [92,52,56,58,93], which is a Python package for machine learning. These are actually wrapper functions for performing SVD with LAPACK subroutines such as `DGESVD` (QR method-based [90]) or `DGESDD` (divide-and-conquer method-based [90]), and both subroutines utilize the Golub-Kahan method [90]. In this method, the covariance matrix is tri-diagonalized by Householder transformation, and then the tri-diagonalized matrix is diagonalized by the QR method or the divide-and-conquer method (**Figure S 3-1a**). Such a two-step transformation is commonly performed by the sequential similarity transformation expressed as $\mathbf{M}_k \ldots \mathbf{M}_2 \mathbf{M}_1 \mathbf{X} \mathbf{M}_1 \mathbf{M}_2 \ldots \mathbf{M}_k$, where $\mathbf{X}$ is an $n$-by-$n$ covariance matrix, $\mathbf{M}_k$ is an $n$-by-$n$ invertible matrix, and $\boldsymbol{k}$ is the step of the transformation. Likewise, when the input data matrix is asymmetric, the matrix is bi-diagonalized and then tri-diagonalized. When the matrix is finally diagonalized at the $\boldsymbol{k}$-th step, the diagonal elements become eigenvalues and $\mathbf{M} = \mathbf{M}_1 \mathbf{M}_2 \ldots \mathbf{M}_k$ becomes the set of corresponding eigenvectors. Although the Golub-Kahan method is the most widely used SVD algorithm, this method has some drawbacks.

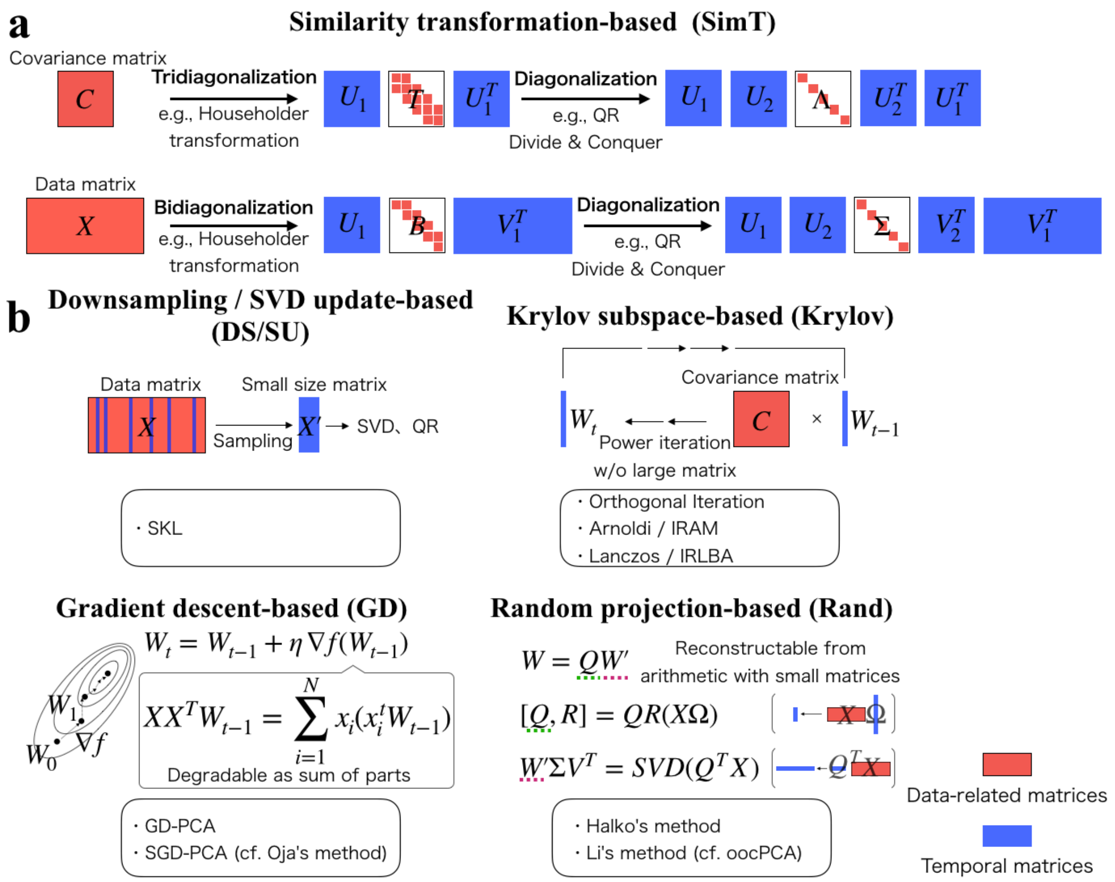


**Supplementary Figure 1-1 | Some fast and memory-efficient PCA algorithms** (a) Full-rank SVD based on LAPACK and (b) truncated SVD based on the recent fast and memory-efficient PCA algorithms.

First, the large dense matrix **M** must be temporarily saved and incrementally updated in each step; therefore, the memory space is filled quickly. Second, when the matrix is large, the data matrix itself is difficult to load and causes an out-of-memory error. For the above reasons, the Golub-Kahan method cannot be directly applied to large-scale scRNA-seq datasets.

There are faster and more memory-efficient PCA algorithms. Contrary to the full-rank SVD solved by LAPACK, such algorithms are formalized as truncated SVD, in which only some of the top PCs are calculated. We classify these methods into five categories (**Figure S 3-1b**).

The first category consists of downsampling-based methods [94]. In these methods, SVD is first performed with a small matrix consisting of cells randomly sampled from the original large matrix. The remaining cells are then projected onto the low-dimensional space spanned by the eigenvectors learned from the small matrix. The effectiveness of this method in scRNA-seq studies has been evaluated by Bhaduri *et al.* [94] (**Table 1**).

The second category is SVD update [95], which repeatedly performs SVD using subsets of the data sampled from the data matrix and incrementally updates the result. Sequential Karhunen-Loève transform (SKL) [95], which is a kind of SVD update, is used in *loompy* (http://loompy.org) (**Table 1**).

The third category consists of Krylov subspace-based methods [96-99]. The most typical method within this category is the power method, which iteratively multiplies a vector **w** with a covariance matrix **X** and normalizes **w**. Within some iterations, **w** converges to the eigenvector (PC1) corresponding to the largest eigenvalue. Since the way to calculate the higher PCs is not obvious, there are some algorithms, such as orthogonal iteration (block power method, subspace iteration, or simultaneous iteration), the Lanczos method, and the Arnoldi method [96]. Orthogonal iteration performs the power method with multiple initial vectors in parallel, and each power iteration step performs QR decomposition for orthonormalization. Contrary to orthogonal iteration, Lanczos and Arnoldi methods respectively introduce Lanczos and Arnoldi processes to generate vectors that are orthogonal to each other. To make the convergence faster, both methods also introduce "restart" strategies such as the augmented implicitly restarted Lanczos bidiagonalization algorithm (IRLBA [100]) and implicitly restarted Arnoldi methods (IRAM [101]), in which new initial vectors are calculated by the accumulated result of Lanczos or Arnoldi processes. In contrast to the Golub-Kahan method, these methods do not generate large dense temporary matrices, and when the data matrix is sparse, these methods are compatible with a sparse matrix format and can be accelerated. *Cellranger* [22], *Seurat2* [49],

*Scanpy* [93], *SAFE* [76], *Scran* [50], *Giniclust2* [59], *MAGIC* [52], *Harmony* [57], and *Scater* [82] use IRLBA for PCA functions (**Table 1**). Although IRAM appears not to have been used in scRNA-seq studies, the effectiveness of its use with population genetic datasets has been argued recently [102].

The fourth category is gradient descent (GD, or steepest descent)-based methods. In this method category, the gradient of the objective function is calculated, and the initial vectors are updated to the reverse direction of the gradient. Although GD utilizes all the data to calculate the gradient (i.e., full gradient), stochastic gradient descent (SGD) calculates the gradient with a subset of the data (i.e., a stochastic gradient). Although these PCA algorithms are sometimes used for situations in which the data are incrementally observed, such as subspace tracking [103], these methods can also be fast and memory-efficient because the calculation of the full/stochastic gradient is decomposable as the sum of the gradient of individual data points. Although these methods appear not to have been used in scRNA-seq studies, in image processing studies, this method is known as Oja's method or the generalized Hebbian algorithm [104-106].

The fifth category comprises random projection-based methods, in which a data matrix is randomly projected onto lower dimensions and basic linear algebraic methods such as QR decomposition and SVD are performed on the smaller matrix. Because most calculations are performed for these random lower dimensions, these methods can be fast and memory efficient. Surprisingly, in this method, the SVD of the original data matrix can be accurately reconstructed from arithmetic with a small matrix with low reconstruction error [107,108]. Although Halko's method is known as a randomized SVD algorithm, Li *et al*. also modified the preconditioning step so that the calculation time is improved (algorithm971 [109]). Halko's method is used in *scanpy* [93], *SIMLR* [75], and *SEQC* [89], and algorithm971 is implemented in *CellFishing.jl* [61] (**Table 1**). Halko's method is also used in population genetic studies [110].

Notably, the acceleration techniques of the above algorithms are based on random row/column selection or random projection of data matrices, both of which are used to make a large matrix smaller. When these processes are used in an out-of-core (also known as, online, incremental, or on-disk) implementation, in which only a subset of the data matrix is loaded into the memory and used to incrementally update the calculation, these algorithms might be scalable for use with even scRNA-seq datasets consisting of millions of cells. For example, in fast Fourier transform-accelerated interpolation-based t-stochastic neighbor embedding (*FIt-SNE* [69]), algorithm971 is implemented in an out-of-core manner and referred to as out-of-core PCA (oocPCA). However, most PCA implements load all the elements of a data matrix into the memory-space simultaneously. Therefore, the order of memory usage of such algorithms is commonly $O(NM)$, where $N$ is the number of genes and $M$ is the number of cells. To extend the scope of algorithms used in the benchmarking, we originally implemented algorithms that included orthogonal iteration, GD, SGD, Halko's method, and algorithm971 in an out-of-core manner. These algorithms were implemented in Julia [113].
