## Additional File 2 for "Benchmarking principal component analysis for large-scale single-cell RNA-sequencing"

### Pseudo-code of all the PCA algorithms

### Golub-Kahan method (EVD)

**Algorithm** Golub-Kahan method (EVD)

**Parameter:** Number of epoch S, Number of dimension K

**Input:** Data matrix $X \in R^{N \times M}$

**Output:** Eigenvectors $W \in R^{M \times K}$

$$C = XX^T / M$$

$$U_1 T U_1^T = \text{Householder (C)}$$

$$U_2 \Lambda U_2^T = \text{Diagonalization (T)}$$

$$U = U_1 U_2$$

$$W = X^T U$$

Ref : Gene H Golub, Matrix Computations, 2012

### Golub-Kahan method (SVD)

**Algorithm** Golub-Kahan method (SVD)

**Parameter:** Number of epoch S, Number of dimension K

**Input:** Data matrix $X \in R^{N \times M}$

**Output:** Eigenvectors $W \in R^{M \times K}$

$$U_1 B W_1^T = \text{Householder}\,(X)$$

$$U_2 \Sigma W_2^T = \text{Diagonalization}\,(B)$$

$$W = W_1 W_2$$

Ref : Gene H Golub, Matrix Computations, 2012

### Downsampling

**Algorithm** Downsampling

**Parameter:** Number of epoch S, Number of dimension K

**Input:** Data matrix $X \in R^{N \times M}$

**Output:** Eigenvectors $W \in R^{M \times K}$

$$X' = \mathrm{RandomColumnSampling}\,(X)$$
$$U'\Sigma' W'^{T} = \mathrm{svd}\,(X')$$
$$W = X^{T} U'$$

Ref : Bhaduri, A et al., Identification of cell types in a mouse brain single-cell atlas using low sampling coverage, BMC Biology, 2018

### SKL

**Algorithm** SKL

**Parameter:** Number of epoch S, Number of dimension K

**Input:** Block matrices by data matrix ($A_0$; $A_1$; ...; $A_{I-1}$) $A_i \in R^{N_i \times M}$, forgetting factor : *ff*

**Output:** Eigenvectors $W \in R^{M \times K}$

$$U_0 D_0 W_0^T = \mathrm{svd}\left(\mathrm{A}_0\right)$$

**for** i = 1, 2, …, I-1 **do**

$$[D'_{i-1}, W'^T_{i-1}] = \mathrm{qr}\left(\mathrm{ff} \cdot \mathrm{D}_{i-1}\mathrm{W}^{\mathrm{T}}_{i-1}; \mathrm{A}_i\right)$$

$$\hat{U}_{i-1}\hat{D}_{i-1}\hat{W}^T = \mathrm{svd}\left(\mathrm{D}'_{i-1}, \mathrm{rank} = \mathrm{K}\right)$$

$$W_{i-1} = W'_{i-1}\hat{W}_{i-1}$$

**end for**

Ref : Levy, A et al., Sequential Karhunen-Loeve Basis Extraction and its Application to Images, IEEE Transactions on Image Processing, 2000

### Orthogonal Iteration

**Algorithm** Orthogonal Iteration

**Parameter:** Number of epoch S, Number of dimension K

**Input:** Data matrix $X \in R^{N \times M}$

Initial matrix $W_0 = (w_1, \ldots, w_K)$ (diagonal element is 1, otherwise 0) $\in R^{M \times K}$

**Output:** Eigenvectors $W \in R^{M \times K}$

**for** $t = 1, 2, \ldots, S$ **do**

$[W_t, \sim] = qr(X^T X W_{t-1})$

**end for**

Ref : Gene H Golub, Matrix Computations, 2012

### Arnoldi method (Arnoldi process)

**Algorithm** Arnoldi method (Arnoldi process)

**Parameter:** Number of epoch S, Number of dimension K

**Input:** Data covariance matrix $C = X^T X/N \in R^{N \times N}$

Initial vector $q_1 \in R^{N \times 1}$

**Output:** Eigenvectors $W \in R^{K \times M}$

**for** t = 1, 2, …, S **do**

$v = Cq_t$ t-th vector generation

**for** i = 1, ..., t **do**

$h_{it} = q_i^T v$

$v = v - h_{it} q_i$ Orthogonalization to 1 - t-th vectors

**end for**

$H_{t+1,t} = \|v\|$

$q_{t+1} = v / H_{t+1,t}$ Normalization

**end for**

$[Y, \Sigma, Y^T] = \mathrm{evd}(H_t)$

$W = Q_t Y$

$$C = W \Sigma W^T$$

$$C = QHQ^T$$

$$C = QY\Sigma Y^T Q^T$$

$H_t$ : Hessenberg matrix

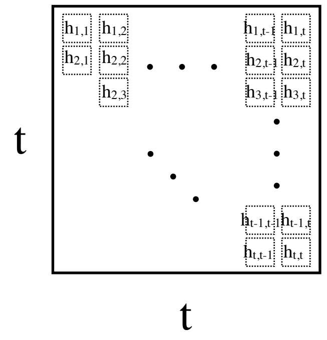


Ref1 : Zhaojun Bai, Templates for the Solution of Algebraic Eigenvalue Problems, 1987
Ref2 : R. B. Lehoucq, et. al., ARPACK User's GUide, 1997

### Implicitly Restarted Arnoldi Method (IRAM)

**Algorithm** Implicitly Restarted Arnoldi Method (IRAM)

**Parameter:** Number of epoch S, Number of dimension K

**Input:** Data covariance matrix $C = X^T X / N \in R^{N \times N}$

Initial vector $q_1 \in R^{N \times 1}$, $m = K - L$ $(K > L)$

$[Q_K, H_K] =$ K step Arnoldi(C, K)

**Output:** Eigenvectors $W \in R^{K \times M}$

**for** $l = 1, 2, \ldots, L$ **do**

$[U, S] =$ K-l step QR($H_K$)

$Q_K^+ = Q_K U, H_K^+ = U^T H_K U$

$\beta_l = \left(H_K^+\right)_{l+1,l}, \sigma_l = U_{Kl}$

$q_{l+1}^+ = q_{l+1}\beta_l + h_{K+1,K} q_{K+1} \sigma_l$

$h_{l+1,l} = \| q_{l+1}^+ \|, q_{l+1}^+ = q_{l+1}^+ / h_{l+1,l}$

$Q_l^+ = Q_K^+ [:,1:l] \quad H_l^+ = H_K^+ [1:l,1:l]$

$[Q_l, H_l] =$ K-l step Arnoldi(C, $Q_l^+$, $H_l^+$, K)

**end for**

$W = Q_L, \Sigma = H_L$

$$C = W \Sigma W^T$$

$$C = Q H Q^T$$

$$C = Q Y \Sigma Y^T Q^T$$

$H_t$ : Hessenberg matrix

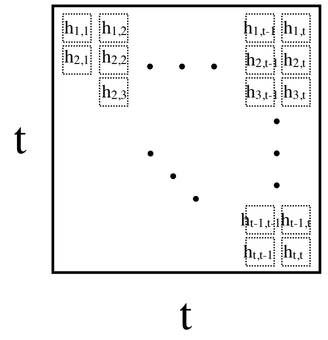

Ref : R. B. Lehoucq, Implicitly Restarted Arnoldi Methods and Subspace Iteration, SIAM J. Matrix Anal. Appl., 2001

### Lanczos method (Lanczos process)

**Algorithm** Lanczos method (Lanczos process)

**Parameter:** Number of epoch S, Number of dimension K

**Input:** Data covariance matrix $C = X^T X/N \in R^{N\times N}$

Initial values $b_0 = 0$, Initial vectors $q_{-1} = 0$, $q_1 \in R^{N\times 1}$

**Output:** Eigenvectors $W \in R^{K\times M}$

**for** t = 1, 2, …, S **do**

$v = Cq_t$ t-th vector generation

$\alpha_t = w_t^T v$

$v = v - \beta_{t-1} q_{t-1} - \alpha_t q_t$ Orthogonalization to t-1and t-th vectors

$\beta_t = \|v\|$

$q_{t+1} = v/\beta_t$ Normalization

**end for**

$[Y, \Sigma, Y^T] = \mathrm{evd}(B_t)$ $\Sigma$ Ritz values $Q$ Ritz vectors

$W = Q_t Y$

$$C = W\Sigma W^T$$
$$= QBQ^T$$
$$= QYBY^TQ^T$$

$B_t$ : Tridiagonal Matrix

t
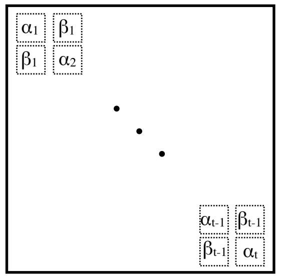
t

Ref : Zhaojun Bai, Templates for the Solution of Algebraic Eigenvalue Problems, 1987

### Augmented implicitly restarted Lanczos bidiagonalization algorithm (IRLBA)

**Algorithm** Augmented implicitly restarted Lanczos bidiagonalization algorithm (IRLBA)

**Parameter:** Number of epoch S, Number of dimension K, Tolerance parameter σ

**Input:** Data matrix $X \in R^{M\times N}$, Initial unit vectors $p_1 \in R^{N\times 1}$, m=min(K+20, 3*K, N)

**Output:** Eigenvectors $W \in R^{K\times M}$

$P_{:1} = p_1, q_1 = Xp_1, \alpha_1 = \|q_1\|, q_1 = q_1/\alpha_1, Q_{:1} = q_1$

**for** t = 1, 2, …, S **do**

$r_t = X^T q_t - \alpha_t p_t$ Orthogonalization to t-1and t-th vectors

$r_t = r_t - w_t \left(w_t^T r_t\right)$ Reorthogonalization

**if** t < m

$\beta_t = \|r_t\|, p_{t+1} = r_t/\beta_t, P_{t+1} = [P_{:t}, p_{t+1}]$

$q_{t+1} = Xp_{t+1} - \beta_t q_t$

$q_{t+1} = q_{t+1} - Q_{:t}\left(Q_{:t}^T q_{t+1}\right)$ Reorthogonalization

$\alpha_{t+1} = \|q_{t+1}\|, q_{t+1} = q_{t+1}/\alpha_{t+1}, Q_{:t+1} = [Q_t, q_{t+1}]$

**end if**

$[Z, \Sigma, Y^T] = \text{svd}(B_t)$

conv = # $\beta_t\Sigma$[m,1:K] < tol*max(Σ)

**if** conv < K

k = min(max(conv + K, k), m - 3)

**end if**

$P_{:,1:k} = P_{:,1:m}Y, B = \Sigma_{1:k,1:k}, Q_{:,1:k} = Q_{:,1:m}Z$ Implicitly restart

**end for**

$W = P, \Sigma = B, V = Q$

$$X = V\Sigma W^T$$

$$X = QBP^T$$

$$XP = QB$$

$$X^T Q = PB^T$$

$$X = QZ\Sigma Y^T P^T$$

$\Sigma$ Ritz values

$P$ Ritz vectors

$B_t$ : Bidiagonal Matrix

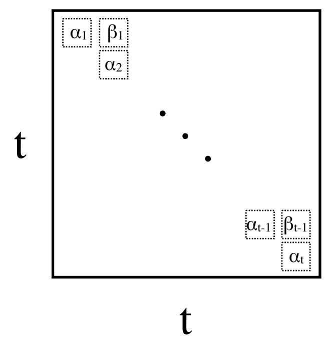


Ref : J. Baglama, et. al., Augmented Implicitly Restarted Lanczos Bidiagonalization Methods, SIAM J. Sci. Comput. 2005
https://github.com/10XGenomics/cellranger/blob/e5396c6c444acec6af84caa7d3655dd33a162852/lib/python/cellranger/analysis/irlb.py

### GD-PCA

**Algorithm** GD-PCA

**Parameters:** Step size η, Number of epoch S, Number of Dimension K

**Input:** Data matrix $X = (x_1, \ldots, x_N) \in R^{M \times N}$

Initial Matrix $W_0 = (w_1, \ldots, w_K)$ (diagonal element is 1, otherwise 0) $\in R^{M \times K}$

Diagonal matrix $D = \mathrm{Diag}(K, \ldots, 1) / 10^5 \in R^{K \times K}$

**Output:** Eigenvectors $W \in R^{K \times M}$

**for** t = 1, 2, …, S **do**

$$W_t' = W_{t-1} + \frac{\eta}{N} \sum_{i=1}^{N} x_i \left( x_i^T W_{t-1} D \right)$$

$$W_t = \mathrm{qr}\left(\mathrm{W}_\mathrm{t}'\right)$$

**end for**

### SGD-PCA

**Algorithm** SGD-PCA (Oja)

**Parameters:** Step size η, Number of epoch S, Number of Dimension K

**Input:** Data matrix $X = (x_1, \ldots, x_N) \in R^{M \times N}$

Initial Matrix $W_0 = (w_1, \ldots, w_K)$ (diagonal element is 1, otherwise 0) $\in R^{M \times K}$

Diagonal matrix $D = \mathrm{Diag}(K, \ldots, 1) / 10^5 \in R^{K \times K}$

**Output:** Eigenvectors $W \in R^{K \times M}$

**for** t = 1, 2, …, S **do**

**for** i = 1, 2, …, N **do**

$$W'_{t,i} = W_{t-1,i} + \frac{\eta}{N} x_i \left( x_i^T W_{t-1,i} D \right)$$

$$W_{t,i} = \mathrm{qr}\left( \mathrm{W}'_{\mathrm{t,i}} \right)$$

**end for**

**end for**

Ref1 : Erkki Oja, et. al., On stochastic approximation of the eigenvectors and eigenvalues of the expectation of a random matrix, Journal of Mathematical Analysis and Applications, 1985

Ref2 : Erkki Oja, Principal components, minor components, and linear neural networks, Neural Networks, 1992

### Halko's method

**Algorithm** Halko's method

**Parameters:** Number of Dimension K, Number of over-sampling L, Number of iterations *its*

**Input:** Data matrix $X = (x_1, \ldots, x_N) \in R^{M\times N}$, Random Matrix $\Omega \in R^{N\times L}$

**Output:** Eigenvectors $W \in R^{K\times M}$

$$Y_0 = X\Omega$$

$$Q_0 = \mathrm{qr}\left(Y_o\right)$$

**for** t = 1 to *its*

$$Y_t = \mathrm{qr}\left(X\mathrm{qr}\left(X^T Q_{t-1}\right)\right)$$

**end for**

$$Q = \mathrm{qr}\left(Y_{its}\right)$$

$$U\Sigma V^T = \mathrm{svd}\left(Q^T X\right)$$

$$W = QU$$

Ref1 : Nathan Halko, et. al., Finding structure with randomness: Probabilistic algorithms for constructing approximate matrix decompositions, SIAM Rev., 2011

Ref2 : Nathan Halko, et. al., An Algorithm for the Principal Component Analysis of Large Data Sets, SIAM J. Sci. Comput., 2011

### Algorithm971

**Algorithm** Algorithm971

**Parameters:** Number of Dimension K, Number of over-sampling L, Number of iterations *its*

**Input:** Data matrix $X = (x_1, \ldots, x_N) \in R^{M \times N}$, Random Matrix $\Omega \in R^{N \times L}$

**Output:** Eigenvectors $W \in R^{K \times M}$

$Y_0 = X\Omega$

$L_0 = \mathrm{lu}(Y_o)$

**for** t = 1 to *its*

$Y_t = XX^T L_{t-1}$

**if** t < its

$L_t = \mathrm{lu}(Y_t)$

**endif**

**end for**

$Q = \mathrm{qr}(Y_{its})$

$U\Sigma V^T = \mathrm{svd}\left(Q^T X\right)$

$W = QU$

Ref1 : Huamin Li, et. al., Algorithm 971 : An Implementation of a Randomized Algorithm for Principal Component Analysis, ACM Trans Math Softw., 2017

Ref2 : George C. Linderman, Fast interpolation-based t-SNE for improved visualization of single-cell RNA-seq dta, Nature Methods, 2019
