## Additional File 3 for "Benchmarking principal component analysis for large-scale single-cell RNA-sequencing"

### Source code of all the PCA algorithms

### normalize (cpmed, centring)

```
normalize <- function(out){
    libsize <- colSums(out)
    out <- median(libsize) * t(t(out) / libsize)
    out <- log10(out + 1)
    m <- rowMeans(out)
    sweep(out, 1, m)
}
```

R : 3.5.0

### prcomp

```
args = commandArgs()
infile = args[1] # Input file
outfile1 = args[2] # Output file (Eigenvectors)
outfile2 = args[3] # Output file (Eigenvalues)
dim = args[4] # Dimension

input = read.csv(infile, header=FALSE)
input = normalize(input)

out = prcomp(input, center=FALSE, scale=FALSE)

write.table(t(out$rotation[,1:dim]), outfile1)
write.table(t(out$sdev[1:dim]^2), outfile2)
```

R : 3.5.0

### Sklearn (LAPACK, v0.20.2)

```
# -*- coding: utf-8 -*-

import sys
import numpy as np
from sklearn.decomposition import PCA

args = sys.argv
infile = args[1] # Input file
outfile1 = args[2] # Output file (Eigenvectors)
outfile2 = args[3] # Output file (Eigenvalues)
dim = int(args[4]) # Dimension

data = np.loadtxt(infile, delimiter=",")
libsize = np.sum(data, axis=0)
med = np.median(np.asarray(libsize))
data = med * (data / libsize)
data = np.log10(data + 1)

pca = PCA(n_components=dim, svd_solver = "full")
pca.fit(data.T)
out = pca.fit_transform(data.T)

np.savetxt(outfile1, out, delimiter=",")
np.savetxt(outfile2, pca.explained_variance_, delimiter=",")
```

Python : 3.6.4

### Multivariate.jl (fit, v0.6.0)

```
using DelimitedFiles
using MultivariateStats
import Statistics: mean

input = ARGS[1]
output1 = ARGS[2]
output2 = ARGS[3]
dim = parse(Int64, ARGS[4])

out = readcsv(input)
libsize = sum(out, dims=1)
out = median(libsize) .* (out ./ libsize)
out = log10.(out .+ 1)

mval = vec(mean(out, dims=2))
res = MultivariateStats.fit(PCA, out, mean=mval, maxoutdim=dim)

writedlm(output1, MultivariateStats.transform(res, out)', ',')
writedlm(output2, principalvars(res), ',')
```

Julia : 1.0.1

### Downsampling

```
# After Downsampling of data matrix by in-house Julia script

args = commandArgs()
infile = args[1] # Down sampled file
outfile1 = args[2] # Output file (Eigenvectors)
outfile2 = args[3] # Output file (Eigenvalues)
dim = args[4] # Dimension

input = read.csv(infile, header=FALSE)
input = normalize(input)

U = svd(input, scale=FALSE)

# V = X'U with incremental row vector loading from X' by in-
house Julia script
```

R : 3.5.0

### Sklearn (Incremental, v0.20.2)

```
# -*- coding: utf-8 -*-

import sys
import numpy as np
from sklearn.decomposition import IncrementalPCA
from sklearn import preprocessing
import itertools

args = sys.argv
infile = args[1] # Input file (CPMED)
outfile1 = args[2] # Output file (Eigenvectors)
outfile2 = args[3] # Output file (Eigenvalues)
dim = int(args[4]) # Dimension

pca = IncrementalPCA(n_components=dim)
i = 1
start = 0
end = dim
eof = False

while eof == False:
  with open(infile) as f:
    start = dim * i
    end = start + dim
    data = np.loadtxt(itertools.islice(f, start, end), delimiter=",")
    if data.shape[0] < dim:
      eof = True
    else:
      data = preprocessing.scale(np.log10(data + 1), axis=1, with_mean=True, with_std=False)
      pca.partial_fit(data)
      i = i + 1

np.savetxt(outfile1, pca.components_.T, delimiter=",")
np.savetxt(outfile2, pca.explained_variance_, delimiter=",")
```

Python : 3.6.4

### irlba (irlba, v2.3.3)

```
library("irlba")

args = commandArgs()
infile = args[1] # Input file
outfile1 = args[2] # Output file (Eigenvectors)
outfile2 = args[3] # Output file (Eigenvalues)
dim = args[4] # Dimension

input = read.csv(infile, header=FALSE)
input = normalize(input)

out = irlba(input, dim)

write.table(t(out$v[,1:dim]), outfile1)
write.table(t(out$d[1:dim]^2/ncol(input)), outfile2)
```

R : 3.5.0

### rARPACK (svds, v0.11-0)

```
library("rARPACK")

args = commandArgs()
infile = args[1] # Input file
outfile1 = args[2] # Output file (Eigenvectors)
outfile2 = args[3] # Output file (Eigenvalues)
dim = args[4] # Dimension

input = read.csv(infile, header=FALSE)
input = normalize(input)

out = rARPACK::svds(input, dim)

write.table(t(out$v[,1:dim]), outfile1)
write.table(t(out$d[1:dim]^2/ncol(input)), outfile2)
```

R : 3.5.0

### RSpectra (svds, v0.14-0)

```
library("RSpectra")

args = commandArgs()
infile = args[1] # Input file
outfile1 = args[2] # Output file (Eigenvectors)
outfile2 = args[3] # Output file (Eigenvalues)
dim = args[4] # Dimension

input = read.csv(infile, header=FALSE)
input = normalize(input)

out = RSpectra::svds(input, dim)

write.table(t(out$v[,1:dim]), outfile1)
write.table(t(out$d[1:dim]^2/ncol(input)), outfile2)
```

R : 3.5.0

### svd (propack.svd, v0.4.3)

```
library("svd")

args = commandArgs()
infile = args[1] # Input file
outfile1 = args[2] # Output file (Eigenvectors)
outfile2 = args[3] # Output file (Eigenvalues)
dim = args[4] # Dimension

input = read.csv(infile, header=FALSE)
input = normalize(input)

out = propack.svd(input, dim)

write.table(t(out$v[,1:dim]), outfile1)
write.table(t(out$d[1:dim]^2/ncol(input)), outfile2)
```

R : 3.5.0

### Sklearn (ARPACK, v0.20.2)

```
# -*- coding: utf-8 -*-

import sys
import numpy as np
from sklearn.decomposition import PCA

args = sys.argv
infile = args[1] # Input file
outfile1 = args[2] # Output file (Eigenvectors)
outfile2 = args[3] # Output file (Eigenvalues)
dim = int(args[4]) # Dimension

data = np.loadtxt(infile, delimiter=",")
libsize = np.sum(data, axis=0)
med = np.median(np.asarray(libsize))
data = med * (data / libsize)
data = np.log10(data + 1)

pca = PCA(n_components=dim, svd_solver = "arpack")
pca.fit(data.T)
out = pca.fit_transform(data.T)

np.savetxt(outfile1, out, delimiter=",")
np.savetxt(outfile2, pca.explained_variance_, delimiter=",")
```

Python : 3.6.4

### Cell Ranger (irlb, v3.0.2)

```
# -*- coding: utf-8 -*-

import sys
import numpy as np
import scipy.sparse as sp
import warnings
import array
import csv
from scipy.sparse import csr_matrix

# copy & paste the functions below
# https://github.com/10XGenomics/cellranger/blob/master/lib/python/cellranger/
analysis/irlb.py

args = sys.argv
infile = args[1] # Input file (LogCPMED)
outfile1 = args[2] # Output file (Eigenvectors)
outfile2 = args[3] # Output file (Eigenvalues)
dim = int(args[4]) # Dimension

data = np.loadtxt(infile, delimiter=",")
A = sp.csr_matrix(data)

out = irlb(A, n=dim)

np.savetxt(outfile1, out[0], delimiter=",")
np.savetxt(outfile2, out[1], delimiter=",")
```

Python : 3.6.4

### Arpack.jl (svds, v0.3.0)

```
using DelimitedFiles
using Arpack
import Statistics: mean

input = ARGS[1]
output1 = ARGS[2]
output2 = ARGS[3]
dim = parse(Int64, ARGS[4])

out = readdlm(input, ',')
libsize = sum(out, dims=1)
out = median(libsize) .* (out ./ libsize)
out = log10.(out .+ 1)
out = out .- vec(mean(out, dims=2))

res = svds(out, nsv=dim)[1]

writedlm(output1, res.Vt', ',')
writedlm(output2, res.S.*res.S/size(res.Vt)[2], ',')
```

Julia : 1.0.1

### OnlinePCA.jl (orthiter, v0.1.0)

```
using OnlinePCA: csv2bin, sumr, orthiter, readcsv, writecsv

wdir = ARGS[1] # Working directory
dim = parse(Int64, ARGS[2]) # Number of dimensions

infile1 = joinpath(wdir, "Data.csv")
infile2 = joinpath(wdir, "Feature_Logmeans.csv")
infile3 = joinpath(wdir, "Sample_NoCounts.csv")
outfile1 = joinpath(wdir, "Data.zst")
outfile2 = joinpath(wdir, "Eigen_vectors.csv")
outfile3 = joinpath(wdir, "Eigen_values.csv")

csv2bin(csvfile=input1, binfile=outfile1)

sumr(binfile=outfile1, outdir= wdir)

libsize = readcsv(infile3)
out = orthiter(input=infile1, dim=dim, rowmeanlist=infile2,
  colsumlist=infile3, cper=median(libsize))

writecsv(output2, out[1])
writecsv(output3, out[2])
```

Julia : 1.0.1

### OnlinePCA.jl (gd, v0.1.0)

```
using OnlinePCA: csv2bin, sumr, orthiter, readcsv, writecsv

wdir = ARGS[1] # Working directory
dim = parse(Int64, ARGS[2]) # Number of dimensions

infile1 = joinpath(wdir, "Data.csv")
infile2 = joinpath(wdir, "Feature_Logmeans.csv")
infile3 = joinpath(wdir, "Sample_NoCounts.csv")
outfile1 = joinpath(wdir, "Data.zst")
outfile2 = joinpath(wdir, "Eigen_vectors.csv")
outfile3 = joinpath(wdir, "Eigen_values.csv")

csv2bin(csvfile=input1, binfile=outfile1)

sumr(binfile=outfile1, outdir= wdir)

libsize = readcsv(infile3)
out = gd(input=infile1, dim=dim, rowmeanlist=infile2,
  colsumlist=infile3, cper=median(libsize))

writecsv(output2, out[1])
writecsv(output3, out[2])
```

Julia : 1.0.1

### OnlinePCA.jl (sgd, v0.1.0)

```
using OnlinePCA: csv2bin, sumr, orthiter, readcsv, writecsv

wdir = ARGS[1] # Working directory
dim = parse(Int64, ARGS[2]) # Number of dimensions

infile1 = joinpath(wdir, "Data.csv")
infile2 = joinpath(wdir, "Feature_Logmeans.csv")
infile3 = joinpath(wdir, "Sample_NoCounts.csv")
outfile1 = joinpath(wdir, "Data.zst")
outfile2 = joinpath(wdir, "Eigen_vectors.csv")
outfile3 = joinpath(wdir, "Eigen_values.csv")

csv2bin(csvfile=input1, binfile=outfile1)

sumr(binfile=outfile1, outdir= wdir)

libsize = readcsv(infile3)
out = sgd(input=infile1, dim=dim, rowmeanlist=infile2,
  colsumlist=infile3, cper=median(libsize))

writecsv(output2, out[1])
writecsv(output3, out[2])
```

Julia : 1.0.1

### rsvd (rsvd, v1.0.0)

```
library("rsvd")

args = commandArgs()
infile = args[1] # Input file
outfile1 = args[2] # Output file (Eigenvectors)
outfile2 = args[3] # Output file (Eigenvalues)
dim = args[4] # Dimension

input = read.csv(infile, header=FALSE)
input = normalize(input)

out = rsvd(input, dim, q=3)

write.table(t(out$v[,1:dim]), outfile1)
write.table(t(out$d[1:dim]^2/ncol(input)), outfile2)
```

R : 3.5.0

### oocRPCA (oocPCA_CSV, v0.0.0.900)

```
library("oocRPCA")

args = commandArgs()
infile = args[1] # Input file (CPMED)
outfile1 = args[2] # Output file (Eigenvectors)
outfile2 = args[3] # Output file (Eigenvalues)
dim = args[4] # Dimension

out = oocPCA_CSV(infile, k=dim, mem=1e+10, its=3,
  centeringRow=TRUE, logTransform=TRUE)

write.table(t(out$v[,1:dim]), outfile1)
write.table(t(out$d[1:dim]^2/ncol(input)), outfile2)
```

R : 3.5.0

### Sklearn (Randomized, v0.20.2)

```
# -*- coding: utf-8 -*-

import sys
import numpy as np
from sklearn.decomposition import PCA

args = sys.argv
infile = args[1] # Input file
outfile1 = args[2] # Output file (Eigenvectors)
outfile2 = args[3] # Output file (Eigenvalues)
dim = int(args[4]) # Dimension

data = np.loadtxt(infile, delimiter=",")
libsize = np.sum(data, axis=0)
med = np.median(np.asarray(libsize))
data = med * (data / libsize)
data = np.log10(data + 1)

pca = PCA(n_components=dim, svd_solver = "randomized")
pca.fit(data.T)
out = pca.fit_transform(data.T)

np.savetxt(outfile1, out, delimiter=",")
np.savetxt(outfile2, pca.explained_variance_, delimiter=",")
```

Python : 3.6.4

### Sklearn (randomized_svd, v0.20.2)

```
# -*- coding: utf-8 -*-

import sys
import numpy as np
from sklearn.utils.extmath import randomized_svd
from sklearn import preprocessing

args = sys.argv
infile = args[1] # Input file
outfile1 = args[2] # Output file (Eigenvectors)
outfile2 = args[3] # Output file (Eigenvalues)
dim = int(args[4]) # Dimension

data = np.loadtxt(infile, delimiter=",")
libsize = np.sum(data, axis=0)
med = np.median(np.asarray(libsize))
data = med * (data / libsize)
data = np.log10(data + 1)
data = preprocessing.scale(data, axis=1,
  with_mean=True,with_std=False)

out = randomized_svd(data, n_components=dim, n_iter=3,
  power_iteration_normalizer="LU")

np.savetxt(outfile1, out[2].T, delimiter=",")
np.savetxt(outfile2, out[1]*out[1]/data.shape[1], delimiter=",")
```

Python : 3.6.4

### dask_ml (PCA, v0.12.0)

```
# -*- coding: utf-8 -*-

import sys
import numpy as np
from dask_ml.decomposition import PCA
import dask.dataframe as dd

args = sys.argv
infile = args[1] # Input file (CPMED)
outfile1 = args[2] # Output file (Eigenvectors)
outfile2 = args[3] # Output file (Eigenvalues)
dim = int(args[4]) # Dimension

data = dd.read_csv(infile, sep=",", header=None)
data = data.to_dask_array(True)
data = np.log10(data + 1)

pca = PCA(n_components=dim, svd_solver="randomized",
 iterated_power=3)
pca.fit(data.T)
out = pca.fit_transform(data.T)

np.savetxt(outfile1, out, delimiter=",")
np.savetxt(outfile2, pca.explained_variance_, delimiter=",")
```

Python : 3.6.4

### OnlinePCA.jl (halko, v0.1.0)

```
using OnlinePCA: csv2bin, sumr, orthiter, readcsv, writecsv

wdir = ARGS[1] # Working directory
dim = parse(Int64, ARGS[2]) # Number of dimensions

infile1 = joinpath(wdir, "Data.csv")
infile2 = joinpath(wdir, "Feature_Logmeans.csv")
infile3 = joinpath(wdir, "Sample_NoCounts.csv")
outfile1 = joinpath(wdir, "Data.zst")
outfile2 = joinpath(wdir, "Eigen_vectors.csv")
outfile3 = joinpath(wdir, "Eigen_values.csv")

csv2bin(csvfile=input1, binfile=outfile1)

sumr(binfile=outfile1, outdir= wdir)

libsize = readcsv(infile3)
out = halko(input=infile1, dim=dim, rowmeanlist=infile2,
  colsumlist=infile3, cper=median(libsize), niter=3)

writecsv(output2, out[1])
writecsv(output3, out[2])
```

Julia : 1.0.1

### OnlinePCA.jl (algorithm971, v0.1.0)

```
using OnlinePCA: csv2bin, sumr, orthiter, readcsv, writecsv

wdir = ARGS[1] # Working directory
dim = parse(Int64, ARGS[2]) # Number of dimensions

infile1 = joinpath(wdir, "Data.csv")
infile2 = joinpath(wdir, "Feature_Logmeans.csv")
infile3 = joinpath(wdir, "Sample_NoCounts.csv")
outfile1 = joinpath(wdir, "Data.zst")
outfile2 = joinpath(wdir, "Eigen_vectors.csv")
outfile3 = joinpath(wdir, "Eigen_values.csv")

csv2bin(csvfile=input1, binfile=outfile1)

sumr(binfile=outfile1, outdir= wdir)

libsize = readcsv(infile3)
out = algorithm971(input=infile1, dim=dim, rowmeanlist=infile2,
  colsumlist=infile3, cper=median(libsize), niter=3)

writecsv(output2, out[1])
writecsv(output3, out[2])
```

Julia : 1.0.1
