## Supplementary figures and images for "Benchmarking principal component analysis for large-scale single-cell RNA-sequencing"

### Additional File 4

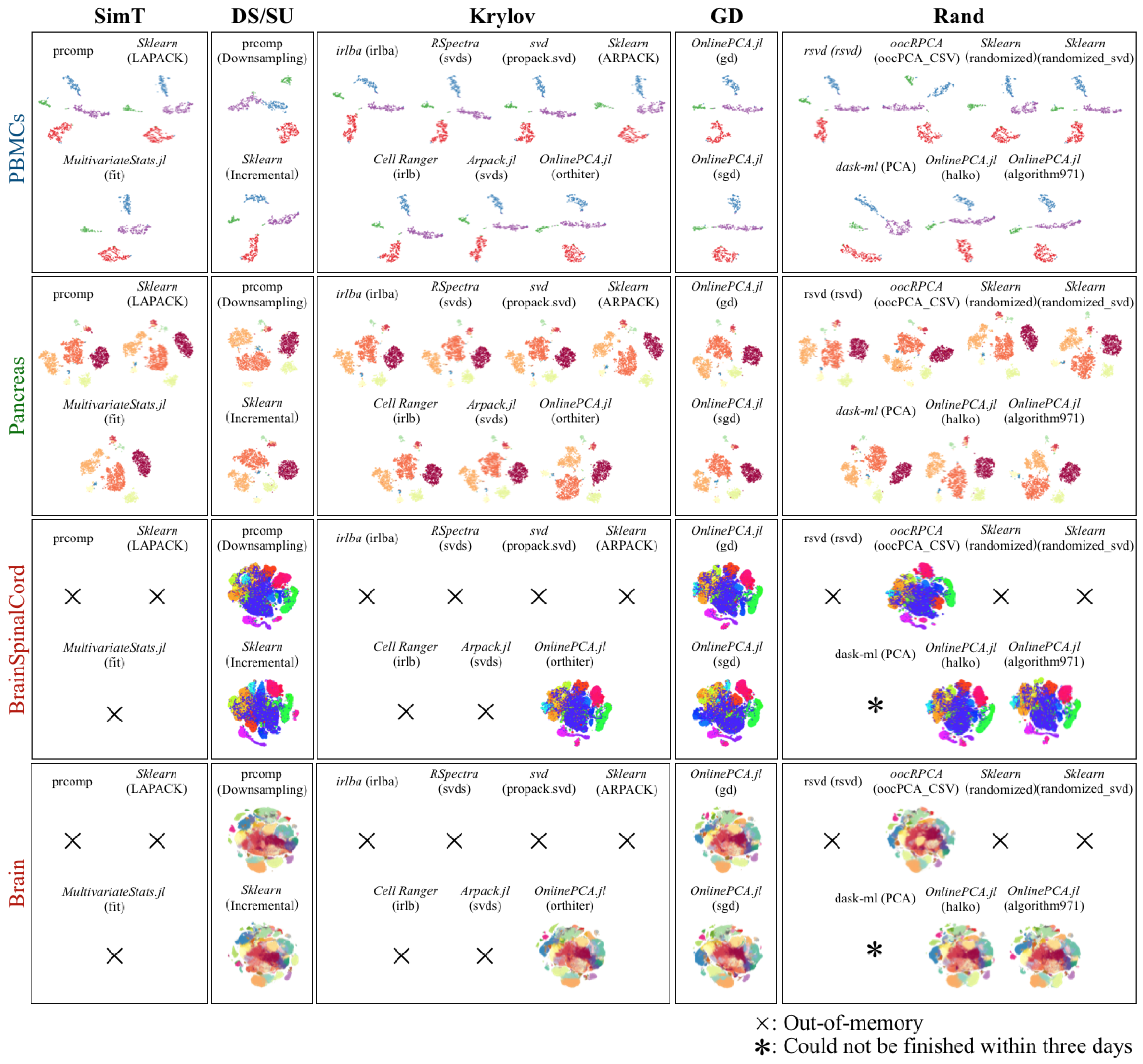

### Additional File 5

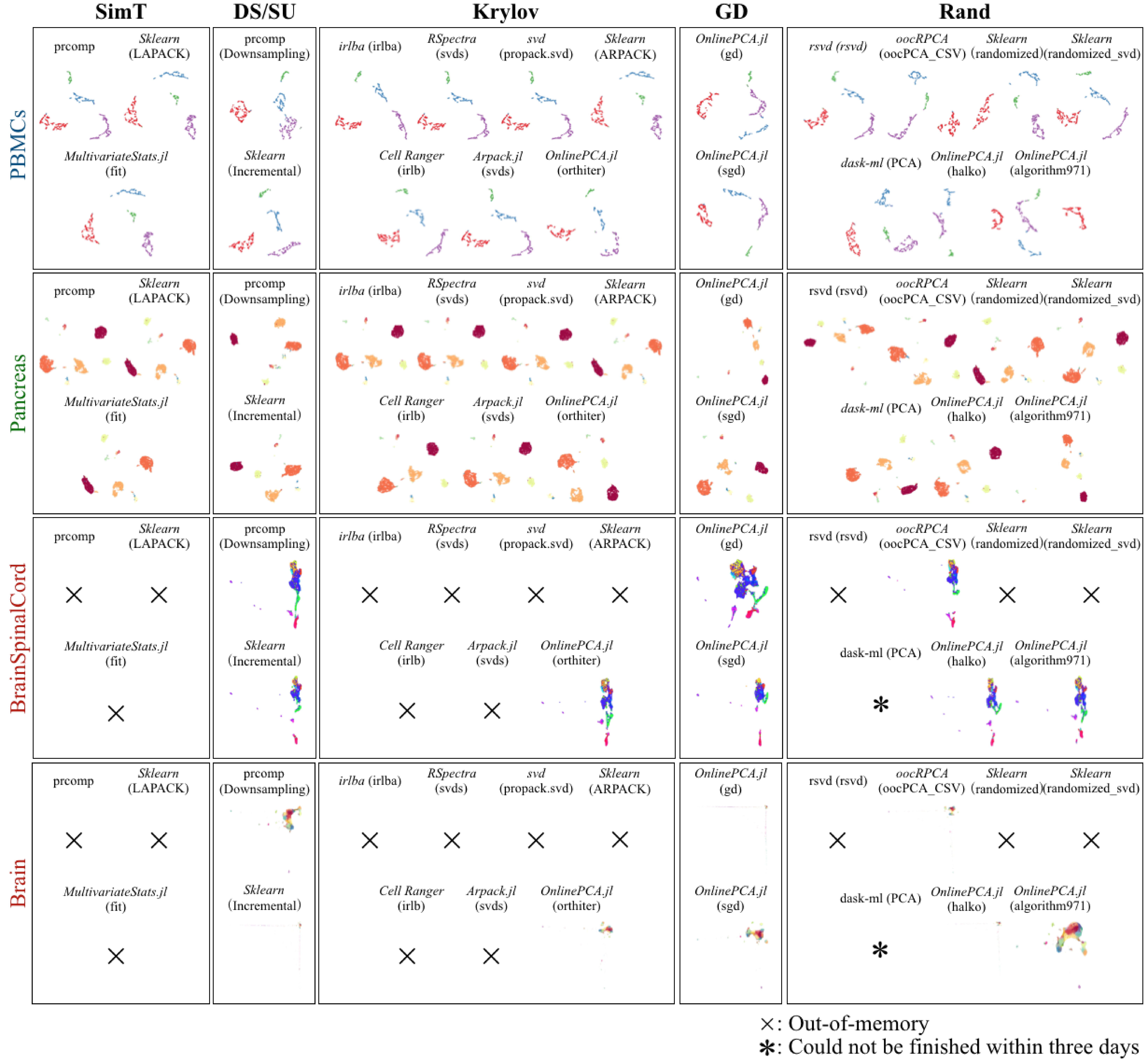

### Additional File 7

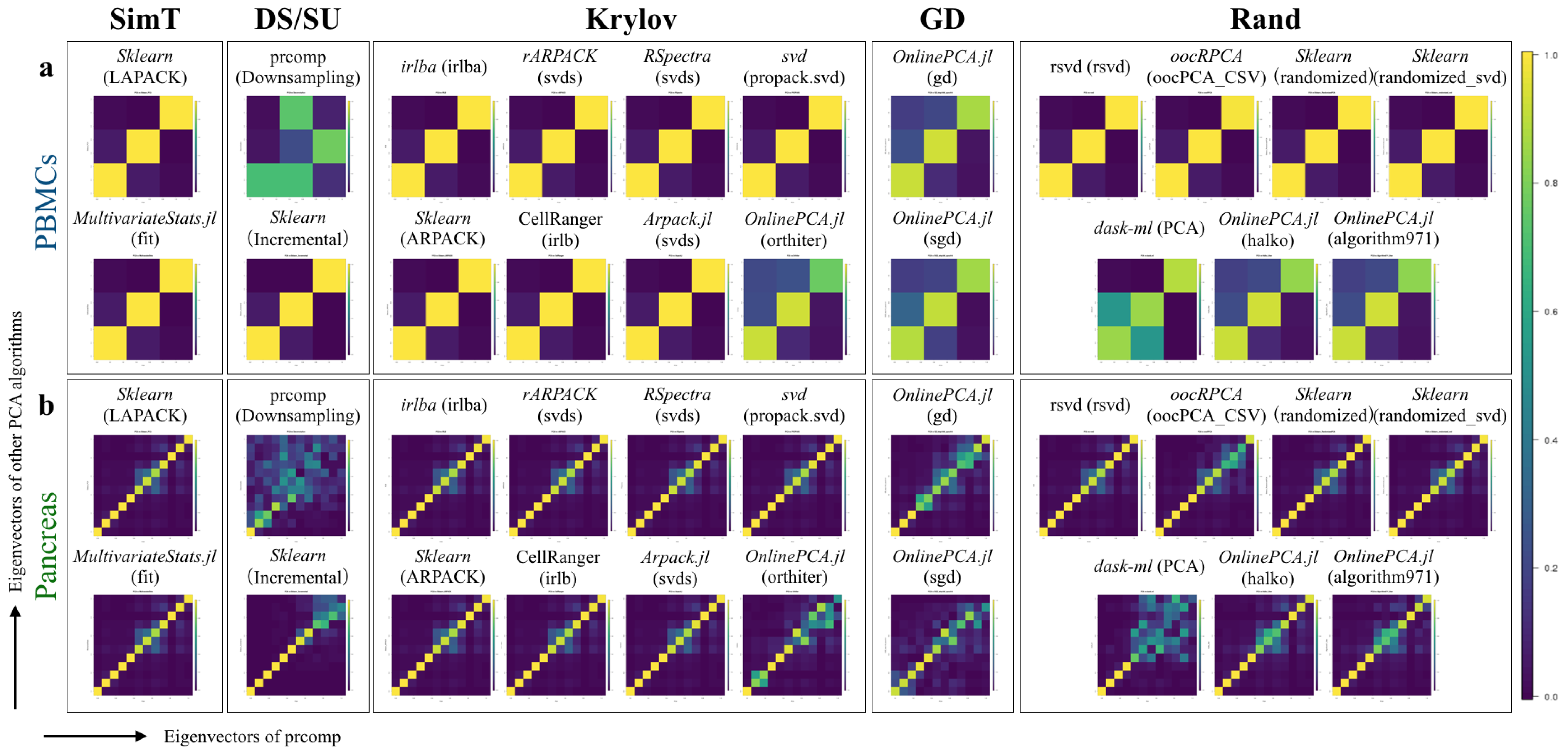

### Additional File 12

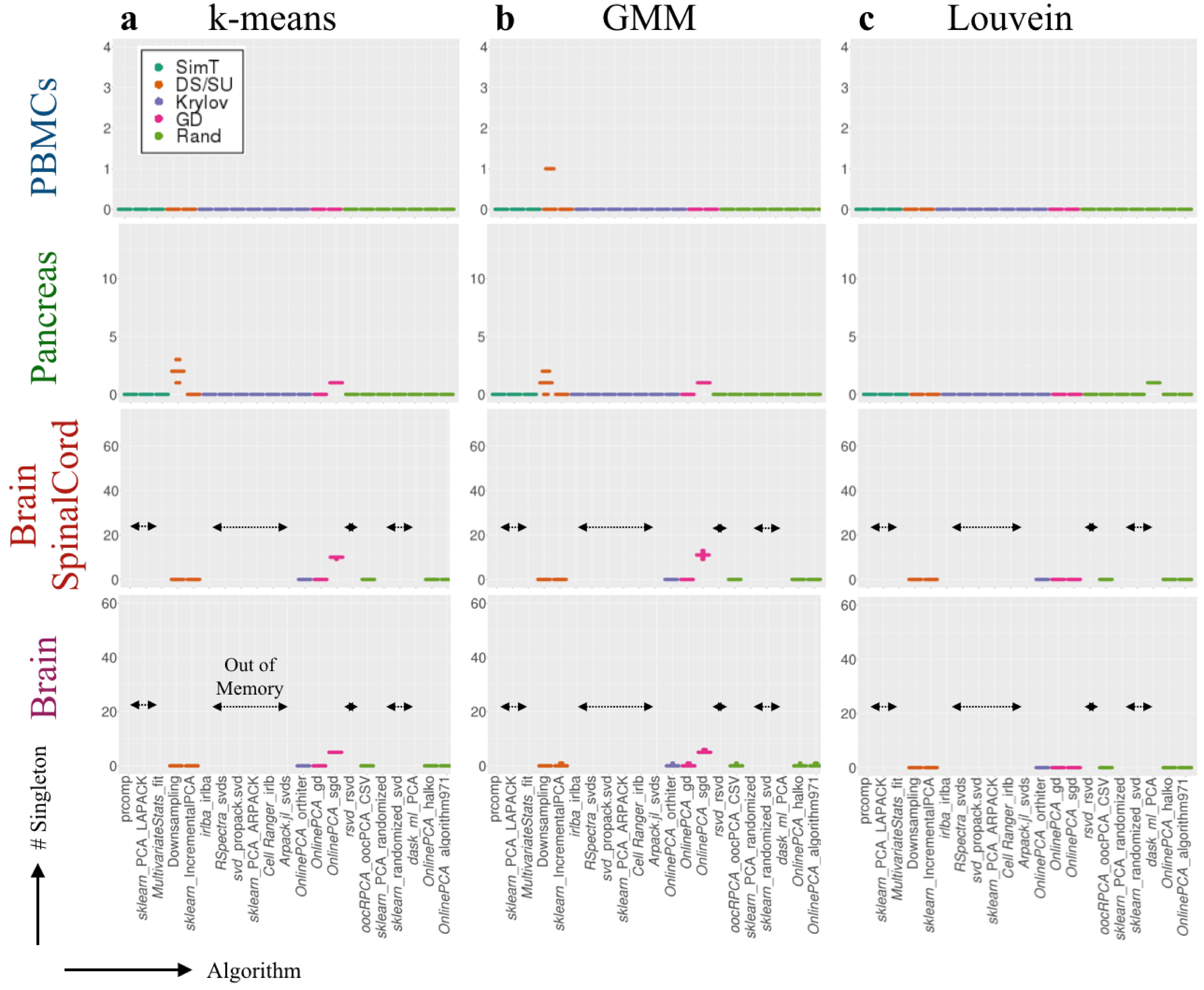

### Additional File 13

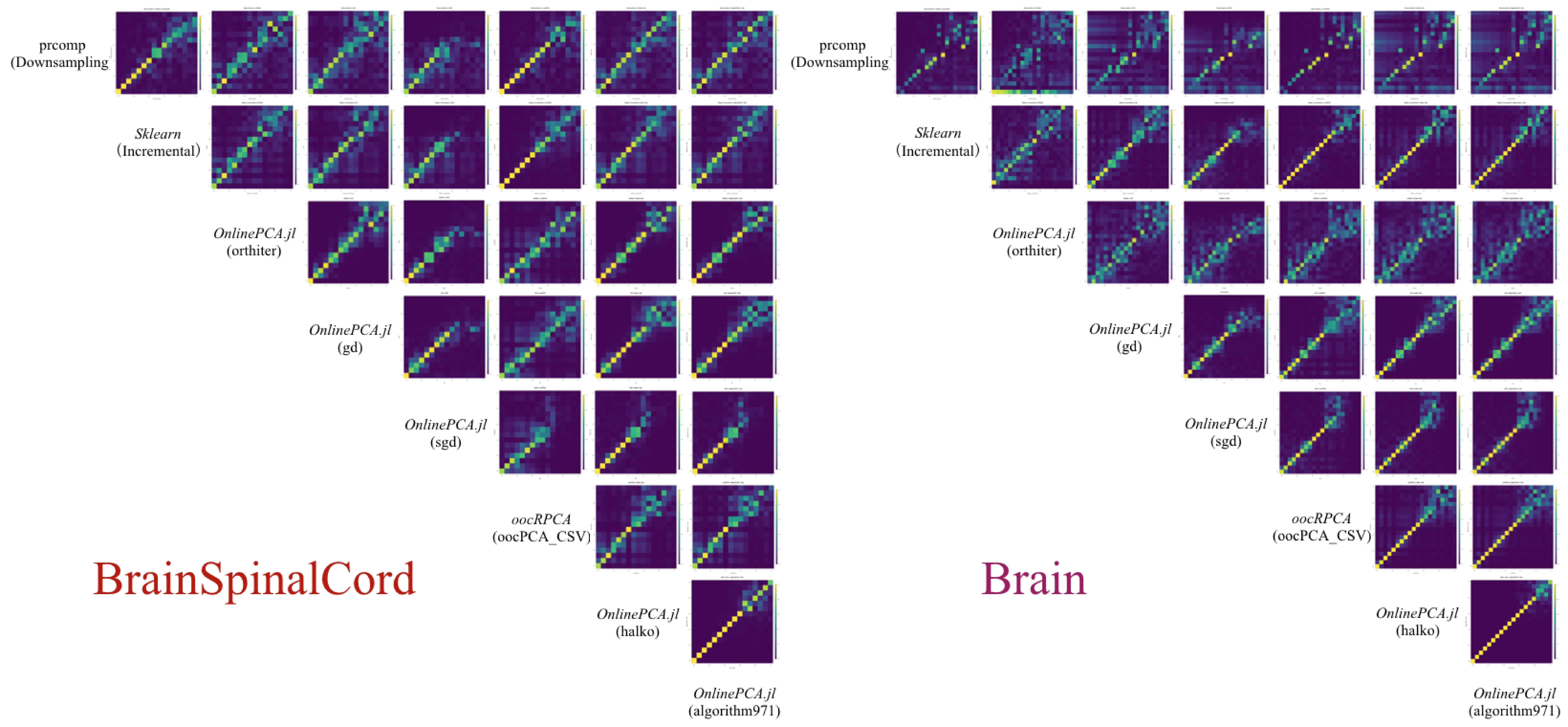

### Additional File 14

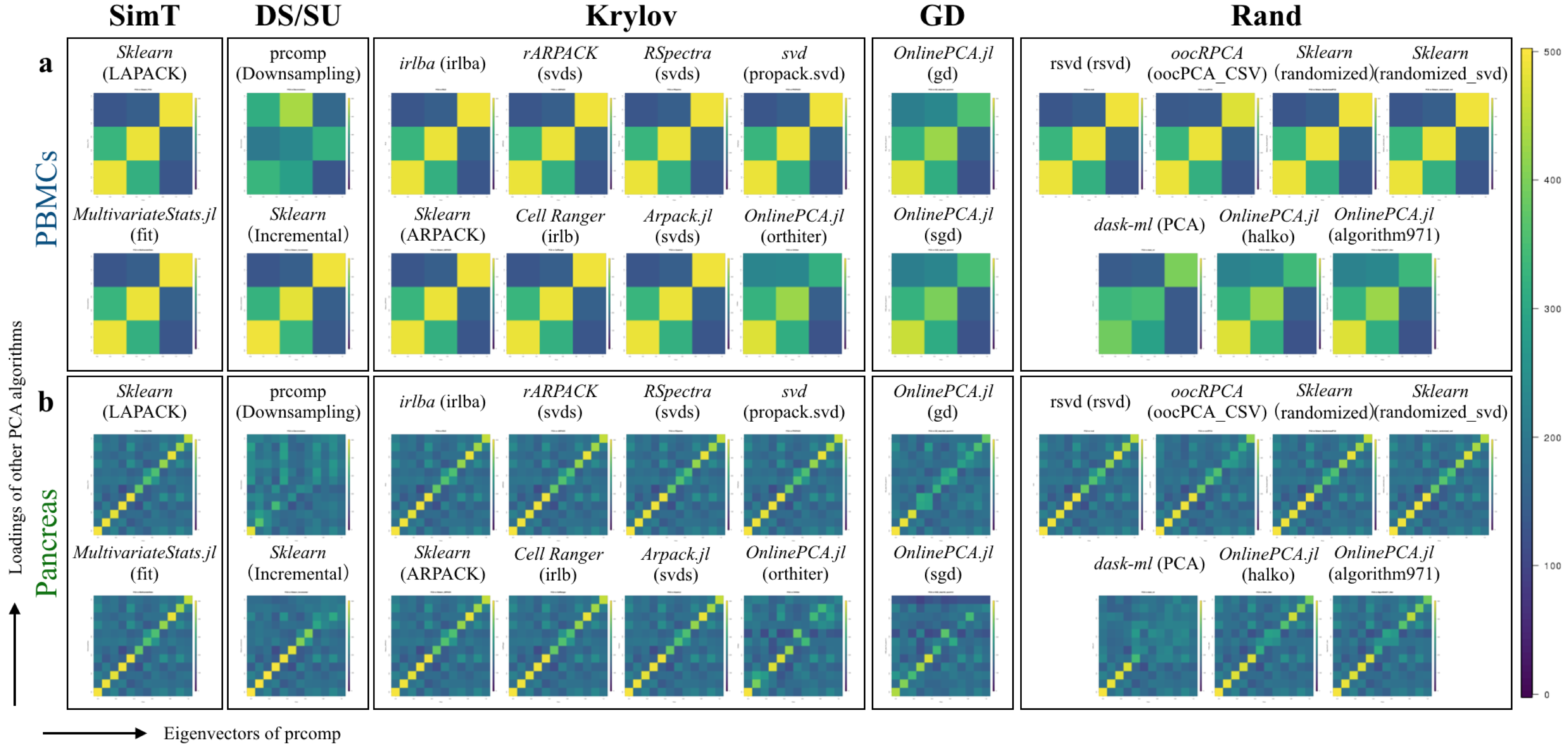

### Algorithm971_3iter.png

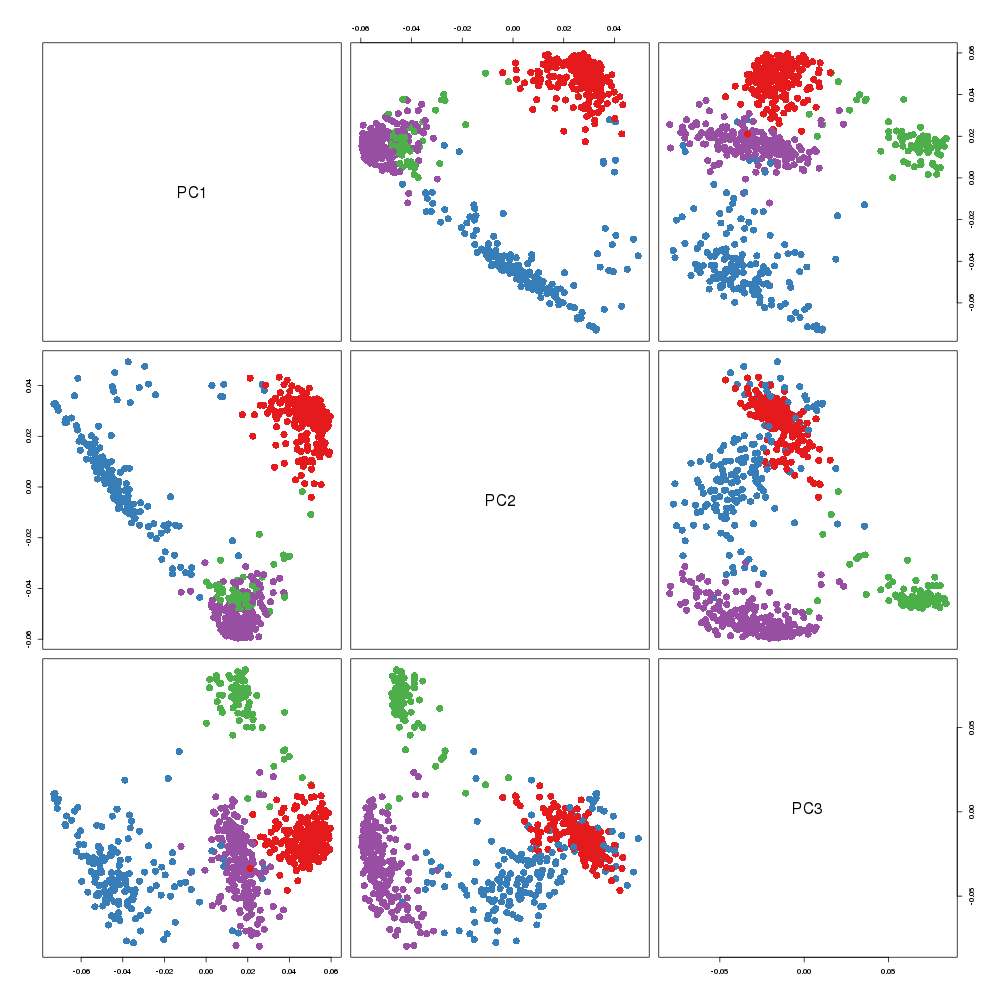

### Algorithm971_3iter.png

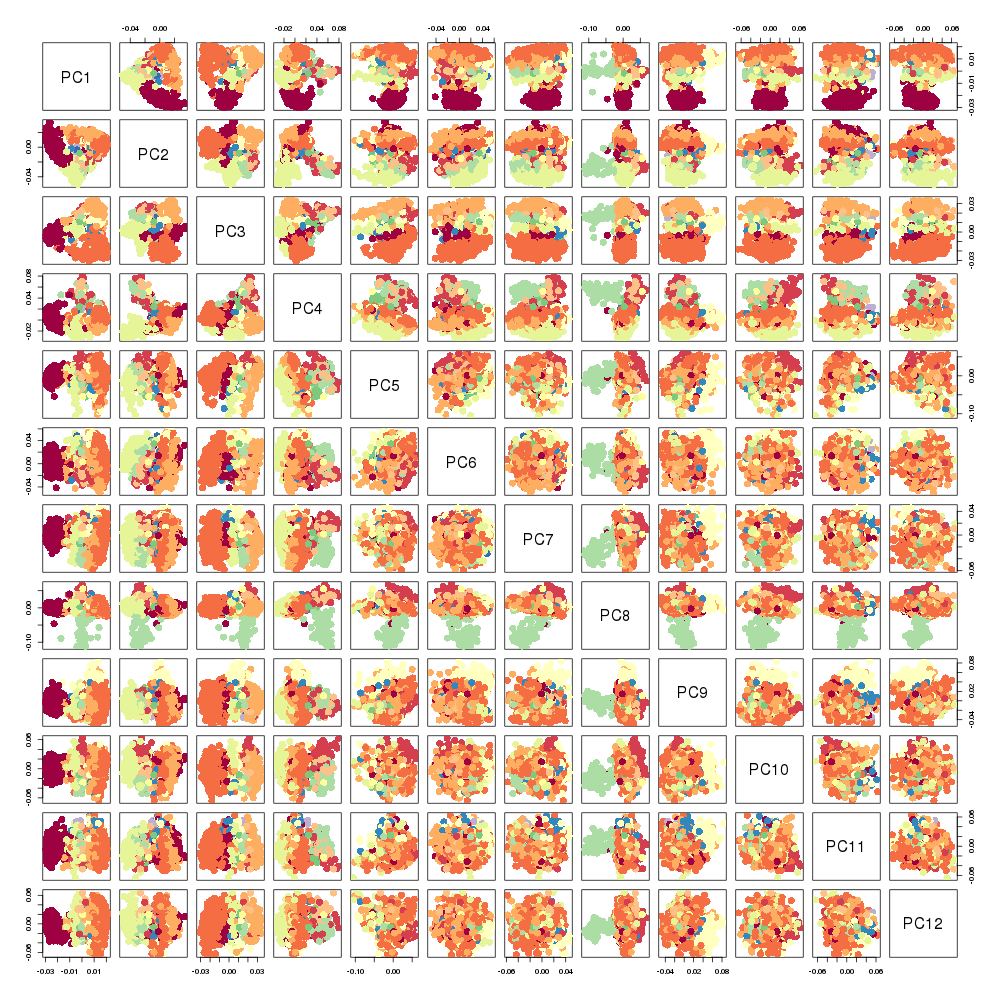

### Algorithm971_3iter.png

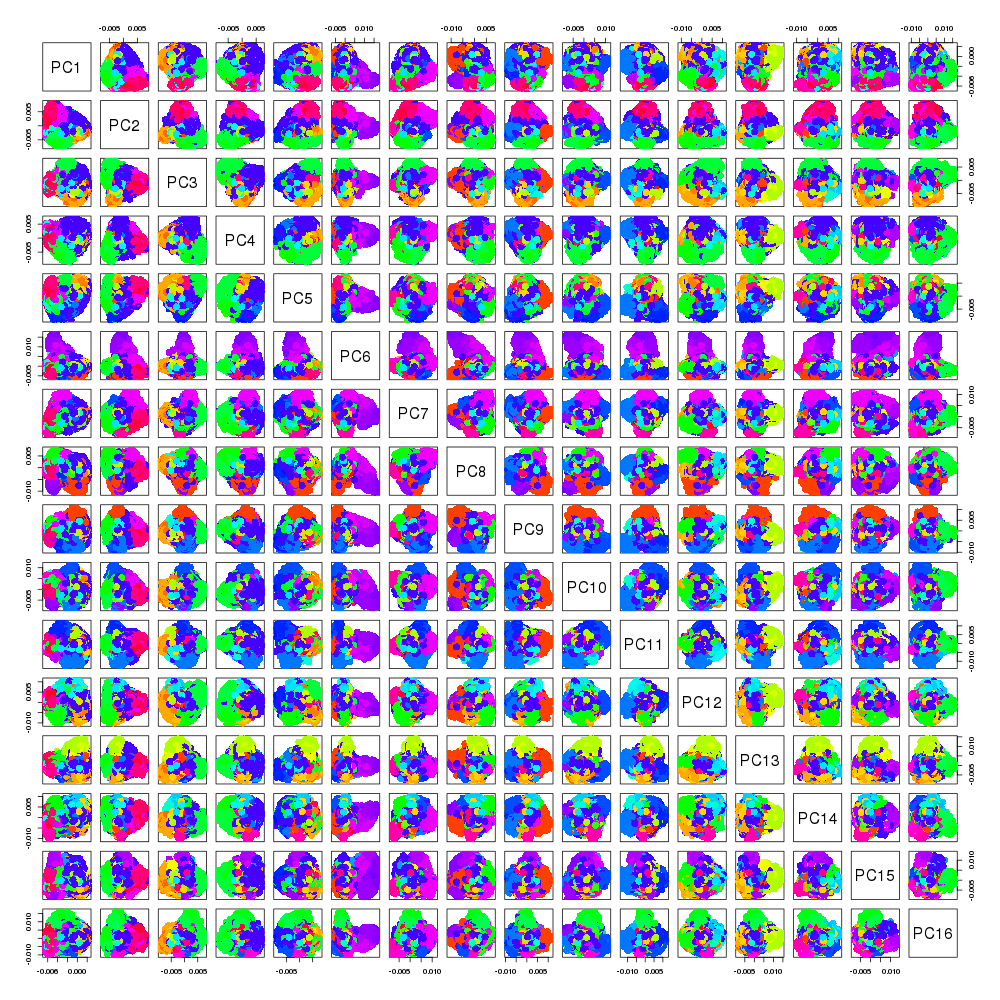

### Algorithm971_3iter.png

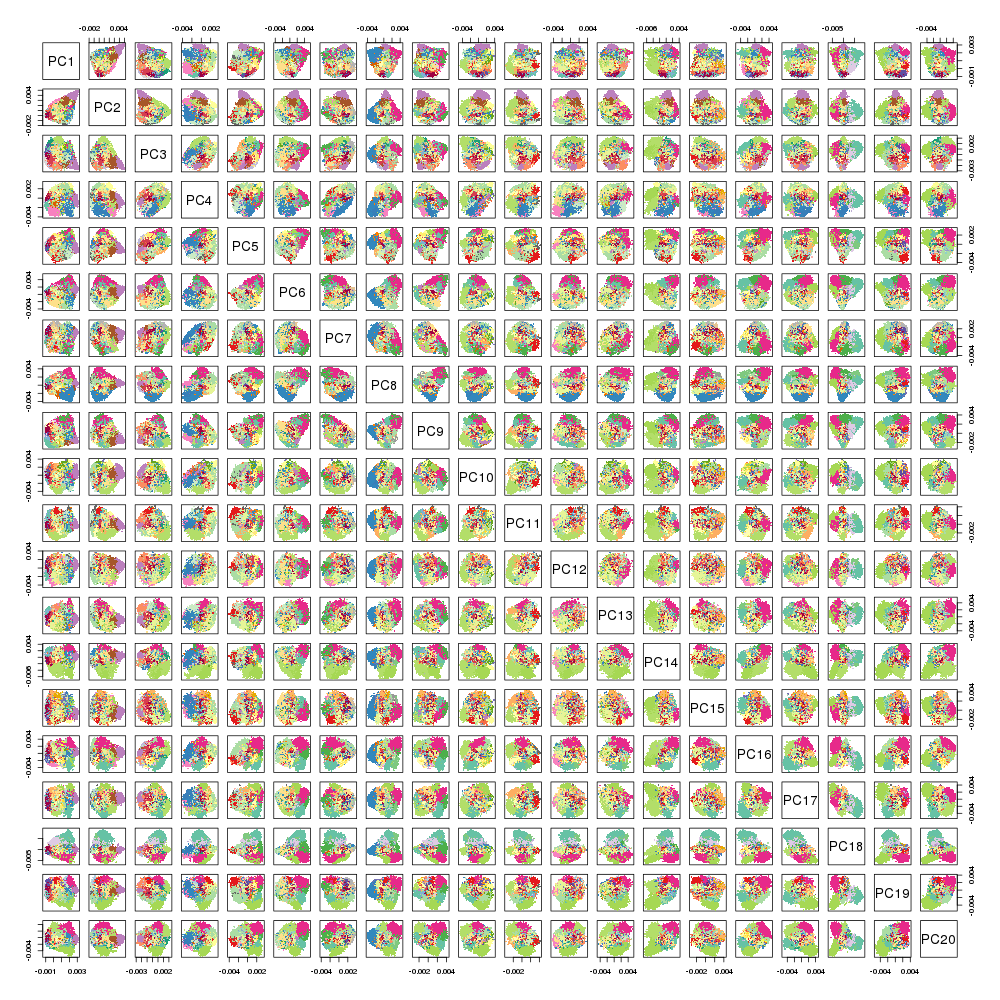

### Arpack.jl.png

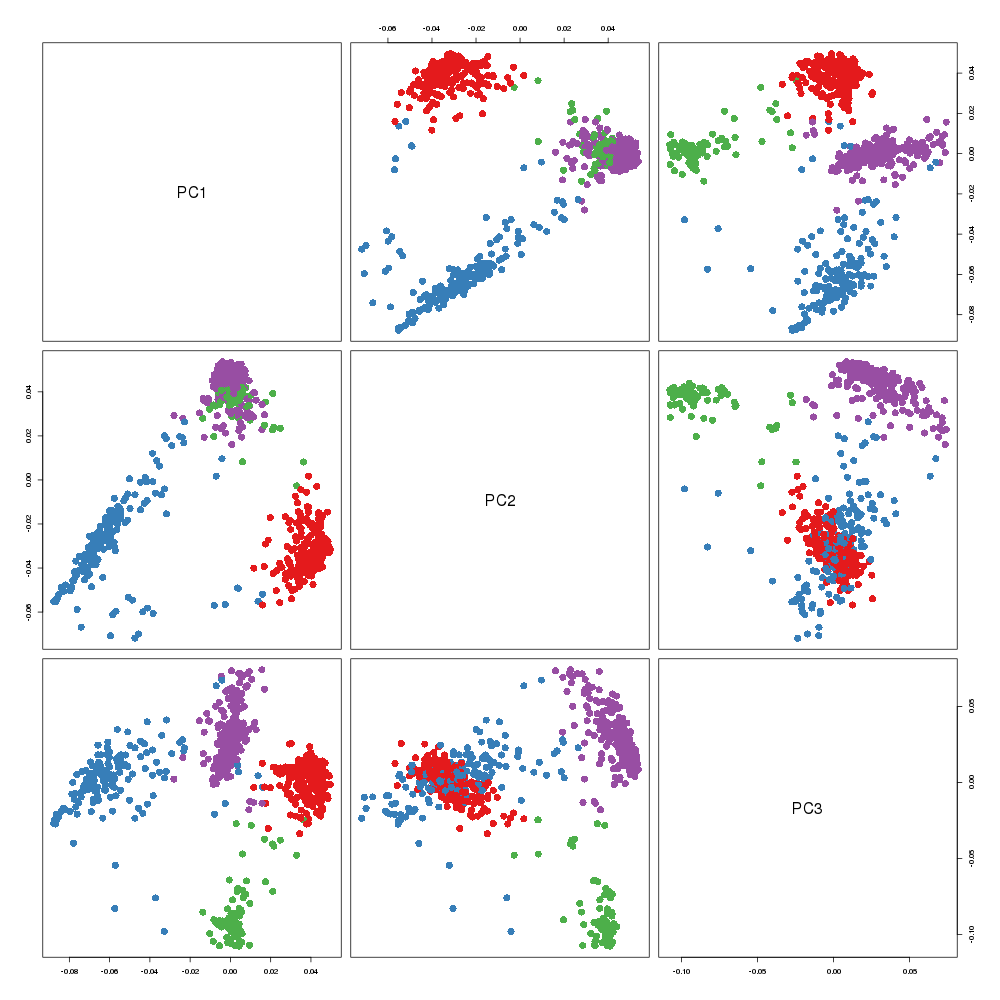

### Arpack.jl.png

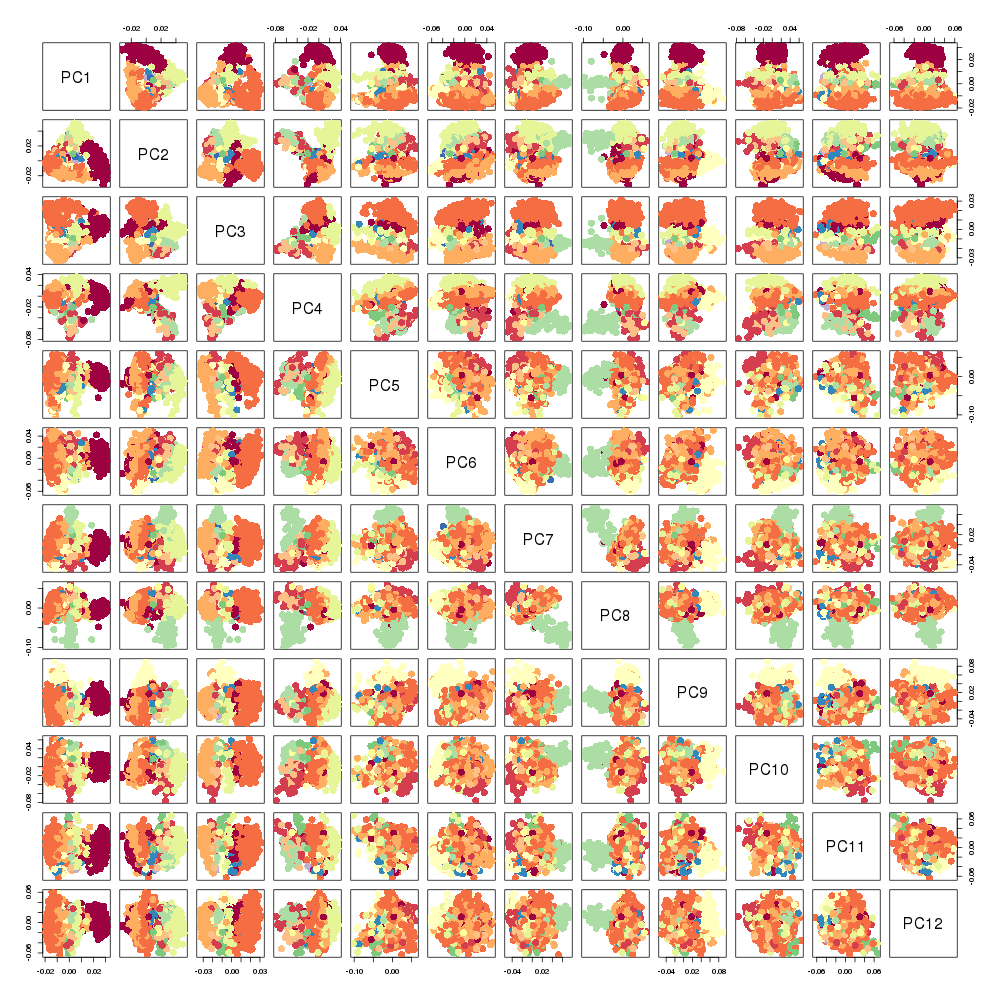

### CellRanger.png

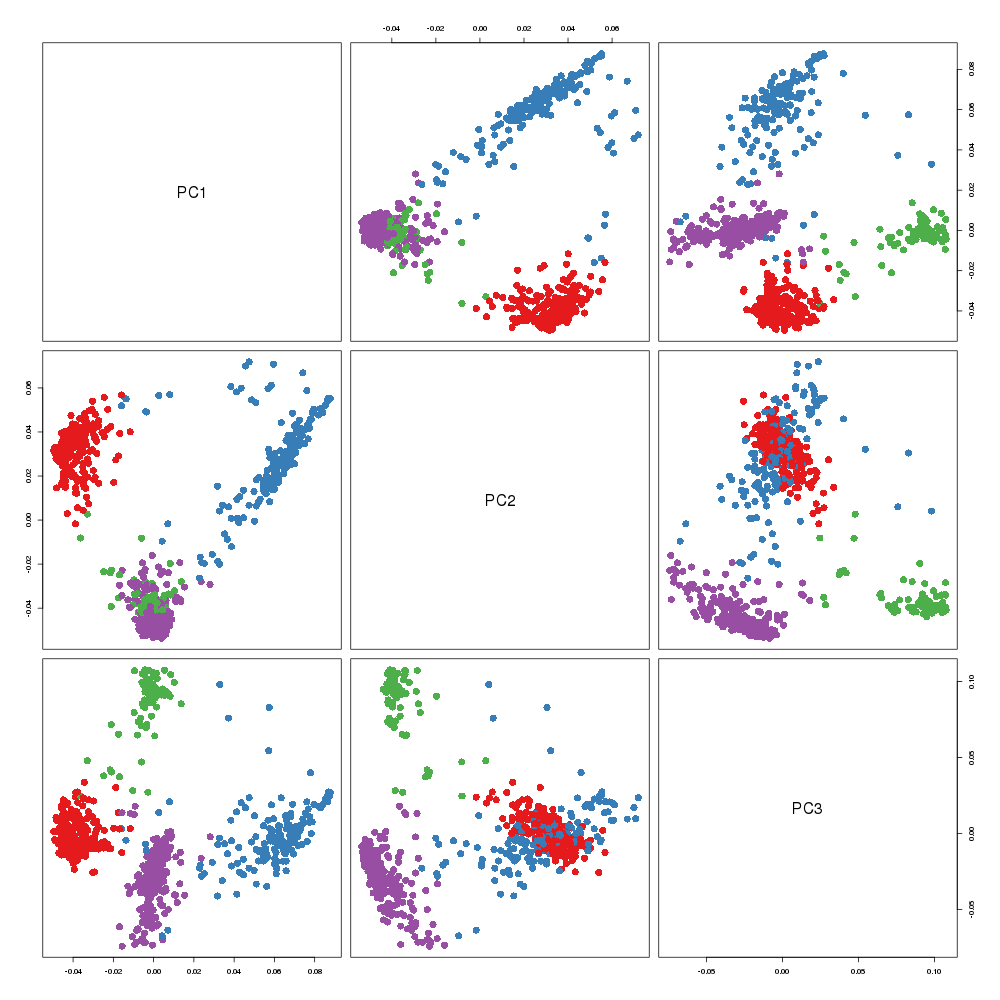

### CellRanger.png

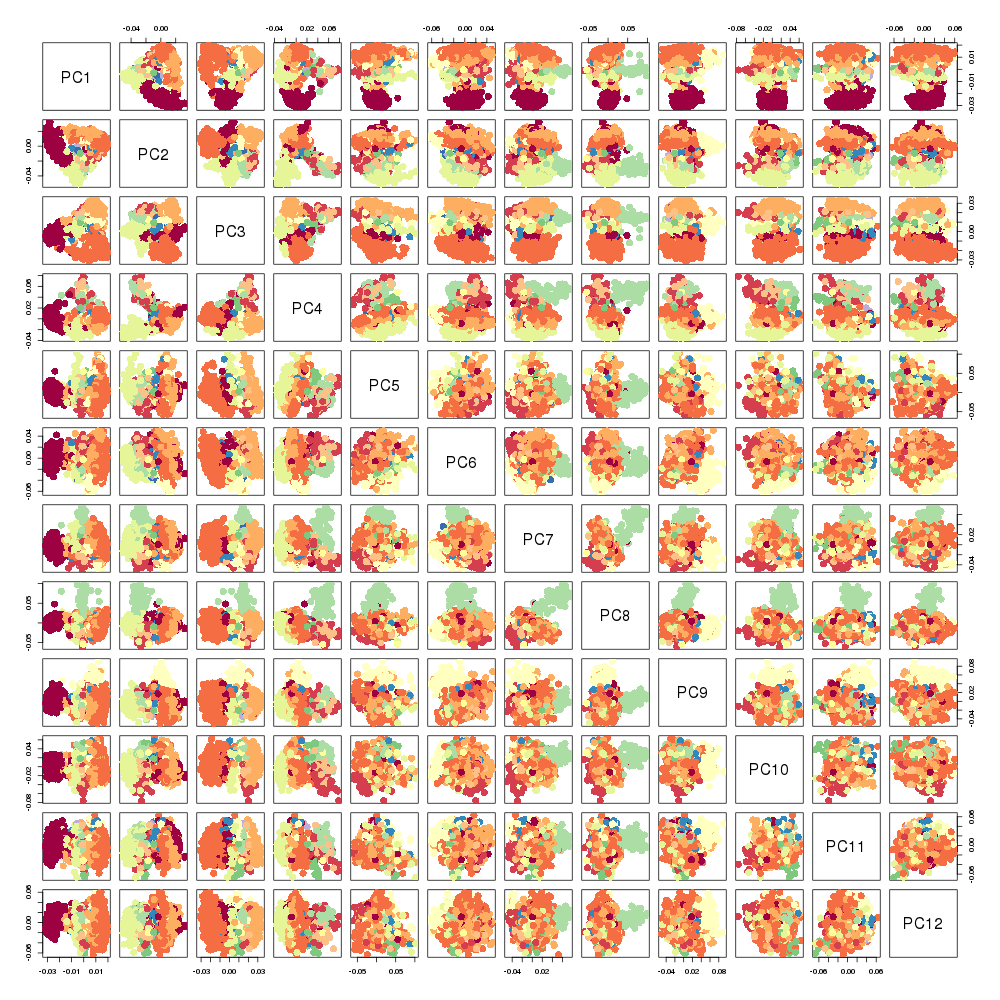

### dask_ml.png

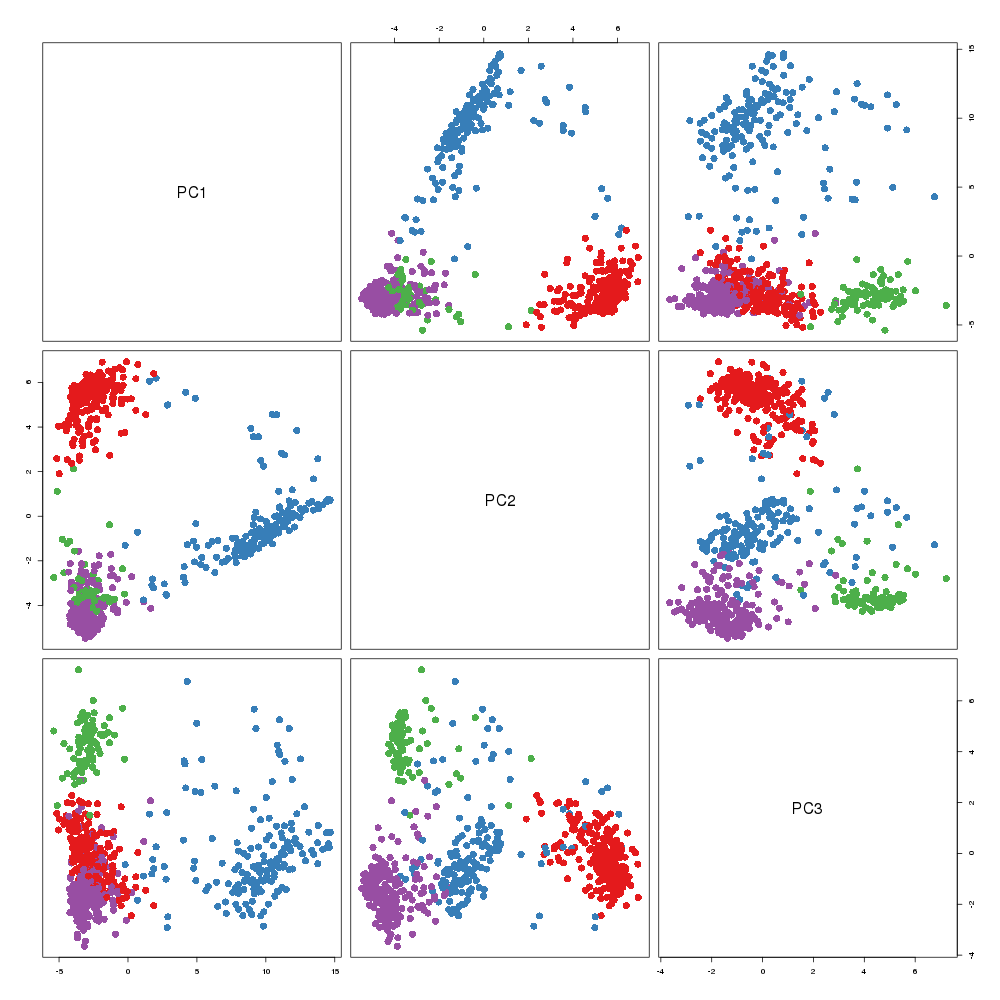

### dask_ml.png

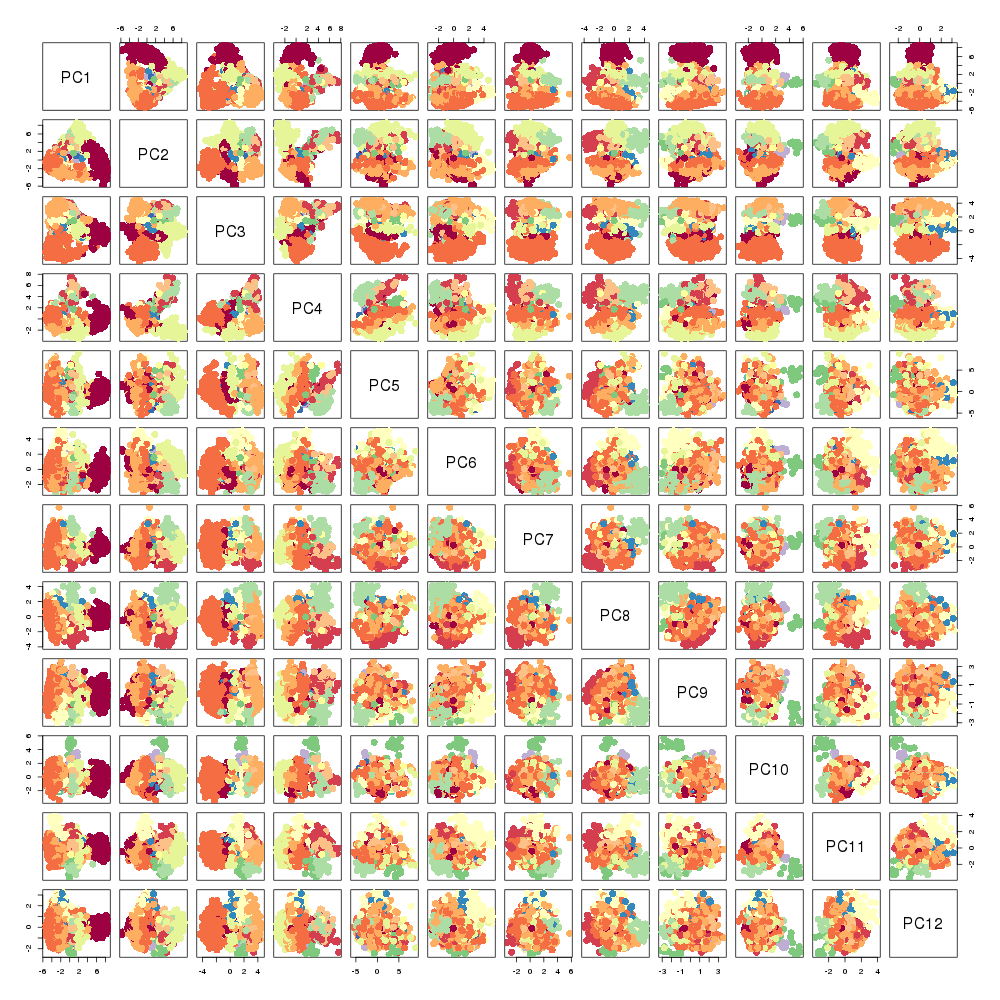

### Downsampling.png

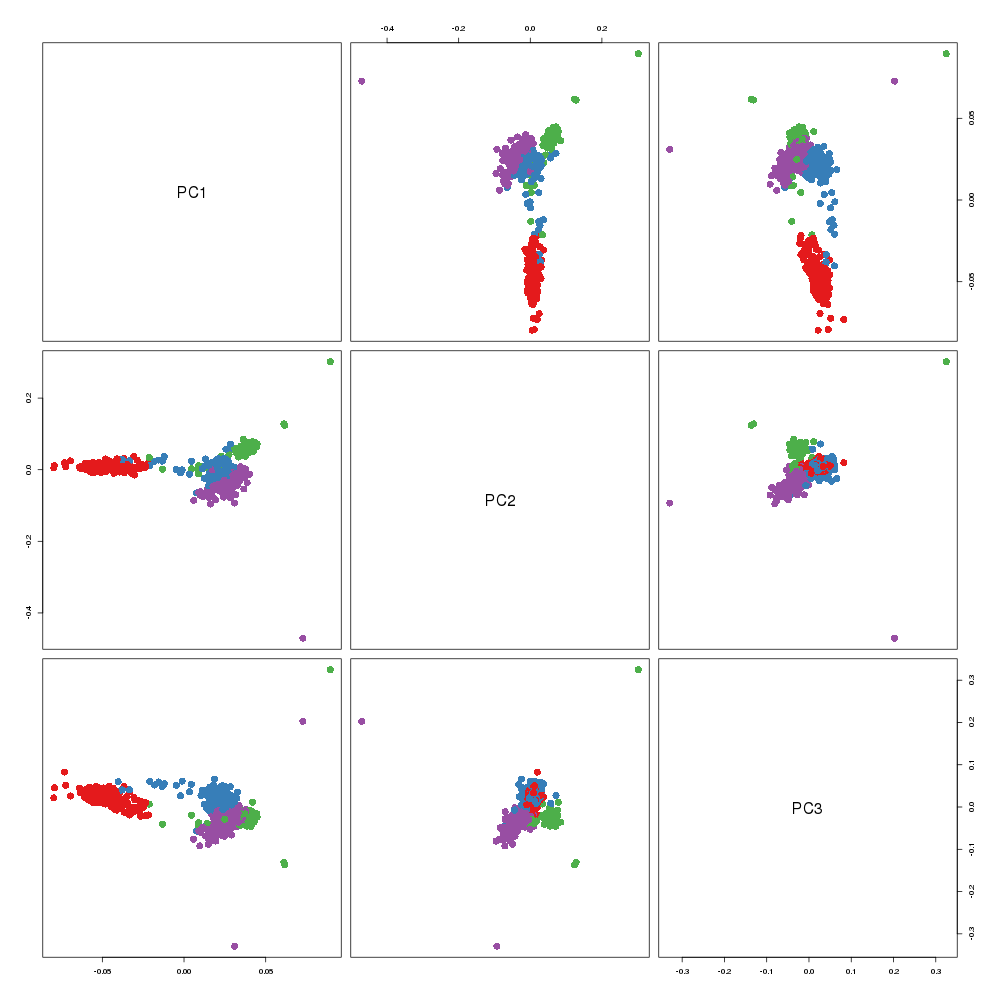

### Downsampling.png

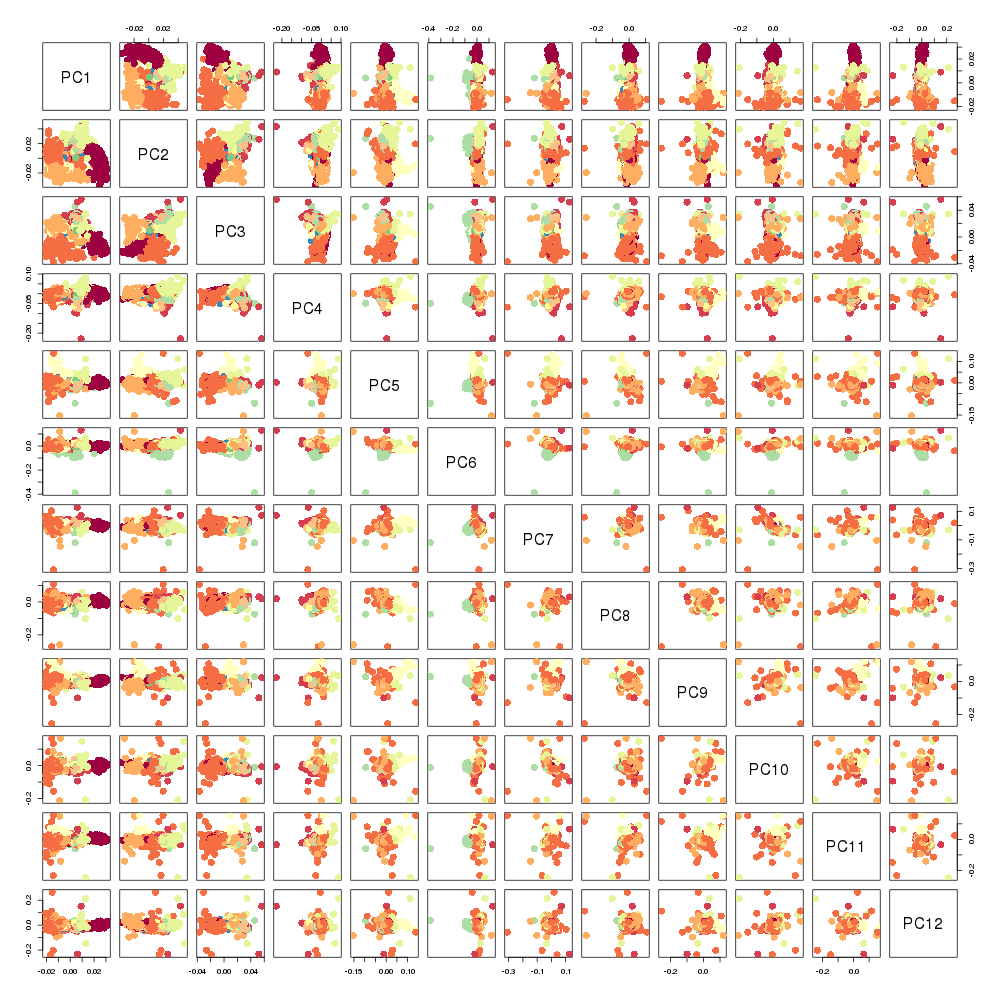

### Downsampling.png

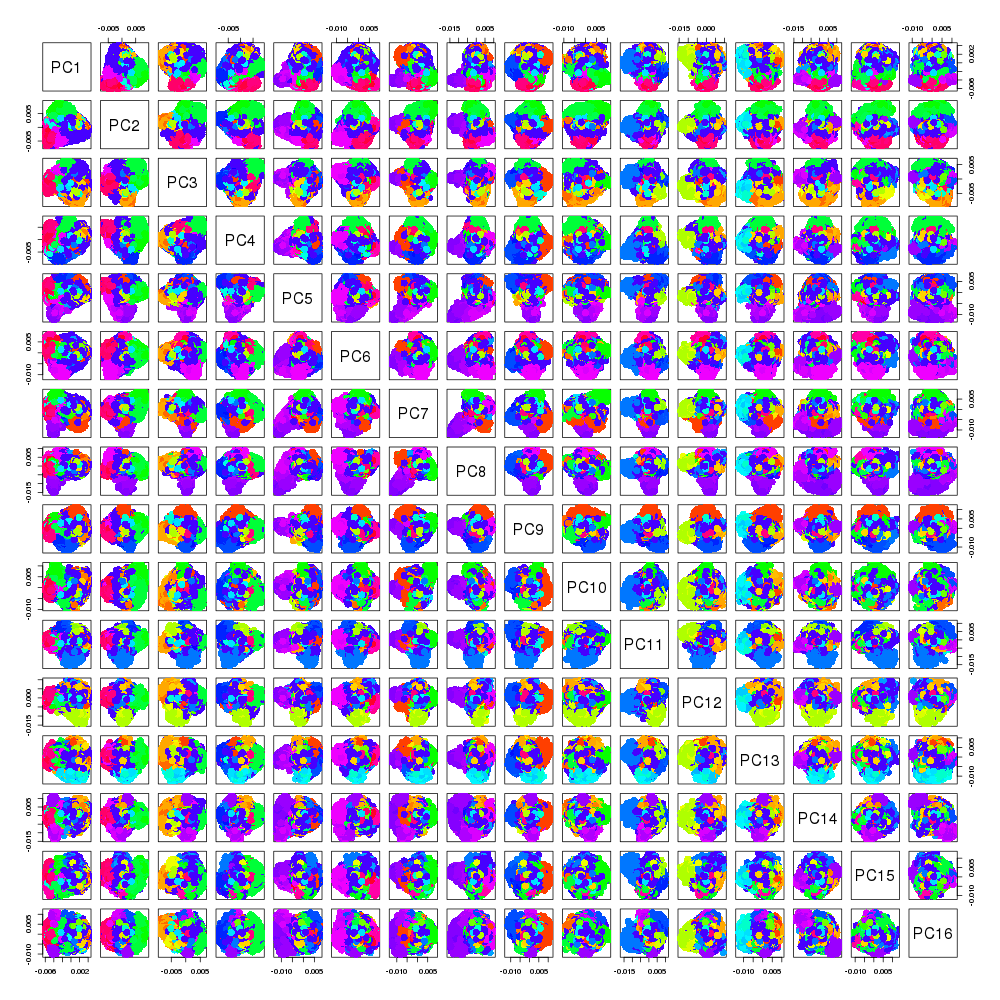

### Downsampling.png

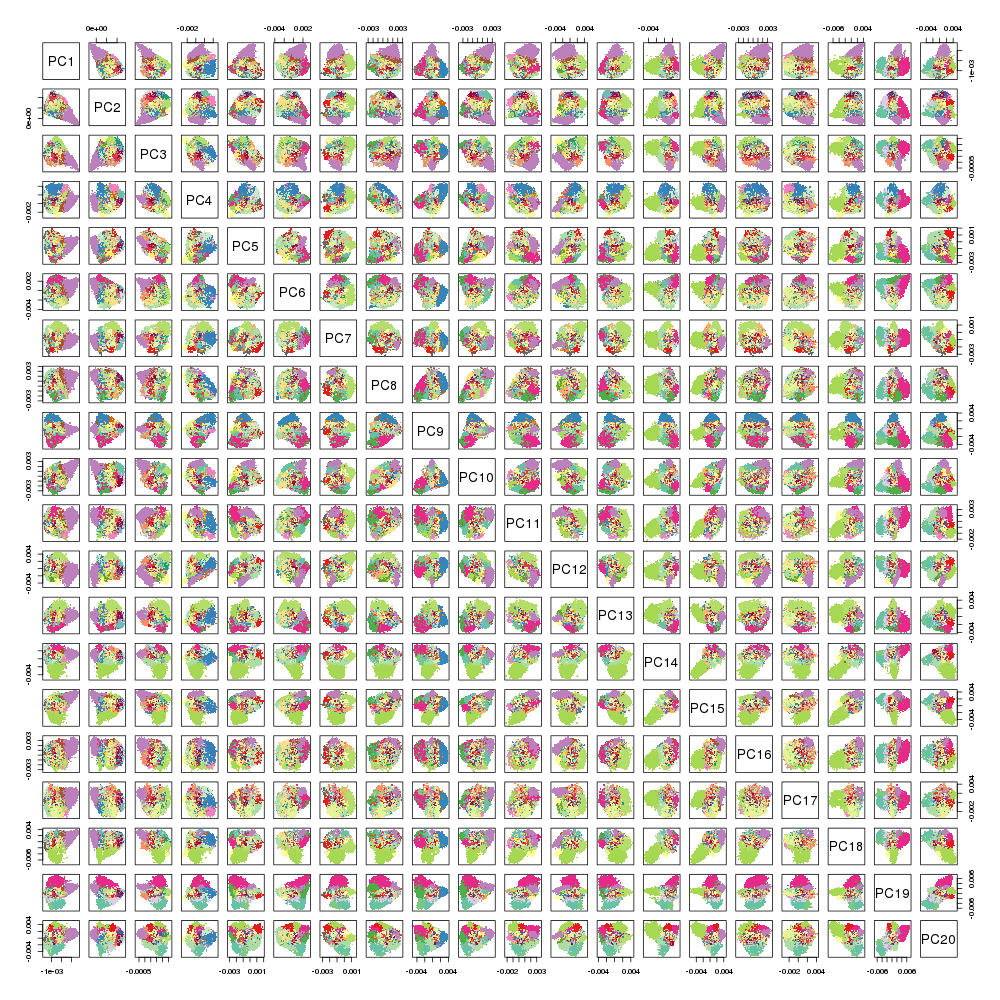

### GD_step1000_epoch10.png

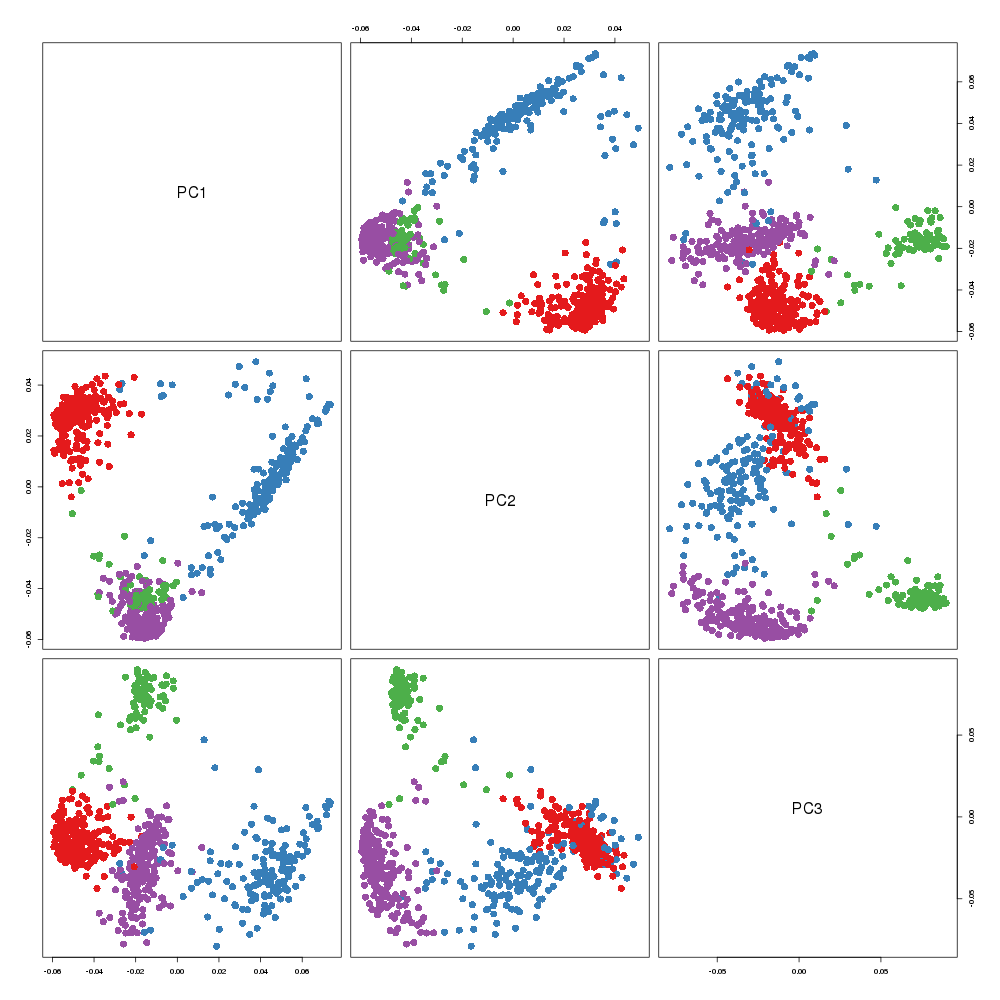

### GD_step1000_epoch10.png

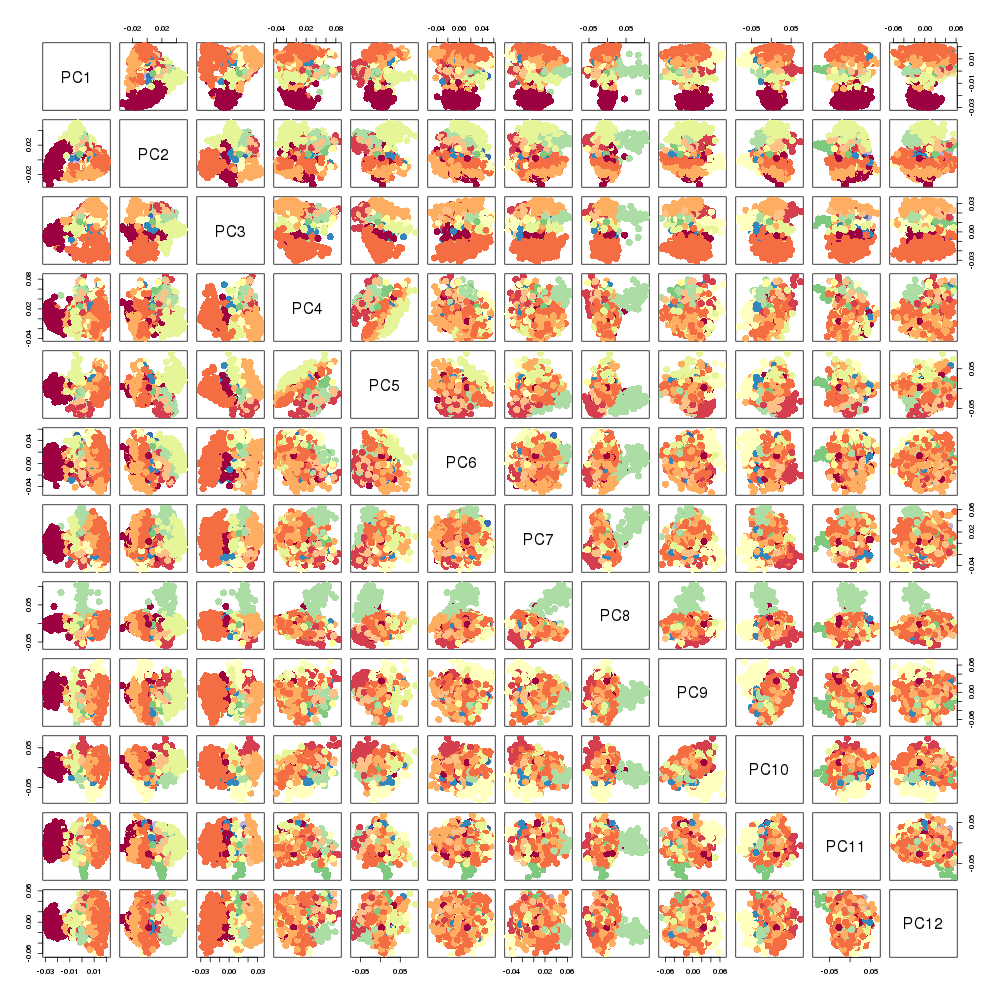

### GD_step1000_epoch10.png

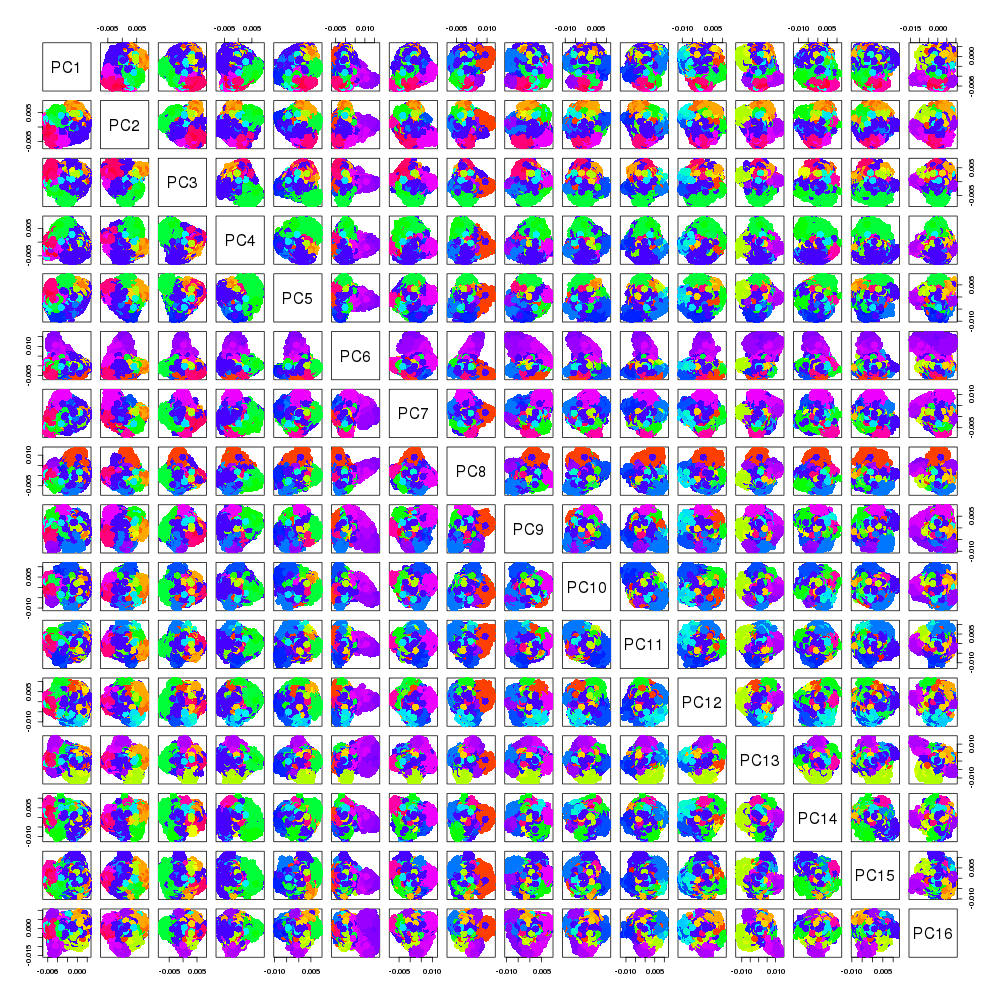

### GD_step1000_epoch10.png

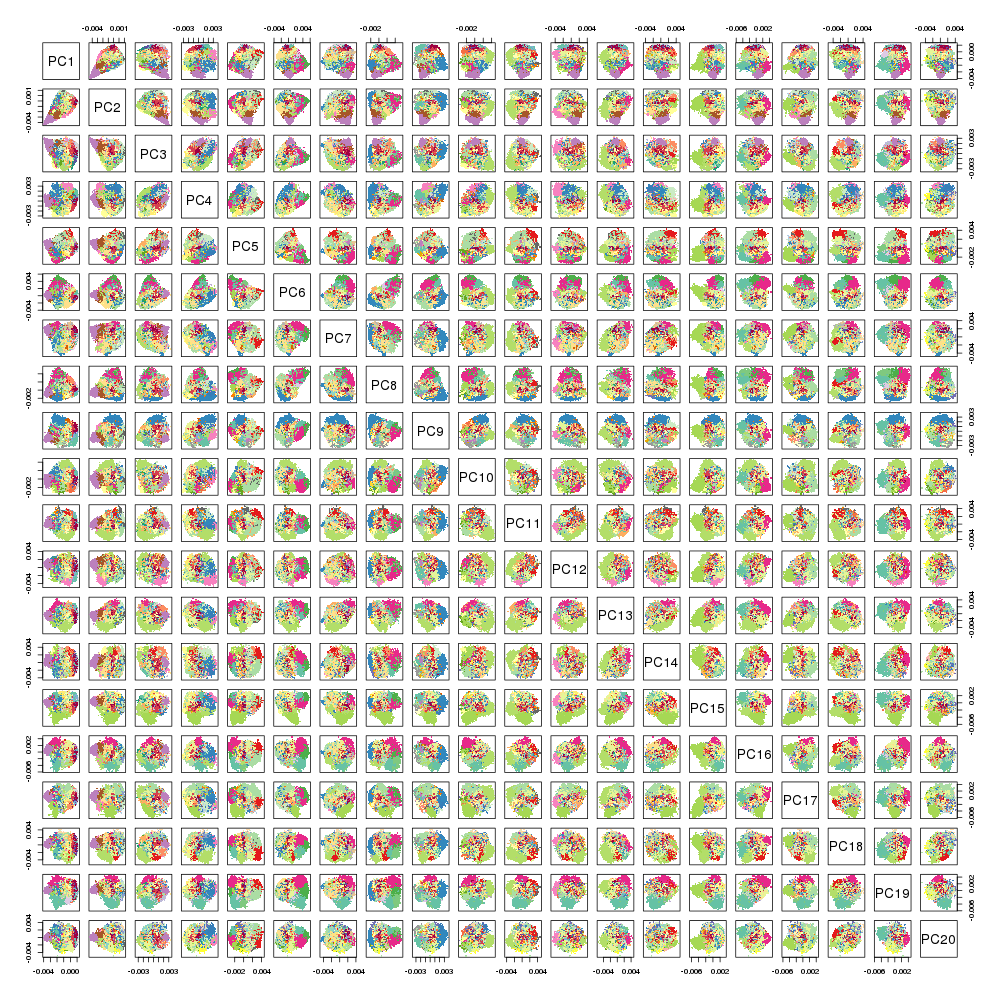

### Halko_0iter.png

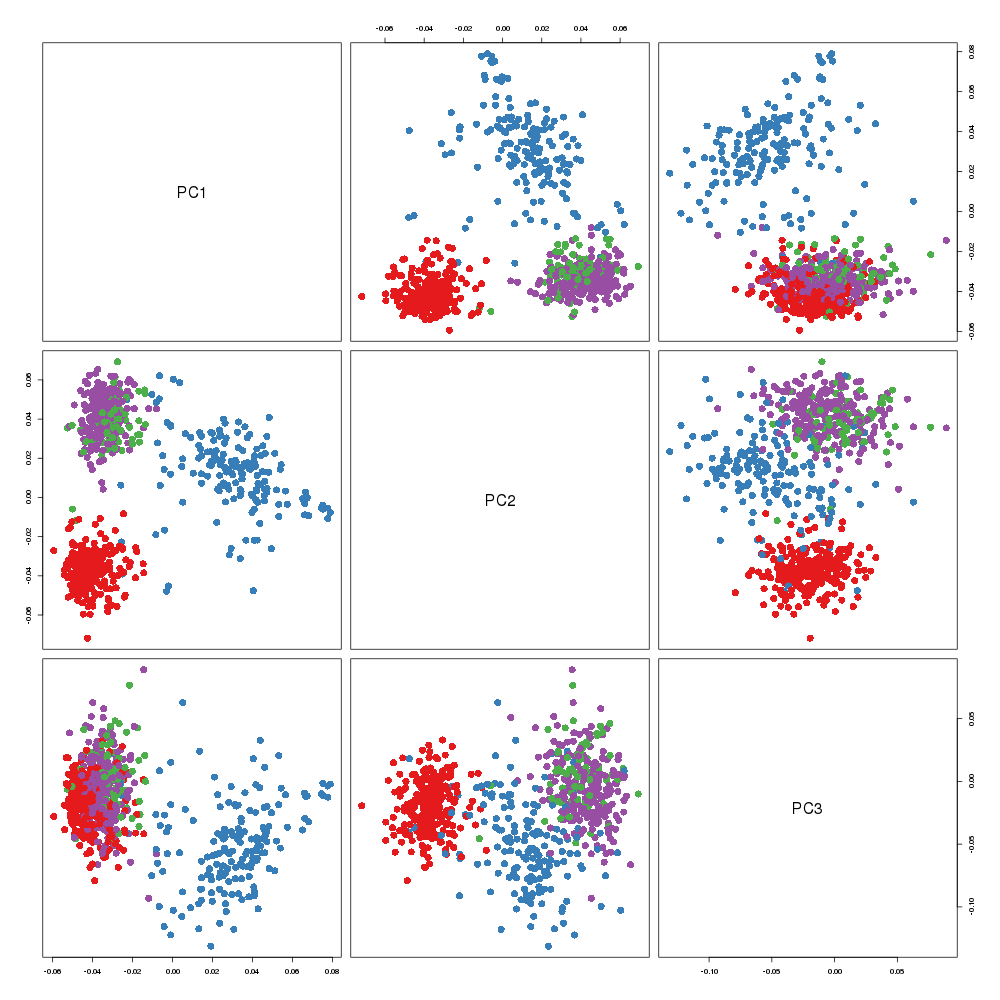

### Halko_0iter.png

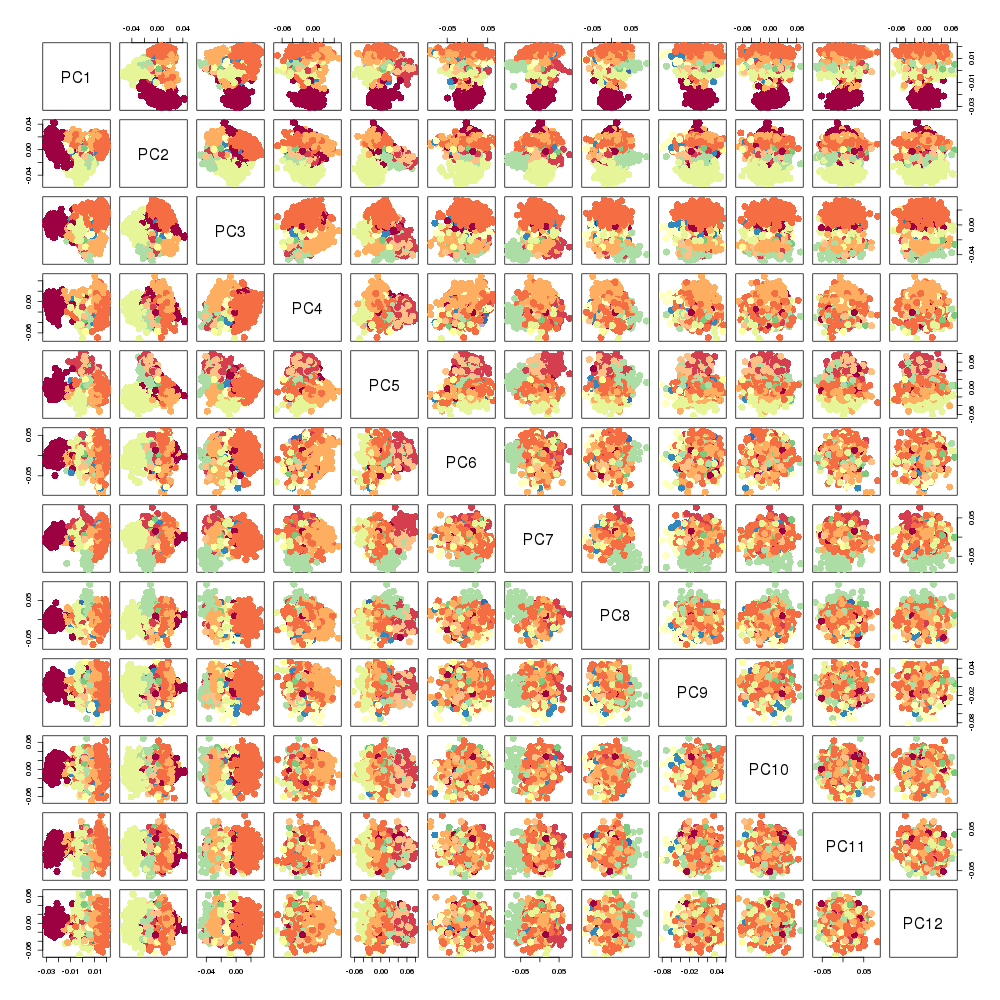

### Halko_0iter.png

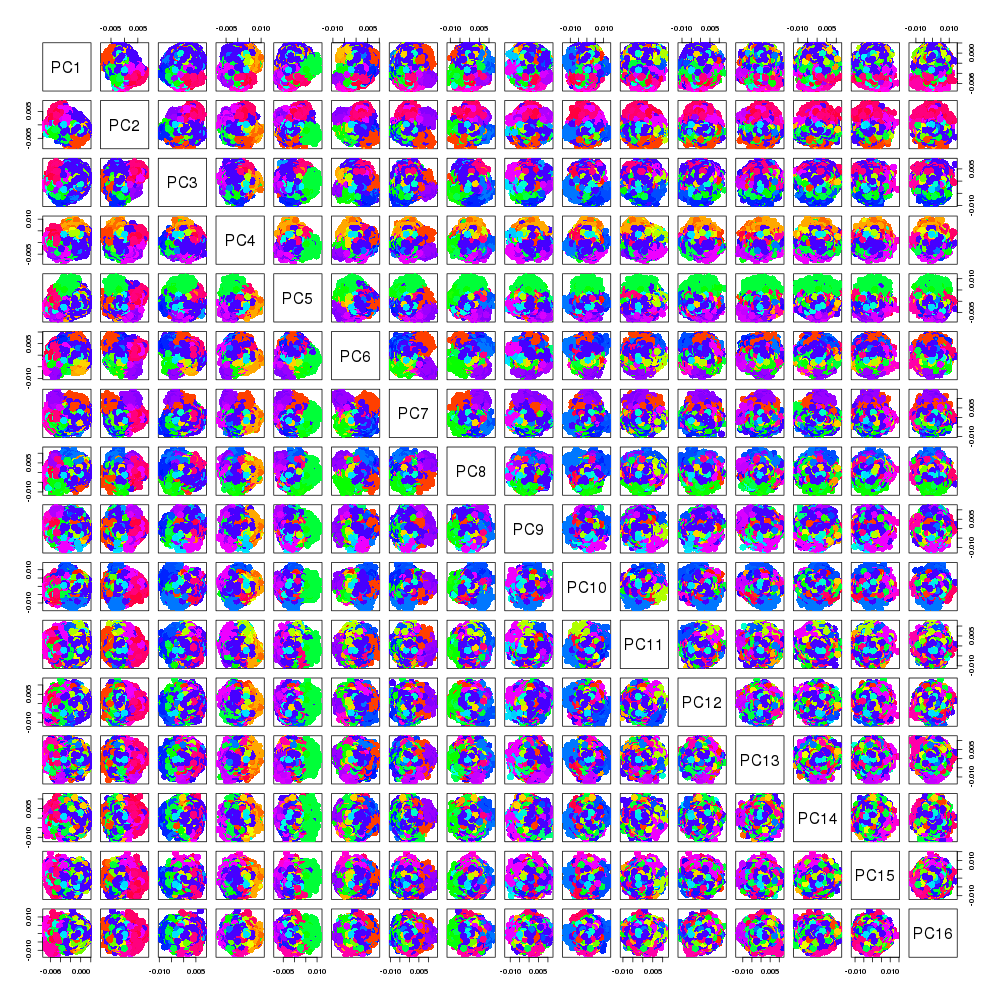

### Halko_0iter.png

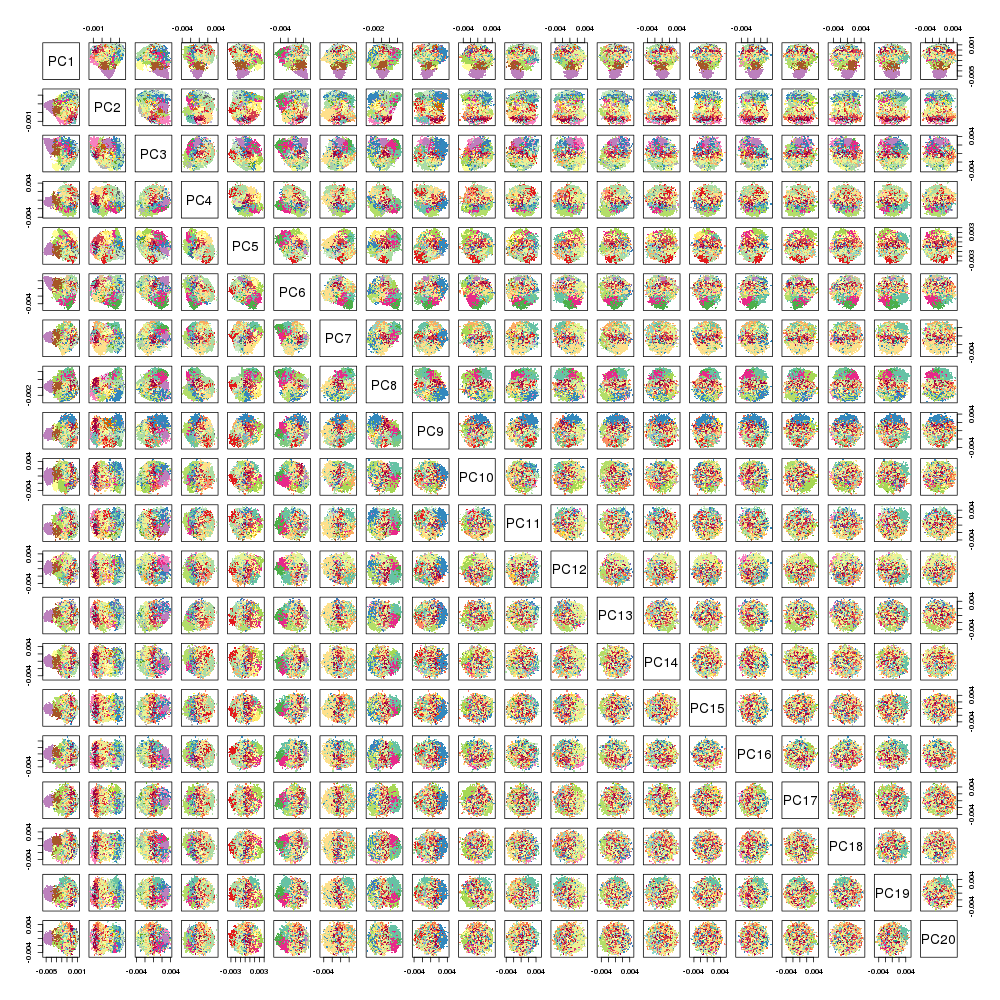

### Halko_1iter.png

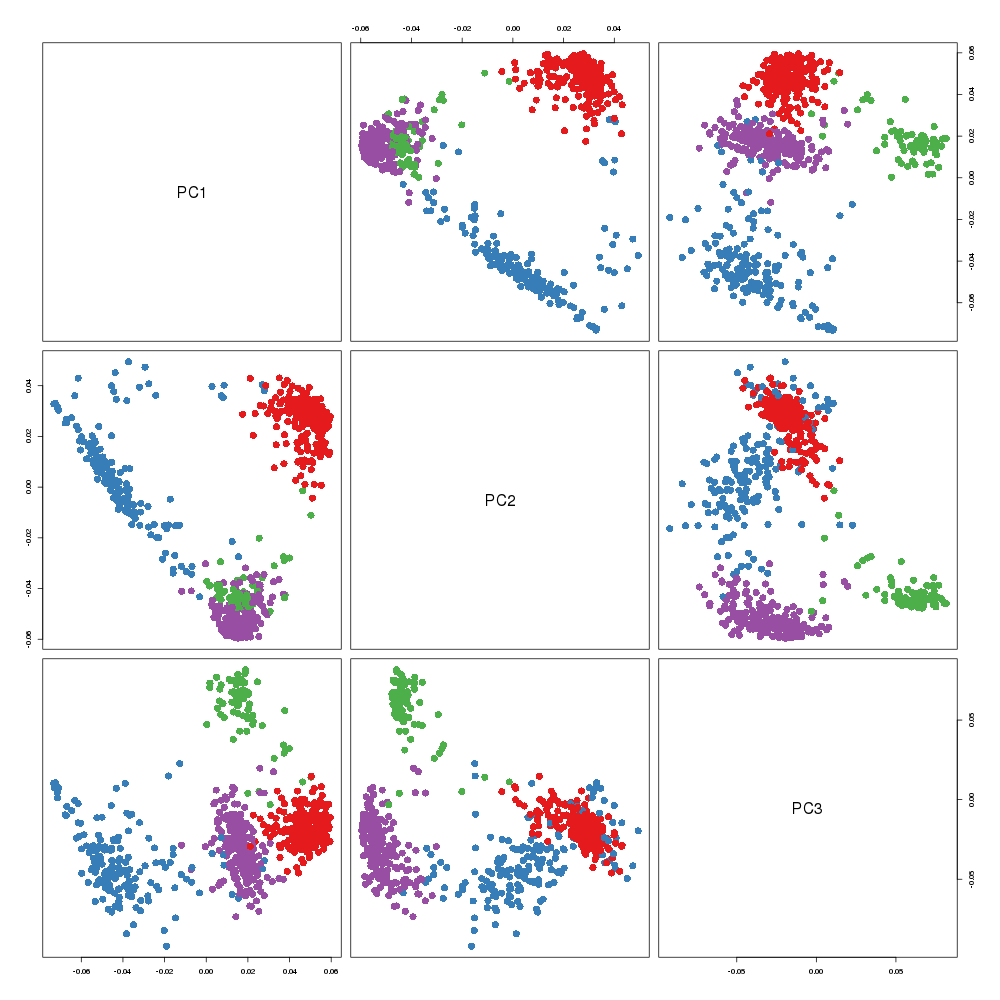

### Halko_1iter.png

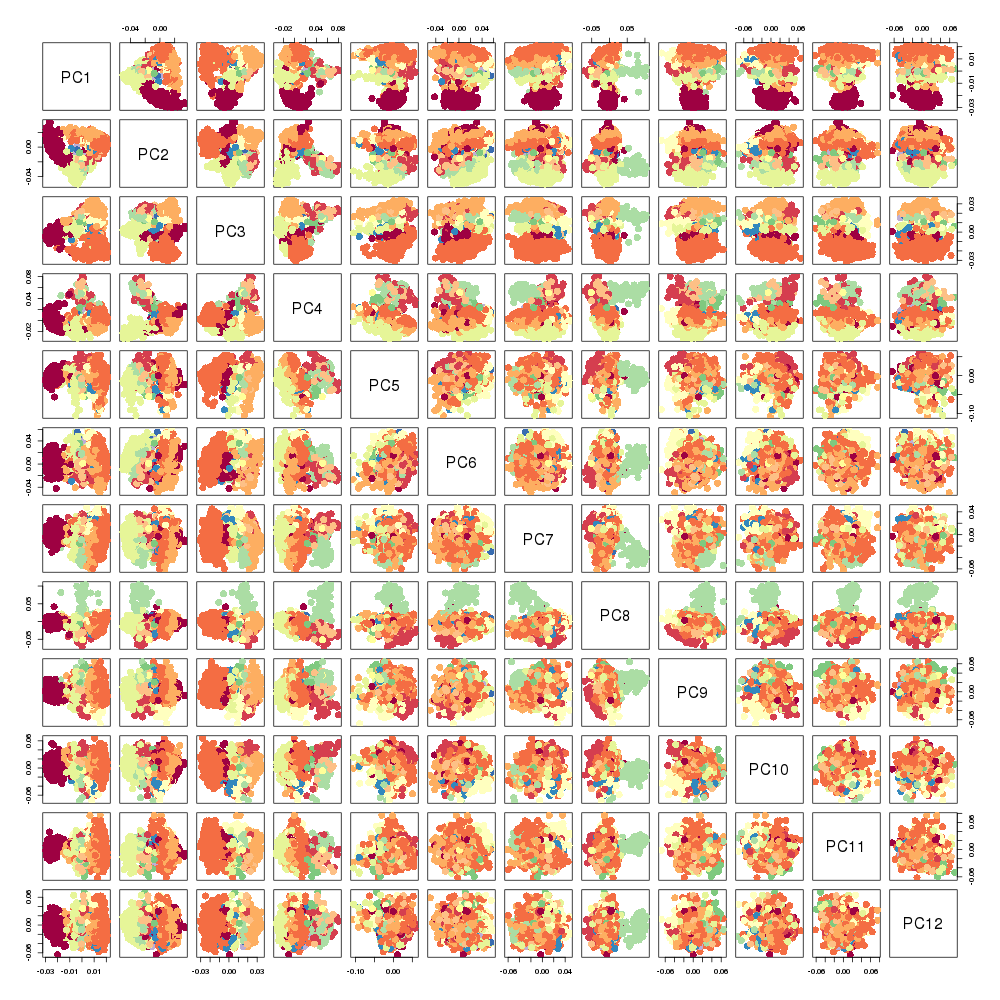

### Halko_1iter.png

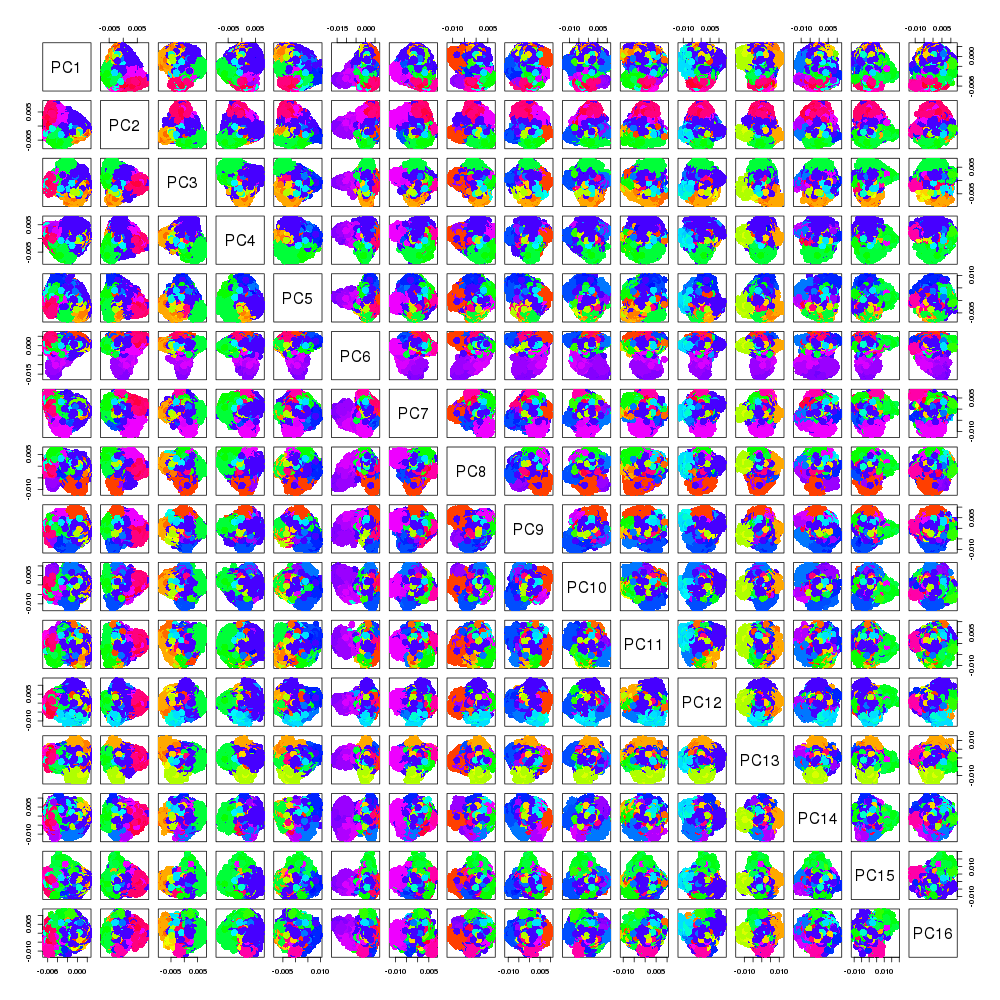

### Halko_1iter.png

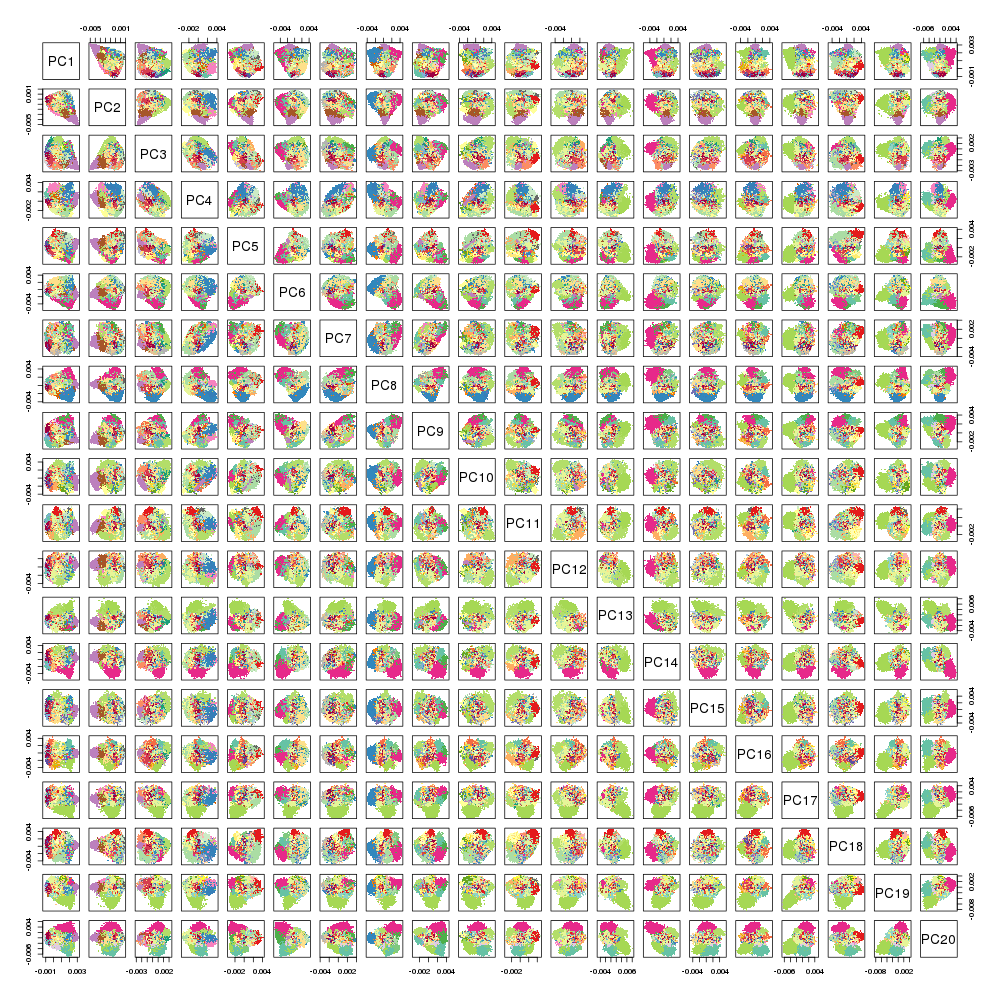

### Halko_2iter.png

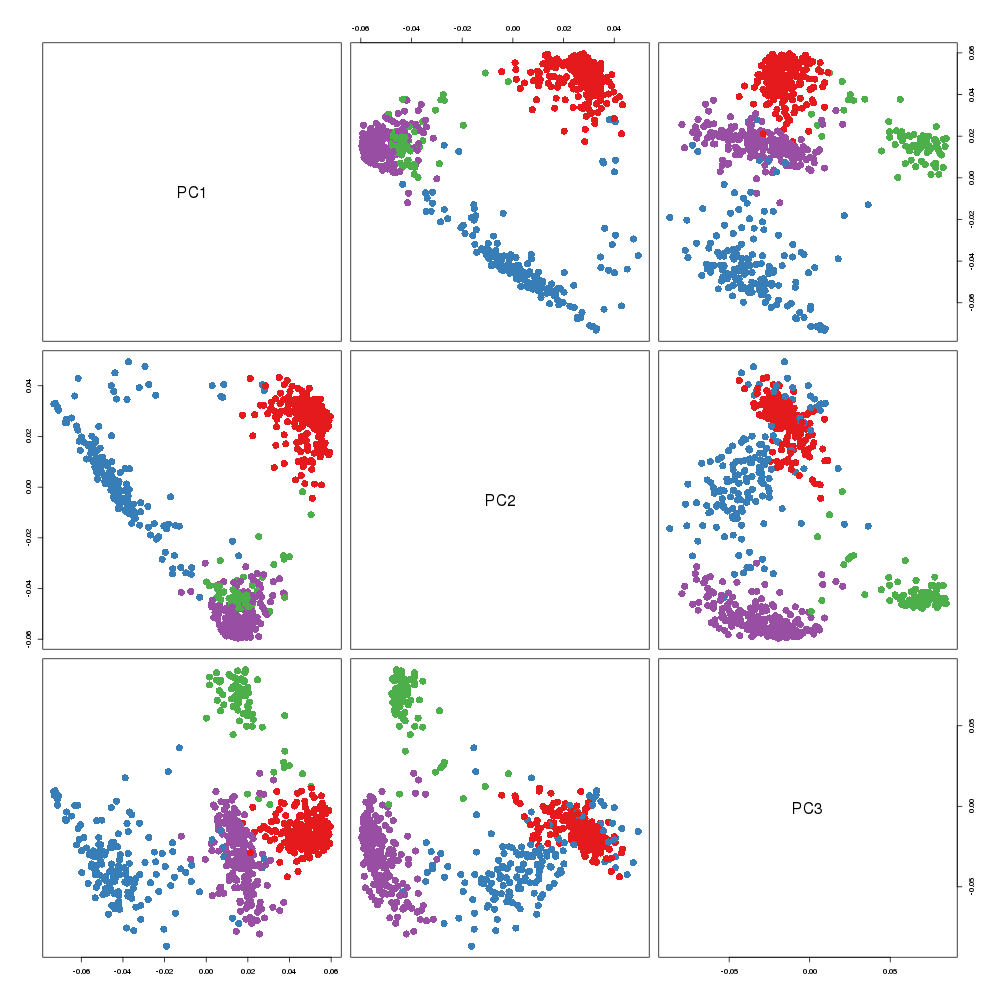

### Halko_2iter.png

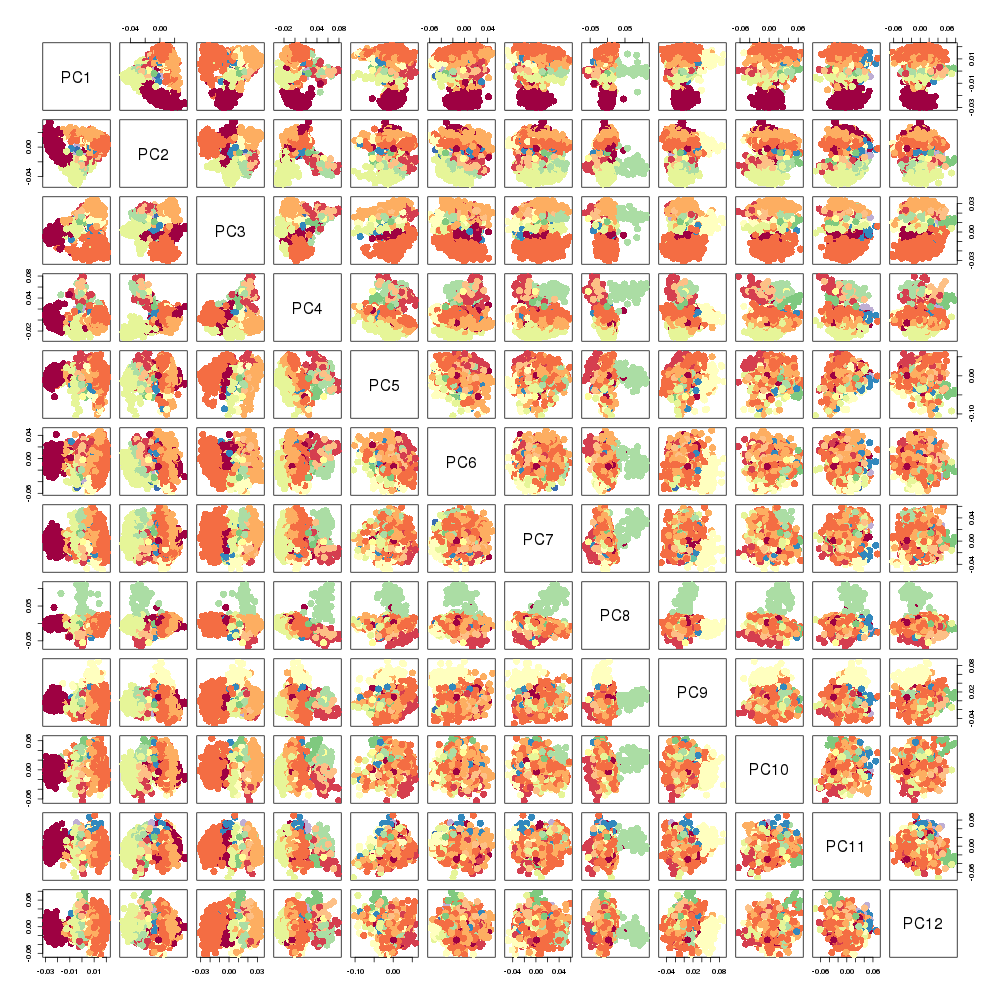

### Halko_2iter.png

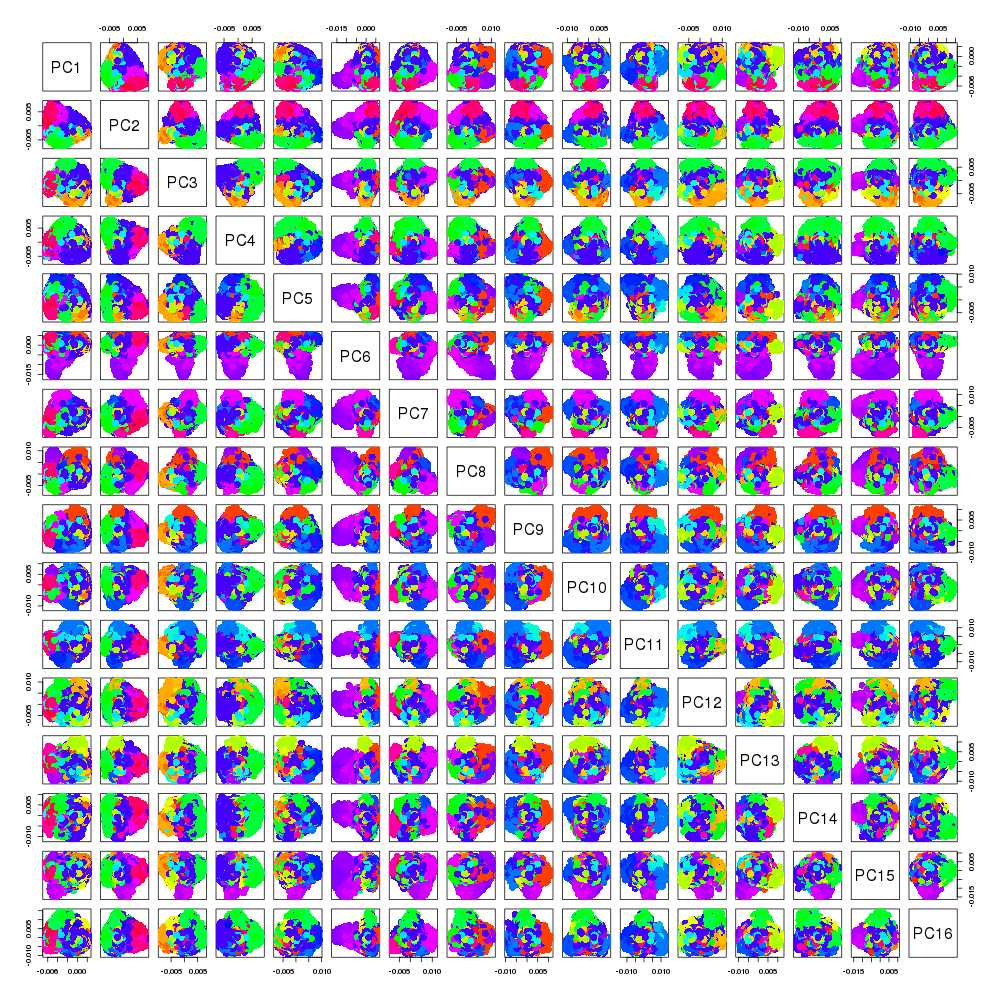

### Halko_2iter.png

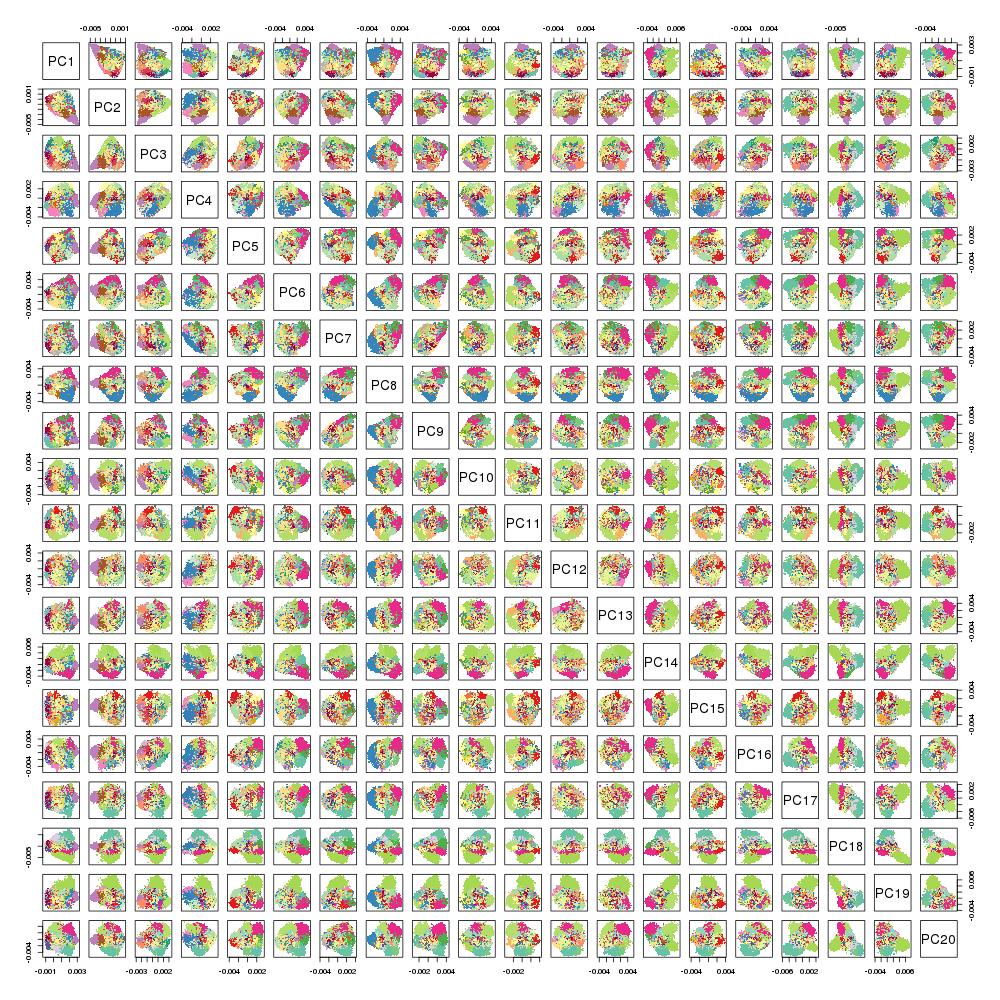

### Halko_3iter.png

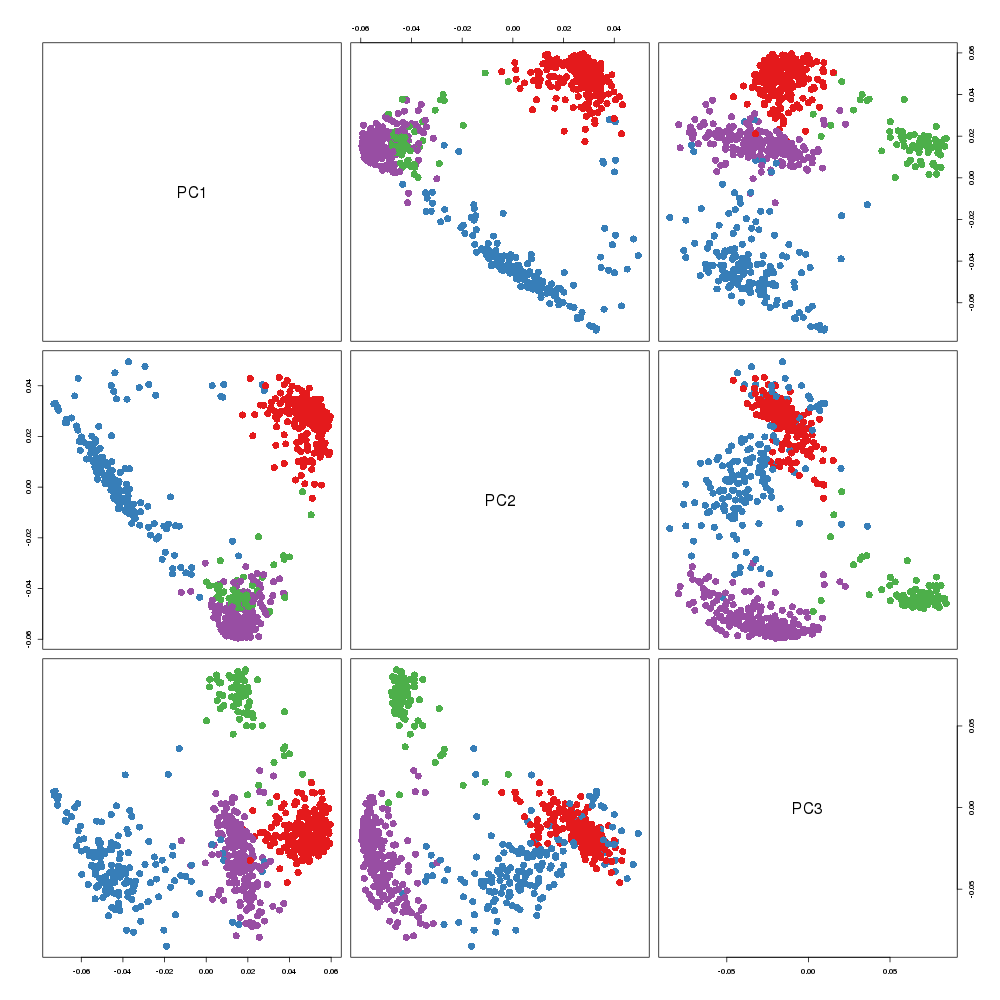

### Halko_3iter.png

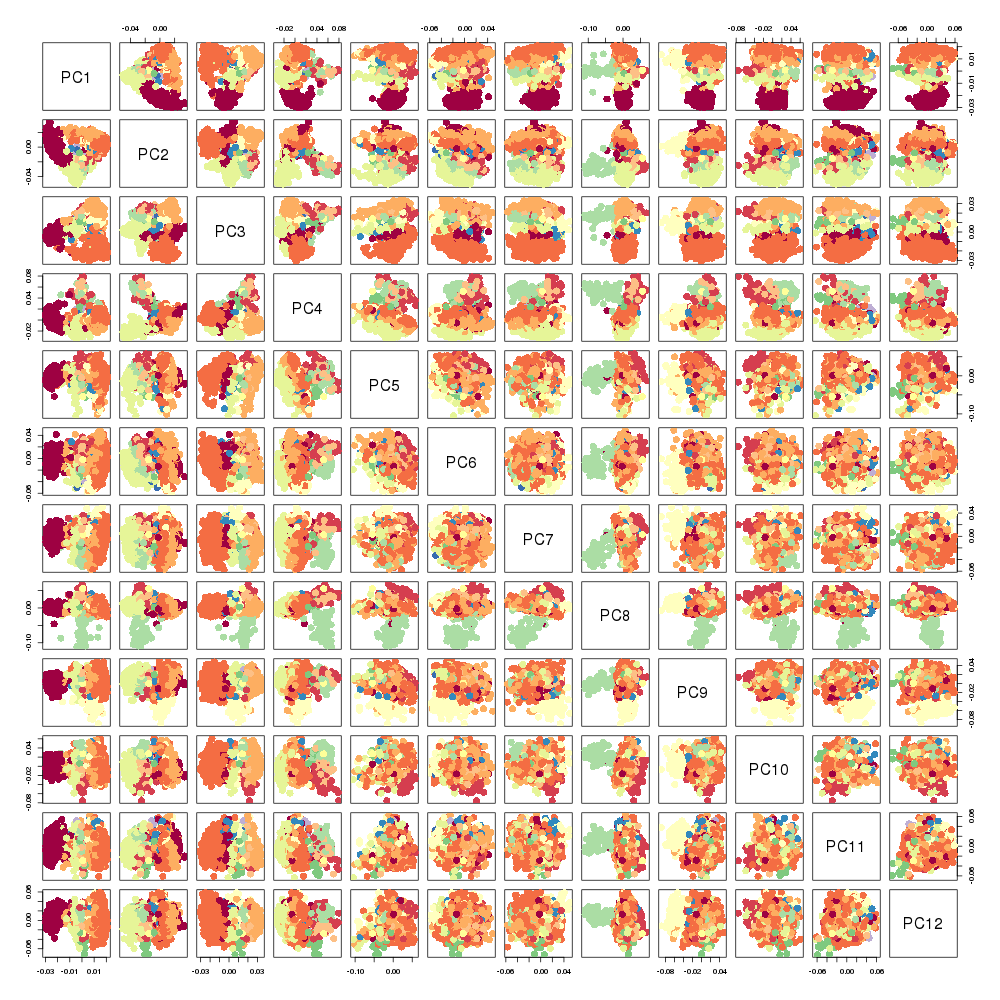

### Halko_3iter.png

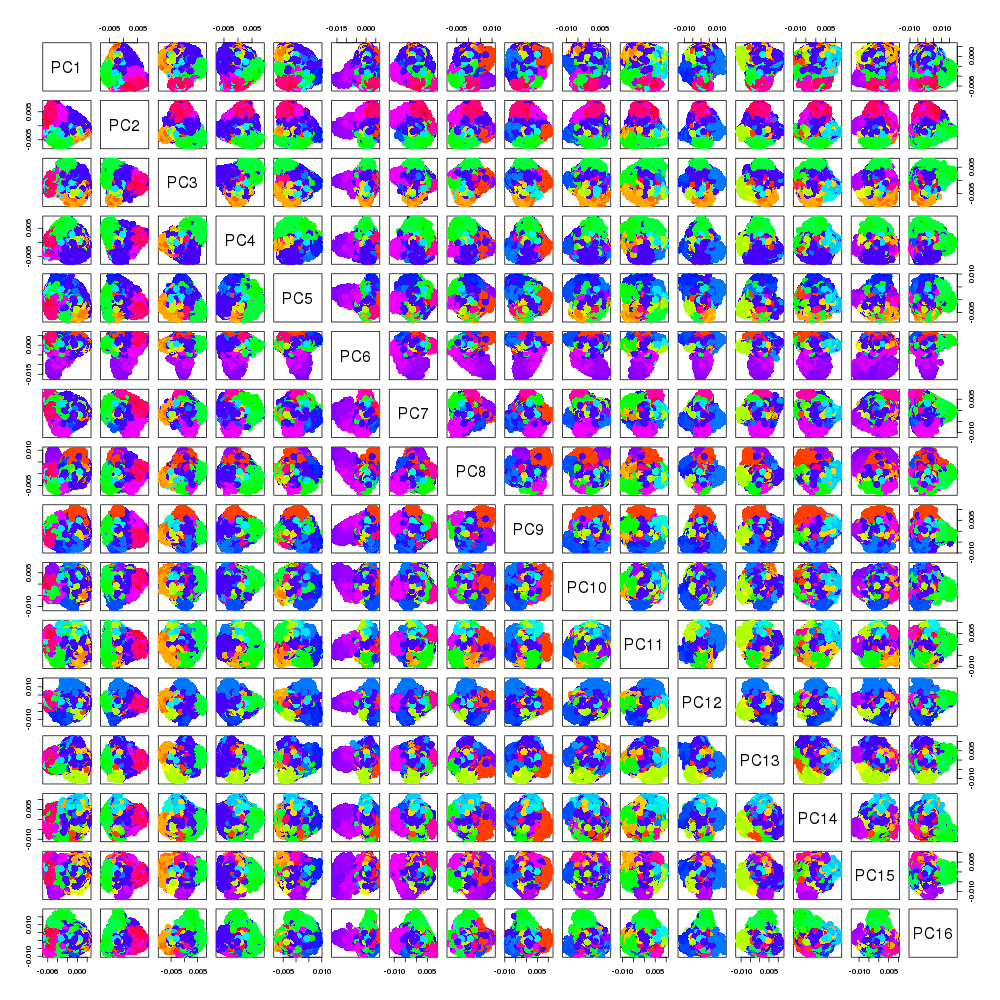

### Halko_3iter.png

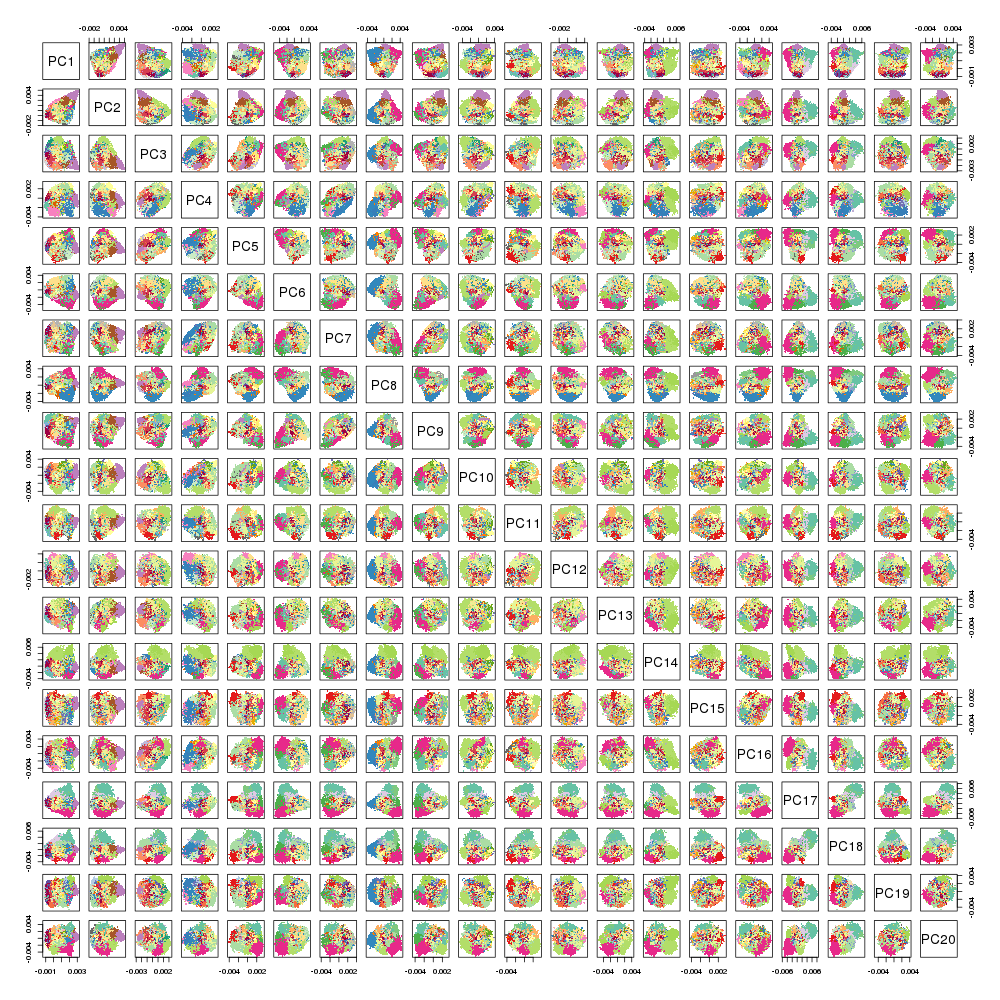

### IRLB.png

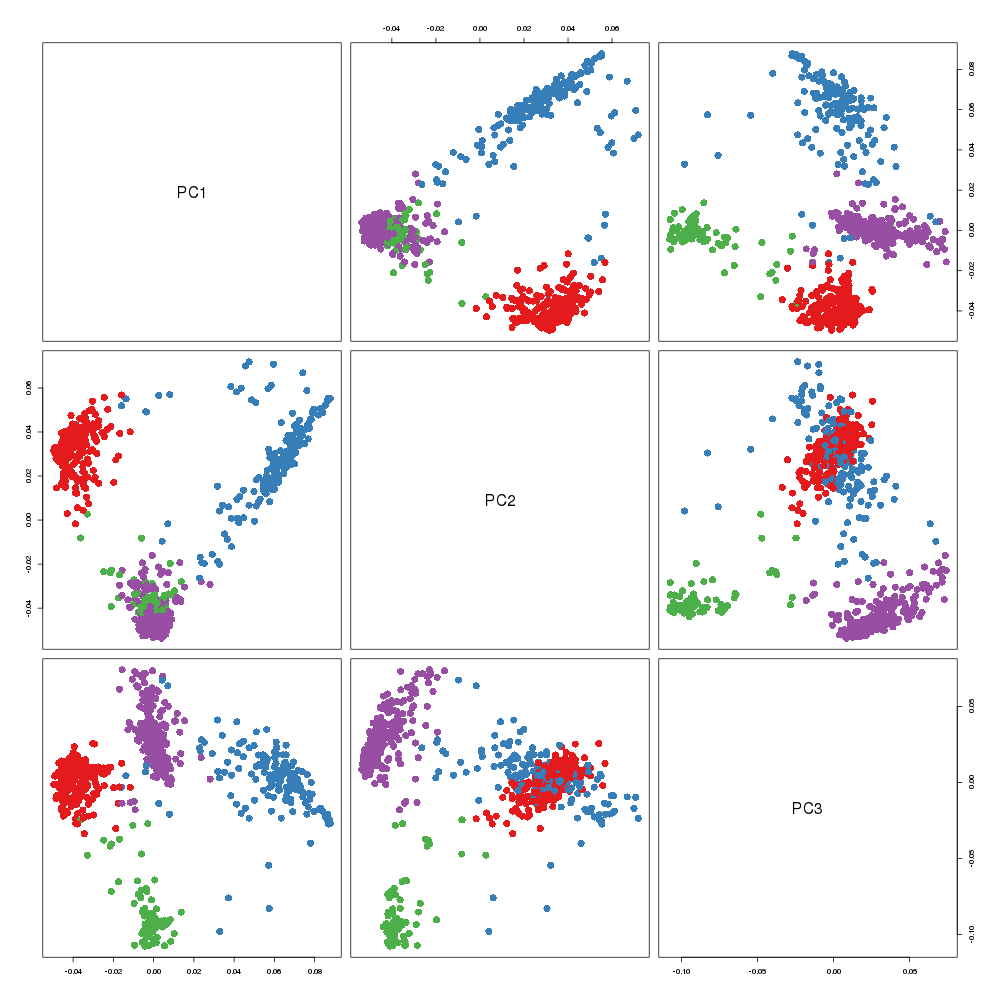

### IRLB.png

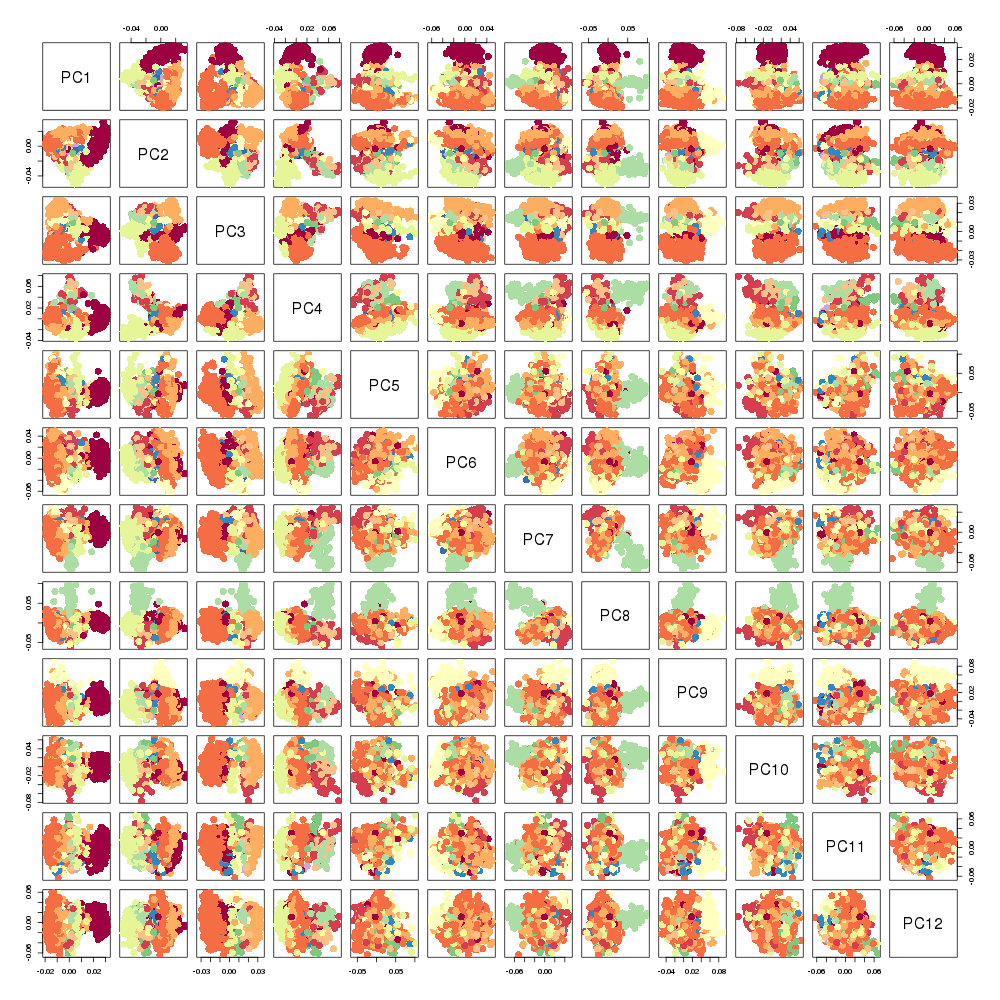

### MultivariateStats.png

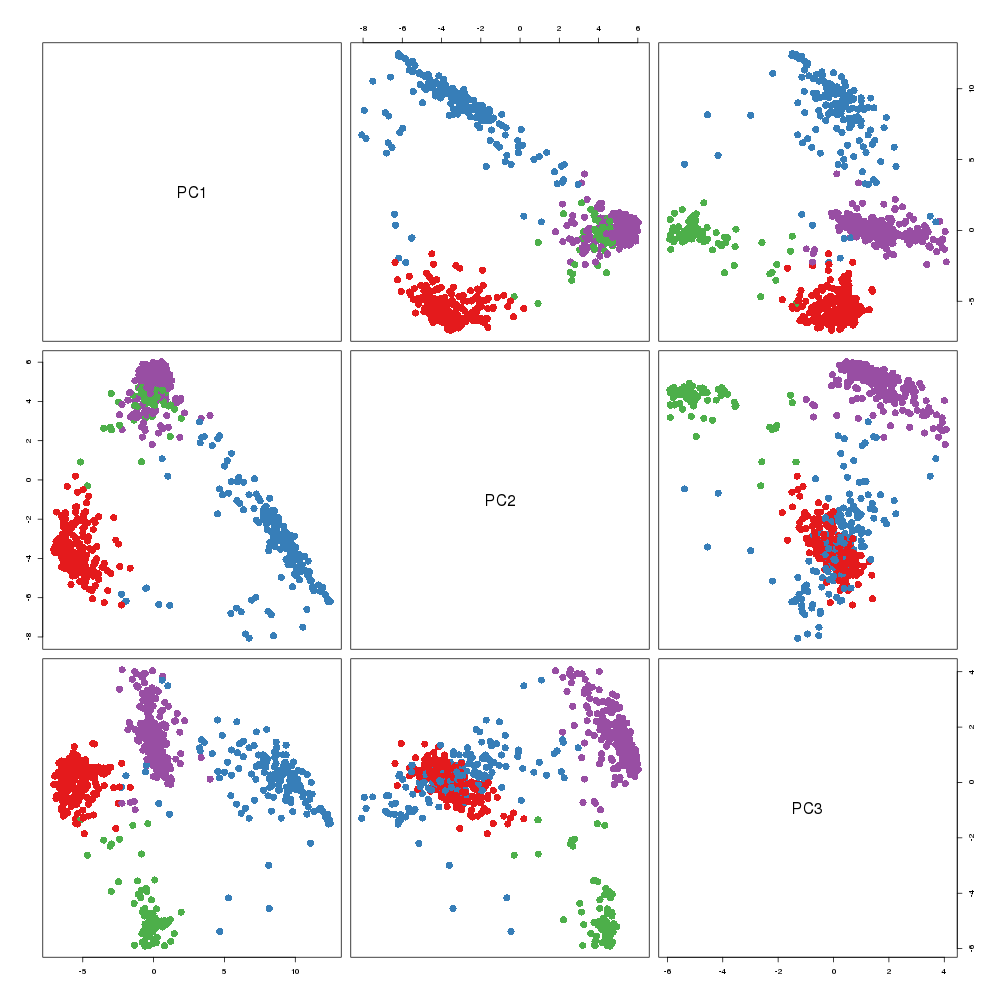

### MultivariateStats.png

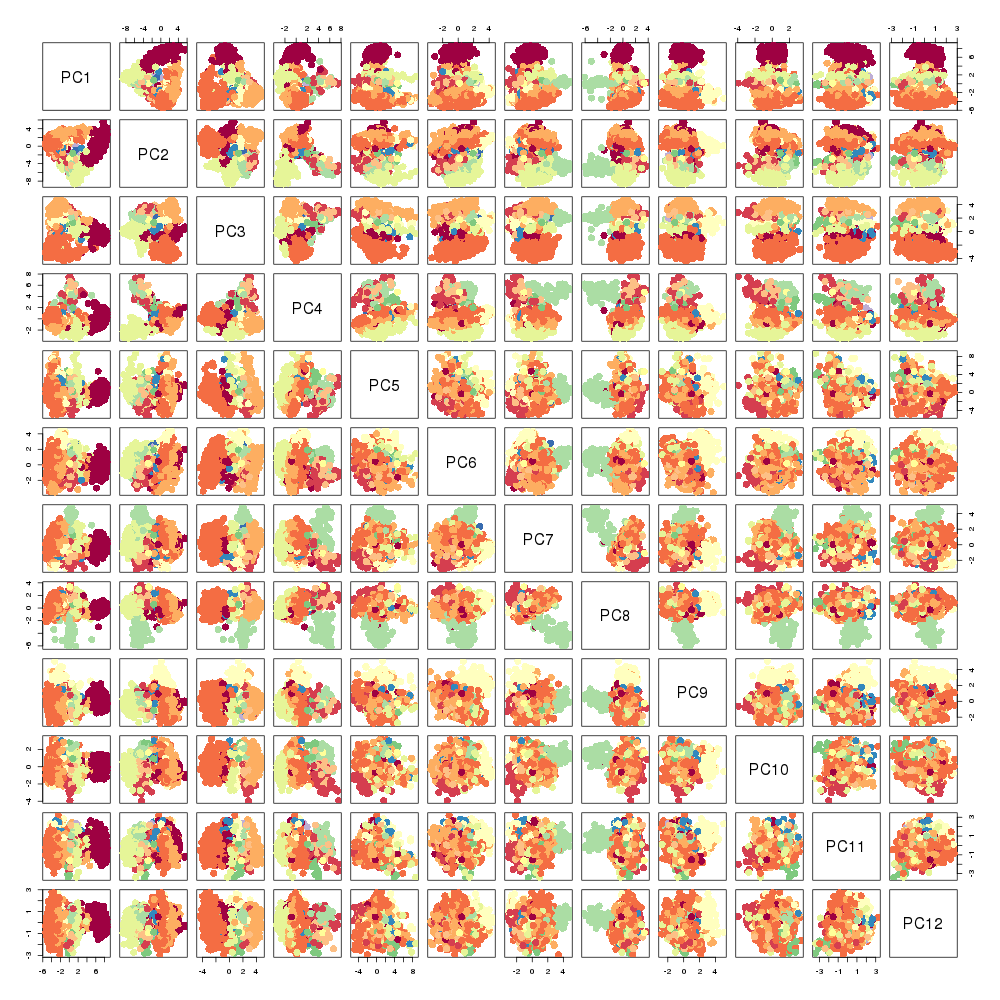

### oocRPCA.png

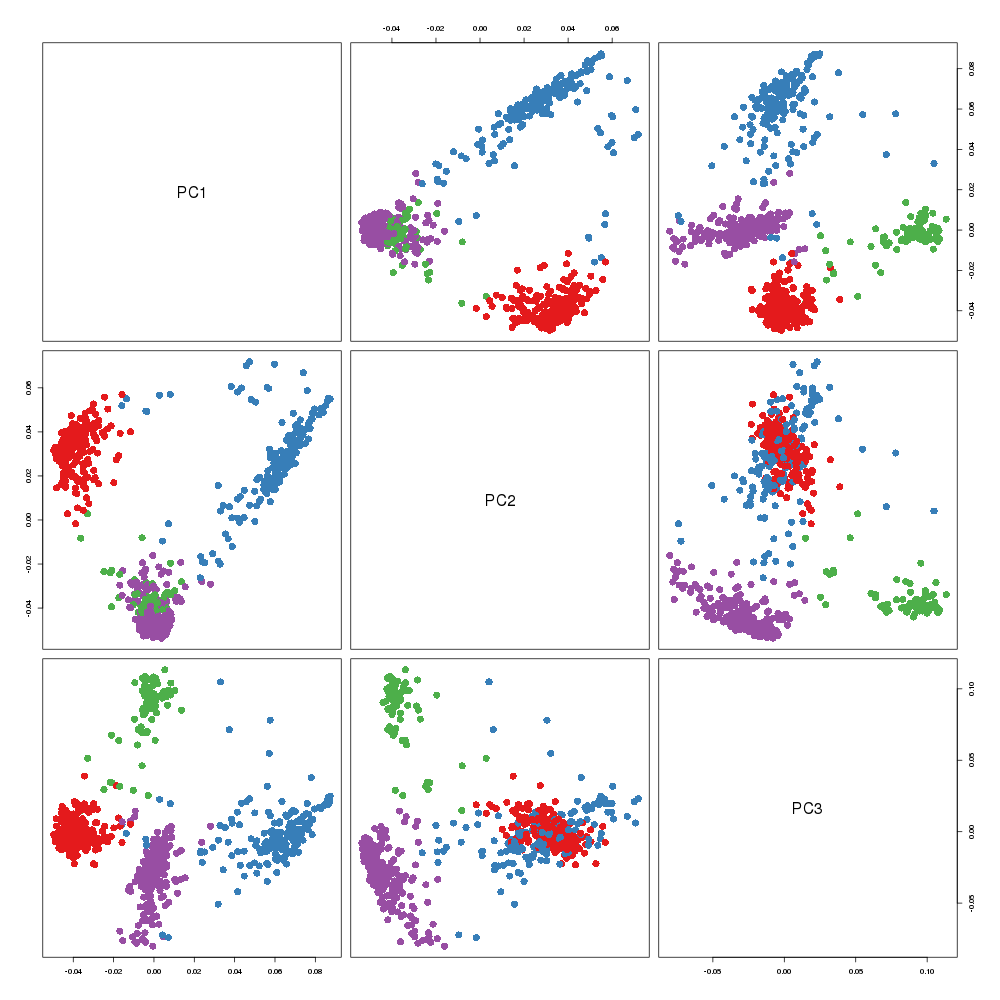

### oocRPCA.png

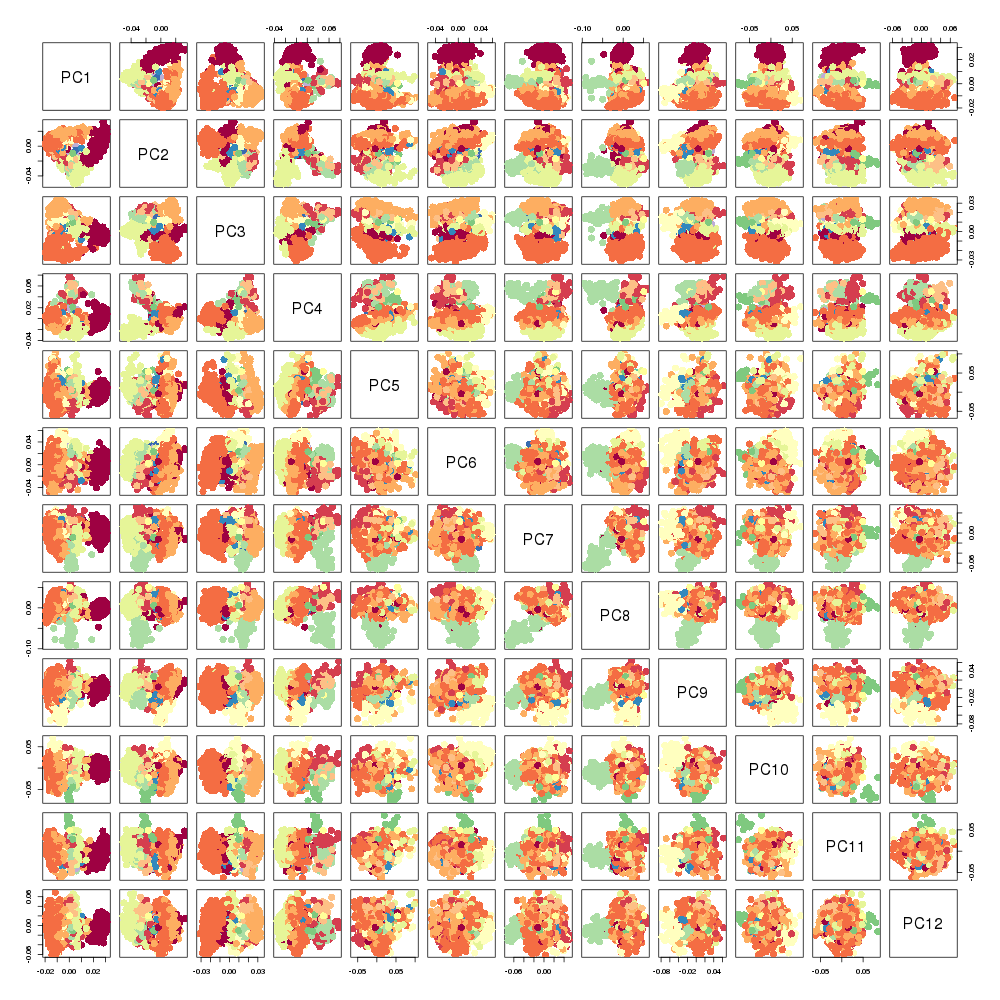

### oocRPCA.png

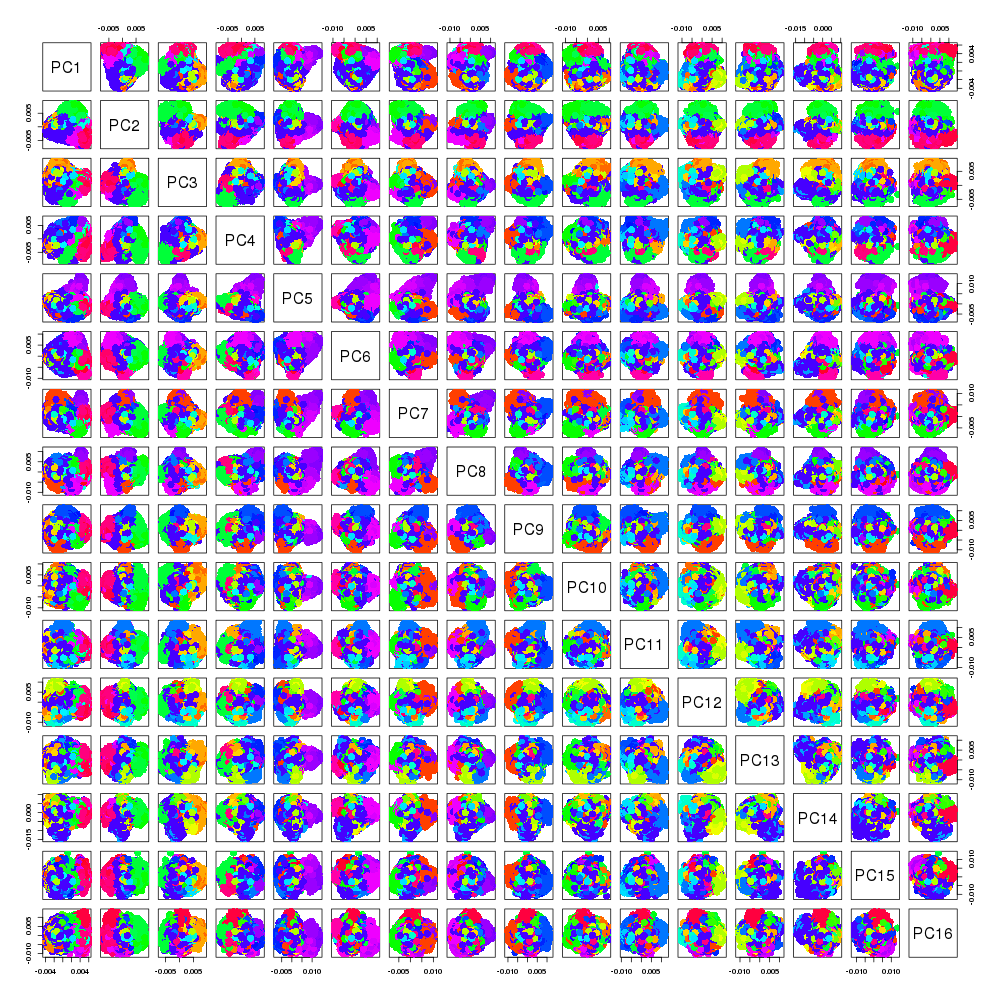

### oocRPCA.png

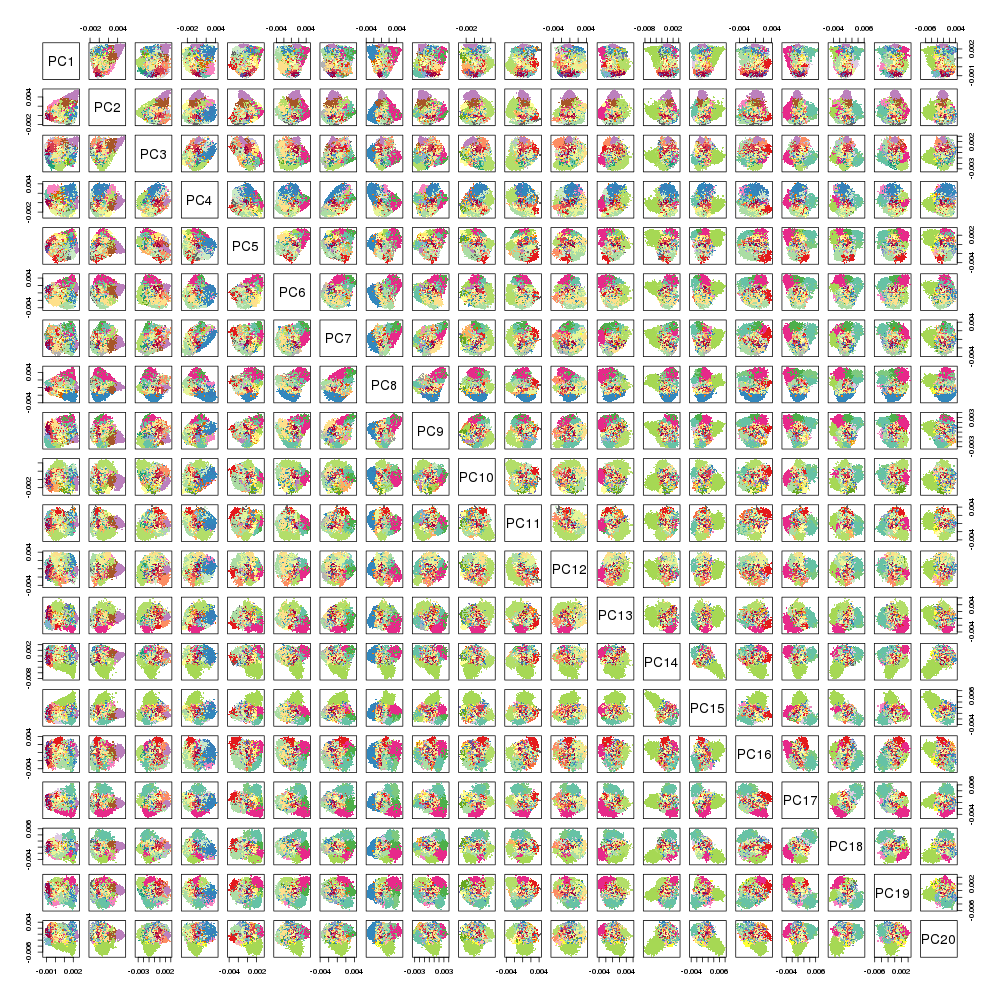

### OrthIter.png

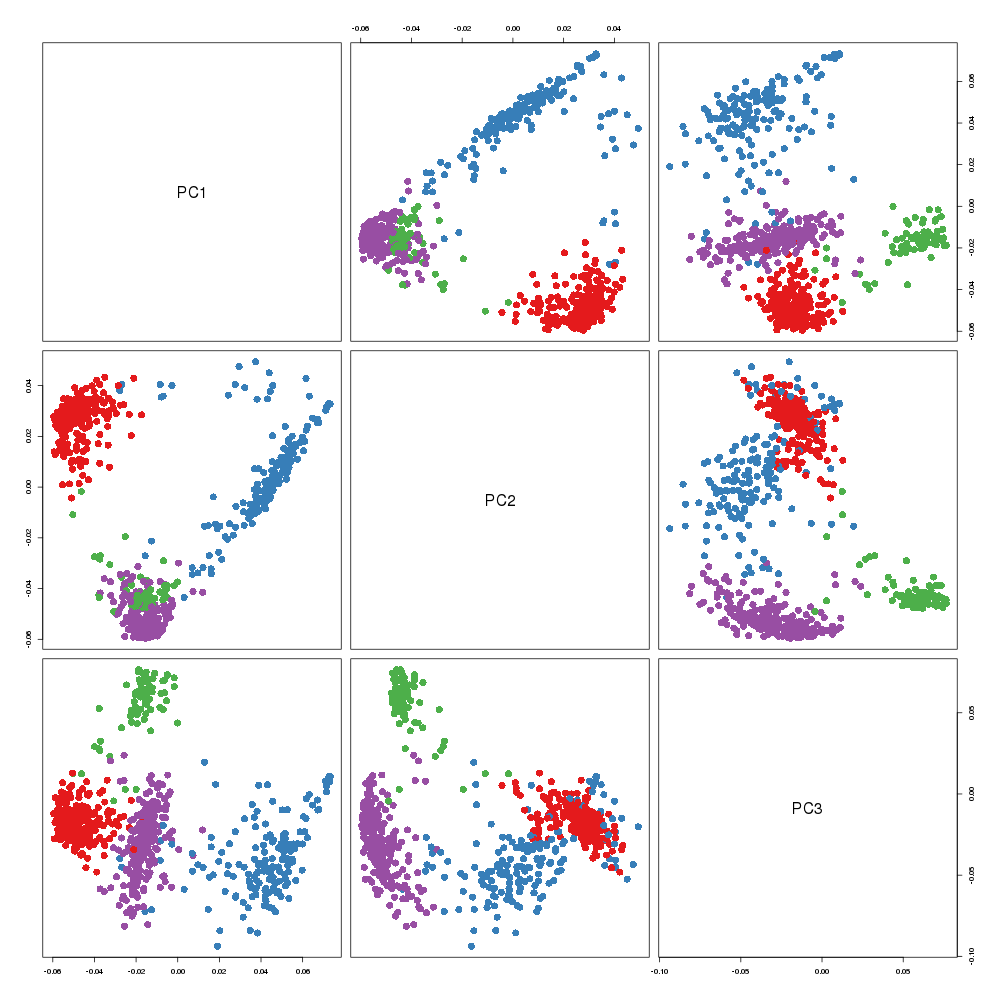

### OrthIter.png

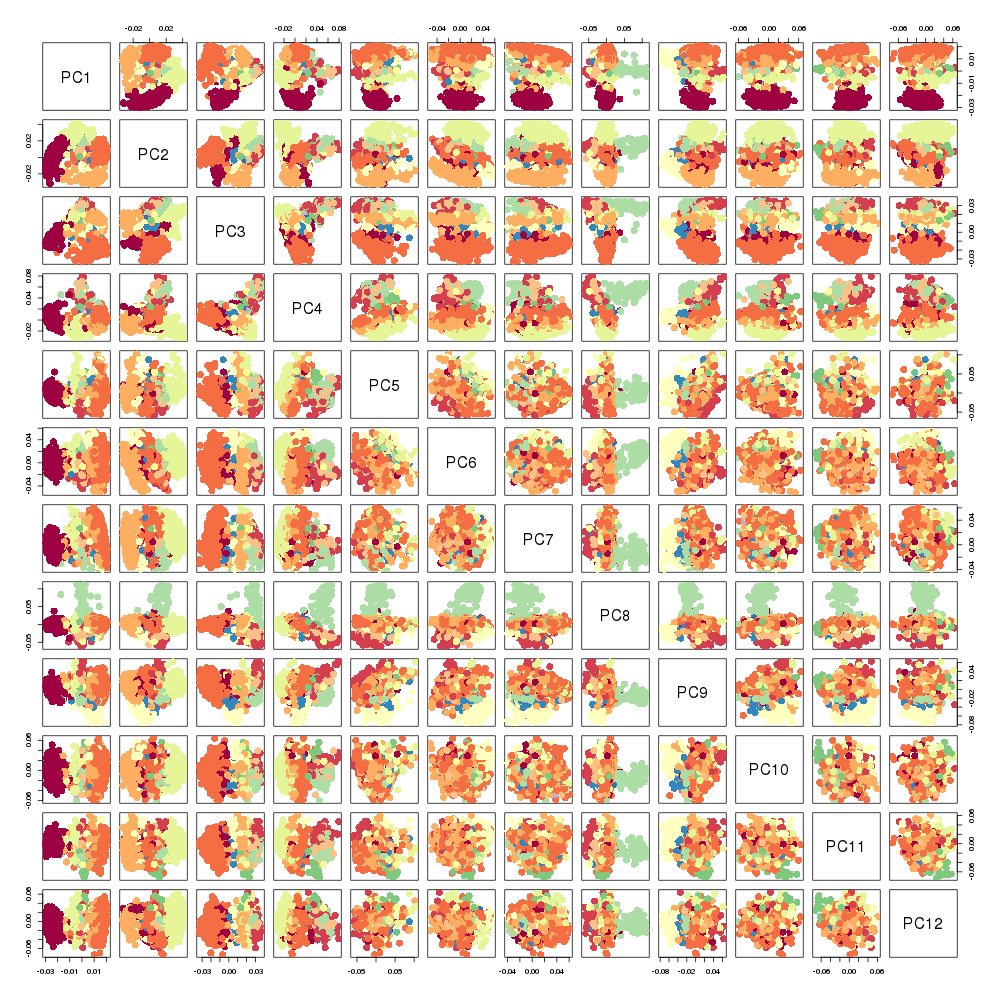

### OrthIter.png

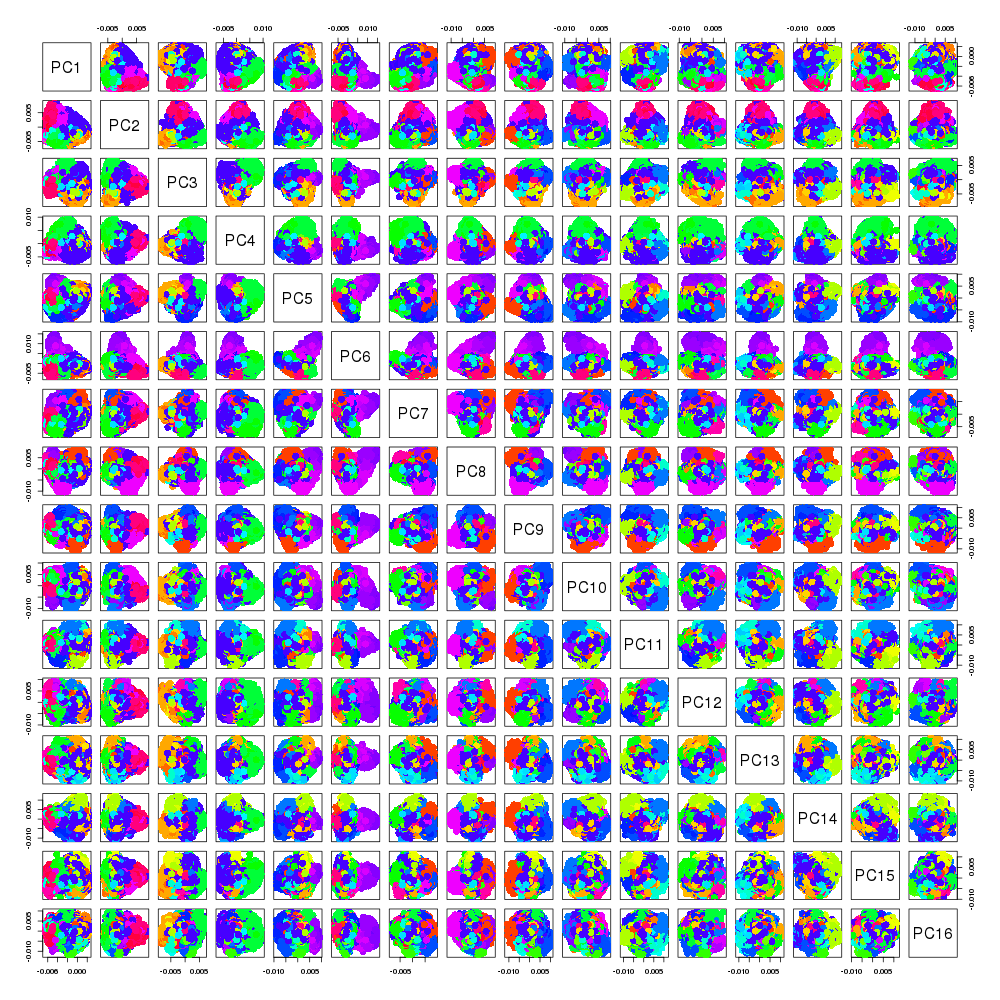

### OrthIter.png

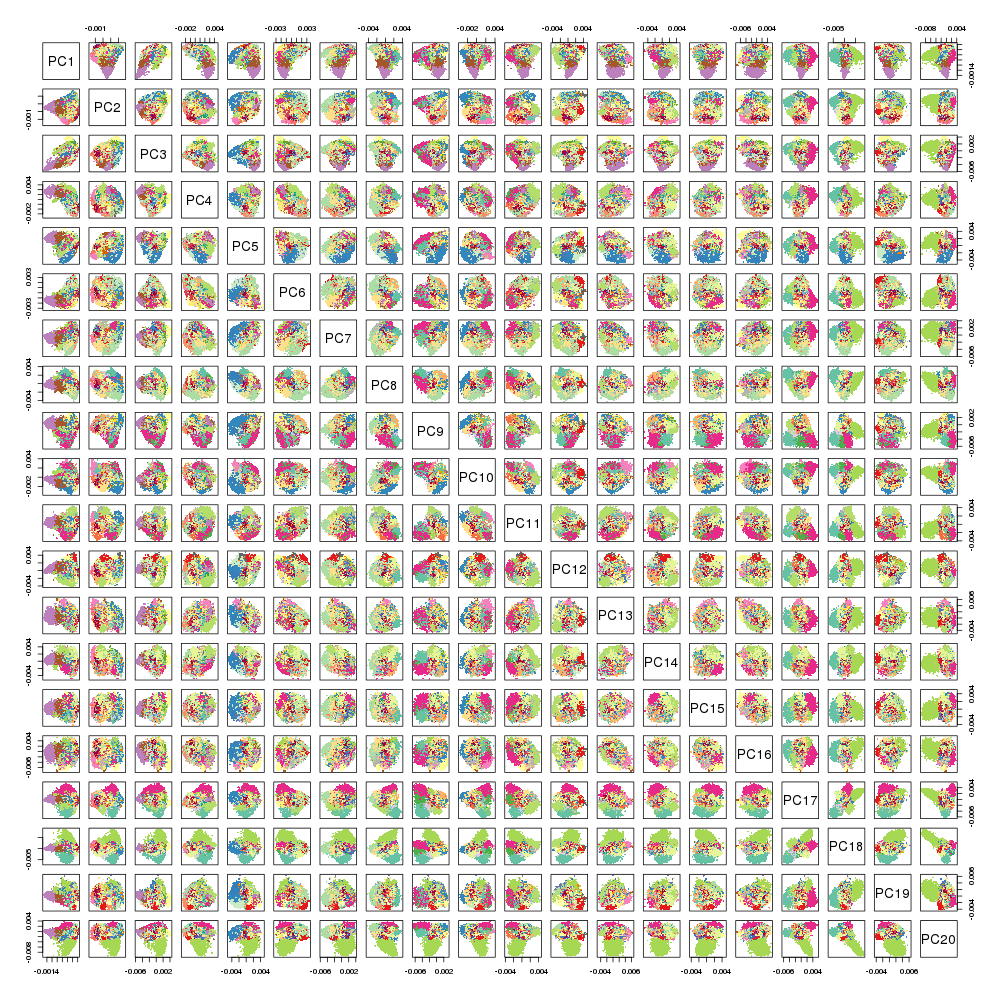

### PCA.png

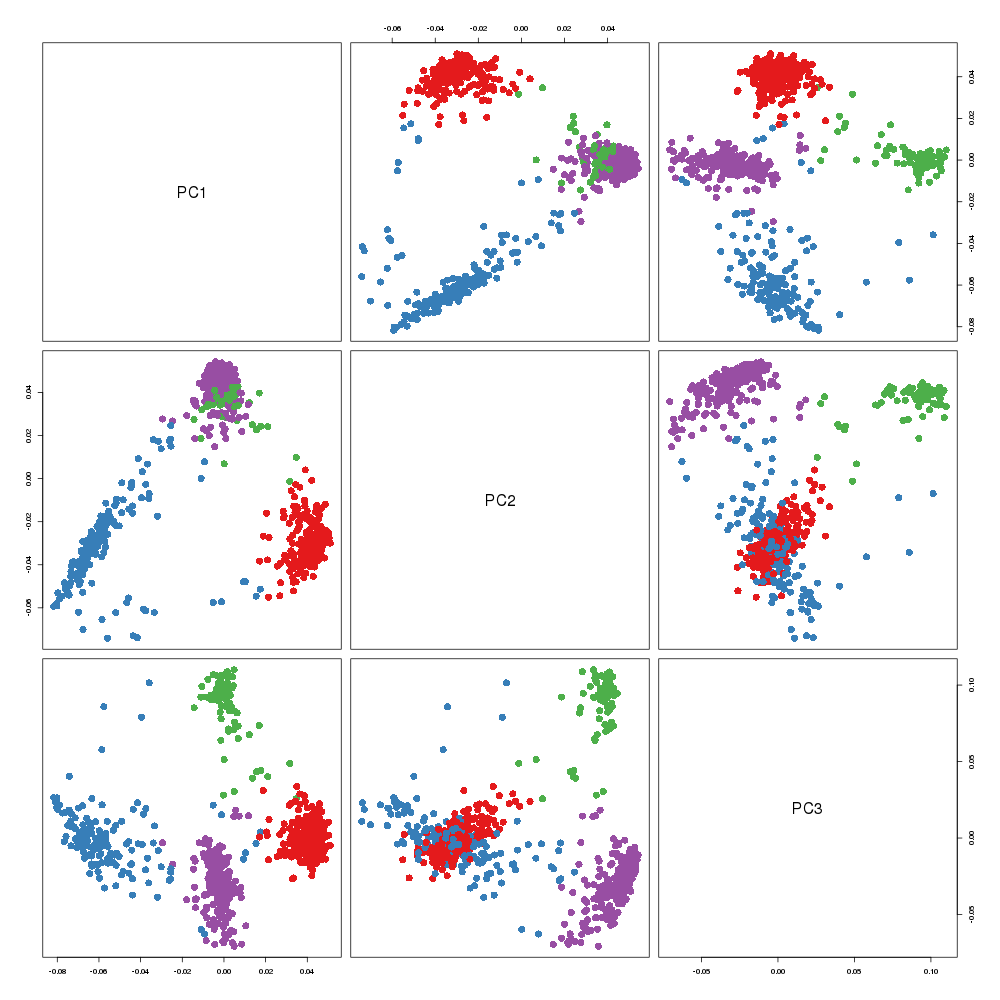

### PCA.png

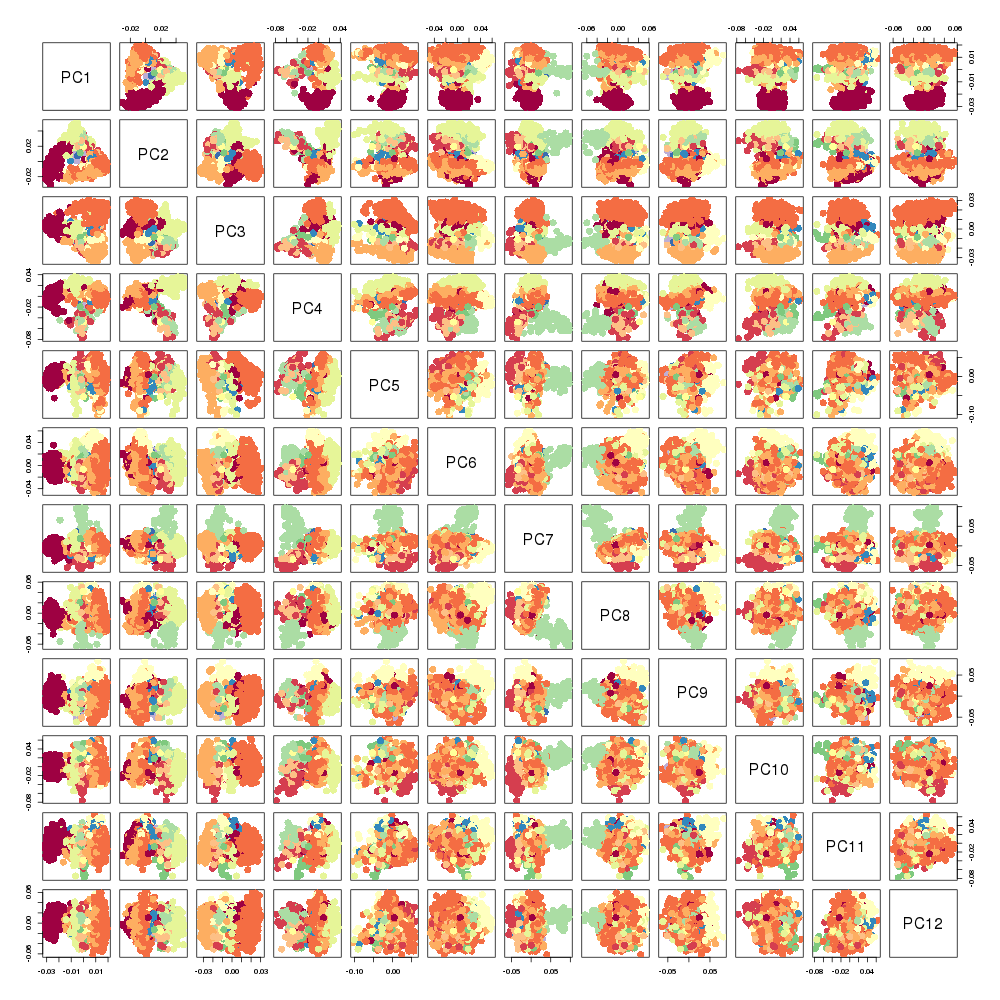

### PROPACK.png

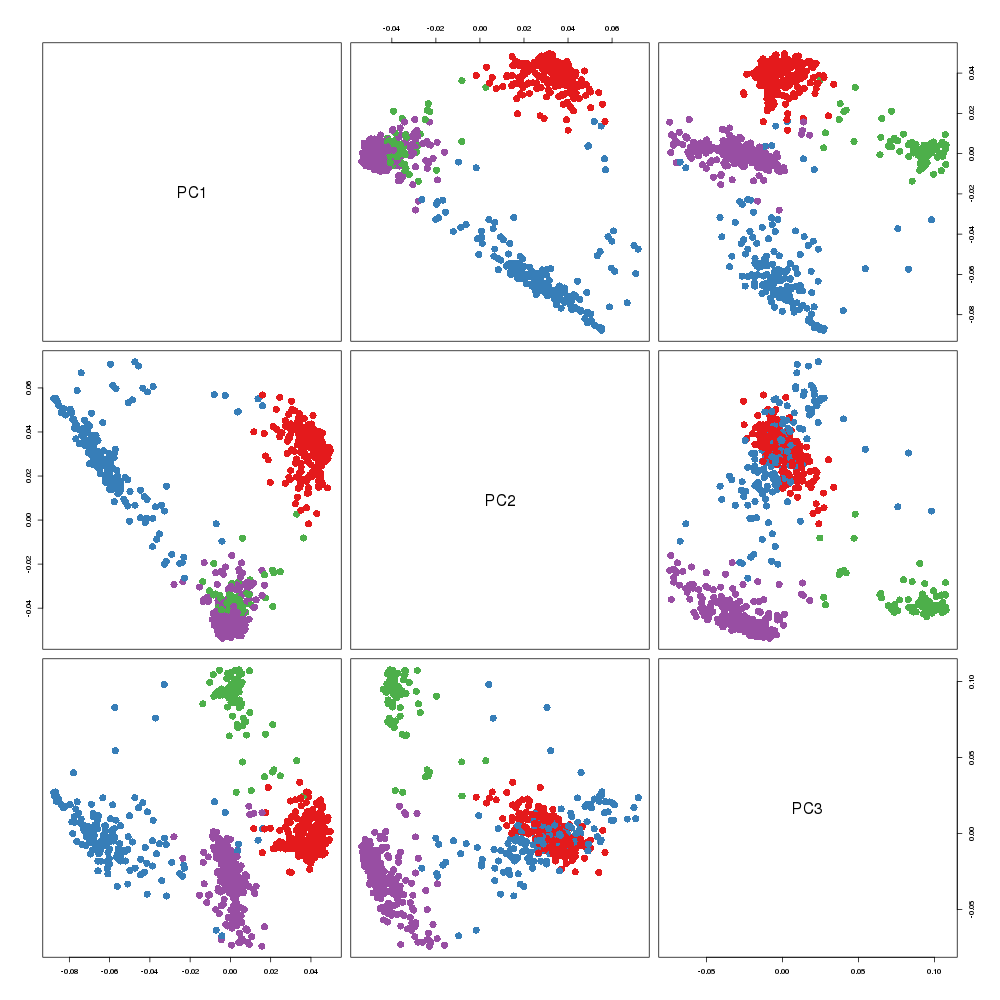

### PROPACK.png

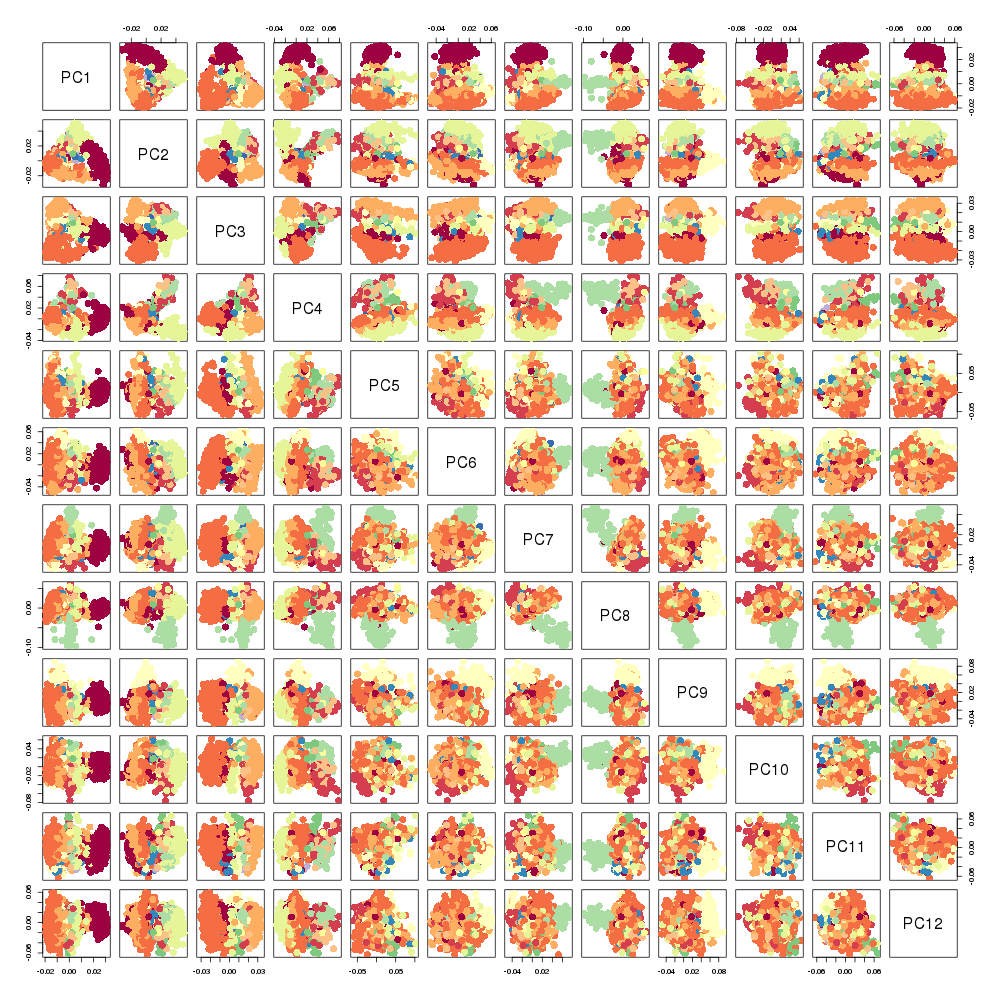

### RSpectra.png

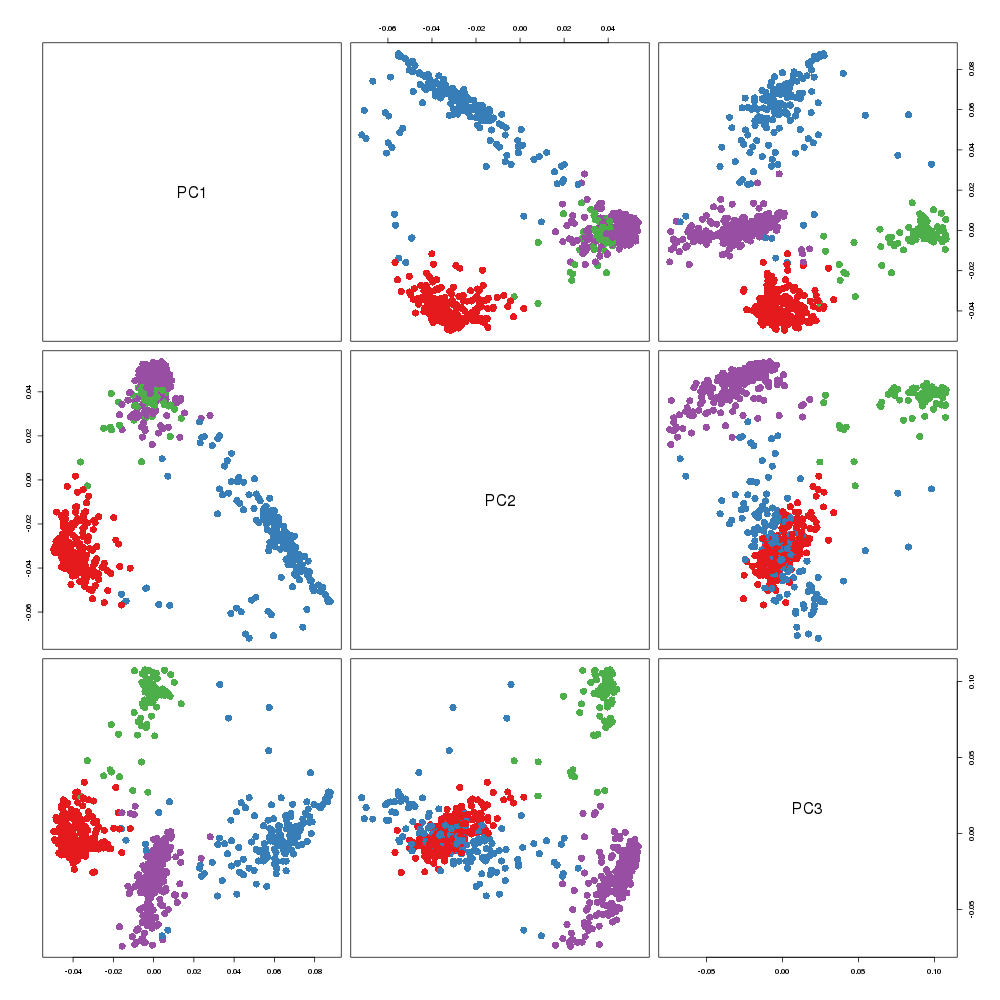

### RSpectra.png

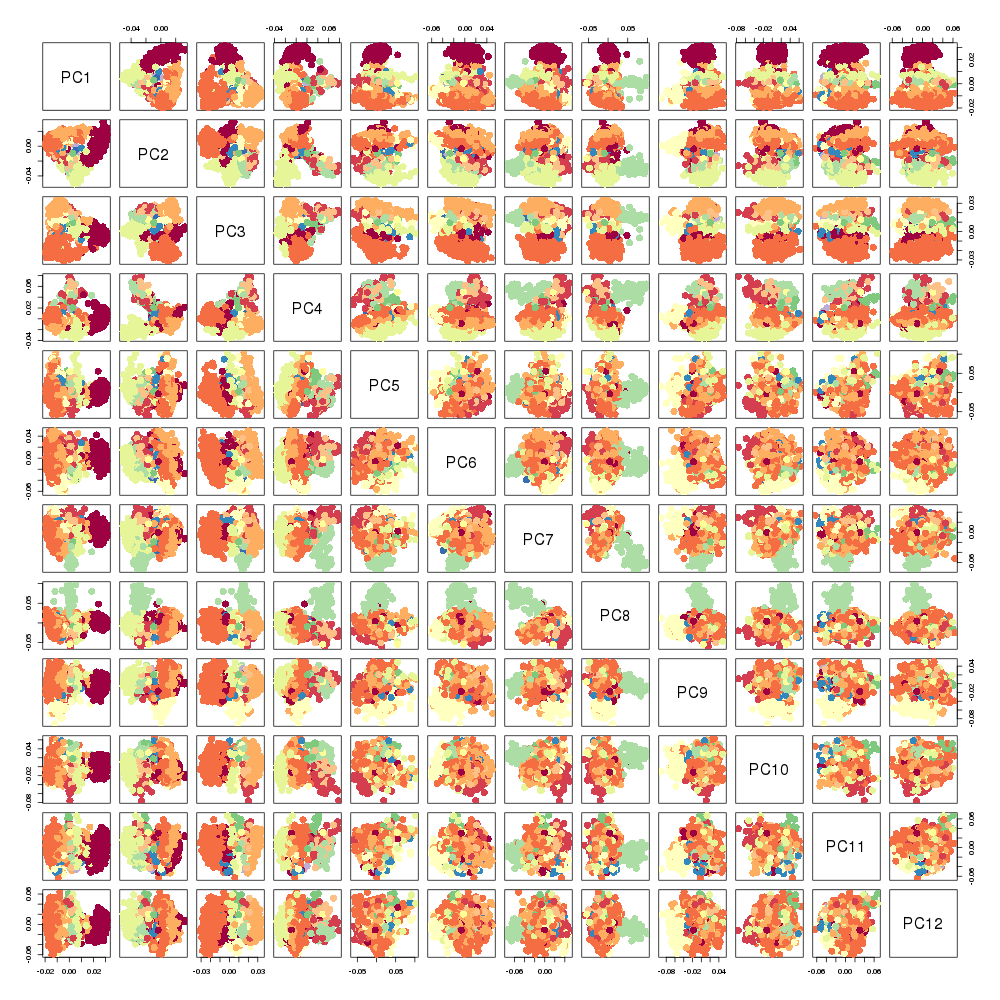

### rsvd.png

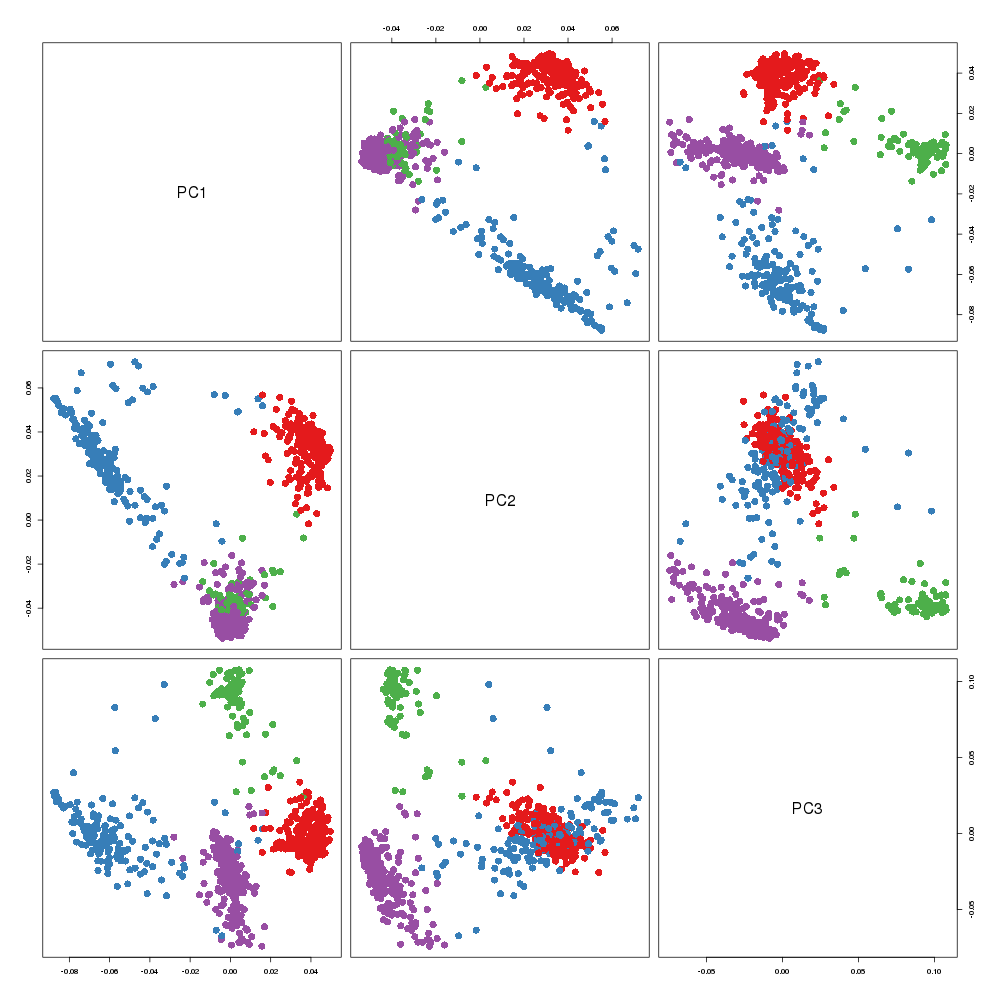

### rsvd.png

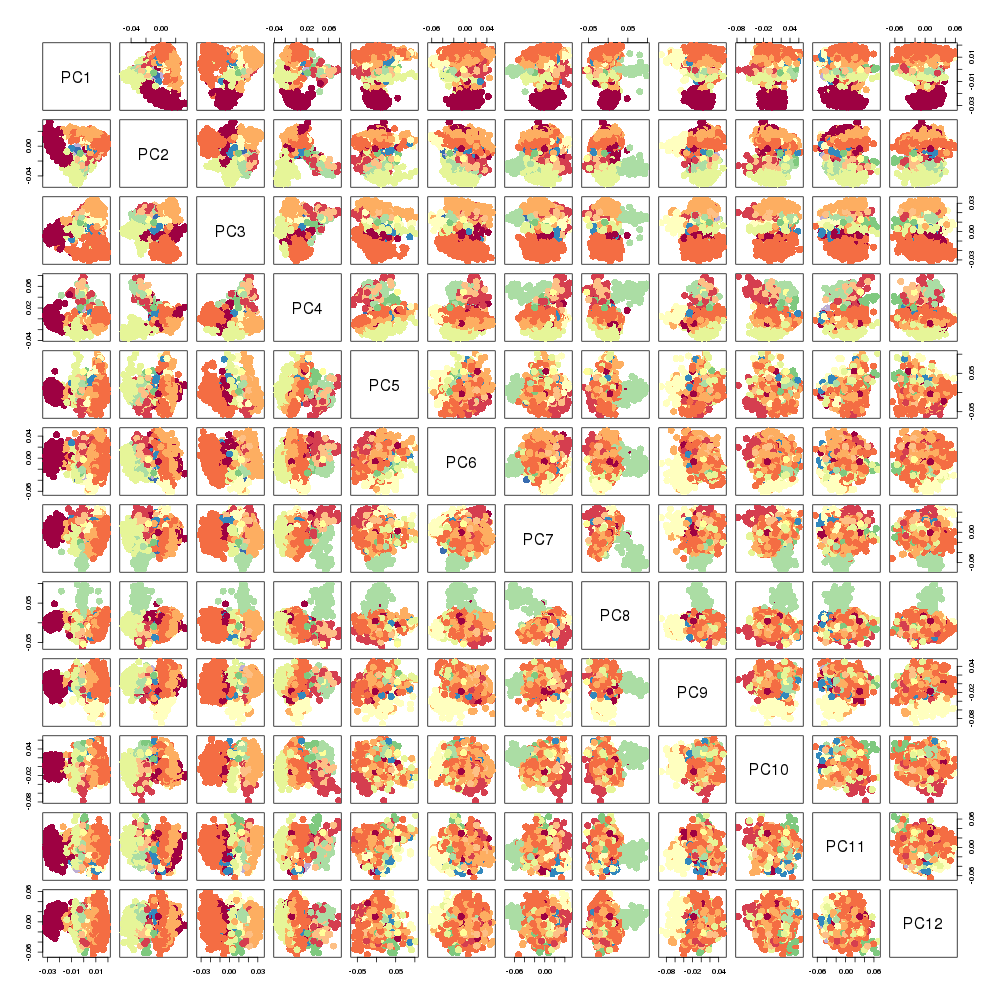

### SGD_step100_epoch10.png

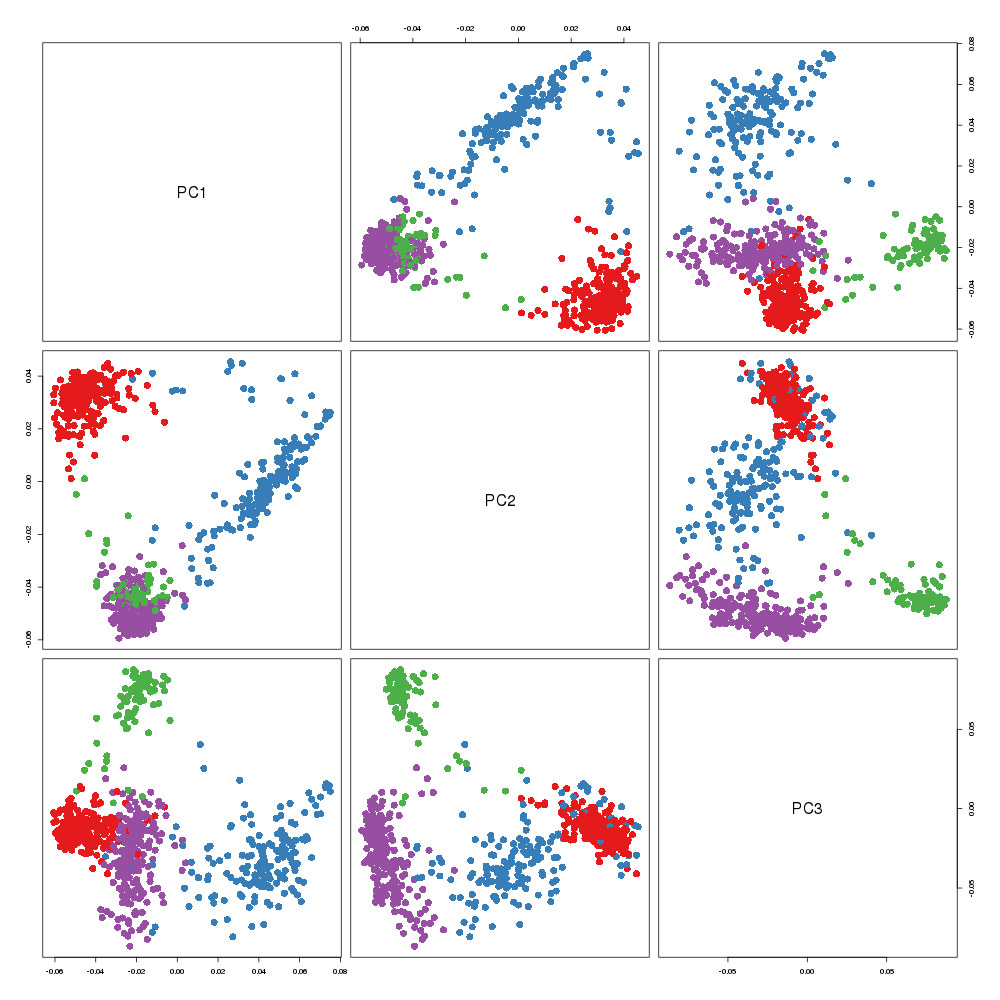

### SGD_step100_epoch10.png

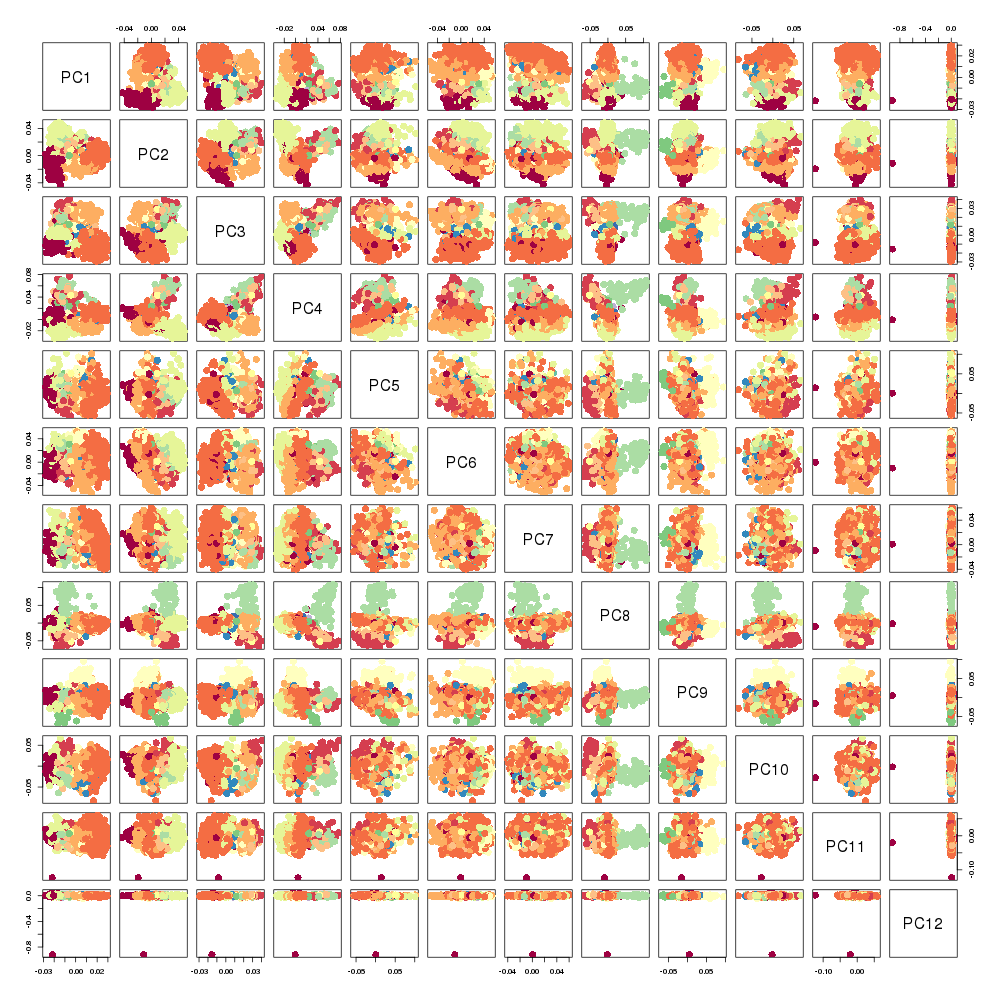

### SGD_step100_epoch10.png

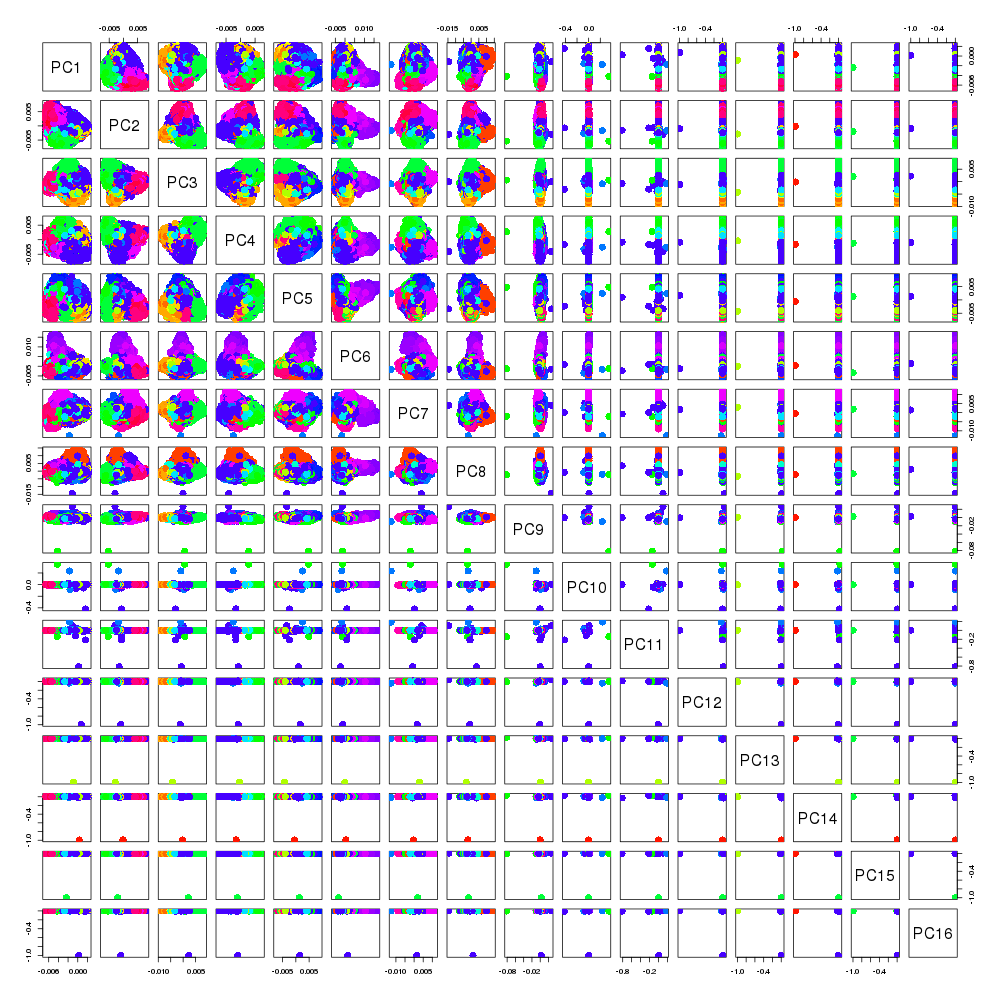

### SGD_step100_epoch10.png

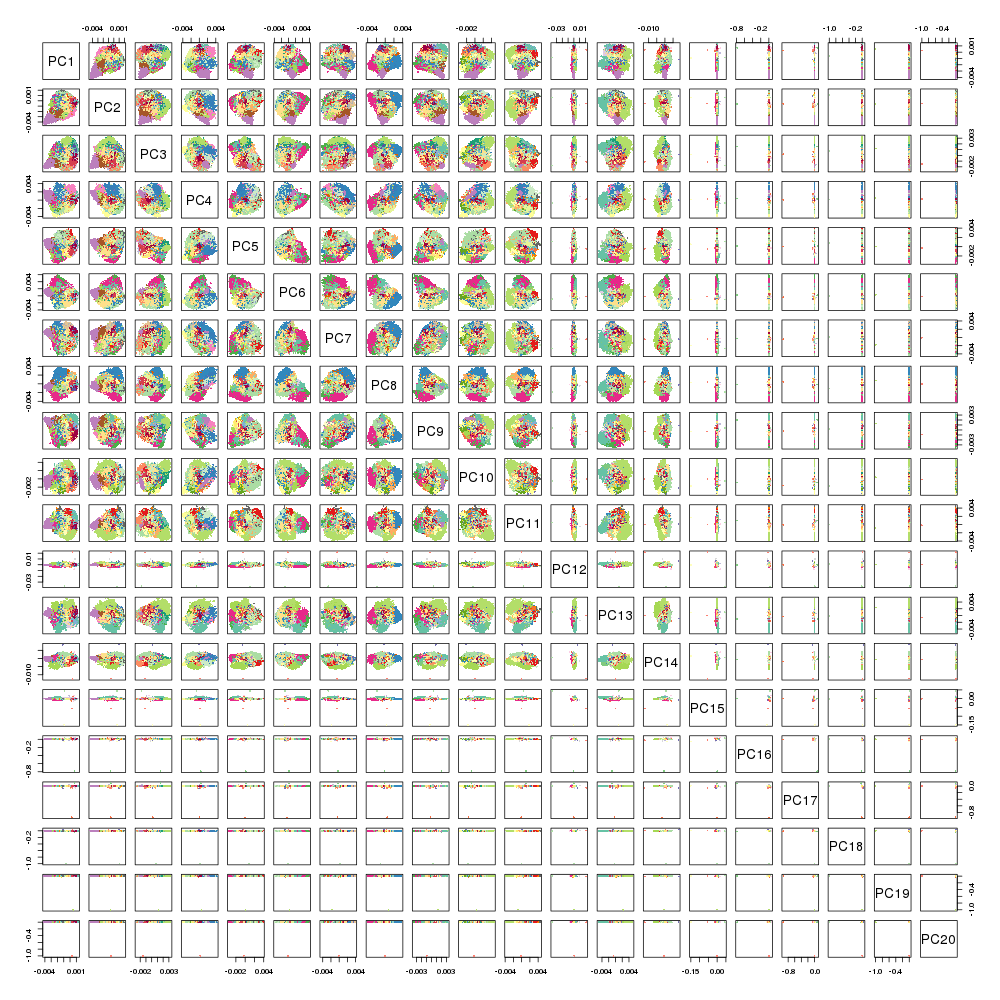

### Sklearn_ARPACK.png

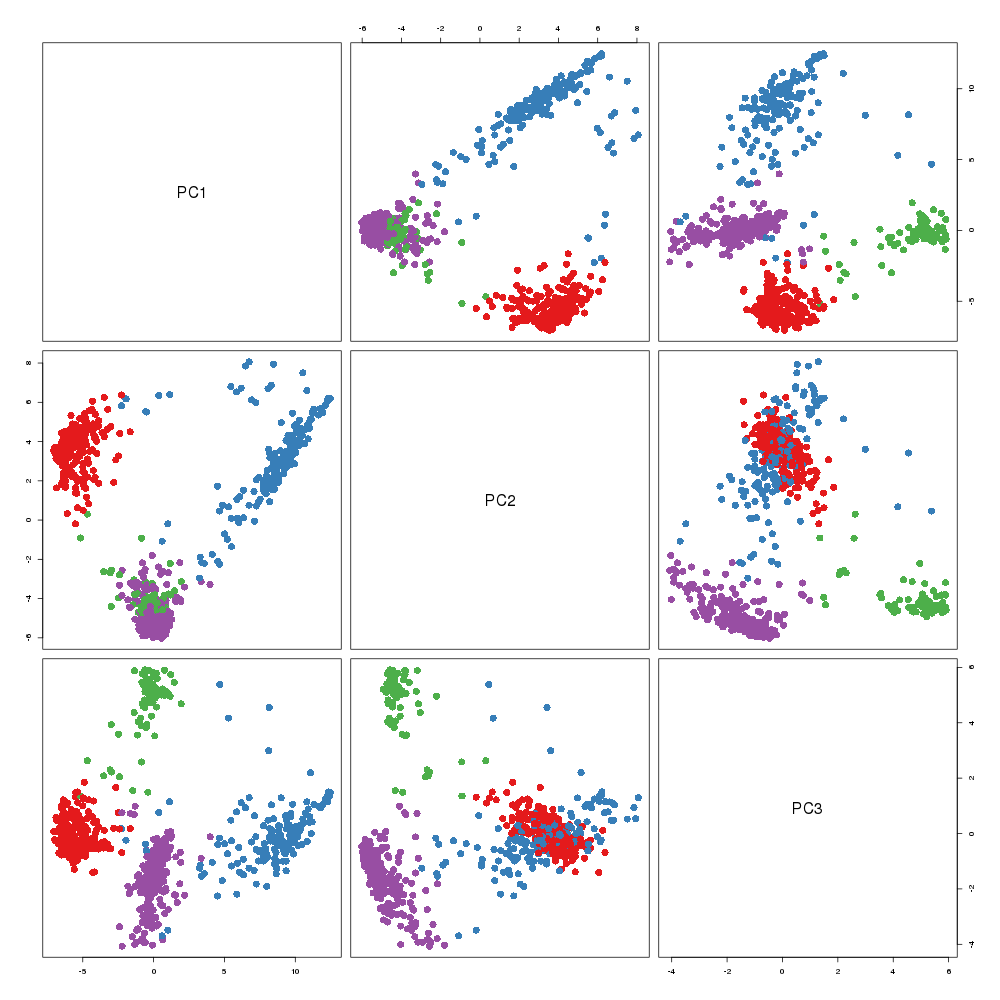

### Sklearn_ARPACK.png

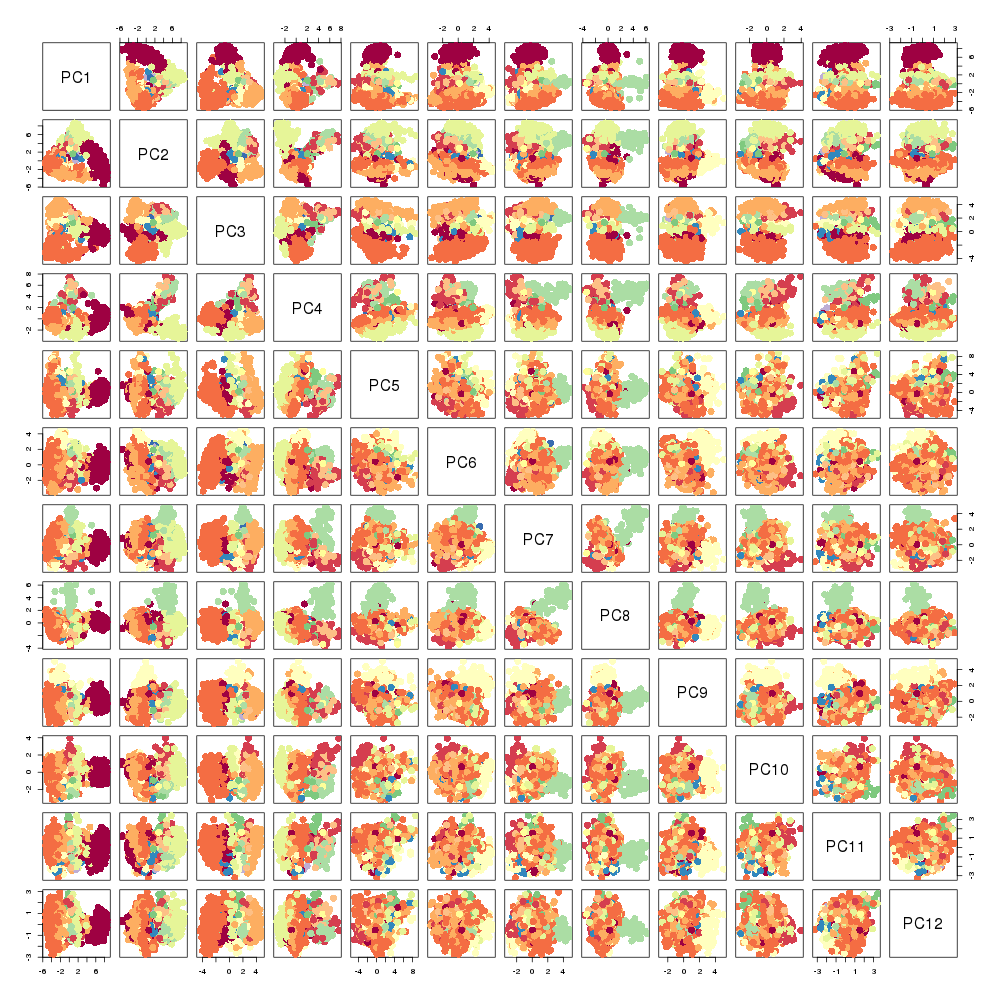

### Sklearn_Incremental.png

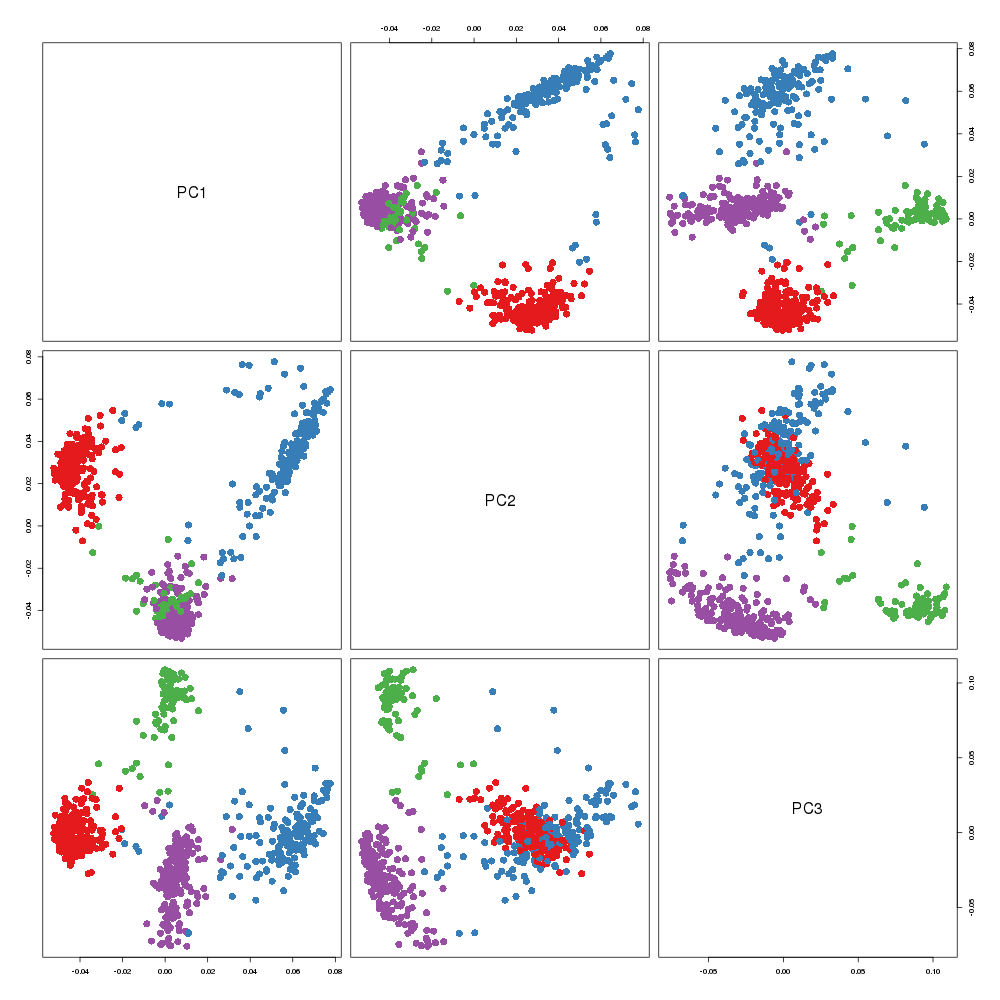

### Sklearn_Incremental.png

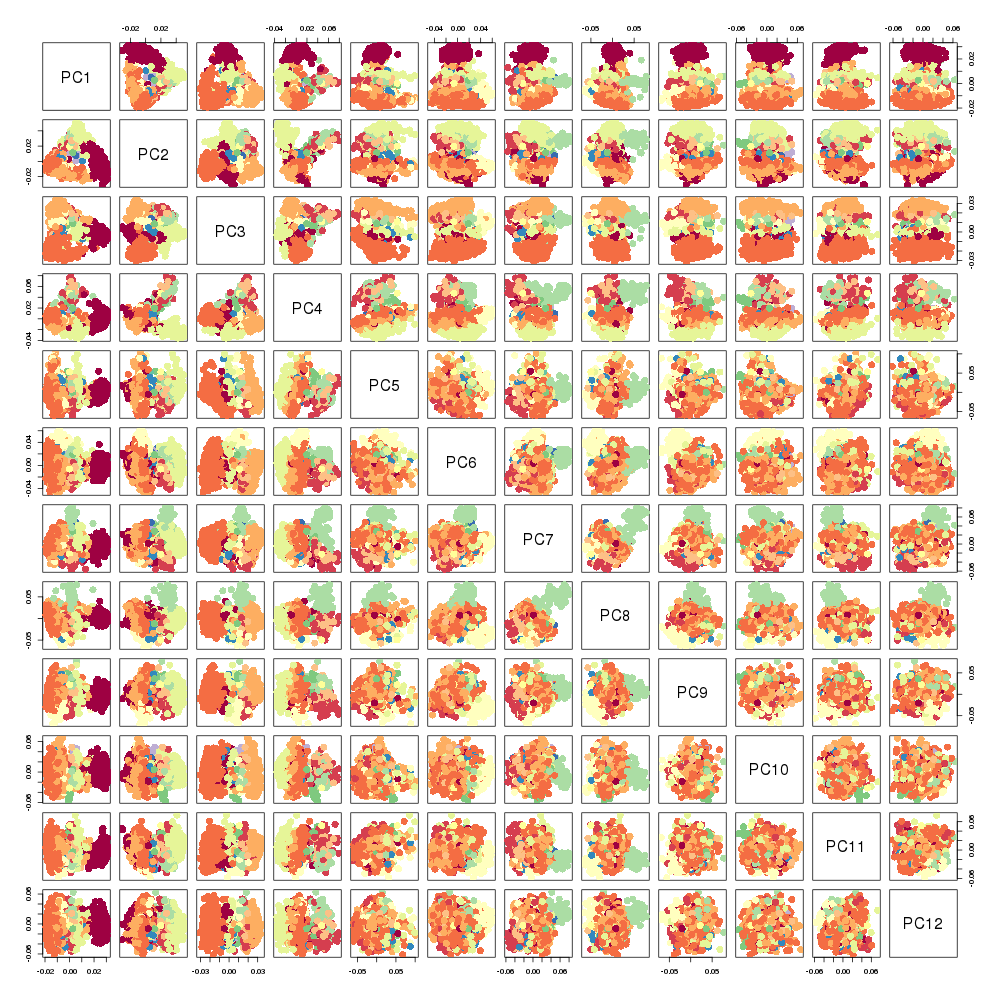

### Sklearn_Incremental.png

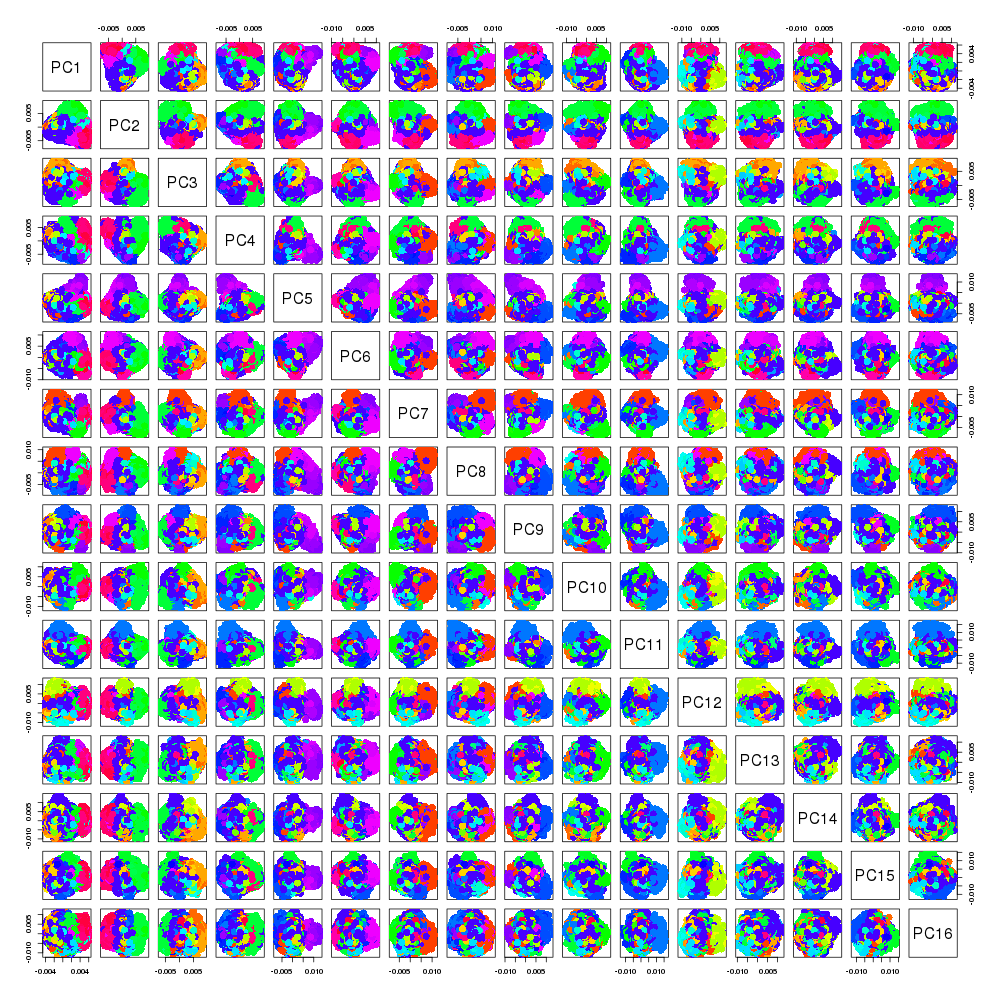

### Sklearn_Incremental.png

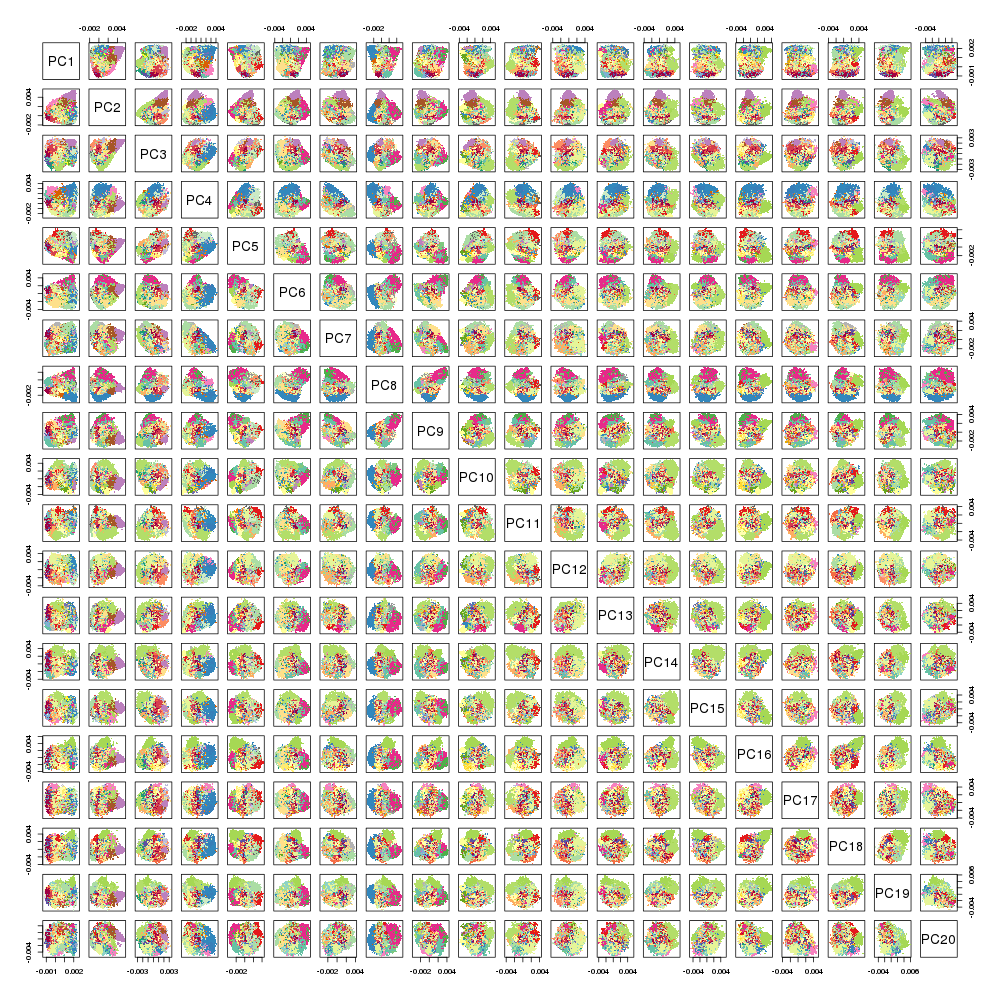

### Sklearn_randomized_svd.png

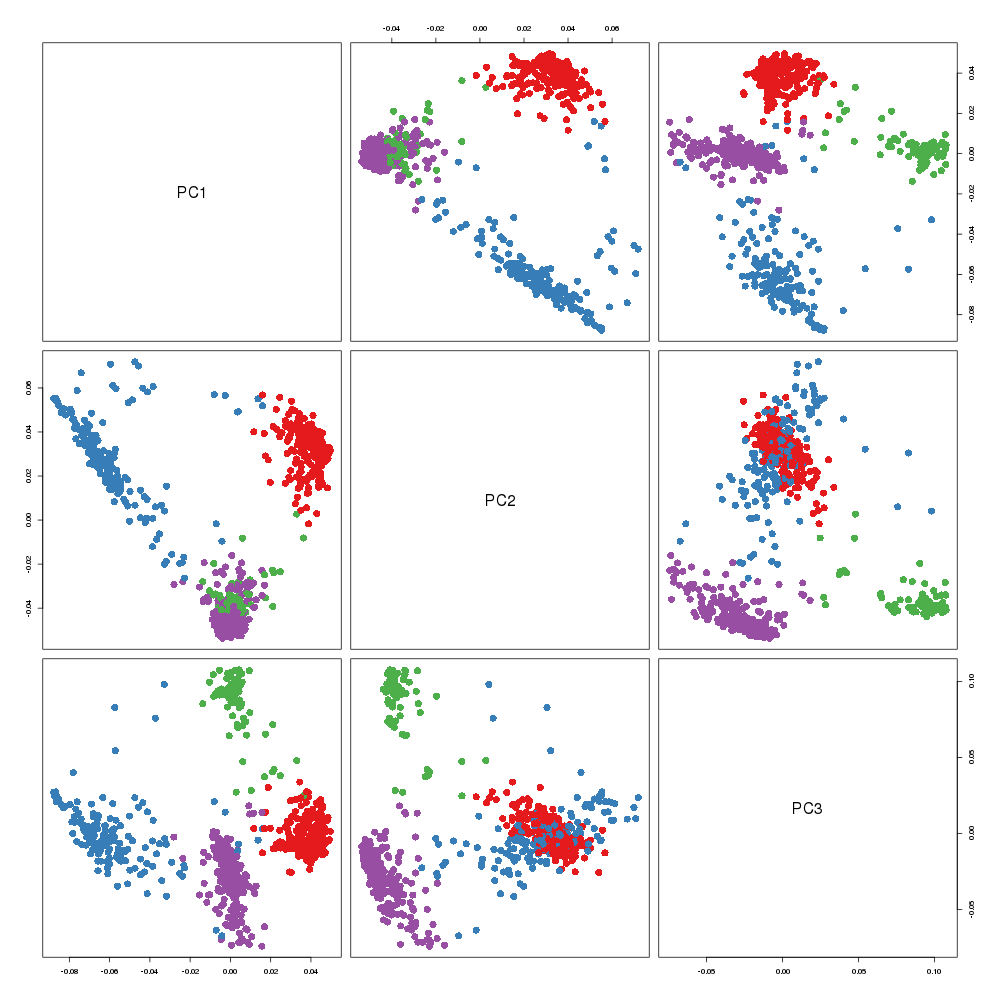

### Sklearn_randomized_svd.png

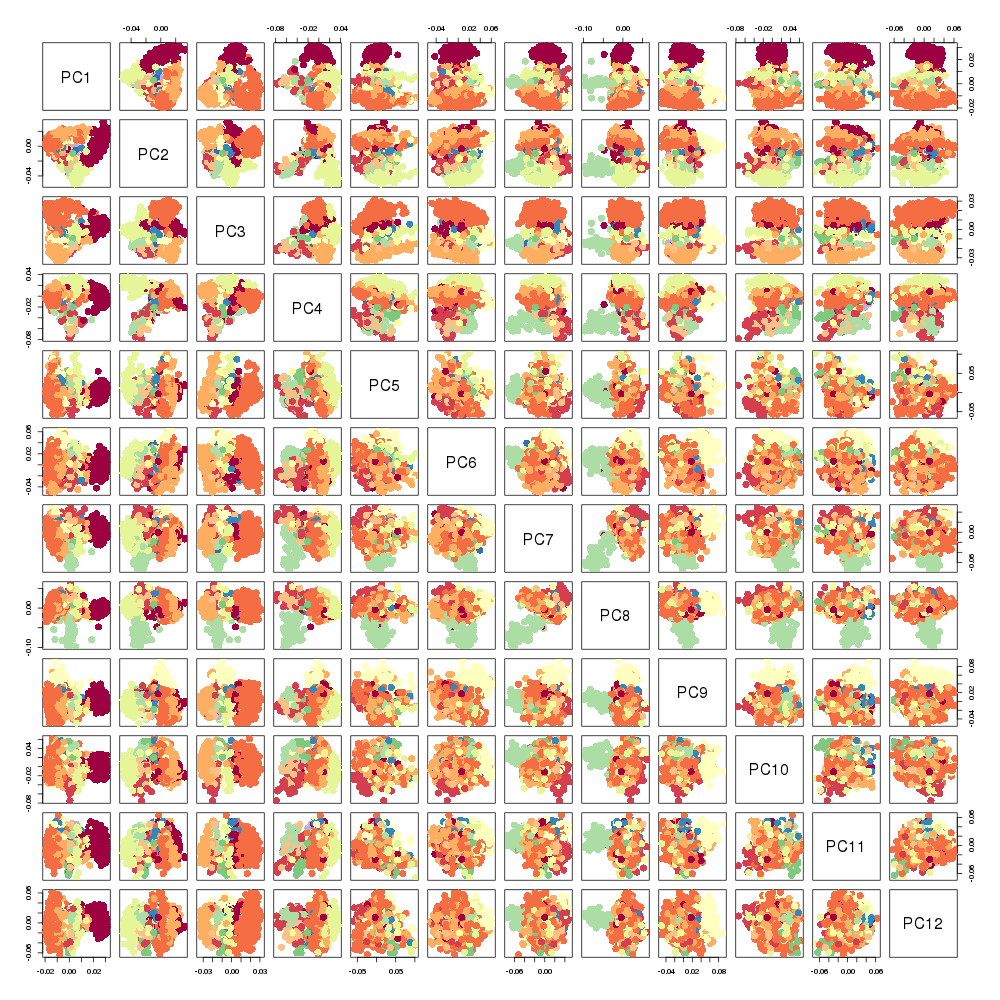

### Sklearn_RandomizedPCA.png

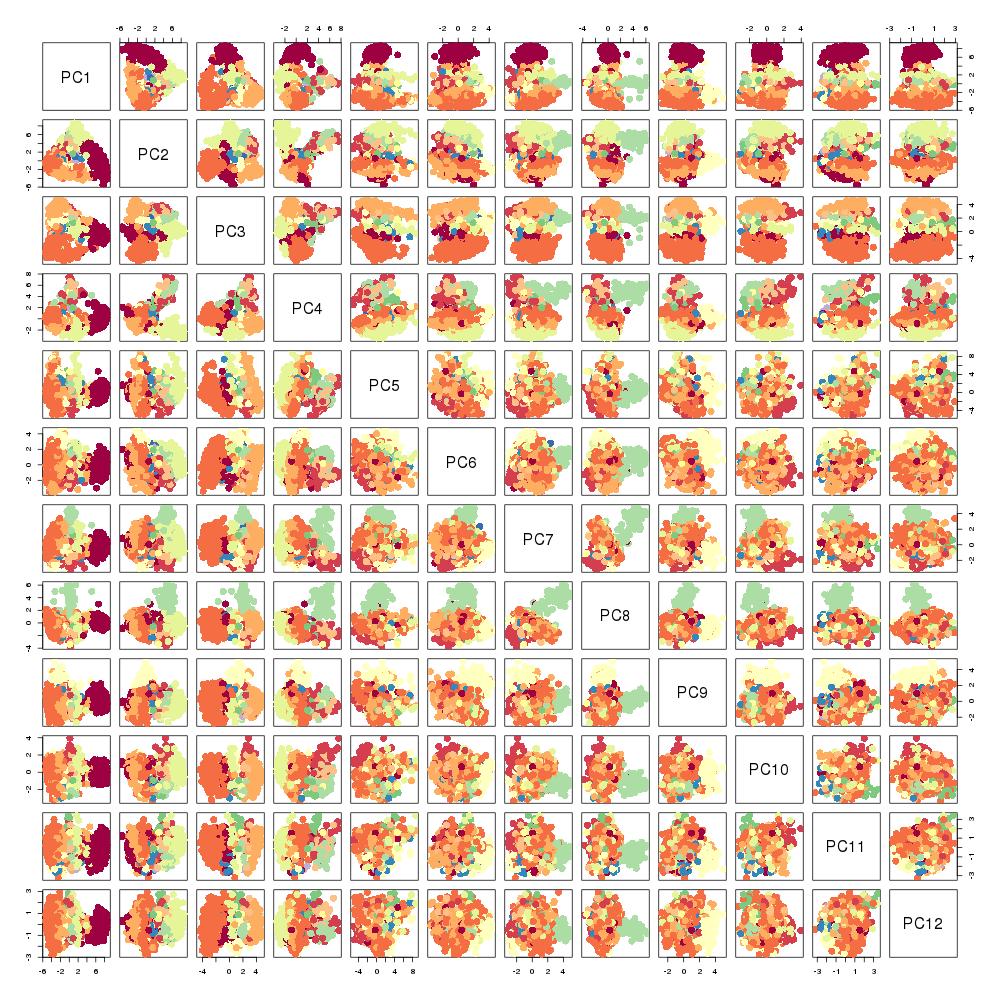
