## Additional File 6 for "Benchmarking principal component analysis for large-scale single-cell RNA-sequencing"

### Results of clustering methods for all PCA implementations

Here, we present the results of clustering performed for all the PCA implementations mentioned in the manuscript: *k*-means, Gaussian mixture model (GMM), HDBScan, and Louvain clustering. When performing HDBScan, principal components (PCs) were first converted to five dimensional UMAP coordinates, and then HDBScan was performed (**Figure S6-1** to **7**). When performing Louvain clustering, the Jaccard index was used as the similarity measure with the *k* nearest neighborhood cells. For the PBMCs, Pancreas, BrainSpinalCord, and Brain datasets, *k* values of 100, 75, 10, and 5 were used, respectively.

The cluster assignment shown in **Figure S6-1** to **7** and the corresponding adjusted Rand index (ARI) values (**Figure S6-8**) show that the Louvain clustering results were stable across all datasets. Therefore, we finally selected Louvain clustering as the measure for benchmarking.

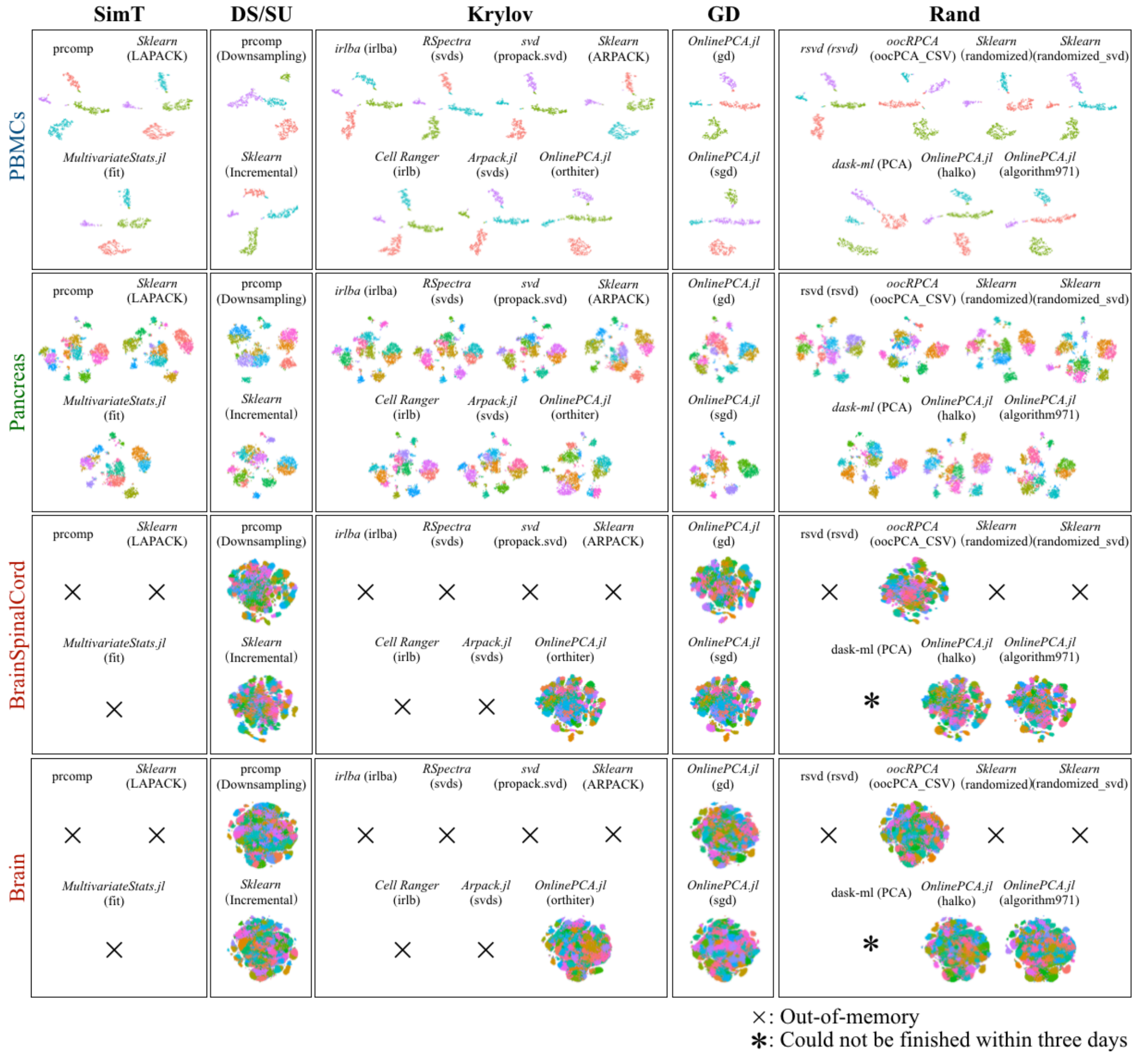


**Supplementary Figure 6-1 | Results of *k*-means clustering** The cluster label is visualized with two-dimensional t-SNE coordinates.

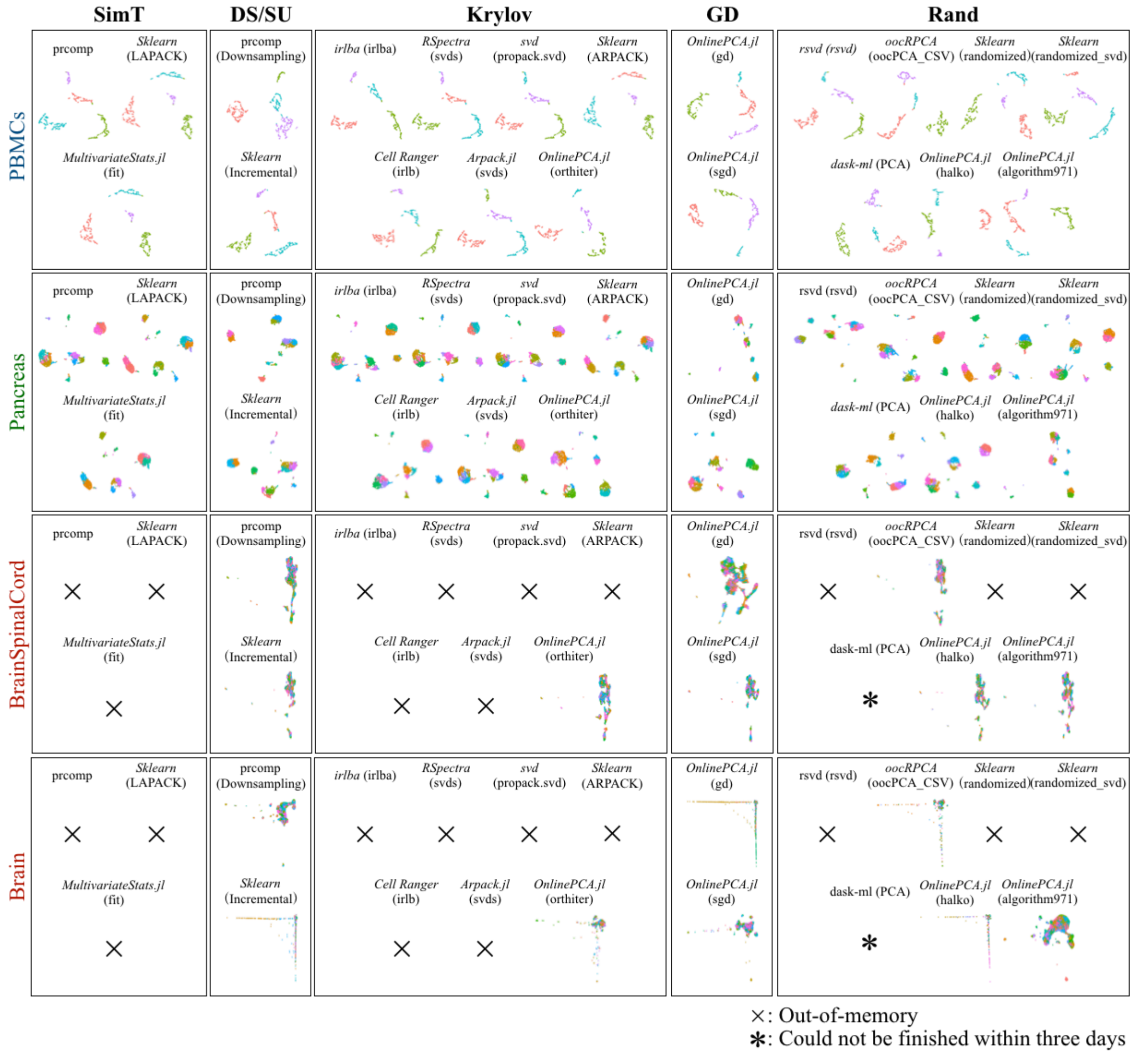


**Supplementary Figure 6-2 | Results of *k*-means clustering** The cluster label is visualized with two-dimensional UMAP coordinates.

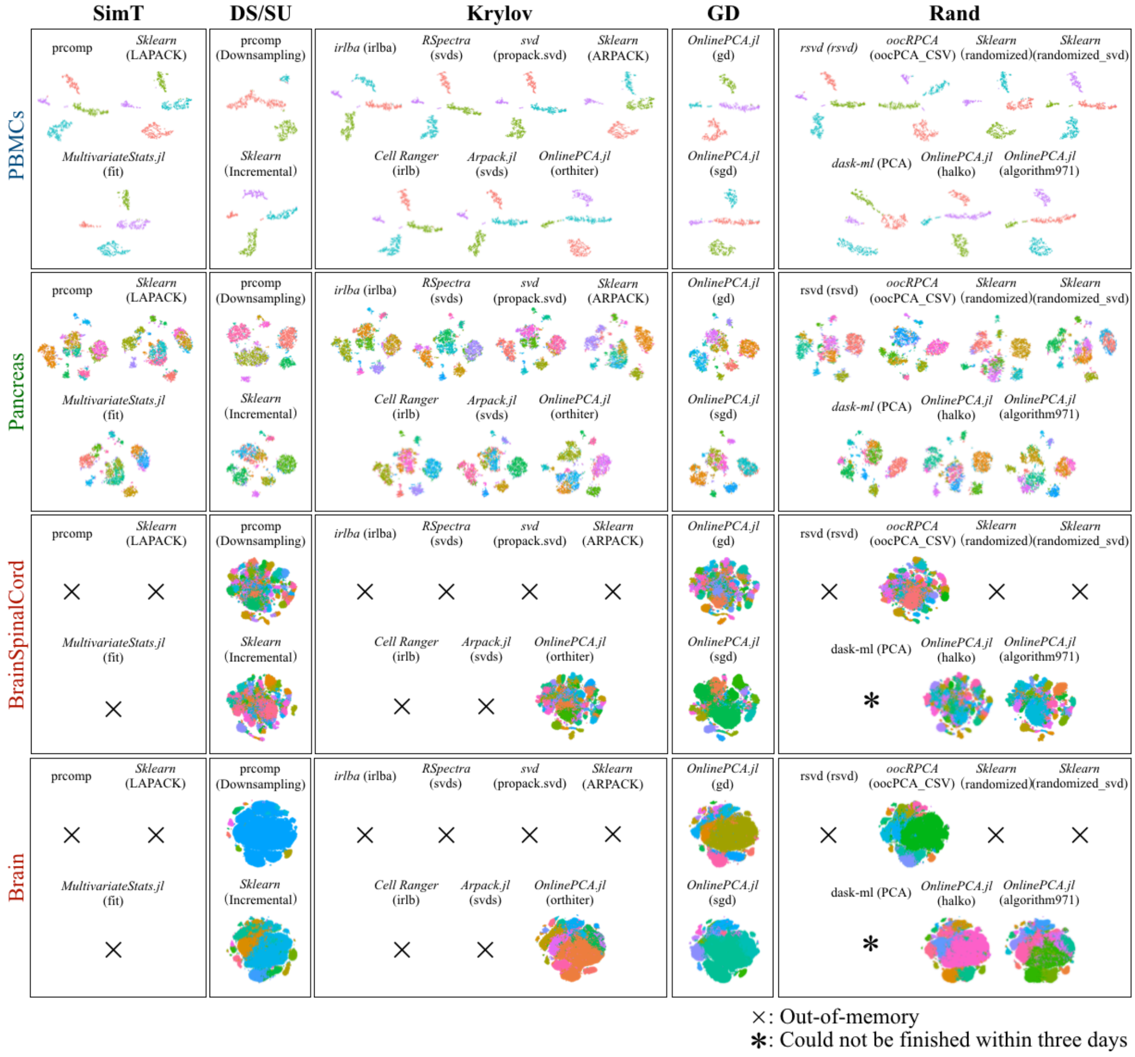


**Supplementary Figure 6-3 | Results of GMM clustering** The cluster label is visualized with two-dimensional t-SNE coordinates.

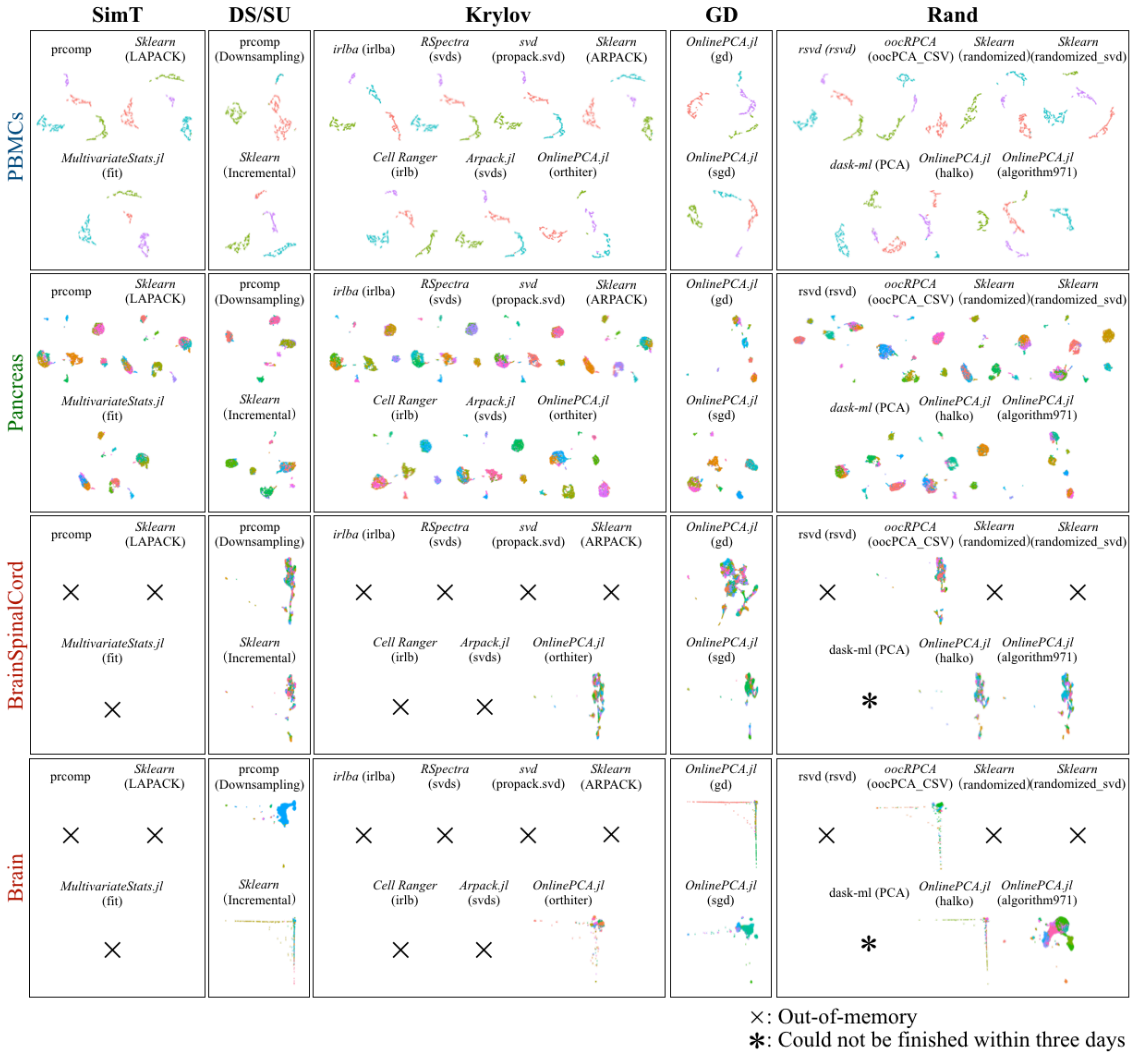


**Supplementary Figure 6-4 | Results of GMM clustering** The cluster label is visualized with two-dimensional UMAP coordinates.

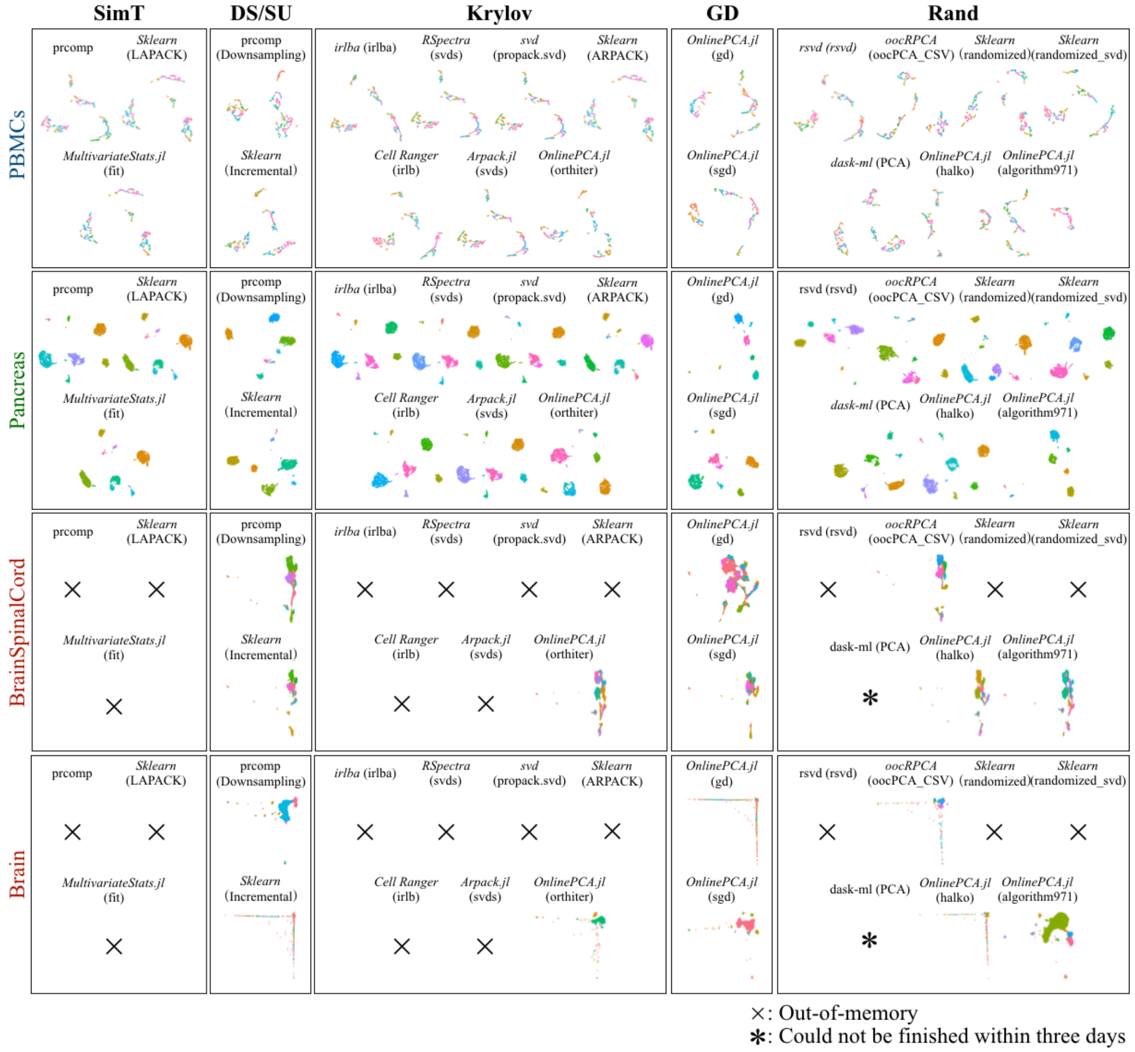


**Supplementary Figure 6-5 | Results of HDBScan clustering** The cluster label is visualized with two-dimensional UMAP coordinates.

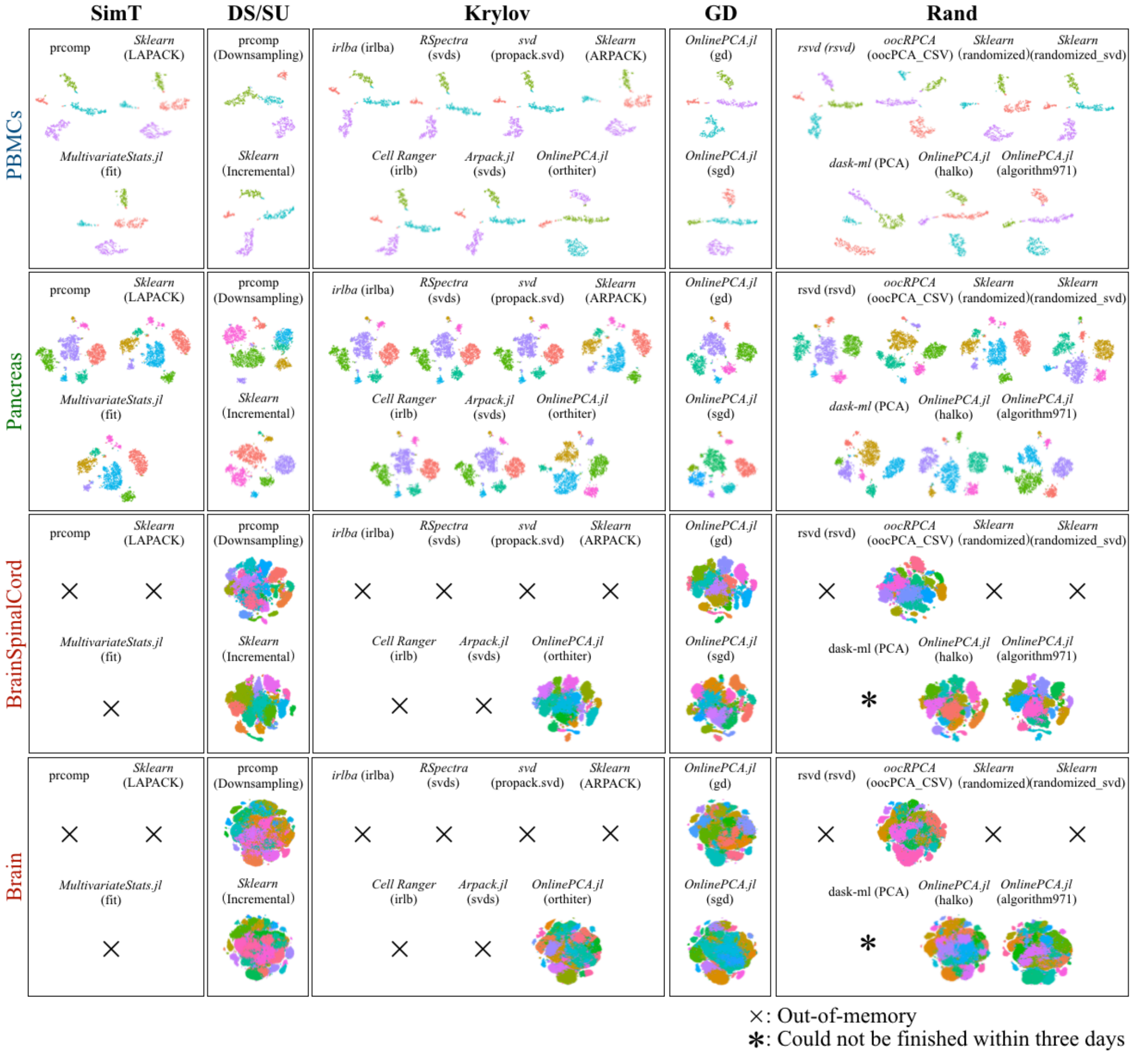


**Supplementary Figure 6-6 | Results of Louvain clustering** The cluster label is visualized with two-dimensional t-SNE coordinates.

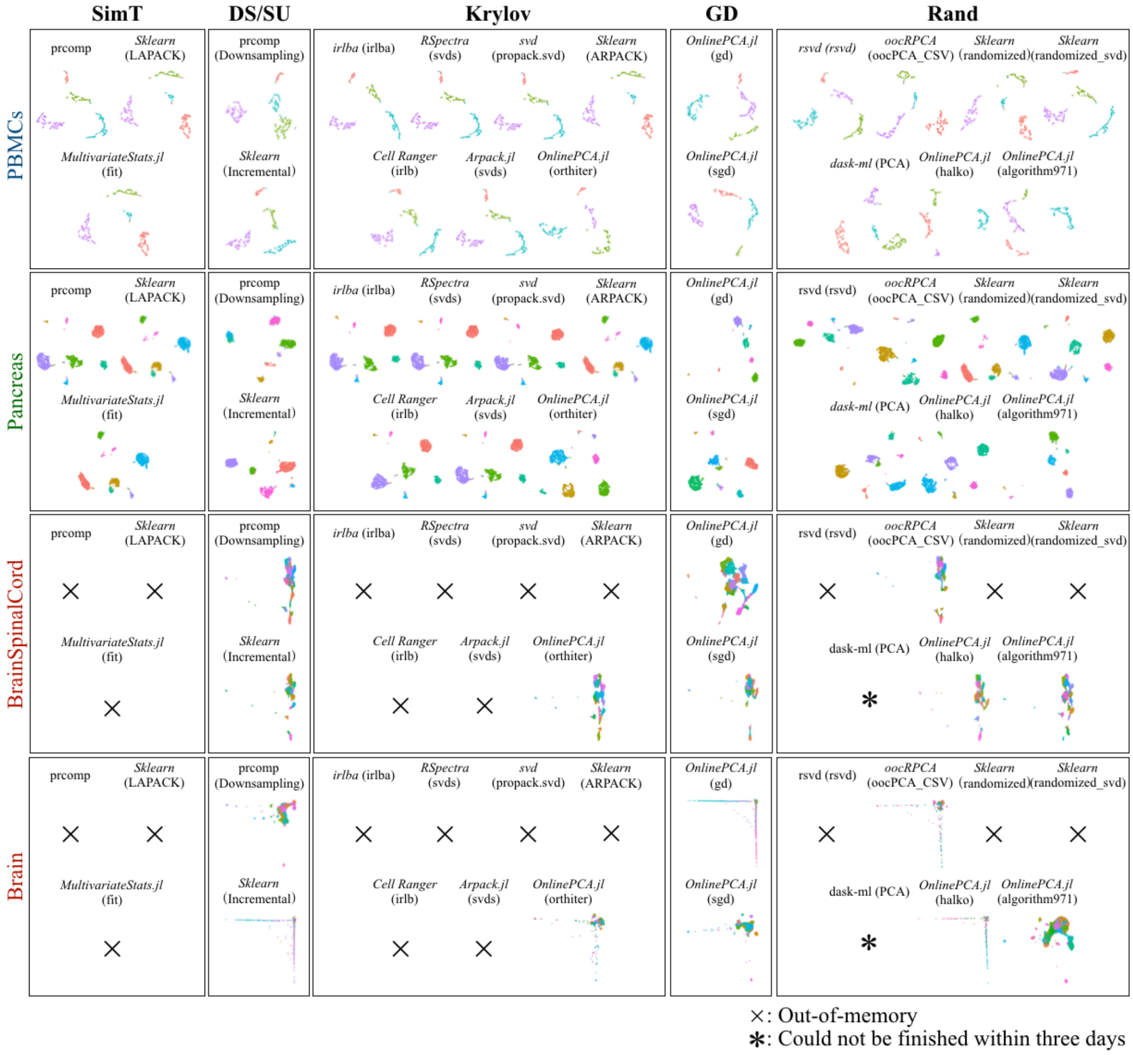


**Supplementary Figure 6-7 | Results of Louvain clustering** The cluster label is visualized with two-dimensional UMAP coordinates.

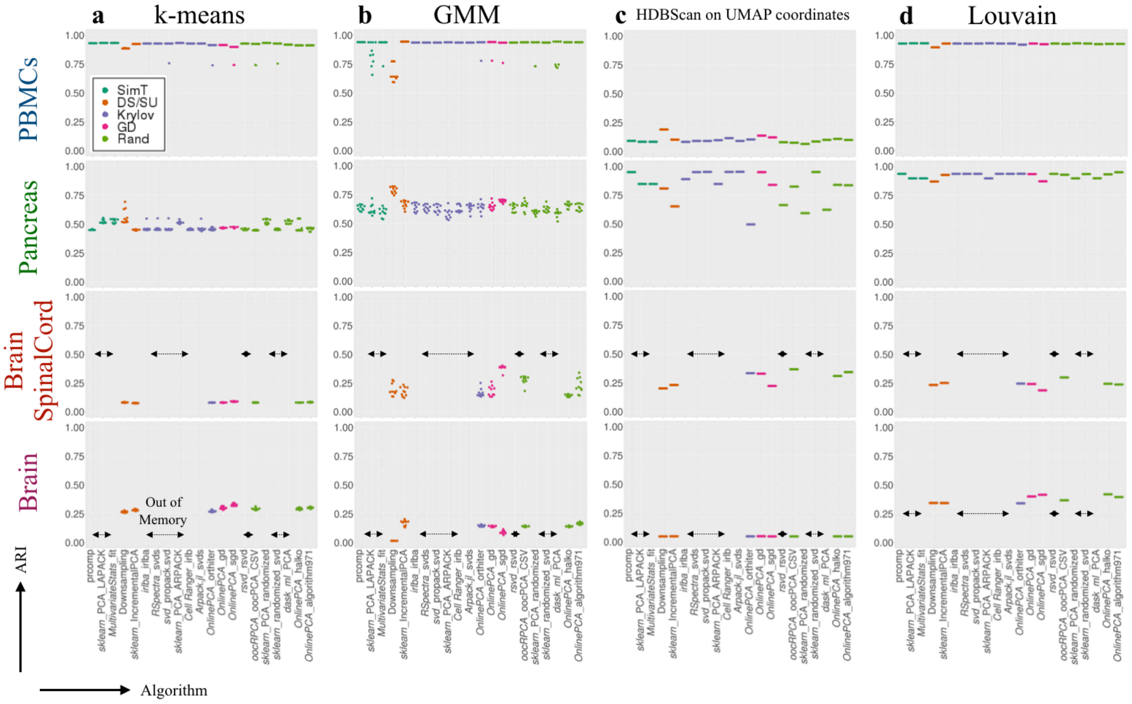


**Supplementary Figure 6-8 | ARI of all clustering results** The algorithms used for benchmarking are listed along the *x*-axes; *y*-axes show associated ARI values.
