## Additional File 16 for "Benchmarking principal component analysis for large-scale single-cell RNA-sequencing"

### Parameter tuning of incremental PCA implementations

Here, we perform parameter tuning of incremental PCA using scikit-learn. In this algorithm, the data matrix is divided into multiple chunks and SVD is performed repeatedly in each chunk to update the SVD result.

Although the smallest chunk size is the number of PCs (**Table 2**), we found that a large chunk size will sometimes improve the accuracy of the PCA results; some subclusters were separated when the chunk size was specified as a large number (**Figure S16-1**, red circle). In addition, the calculation time was also improved when the chunk size was larger (**Figure 8**). Therefore, we set the chunk size to 100 in the analyses of all real datasets. The much larger value might have improved the accuracy and calculation time, though more memory space was consumed.

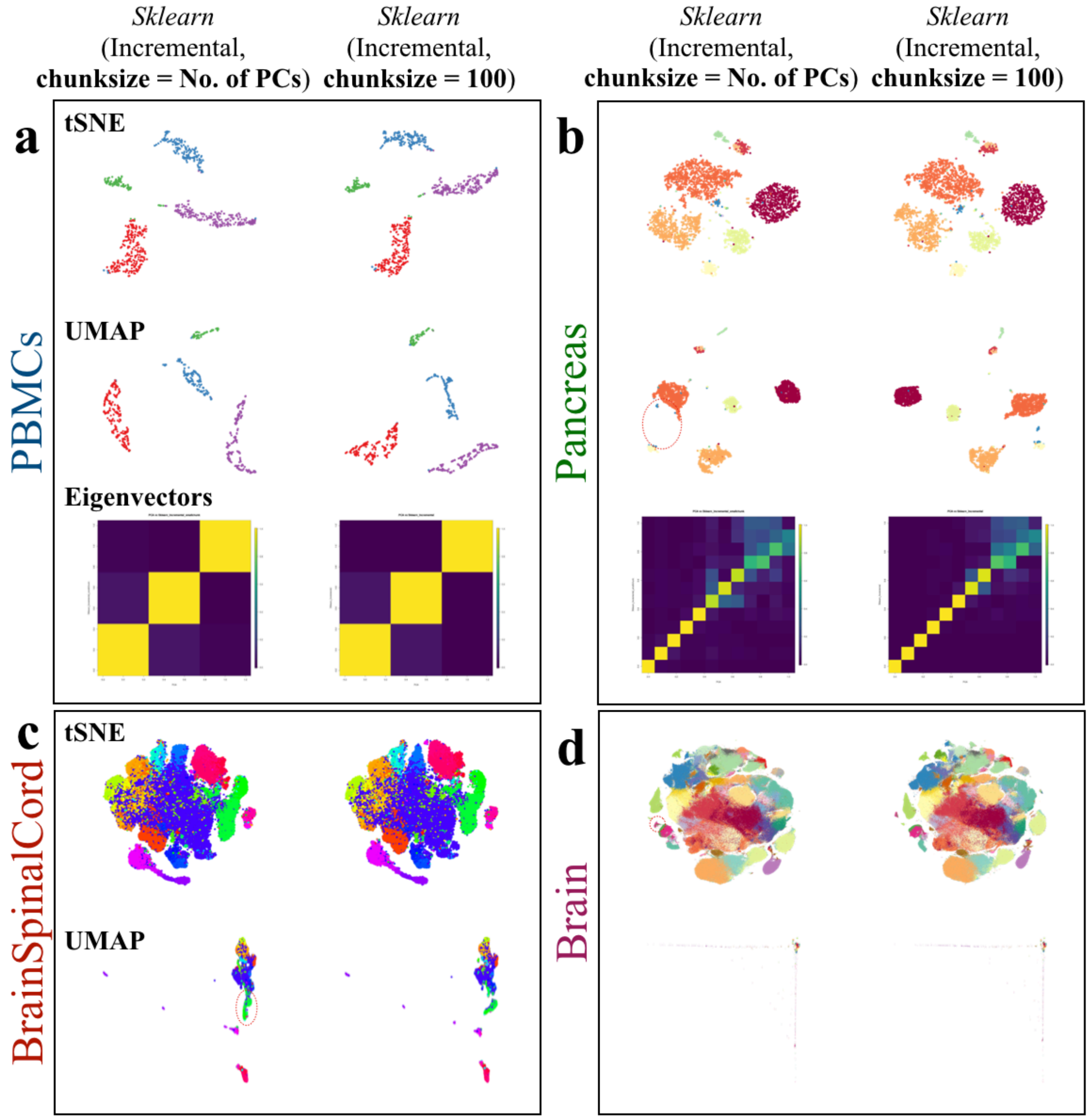


**Supplementary Figure 16-1 | Parameter tuning of incremental PCA (*scikit-learn*)** The chunk size was set at integer values for each number of PCs up to 100.
