## Additional File 17 for "Benchmarking principal component analysis for large-scale single-cell RNA-sequencing"

### Parameter tuning of the orthogonal iteration, gradient descent, and stochastic gradient descent implementations

Here, we evaluate the convergence of three PCA implementations, **orthogonal iteration** (**orthiter**), **gradient descent** (**gd**), and **stochastic gradient descent** (**sgd**), from the *OnlinePCA.jl* package. Both **gd** and **sgd** have a step size parameter, with different parameter values causing different behaviors; too small of a step size value causes slow convergence and too large of a value causes divergence from the optimal solution with each step. In both cases, the result does not converge and is unreliable. Therefore, we performed grid search with multiple step size parameters ($10^{-1}$, $10^{0}$, …, $10^{7}$) and estimated the appropriate values. Because the PCA objective function can be written as the square error between a data matrix and the matrix reconstructed by eigenvectors (reconstruction error), we calculated the error in each of the 5000 rows (**Figure S17-1,2,** and **3**) and confirmed the calculations converged within 10 epochs (1 epoch is equal to the number of all rows in a data matrix). We also calculated the relative change of the error and the proportion of explained variance from the estimated eigenvectors in each of the 5000 rows. Finally, for all the real datasets, we set the step size of **gd** and **sgd** as 1000 and 100, respectively.

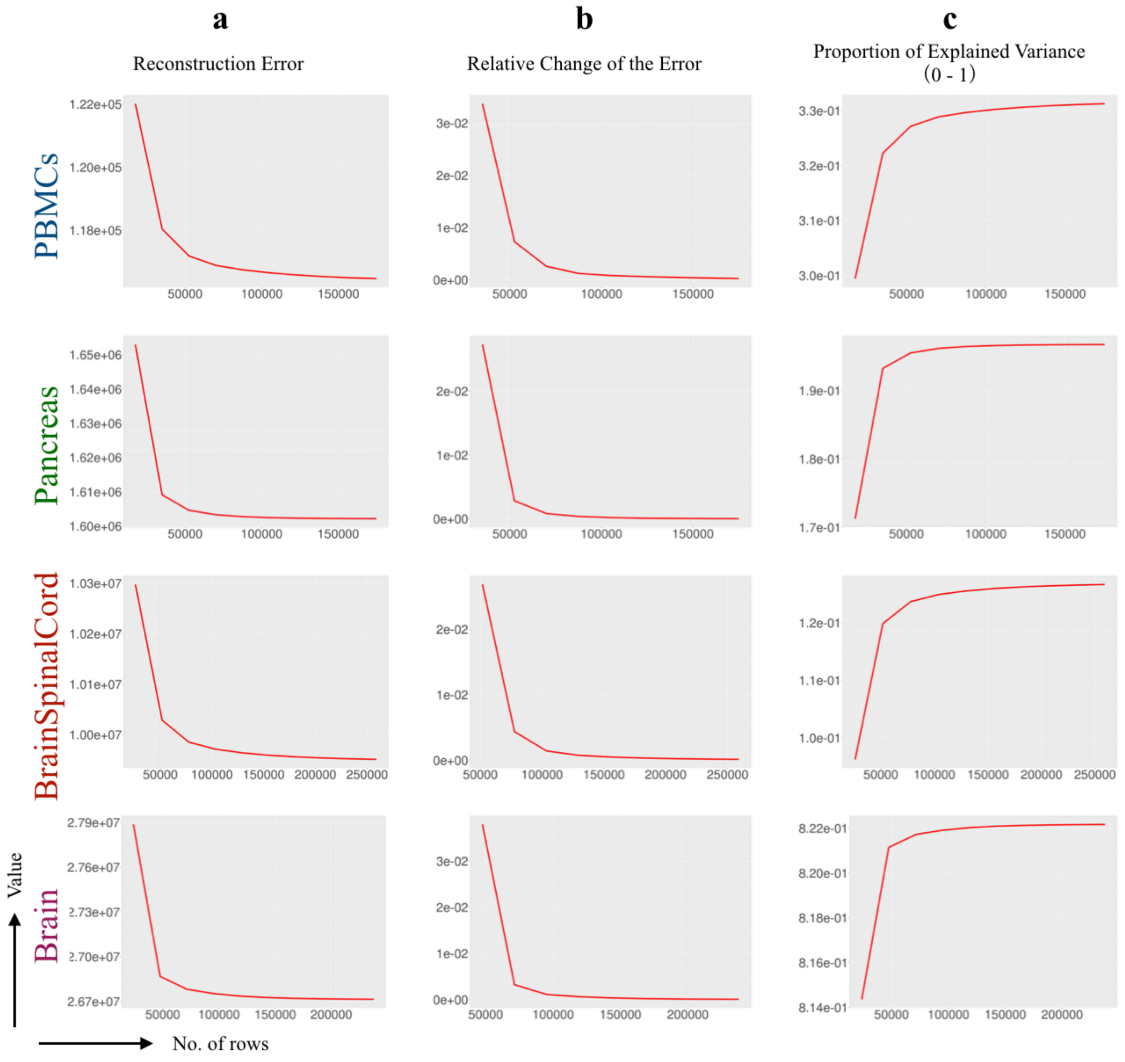


**Supplementary Figure 17-1 | Convergence analysis of orthiter (*OnlinePCA.jl*)** To validate that calculation by **orthiter** (*OnlinePCA.jl*) converges, reconstruction error and relative change of the error were calculated for each of 10 epochs. In all panels, the *x*-axis indicates the number of rows and the y-axis indicates the metrics to evaluate the reconstruction error.

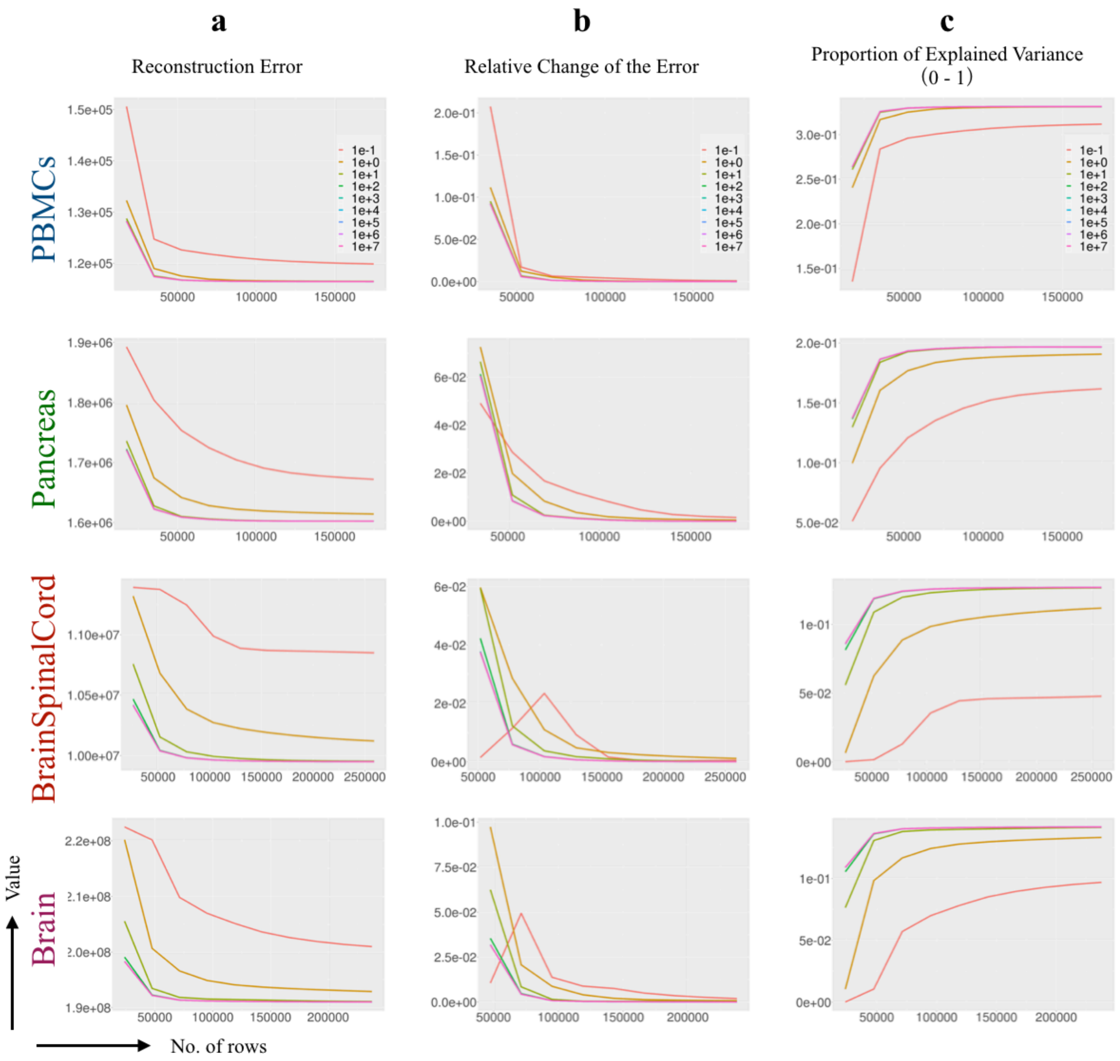


**Supplementary Figure 17-2 | Convergence analysis of gd (*OnlinePCA.jl*)** To validate that calculation by **gd** (*OnlinePCA.jl*) converges, reconstruction error and relative change of the error were calculated for each of 10 epochs. In all panels, the *x*-axis indicates the number of rows the y-axis indicates the metrics to evaluate the reconstruction error.

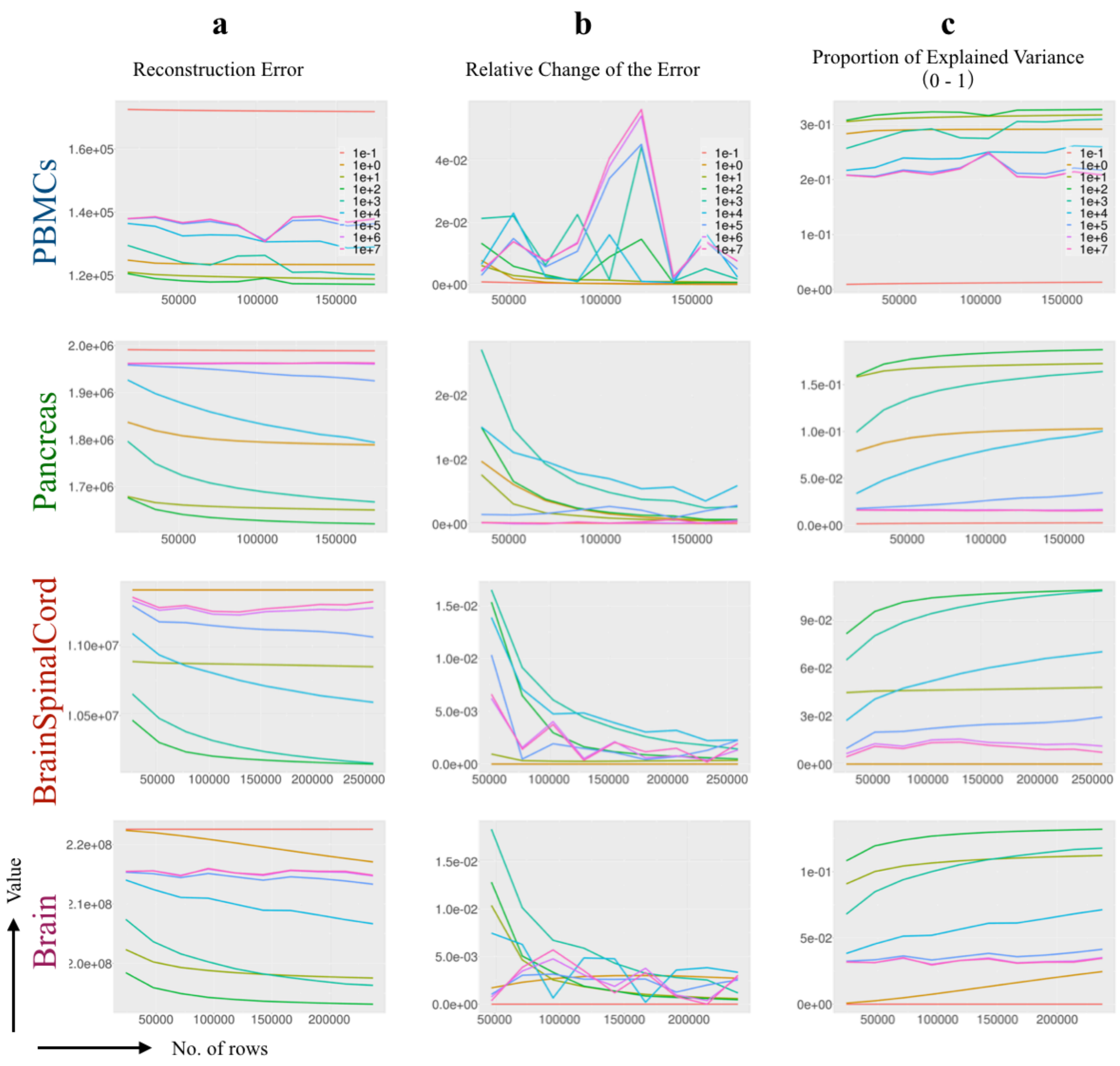


**Supplementary Figure 17-3 | Convergence analysis of sgd (*OnlinePCA.jl*)** To validate that calculation by **sgd** (*OnlinePCA.jl*) converges, reconstruction error and relative change of the error were calculated for each epoch of 5000 rows until a total of 10 epochs was reached. In all panels, the *x*-axis indicates the number of rows the y-axis indicates the metrics to evaluate the reconstruction error.
