## Additional File 18 for "Benchmarking principal component analysis for large-scale single-cell RNA-sequencing"

### Parameter tuning of the randomized SVD algorithms

Here, we perform parameter tuning of the randomized SVD algorithms examined, including Halko's method and algorithm971 [107-109]. These algorithms commonly use the preconditioning process called *power iteration*. This step sharpens the distribution of eigenvalues and enforces a more rapid decay of the singular values ([111] and **Additional file 2**).

We found that the number of iterations (***niter***) is critical to accuracy and at least three or more values are needed (**Figure S18-1** and **2**). Accordingly, for all randomized SVD implementations mentioned in the main manuscript, a value of 3 was used for the *niter* parameter.

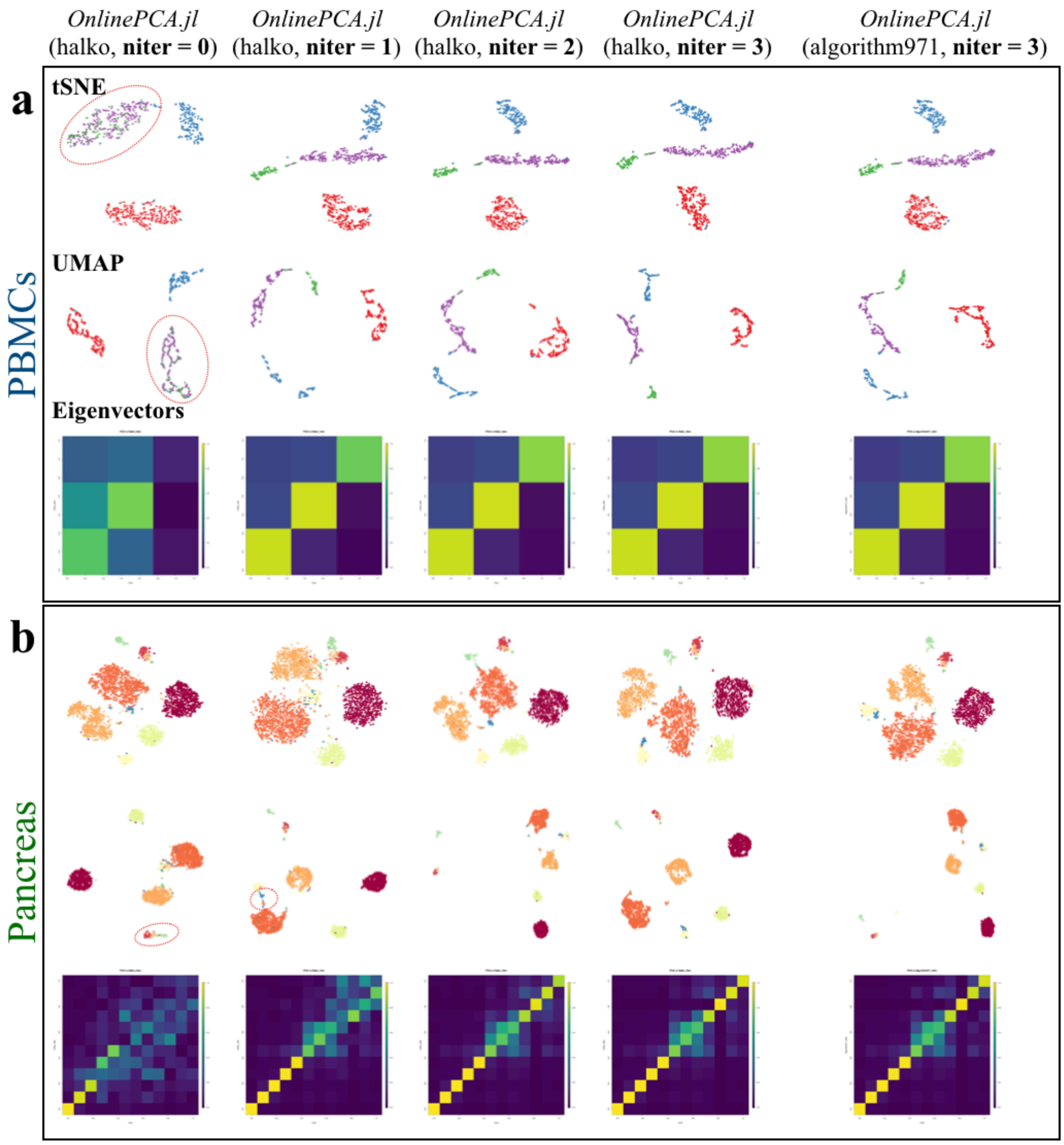


**Supplementary Figure 18-1 | Parameter tuning of Halko's method and algorithm971 (OnlinePCA.jl) (PBMCs and Pancreas datasets)** The *niter* parameter was set at integer values from 0 to 3; the t-SNE appearance and cross-product plot used in the main manuscript were confirmed.

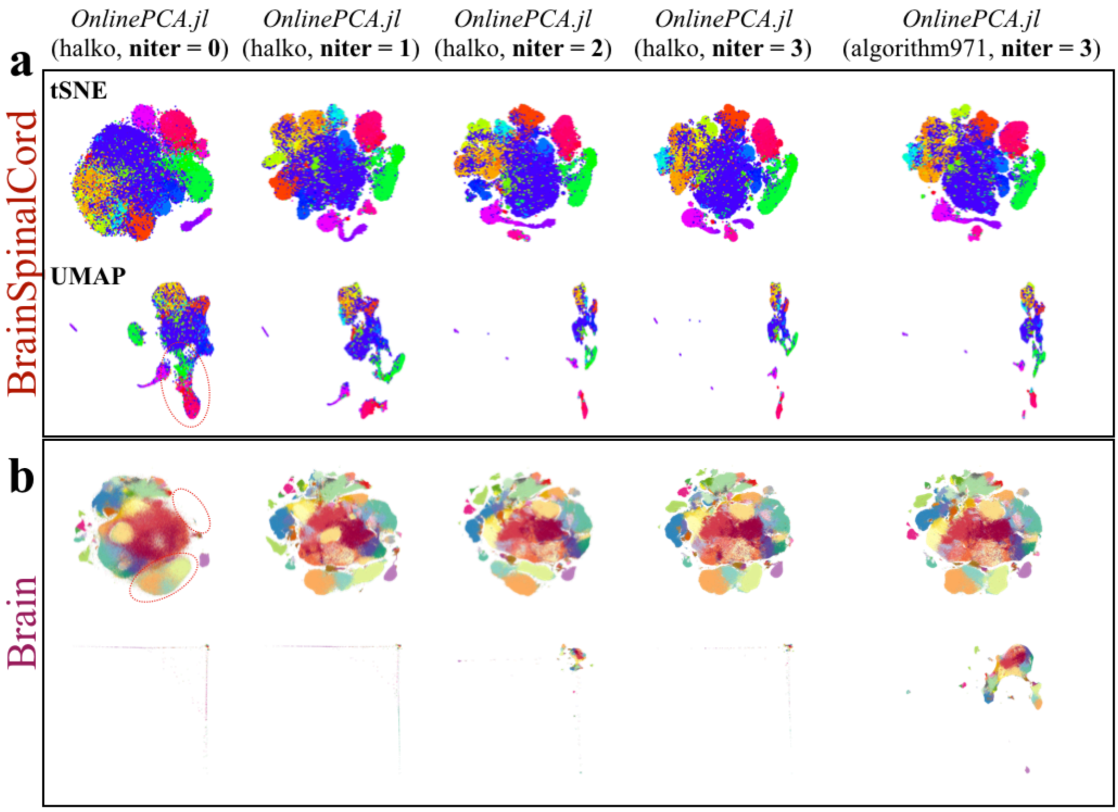


**Supplementary Figure 18-2 | Parameter tuning of Halko's method and algorithm971 (OnlinePCA.jl) (BrainSpinalCord and Brain datasets)**

The *niter* parameter was set at integer values from 0 to 3; the t-SNE appearance and cross-product plot used in the main manuscript were confirmed.
