## Additional File 19 for "Benchmarking principal component analysis for large-scale single-cell RNA-sequencing"

### Data

**• Out-of-core data format**
HDF5, Loom

**• Binding of out-of-core data**
- HDF5 : *rhdf5* (R), *h5py* (Python), *HDF5.jl* (Julia)
- Loom : *LoomExperiment* (R), *loomR* (R), *loompy* (Python)
- Other :
  - R : *bigmemory*, *ff*, *RevoScaleR*, *disk.frame*
  - Python : *Dask*, *Dask-ML*, *H2O*, *Odo*, *Ray*, *Vaex*, *Blaze*, *DistArray*, *bolt*
  - Julia : *ComputeFramework.jl*, *JuliaDB.jl*, *Dagger.jl*, *Elemental.jl*, *DistributedArray.jl*

**• Data compression**
Gzip, Bzip2, Lz4, Zlib, Zstd

**• Sparse data format**
DOK, LIL, COO, CSR, CSC, DIA, BSR, Matrix Market

**• Data type**
Float32/16/8, Int32/16/8, Uint32/16/8

### Algorithm

**• Eigensolver (Dense matrix)**
*Intel MKL*, *ScaLAPACK*

**• Eigensolver (Sparse matrix)**
*ARPACK*, *P_ARPACK*, *Spectra*, *Eigen*, *Armadillo*, *redSVD*, *PROPACK*

**• Binding of eigensolver**
*pbdSLAP* (R), *scalapy* (Python), *ScaLAPACK.jl* (Julia), *rARPACK* (R), *Arpack.jl* (Julia), *RSpectra* (R), *RcppEigen* (R), *RcppArmadillo* (R), *RRedSVD* (R), *pyredsvd* (Python), *svd* (R)

**• Available packages (cf. Figure 13)**

**• Fast / memory-efficient algorithm**
- **DS** : CX / CUR decomposition
- **Krylov** : Randomized Block Lanczos
- **GD** : Learning Rate Scheduler, Variance Reduction, Riemannian Gradient
- **Rand** :
  - Sampling : Uniform distribution, Subsampled Randomized Hadamard Transform (SRHT), Count sketch
  - Pass-efficient / One-pass PCA

### Framework/Environment

**• Distribution/Parallel framework**
- Grid Scheduler (*qsub* (R))
- MPI (*Rmpi*/*pbdMPI* (R), mpi4py (Python), *MPI.jl* (Julia))
- OpenMP (*romp* (R))
- std::thread, TinyThread, Interl TBB (*RcppParallel* (R), *RcppThread* (R))
- Apache *Spark*/*MLib* (Scala, Java, Python, R)
- Apache *Hadoop*/*Mahout* (Java)
- *libskylark* (Python)

**• Distribution/Parallel environment**
- GPU or GPU/CPU-hybrid Cluster Machine
- Accelerator (FPGA, SIMD, MIMD)
- Cloud Computing (Google Cloud, AWS, Azure)

**• C / C++ interface**
*Rcpp* (R), *Cython* (Python), Cxx.jl (Julia)

**• JIT compiler**
*compiler* (R), *Numba* (Python)
