## Additional File 30 for "Benchmarking principal component analysis for large-scale single-cell RNA-sequencing"

### Effect of feature selection on clustering accuracy

Here, we investigated the relationship between the number of genes extracted by highly variable genes (HVGs) and the effect on clustering accuracy. Assuming a situation in which the cluster label and the cluster specific marker genes were not defined yet, we extracted HVGs (http://pklab.med.harvard.edu/scw2014/subpop_tutorial.html) from the Brain dataset (23,771 genes × 1,306,127 cells) with nine different thresholds (top 50, 100, 200, 500, 1000, 2000, 5000, 10,000, and 20,000 genes according to their *p*-values ranked in ascending order, **Figure S20-1**). Because the dataset is huge, and it is hard to load all the elements of the matrix into memory space at once, we originally re-implemented this method in an out-of-core manner using Julia.

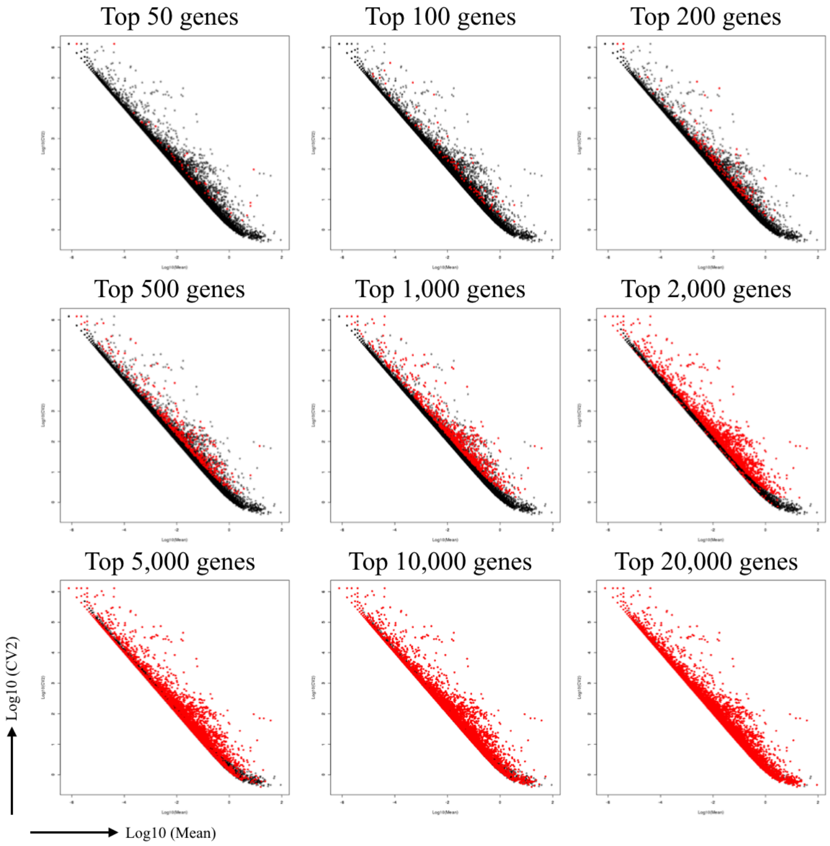


**Supplementary Figure 20-1 | Mean-CV2 plots** The red points are the HVGs and the black points are not HVGs, respectively.

Four clustering methods (*k*-means, GMM, HDBScan, and Louvain) were then performed with each dataset, and clustering accuracy was evaluated using the adjusted Rand index (ARI) values (**Figure S20-2,3,4**, and **5**).

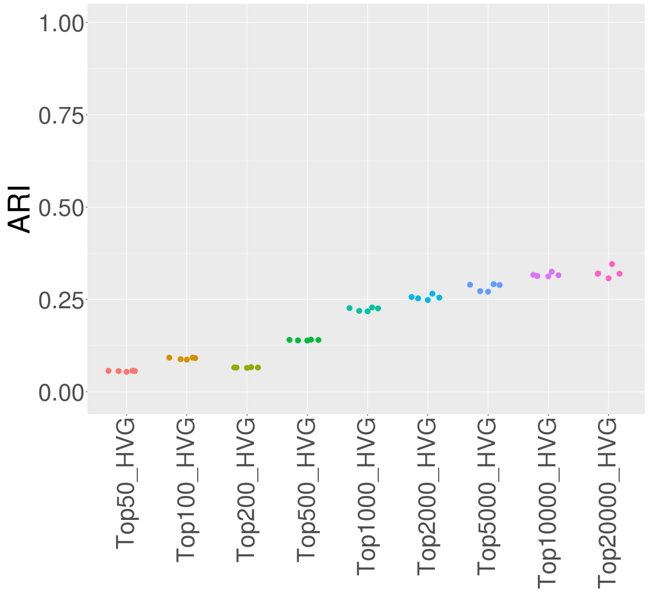


**Supplementary Figure 20-2 | ARI of *k*-means** For each dataset, five trials of *k*-means clustering were performed.

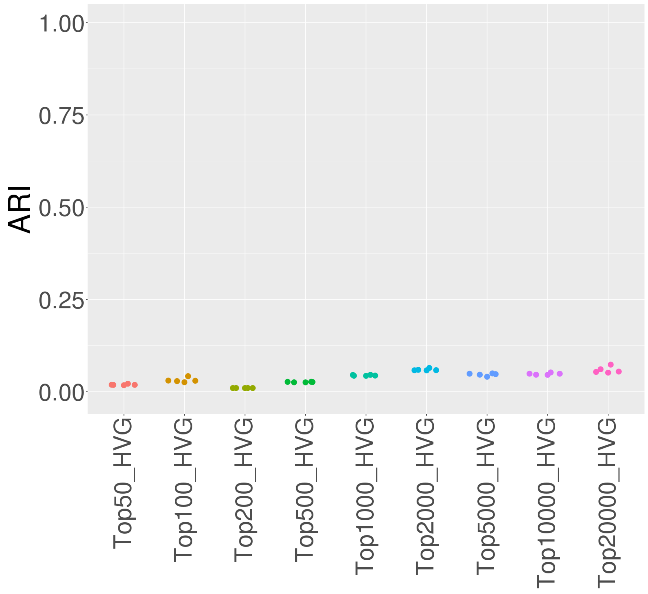


**Supplementary Figure 20-3 | ARI of GMM** For each dataset, five trials of GMM clustering were performed.

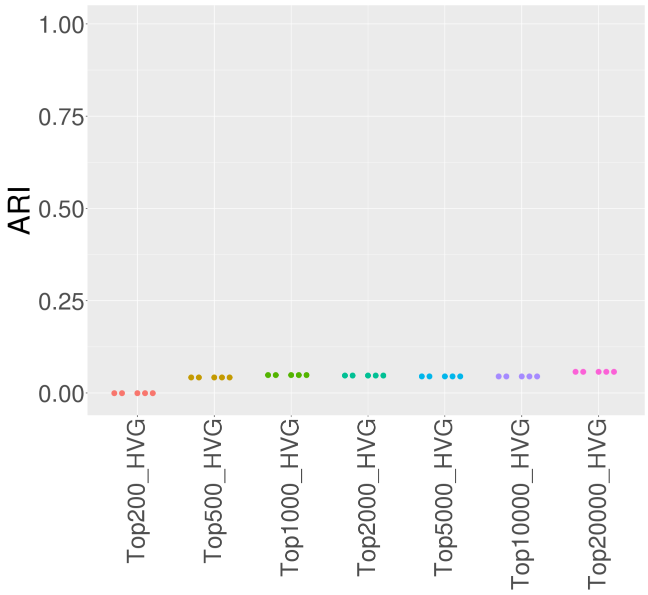


**Supplementary Figure 20-4 | ARI of HDBScan on five-dimensional UMAP coordinates** For each dataset, five trials of HDBScan clustering were performed.

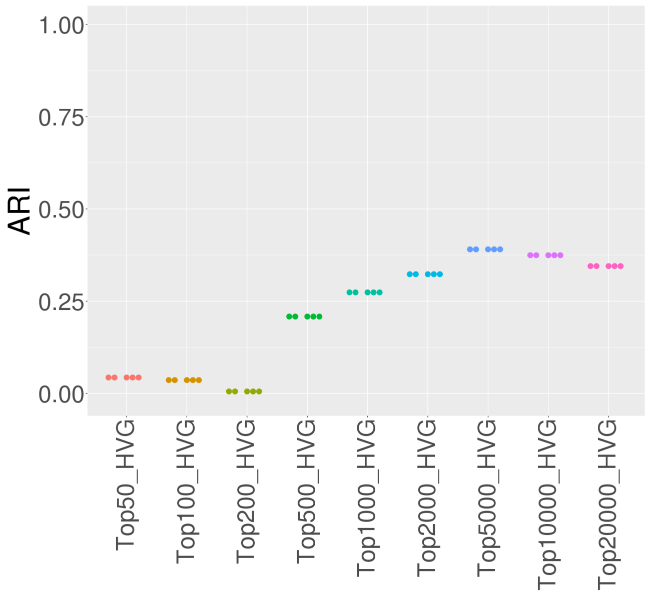


**Supplementary Figure 20-5 | ARI of Louvain** For each dataset, five trials of Louvain clustering were performed.

These results show that ARI values become saturated at around 5000 to

20,000 genes. On the other hand, file size increases with the number of genes (**Figure S20-6**), but such large datasets cannot be loaded as a matrix or data frame in R; in this language, the product of the number of rows and columns cannot exceed 2,147,483,647 (cf. Machine$integer.max, https://stackoverflow.com/questions/14589354/struggling-with-integers-maximum-integer-size). In the case of the Brain dataset, therefore, only 1644 genes (= 2,147,483,647 / 1,306,127) can be imported into R. The results above show that such a small number of genes is insufficient to reach large ARI values in the case of the Brain dataset. Although Python and Julia seem not to define such a limitation, computer specifications impose their own limitations. Indeed, even our large memory machine (512 GB) was unable to load the Brain dataset as a matrix object in either language.

Given the above issues, a scalable PCA implementation is obviously necessary, and as mentioned in the main manuscript, actual data analysis involves trial-and-error cycles (e.g., feature selection or data deletion). Thus, faster and lighter PCA implementations are of value.

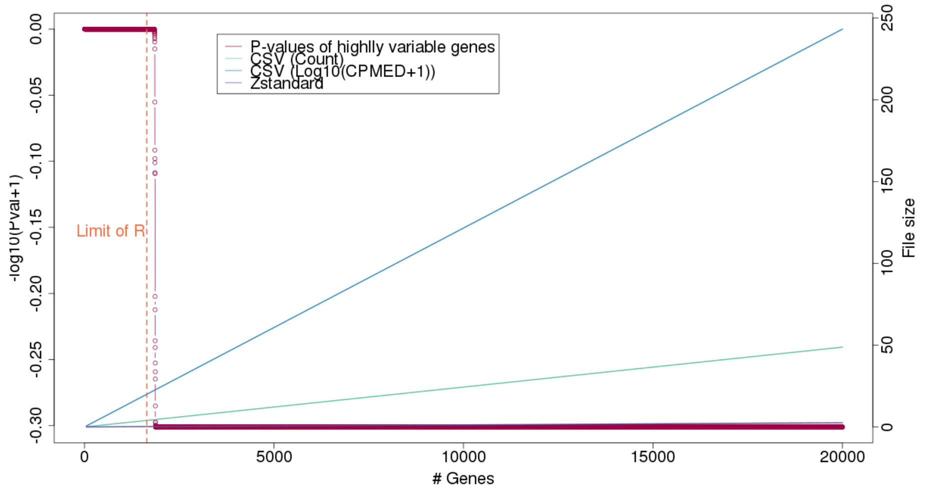


**Supplementary Figure 20-6 | Relationship between the number of genes, *p*-values of HVGs, and the file size of certain file formats** The *x*-axis indicates the number of genes, while the left and right *y*-axes respectively show the logarithms of *p*-values of HVGs and the file size of each dataset.
