## Additional File 21 for "Benchmarking principal component analysis for large-scale single-cell RNA-sequencing"

Comparison of normalizing size factors


### Comparison of normalizing size factors

```
library(SingleCellExperiment)
```

```
## Loading required package: SummarizedExperiment
```

```
## Loading required package: GenomicRanges
```

```
## Loading required package: stats4
```

```
## Loading required package: BiocGenerics
```

```
## Loading required package: parallel
```

```
## 
## Attaching package: 'BiocGenerics'
```

```
## The following objects are masked from 'package:parallel':
## 
##     clusterApply, clusterApplyLB, clusterCall, clusterEvalQ,
##     clusterExport, clusterMap, parApply, parCapply, parLapply,
##     parLapplyLB, parRapply, parSapply, parSapplyLB
```

```
## The following objects are masked from 'package:stats':
## 
##     IQR, mad, sd, var, xtabs
```

```
## The following objects are masked from 'package:base':
## 
##     anyDuplicated, append, as.data.frame, basename, cbind,
##     colnames, dirname, do.call, duplicated, eval, evalq, Filter,
##     Find, get, grep, grepl, intersect, is.unsorted, lapply, Map,
##     mapply, match, mget, order, paste, pmax, pmax.int, pmin,
##     pmin.int, Position, rank, rbind, Reduce, rownames, sapply,
##     setdiff, sort, table, tapply, union, unique, unsplit, which,
##     which.max, which.min
```

```
## Loading required package: S4Vectors
```

```
## 
## Attaching package: 'S4Vectors'
```

```
## The following object is masked from 'package:base':
## 
##     expand.grid
```

```
## Loading required package: IRanges
```

```
## Loading required package: GenomeInfoDb
```

```
## Loading required package: Biobase
```

```
## Welcome to Bioconductor
## 
##     Vignettes contain introductory material; view with
##     'browseVignettes()'. To cite Bioconductor, see
##     'citation("Biobase")', and for packages 'citation("pkgname")'.
```

```
## Loading required package: DelayedArray
```

```
## Loading required package: matrixStats
```

```
## 
## Attaching package: 'matrixStats'
```

```
## The following objects are masked from 'package:Biobase':
## 
##     anyMissing, rowMedians
```

```
## Loading required package: BiocParallel
```

```
## 
## Attaching package: 'DelayedArray'
```

```
## The following objects are masked from 'package:matrixStats':
## 
##     colMaxs, colMins, colRanges, rowMaxs, rowMins, rowRanges
```

```
## The following objects are masked from 'package:base':
## 
##     aperm, apply, rowsum
```

```
sce <- readRDS("real/zheng_2017_monocytes/data/01_sce_all_genes_all_cells.rds")

# raw counts
Y_raw <- as.matrix(assay(sce, "counts"))
libsize <- colSums(Y_raw)

# Count per million (cpm)
Y_cpm <- 1e6 * t(t(Y_raw) / libsize)

# Count per 10k (cp10k)
Y_cp10k <- 1e4 * t(t(Y_raw) / libsize)

# Count per median (cpmed)
Y_cpmed <- median(libsize) * t(t(Y_raw) / libsize)
```

```
library(ggplot2)
library(dplyr)
```

```
## 
## Attaching package: 'dplyr'
```

```
## The following object is masked from 'package:matrixStats':
## 
##     count
```

```
## The following object is masked from 'package:Biobase':
## 
##     combine
```

```
## The following objects are masked from 'package:GenomicRanges':
## 
##     intersect, setdiff, union
```

```
## The following object is masked from 'package:GenomeInfoDb':
## 
##     intersect
```

```
## The following objects are masked from 'package:IRanges':
## 
##     collapse, desc, intersect, setdiff, slice, union
```

```
## The following objects are masked from 'package:S4Vectors':
## 
##     first, intersect, rename, setdiff, setequal, union
```

```
## The following objects are masked from 'package:BiocGenerics':
## 
##     combine, intersect, setdiff, union
```

```
## The following objects are masked from 'package:stats':
## 
##     filter, lag
```

```
## The following objects are masked from 'package:base':
## 
##     intersect, setdiff, setequal, union
```

```
library(tidyr)
```

```
## 
## Attaching package: 'tidyr'
```

```
## The following object is masked from 'package:S4Vectors':
## 
##     expand
```

```
library(viridis)
```

```
## Loading required package: viridisLite
```

```
theme_set(theme_bw())
#g <- match(20, apply(Y_raw, 1, max))
#tibble(umi = Y_raw[g,]) %>% ggplot(aes(x = umi)) + geom_bar()
```

```
Y_raw_l2 <- log2(Y_raw+1)
Y_cpm_l2 <- log2(Y_cpm+1)
Y_cp10k_l2 <- log2(Y_cp10k+1)
Y_cpmed_l2 <- log2(Y_cpmed+1)
plot_histgrams <- function (g) {
  tibble(
    raw = Y_raw_l2[g,],
    cpm = Y_cpm_l2[g,],
    cp10k = Y_cp10k_l2[g,],
    cpmed = Y_cpmed_l2[g,],
  ) %>%
  gather(key = "norm", val = "value") %>%
  mutate(norm = factor(norm, levels = c("raw", "cpm", "cp10k", "cpmed"))) %>%
  ggplot(aes(x = value)) +
    geom_histogram(bins = 80) +
    facet_wrap(~norm) +
    ggtitle(rownames(Y_raw)[g])
}
```

```
gg <- match(c(10, 15, 20, 25, 30), apply(Y_raw, 1, max))
plot_histgrams(gg[1])
```

```
plot_histgrams(gg[2])
```

```
plot_histgrams(gg[3])
```

```
plot_histgrams(gg[4])
```

```
plot_histgrams(gg[5])
```

```
pc_raw <- prcomp(t(Y_raw_l2), rank = 10)
pc_cpm <- prcomp(t(Y_cpm_l2), rank = 10)
pc_cp10k <- prcomp(t(Y_cp10k_l2), rank = 10)
pc_cpmed <- prcomp(t(Y_cpmed_l2), rank = 10)
```

```
zero_fraction <- colMeans(Y_raw == 0)
log_total_umi <- log10(colSums(Y_raw))
tibble(
  raw = pc_raw$x[,1],
  cpm = pc_cpm$x[,1],
  cp10k = pc_cp10k$x[,1],
  cpmed = pc_cpmed$x[,1]
) %>%
  gather(key = "norm", value = "pc1") %>%
  mutate(
    norm = factor(norm, levels = c("raw", "cpm", "cp10k", "cpmed")),
    zero_fraction = rep(zero_fraction, 4),
    log_total_umi = rep(log_total_umi, 4)) %>%
  ggplot(aes(x = zero_fraction, y = pc1, color = log_total_umi)) +
    geom_point(size = 0.2) +
    scale_color_viridis() +
    facet_wrap(~norm, scales = "free_y")
```
